# Reconstructing the ancestral vertebrate brain using a lamprey neural cell type atlas

**DOI:** 10.1101/2022.02.28.482278

**Authors:** Francesco Lamanna, Francisca Hervas-Sotomayor, A. Phillip Oel, David Jandzik, Daniel Sobrido-Cameán, Megan L. Martik, Stephen A. Green, Thoomke Brüning, Katharina Mößinger, Julia Schmidt, Celine Schneider, Mari Sepp, Florent Murat, Jeramiah J. Smith, Marianne E. Bronner, María Celina Rodicio, Antón Barreiro-Iglesias, Daniel M. Medeiros, Detlev Arendt, Henrik Kaessmann

**Affiliations:** Center for Molecular Biology of Heidelberg University (ZMBH), DKFZ-ZMBH Alliance, D-69120 Heidelberg, Germany; Developmental Biology Unit, European Molecular Biology Laboratory, D-69012 Heidelberg, Germany; Department of Ecology and Evolutionary Biology, University of Colorado, Boulder, Colorado 80309, USA; Department of Zoology, Comenius University, Bratislava, Slovakia; Department of Functional Biology, CIBUS, Faculty of Biology, Universidade de Santiago de Compostela, 15782 Santiago de Compostela, Spain; Division of Biology and Biological Engineering, California Institute of Technology, Pasadena, CA, USA; Department of Molecular and Cell Biology, University of California, Berkeley, CA, USA; INRAE, LPGP, 35000 Rennes, France; Department of Biology, University of Kentucky, Lexington, Kentucky

## Abstract

The vertebrate brain emerged more than ∼500 million years ago in common evolutionary ancestors. To systematically trace its cellular and molecular origins, we established a spatially resolved cell type atlas of the entire brain of the sea lamprey – a jawless species whose phylogenetic position affords the reconstruction of ancestral vertebrate traits – based on extensive single-cell RNA-seq and *in situ* sequencing data. Comparisons of this atlas to neural data from the mouse and other jawed vertebrates unveiled various shared features that enabled the reconstruction of the core cell type composition, tissue structures, and gene expression programs of the ancestral brain. However, our analyses also revealed key tissues and cell types that arose later in evolution. For example, the ancestral vertebrate brain was likely devoid of cerebellar cell types and oligodendrocytes (myelinating cells); our data suggest that the latter emerged from astrocyte-like evolutionary precursors on the jawed vertebrate lineage. Our work illuminates the cellular and molecular architecture of the ancestral vertebrate brain and provides a foundation for exploring its diversification during evolution.

---

The brain of vertebrates is a structurally complex and preeminent organ because of its central functions in the body. Its most fundamental divisions are the forebrain (prosencephalon; traditionally divided into the telencephalon and diencephalon), the midbrain (mesencephalon), and hindbrain (rhombencephalon) (Fig. 1a). This regionalization is shared across all extant jawed vertebrates, and present even in jawless vertebrates (i.e., the extant cyclostomes: lampreys and hagfishes), the sister lineage of jawed vertebrates (gnathostomes)^1^ (Fig. 1a), which have overall less complex brains than jawed vertebrates^2^. While a basic molecular regionalization has been described for the substantially simpler central nervous systems of the closest evolutionary relatives of vertebrates (urochordates and cephalochordates)^3,4^, the anatomical complexity of the four major divisions of the vertebrate brain evolved in common vertebrate ancestors ∼515-645 MYA^5^ (Fig. 1a), likely as part of the cephalic expansion that commenced around the emergence of this animal lineage (“new head” hypothesis)^6^.

**Fig. 1.**
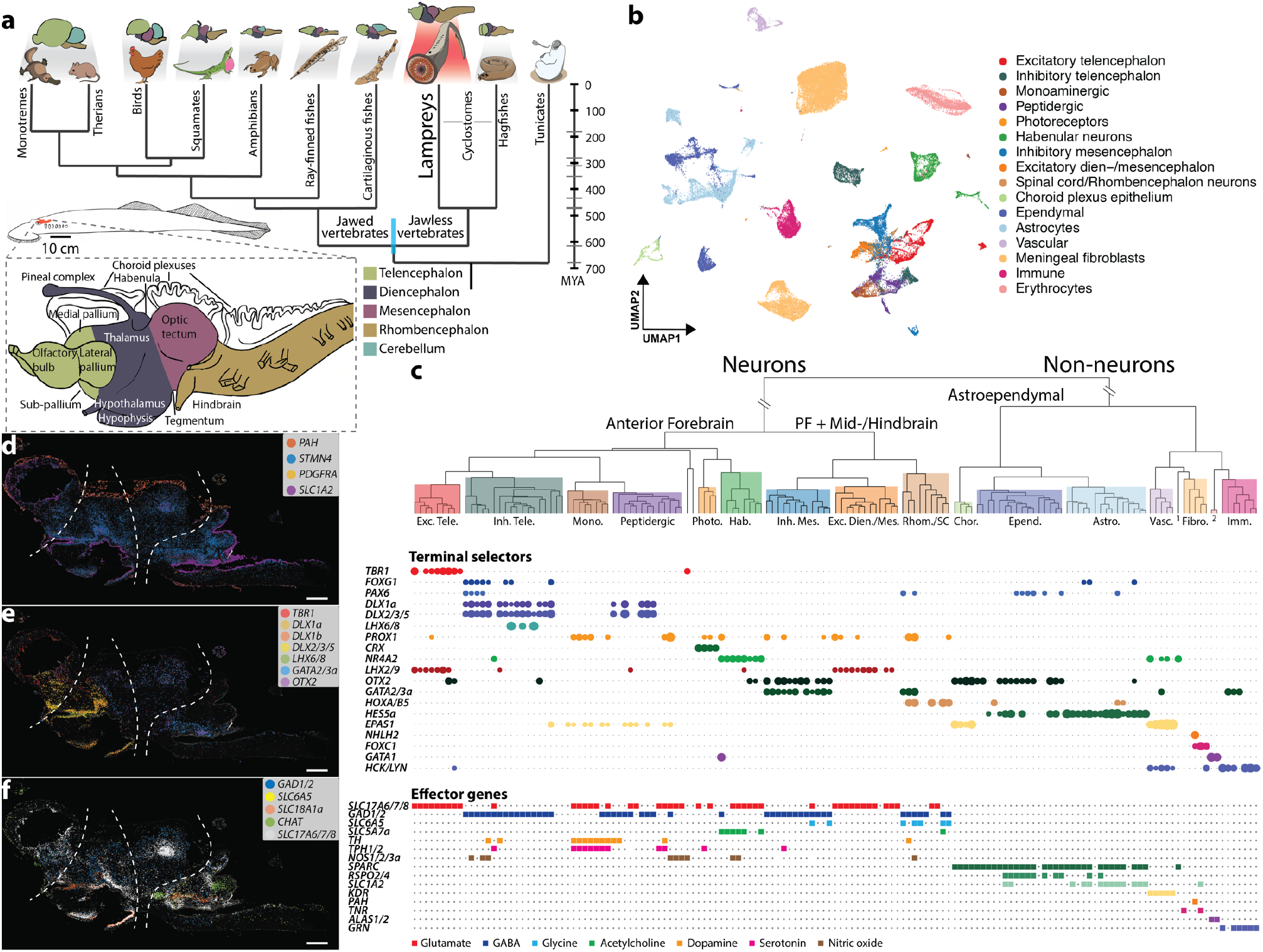
Adult brain atlas overview. **a**, Upper panel: phylogenetic tree displaying the main vertebrate lineages and their approximate brain anatomies; blue bar indicates the estimated confidence interval for cyclostomes/gnathostomes divergence times^5^. Lower panel: schematic of the adult sea lamprey brain showing the different regions dissected for this study. **b**, UMAP of brain cells colored according to their corresponding cell type group. **c**, Dendrogram describing the relationships between the identified cell types. Colored boxes correspond to highlighted cell type groups in b. Upper panel: expression of terminal selector marker genes within each cell type; circle sizes are proportional to the number of cells expressing the gene. Lower panel: binary expression (presence/absence; based on whether a given gene is differentially expressed in the corresponding cell type; see Methods) of effector genes (neurotransmitters for neuronal types). PF, Posterior Forebrain; 1, PNS glia; 2, Erythrocytes. **d, e, f**, Sagittal sections (same orientation as in panel a) of the adult brain showing ISS maps of genes marking: neurons (*STMN4*), astroependyma (*SLC1A2*), and meningeal fibroblasts (*PAH, PDGFRA*) (d); anterior forebrain vs. posterior forebrain and midbrain neuronal factors (e); and neurotransmitter genes (f). Dashed lines separate the main four brain regions illustrated in panel a. See Extended Data Fig. 12 for ISS section schemes; scale bars, 500µm. N.B.: lamprey gene symbols throughout this study are based on corresponding mouse ortholog names.

Previous anatomical and molecular studies of the vertebrate brain have yielded intriguing insights and hypotheses pertaining to its structural and functional evolution^7,8^. However, its ancestral cellular composition and underlying gene expression programs, as well as its subsequent diversification, have not been systematically explored.

To fill this critical gap, we generated a comprehensive cell type atlas of the adult and larval (ammocoete) brain of the sea lamprey (*Petromyzon marinus*), based on extensive transcriptomic and spatial expression data at single-cell resolution (https://lampreybrain.kaessmannlab.org/). Integrated comparative analyses of this atlas unveiled details of the cell type repertoire and molecular architecture of the ancestral vertebrate brain, but also revealed distinct cell types, gene expression programs, and tissue structures that emerged during the evolution of the brain in jawed and jawless vertebrates, respectively.

## Cellular and molecular organization of the lamprey brain

We generated scRNA-seq data (21 libraries in total) for whole adult and ammocoete brains, and separately for their four major anatomical regions (telencephalon, diencephalon, mesencephalon, rhombencephalon), to facilitate cell type assignments (Fig. 1a; Supplementary Tables 1, 2). To ensure optimal scRNA-seq read mapping, we substantially refined and extended previous annotations of the lamprey germline genome^9^ (Extended Data Fig. 1; Supplementary Data 1) based on 63 deeply sequenced RNA-seq libraries covering six major organs, including different brain regions (Supplementary Table 1; Methods). After quality control and data filtering (Methods), we obtained transcriptomes for a total of 159,381 high-quality cells (adult: 72,810; ammocoete: 86,571). Using a detailed clustering approach and iterative marker gene-based annotation procedure (Methods), we identified 151 (95 neuronal) distinct cell types in the adult and 120 (92 neuronal) in the larval datasets, respectively (Supplementary Table 3; see online atlas). To spatially localize cell types across the brain, we generated *in situ* sequencing (ISS)^10^ data for 93 selected marker genes in both lamprey life stages, and single-molecule RNA-FISH (smFISH) images for four genes in the larval stage (Supplementary Table 4; Supplementary Data 2).

Overall, the cell type compositions and gene expression patterns are similar between the adult and ammocoete datasets (Extended Data Figs. 2, 3). However, the adult brain is characterized by larger numbers of expressed genes per cell and greater cell type specificities^11^ of gene expression compared to the ammocoete brain (Extended Data Fig. 4b, c). This result is robust to controls for technical differences between the datasets (Methods; Extended Data Fig. 4a, b, c). Additionally, the ammocoete brain displays the expression of several transcription factor genes (TFs) marking periventricular neurons (e.g., *ZFP704*; Extended Data Fig. 4d-f), which might indicate the presence of not fully differentiated cells in the larval brain.

A cell type tree derived from the adult dataset, which reflects cell type relationships based on gene expression distances, unveils the hierarchical organization of cell types in the lamprey brain (Fig. 1b, c). The primary division is between neuronal and non-neuronal cell types, which are in turn split into astroependymal cells (i.e., neural tube-derived glia) and other cells (i.e., vascular cells, meningeal fibroblasts, hematopoietic cells, and glial cells from the peripheral nervous system, PNS). Our spatial ISS data illustrate that these three major cell type classes occupy very distinct areas of the brain (Fig. 1d).

At a secondary hierarchical level, non-neuronal cells are organized according to their cell class identity (e.g., astrocytes, ependymal cells, erythrocytes, immune cells), in agreement with their molecular phenotype (Fig. 1c). By contrast, the organization of neuronal types primarily reflects their anatomical origin. Thus, a first separation is evident between telencephalic, anterior diencephalic (i.e., hypothalamus and pre-thalamus), pineal, and habenular neurons on one side of the neuronal clade, and posterior diencephalic (i.e., thalamus and pre-tectum), mesencephalic and rhombencephalic/spinal cord neurons on the other. Within each developmental subdivision, neurons are organized according to their neurotransmitter phenotype (Fig. 1c).

The overall hierarchical cell type organization of the lamprey brain is supported by the expression patterns of terminal selectors (i.e., sets of TFs that determine and maintain cell type identity^12,13^) and effector genes (i.e., sets of genes that characterize the molecular phenotype of cells) (Fig. 1c). Inhibitory neurons, for instance, are regulated mainly by *DLX* genes in the anterior forebrain^14^ but by *GATA2/3, OTX2* and *TAL* genes in the posterior forebrain, midbrain, hindbrain and spinal cord^15,16^ (Fig. 1c; Extended Data Fig. 7a). Our ISS data confirm this strict compartmentalization of neuronal regulators (Fig. 1e; Extended Data Fig. 7h, i, l, m). Conversely, neurotransmitter-related genes are expressed across different brain regions (Fig. 1c, f).

The hierarchical relationship of cell types in the lamprey brain is very similar to that observed for a reference mammalian brain atlas (i.e., that of the mouse^17^), which suggests that all vertebrates share a common general cellular and molecular organization of neural tissues that was established during the evolution of the vertebrate stem lineage.

## Vertebrate cell type families

To illuminate the cell type composition and molecular architecture of the ancestral vertebrate brain and to uncover differences between the central nervous systems (CNS) of cyclostomes and gnathostomes, we performed detailed comparative analyses of our adult lamprey atlas with a corresponding atlas established for the mouse^17^. The neuronal and non-neuronal cells of the two atlases were contrasted separately using a dedicated method for homologous cell type detection (self-assembling manifold mapping, SAMap)^18^ and a correlation-based analysis of gene expression that also considers paralogous genes and was adapted from a previous approach^19^ (Methods).

The SAMap results show a great degree of correspondence between the two species for groups of cell types belonging to the same class (e.g., vascular cells, astrocytes, excitatory neurons of the telencephalon), as indicated by the UMAP projection of the inter-species manifold (Fig. 2a, b) and the distribution of mapping scores between the two atlases (Fig. 2e, f; Extended Data Fig. 9; Supplementary Table 5). This high-level similarity is confirmed by cross-species dendrograms based on the correlation approach applied to orthologous TF genes (Fig. 2c, d; Extended Data Fig. 10). These observations suggest that many of the corresponding cell classes share evolutionarily related gene expression programs (Extended Data Fig. 11a). We propose that the matching groups of cell types uncovered in these analyses might constitute homologous cell type ‘families’^20^ that were already present in the brain of the last common ancestor of jawless and jawed vertebrates more than ∼515-645 MYA^5^.

**Fig. 2.**
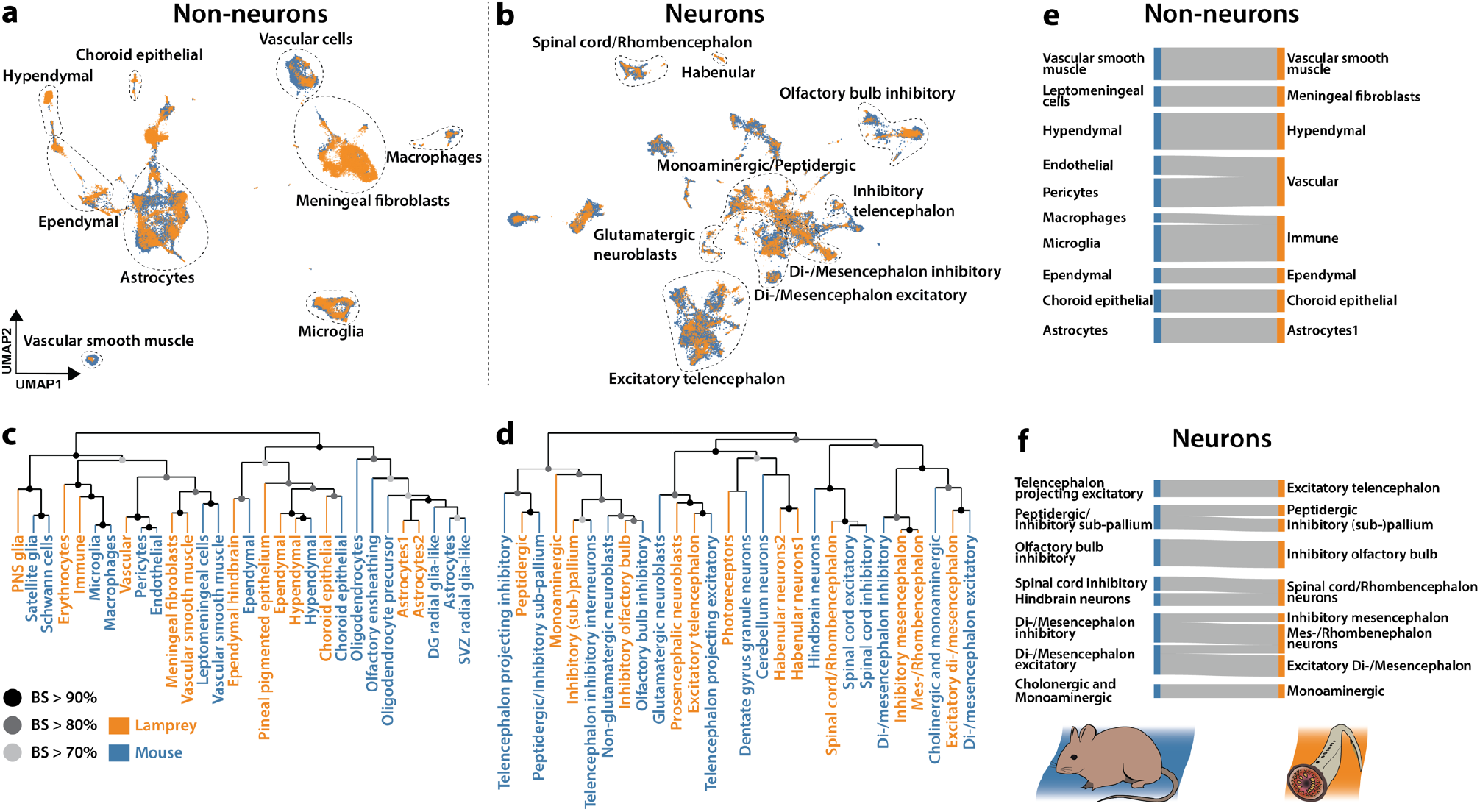
Comparisons between lamprey and mouse brain atlases. **a, b**, SAMap results displaying UMAP projections of non-neuronal (a) and neuronal (b) cells from both species. Erythrocytes and oligodendrocytes were removed from the lamprey and mouse datasets respectively. **c, d**, Dendrograms reporting gene expression distance (Pearson’s *r*) of TF genes of non-neuronal (c) and neuronal (d) cell type groups from the two species. BS, bootstrap support (n = 1,000). **e, f**, Sankey diagrams relating non-neuronal (e) and neuronal (f) cell type groups between the two species based on SAMap mapping scores (min = 0.1; max = 0.65); link width proportional to mapping score.

## Blood, vascular, and PNS cells

The hematopoietic cells found in the lamprey brain can be classified into erythrocytes, characterized by the massive expression of hemoglobin and heme-related genes (e.g., *ALAS1/2*), and immune cells, which are mainly composed of microglia/macrophages and lymphocytes (Extended Data Fig. 5a). The microglia/macrophage cell types are highly correlated to mammalian perivascular macrophages and microglia (Fig. 2c, e; Supplementary Table 5), and express genes that are typically related to the non-specific immune response (e.g., *GRN, CSF1R, HCK/LYN*; Extended Data Fig. 5a-c) both outside (macrophages) and inside (microglia) the brain (Extended Data Fig. 5d). We also identified a lymphocytic cell population (type: Lympho2) expressing one of the two known cyclostome-specific variable lymphocyte receptor genes (*VLRA*; Extended Data Fig. 5a), which is part of a distinct adaptive immune system that emerged on the cyclostome lineage in parallel to that of gnathostomes^21^.

We identified several vascular cell types, corresponding to endothelial cells/pericytes, which express typical vascular markers (e.g., *EPAS1, KDR*; Extended Data Fig. 5a-c) and are principally localized at the innermost meningeal layer (Extended Data Fig. 5d), forming the perineural vascular plexus^22^. The inner and outer leptomeningeal layers are populated by fibroblast-like cells (type: Fibro1) that are likely homologous to the meningeal vascular fibroblasts described in the mouse brain^17,23^ given their high respective homology mapping scores (Fig. 2c, e; Supplementary Table 5) and the expression of key orthologous marker genes (e.g., *PDGFRA, FOXC1, LUM*; Extended Data Fig. 5a-d). A second fibroblast type (Fibro2) occupies the space between the leptomeningeal boundaries (Extended Data Fig. 5d) and is characterized by the expression of genes involved in the metabolism of glucose (*G6PC*)^24^, fatty acids (*FABP3*), cholesterol (*SOAT1/2*), and aromatic aminoacids (*PAH*) (Extended Data Fig. 5a) This cell type might correspond to meningeal round cells, which form a metabolically active tissue typical of lamprey that is not present in the meninges of other vertebrates^22,24^.

PNS glia are represented by a small cluster (n = 53) expressing the TF genes *SOXE2* and *SOXE3* (orthologous to mammalian *Sox10* and *Sox9*, respectively^25^); they co-localize within cranial nerve roots (Extended Data Fig. 5a, e). This group of cells, which most likely corresponds to the peripheral ensheathing glia described by Weil and colleagues^26^, expresses some markers whose mouse orthologs are characteristic of satellite glia (*SOXE2*) and Schwann cells (*EGR2/3/4, PMP22*/*EMP3*) (Extended Data Fig. 5a) However, they lack the expression of key peripheral myelin constituent genes like *MPZ* and *PMP2*, confirming the absence of myelin from the lamprey PNS^27^. The co-localization of this cell type together with meningeal fibroblasts and vascular smooth muscle cells within the cell type tree (Extended Data Fig. 5a) likely reflects their common developmental origin from the neural crest/placodes^17^.

## Astroependymal cells and the origin of myelination

Astroependymal cells (i.e., CNS glia) are divided into two main, developmentally related, cell classes: ependymal cells and astrocytes. Ependymal cells are ciliated, epithelial-like cells that populate the ventricular system of the brain, the circumventricular organs (CVO)^28^, and the choroid plexuses^29^, and are characterized by the expression of the ciliogenesis-related TF *FOXJ1* and the extracellular matrix component *CCN2/3/5* (Extended Data Fig. 6a, e, f, i). We identified two types of specialized secretory ependymal types in the lamprey brain: choroid plexus epithelial cells (*OTX2*^+^), responsible for the production of cerebrospinal fluid (CSF), and hypendymal cells of the sub-commissural organ (SCO), which massively express the main Reissner’s Fibers component SCO-spondin (*SSPO*)^30^ (Extended Data Fig. 6a, b, g, i). Two additional types of specialized ependymal cells are the pigmented pineal epithelial cells, defined by markers that are common to the retina pigment epithelium (e.g., *RPE65, RRH*; Extended Data Fig. 6a), and the *KERA*-expressing ependymal cells of the hindbrain and spinal cord (types: ReEpen1, ReEpen3; Extended Data Fig. 6a, c, d). The large number of detected ependymal cells and cell types in the adult dataset (Extended Data Fig. 3b) likely reflects the large relative sizes of the ventricles and choroid plexuses of the lamprey brain (Extended Data Fig. 6i)^31^.

Lamprey astrocytes are highly comparable to those from mouse in terms of their overall transcriptome (Fig. 2c, e). They share key marker genes that are fundamental for the development and function of astrocytes, such as *SOXE3* (*Sox9*), *HES5*, and *SLC1A2* (Fig. 3a; Extended Data Fig. 6a). Like in other anamniotes (e.g., fishes, amphibians), lamprey astrocytes are mainly localized around the ventricles (Fig.3; Extended Data Fig. 6h), forming the so-called ependymo-radial glia^32^.

**Fig. 3.**
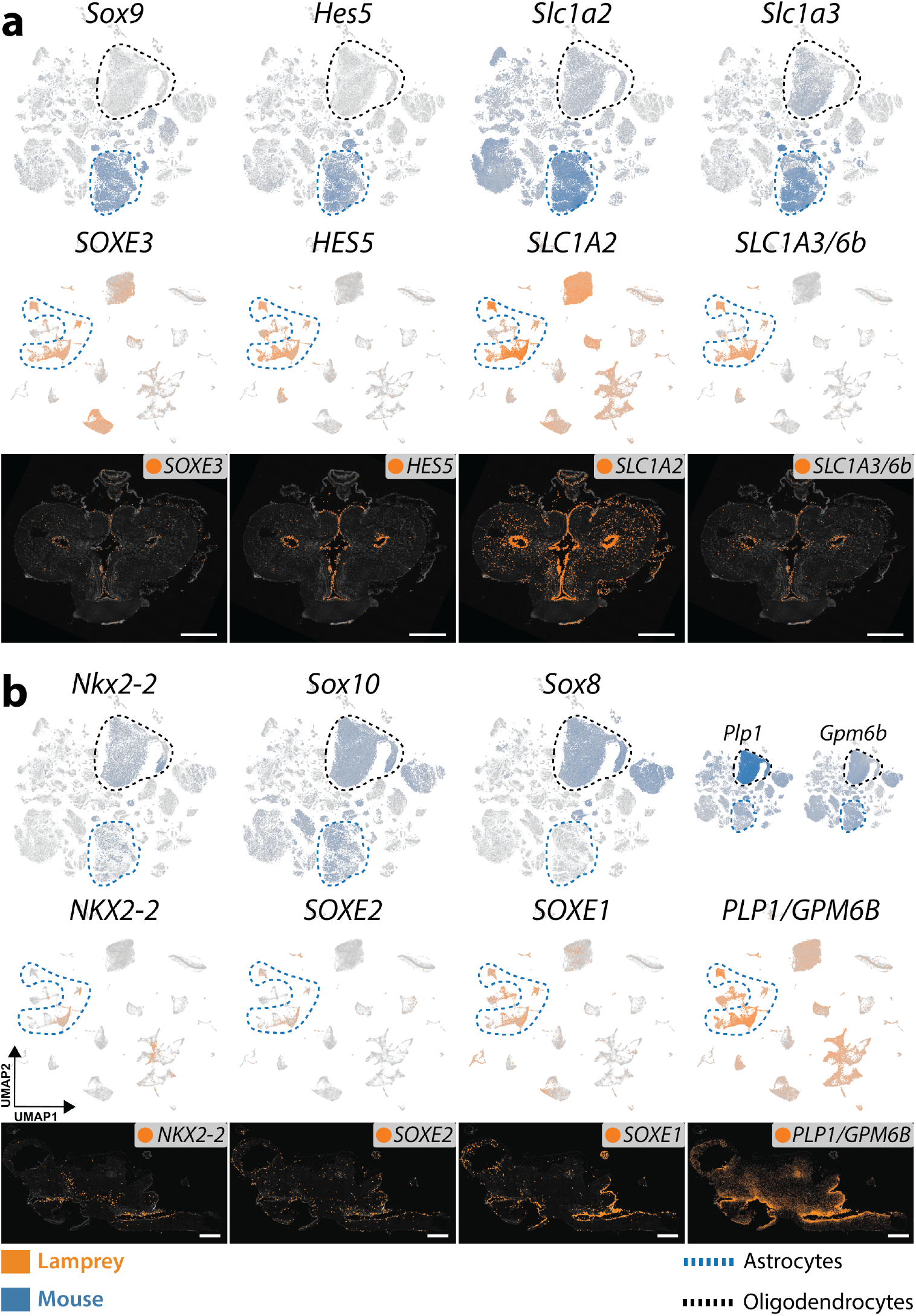
Expression of astrocyte- and oligodendrocyte-specific genes. **a, b**, UMAPs showing the expression of astrocyte- (a) and oligodendrocyte- (b) specific orthologous genes in the mouse (upper panels) and lamprey (middle panels) atlases. Lower panels: ISS maps of the adult lamprey brain for the same genes, showing coronal sections of the telencephalon (a) and sagittal sections of the whole brain (b; same orientation as in Fig. 1a). See Extended Data Fig. 12 for ISS section schemes; scale bars, 500µm.

Like in the PNS, lamprey CNS axons are not myelinated^27^, consistent with the absence of key master regulators of oligodendrocyte identity (*OLIG1, OPALIN*) and myelin specific genes (*MOBP, TSPAN2*) from its genome. Other myelin-related genes are present in the genome, but they are not expressed in glial cells (e.g., *PDGFRA* and *NKX6-1/2* are expressed in meningeal fibroblasts; Extended Data Fig. 5a). Notably, despite the lack of myelination, lamprey astrocytes express several oligodendrocyte-specific genes, such as the TFs *NKX2-2* and *SOXE2* (*Sox10*)^33^ (Fig. 3b; Extended Data Fig. 6a), the proteolipid gene *PLP1/GPM6B* (orthologous to the myelin components *Plp1* and *Gpm6b*) (Fig. 3b; Extended Data Fig. 6a), and the extracellular matrix glycoproteins *TNR* and *HEPACAM* (Extended Data Fig. 6a, j, k). Given the expression of crucial TFs of oligodendrocyte identity and the presence of myelin-related genes within lamprey astrocytes, our findings lend strong support to the hypothesis that oligodendrocytes originated from astrocyte-like glia in gnathostome ancestors^26^.

## Neuronal diversity across brain regions

Finally, we scrutinized neuronal cell types across the different brain regions. Hindbrain and spinal cord neurons are defined by the expression of several *HOX* genes (*HOXA/B3, HOXA/B4, HOXA/B5*; Extended Data Fig. 7a, b). Two types of hindbrain glycinergic cells (types: ReInh5, ReInh6), likely corresponding to inhibitory reticulospinal neurons^34^, are highly correlated to reticular neurons of the medulla in mouse^17^ (Supplementary Table 5) and express related markers (*SLC6A5, SLC32A1b, EBF2/3*; Extended Data Fig. 7a, b, f, g). Cholinergic neurons expressing the TF gene *TBX6/20* show very localized expression within the hindbrain, likely corresponding to afferent nuclei of cranial nerves^35^ (Extended Data Fig. 7c-e). None of the detected midbrain/hindbrain clusters specifically express markers related to Purkinje (e.g., *ALDOC, PCP2, SLC1A6, CAR8*) or granule (e.g., *NEUROD1, CBLN1, GABRA6*) neurons of the cerebellum, nor are these markers expressed in the dorsal isthmic region (Supplementary Data 2). We did also not detect the expression of marker genes in this region that are associated with neurons of inferred ancestral cerebellar nuclei^36^, which were shown to have diversified in the gnathostome lineage through duplications^36^. These observations confirm the absence of proper cerebellar nuclei in the lamprey brain^36,37^. Within the rostral spinal cord we identified two types of GABAergic CSF-contacting (CSF-c) cells^38^ (types: ReInh1, ReInh2); these are ciliated neurons that are homologous to the gnathostome CSF-c neurons of the spinal cord central canal, and express genes coding for channels that respond to changes in CSF pH (*PKD2L1, PKD2L2*), and for proteins that remove toxic oxidative compounds from the CSF (*AMBP*) (Extended Data Fig. 7a, b).

Thalamic, pre-tectal and tectal neurons are divided into excitatory and inhibitory classes (Extended Data Fig. 7a) and express TFs that are typical of homologous anatomical regions in mouse (i.e., thalamus, pre-tectum, and superior colliculus)^17^. In fact, like in the murine brain, glutamatergic neurons are characterized by the expression of *SHOX2, EBF1, EBF2/3*, whereas GABAergic neurons express *GATA2/3*a, *GATA2/3*b, *TAL1, OTX1/2* (Fig. 1e; Fig. 4a; Extended Data Fig. 7a, f-m).

**Fig. 4.**
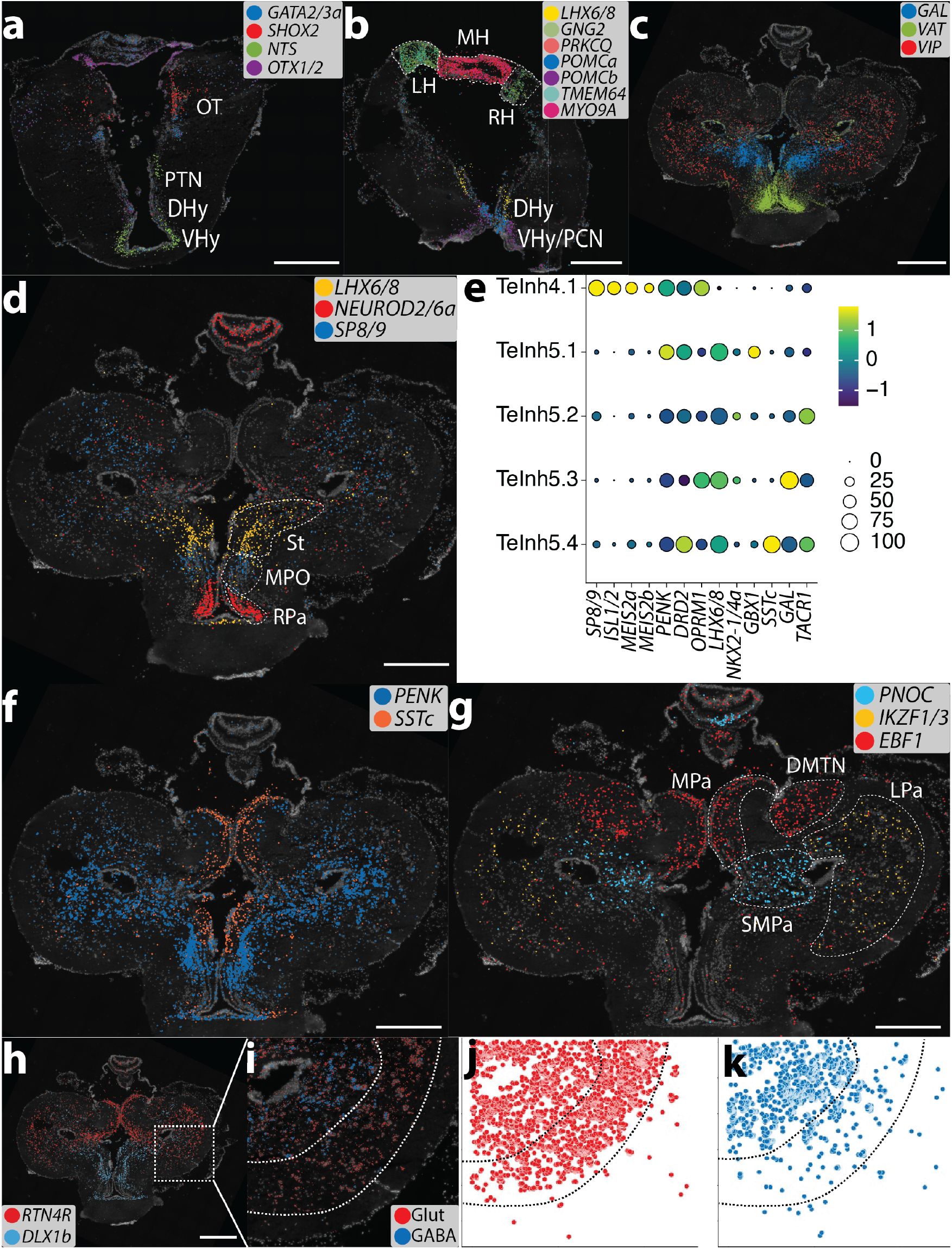
Neuronal spatial maps. **a-d, f-h** ISS maps of selected neuronal marker genes across various adult brain coronal sections. **e**, Dotplot displaying the expression of marker genes for subtypes of LGE- (TeInh4) and MGE- (TeInh5) derived (sub-)pallial inhibitory neurons. **i**, Magnification from the dashed square in h showing the layered organization of the lateral pallium; each neuronal class is highlighted by plotting the expression of multiple specific marker genes: GABA (*GAD1/2, DLX1a, DLX1b, DLX2/3/5*), Glut (*SLC17A/6/7/8, TBR1*). **j, k**, Spatial scatter plot (same coordinates and markers as in i) highlighting the position of GABAergic and glutamatergic neurons within the lateral pallium. DHy, dorsal hypothalamus; DMTN, dorsomedial telencephalic nucleus; LH, left habenula; LPa, lateral pallium; MH, medial habenula; MPa, medial pallium; MPO, medial preoptic nucleus; OT, optic tectum; PCN, postoptic commissure nucleus; PTN, posterior tubercle nucleus; RH, right habenula; RPa, rostral paraventricular area; SMPa, sub-medial pallium; St, striatum; VHy, ventral hypothalamus. See Extended Data Fig. 12 for ISS section schemes; scale bars, 500µm.

Epithalamic neurons (i.e., neurons stemming from the dorsal-most region of the diencephalon) are divided into habenular types and pineal/parapineal photoreceptors, like in the gnathostome brain^39^. All habenular neurons express the same TFs (*NR4A2, ETV1, IRX2/5*; Extended Data Fig. 8a, d), with the medial and lateral nuclei showing very distinct expression patterns for several genes (e.g., *MYO9A, PRKCQ, GNG2, TMEM64*; Fig. 4b; Extended Data Fig. 8a, c). The medial habenula is occupied by glutamatergic, nitrergic, and cholinergic neurons^40,41^ (Fig. 1c; Extended Data Fig. 8a, c), with a cell type expressing neuropeptide Y (*NPY*; Extended Data Fig. 8a, c). The lateral habenulae are molecularly related to each other; they co-express several markers (*GNG2, TMEM64, SLC1A3/6a*; Fig. 4b; Extended Data Fig. 8a, c) and can be distinguished by the differential expression of two neuropeptide genes: proenkephalin (*PENK*; right) and cholecystokinin-like (*CCK*-like; left) (Extended Data Fig. 8e).

The pineal and parapineal organs of the lamprey are characterized by the presence of cells that are both photoreceptive and neuroendocrine^42^. Such a dual nature is confirmed by our data at the molecular level by the expression of the genes *CRX* (necessary for the differentiation of photoreceptors), *GUCA1B* (involved in visual phototransduction), *LHX3/4*, and *ISL1/2* (required for the development of neuroendocrine cells in the mammalian anterior pituitary)^43^ in all detected cell types (Extended Data Fig. 8a, n, r). Within the pineal organ, we identified cone- (type: Photo1) and rod-like (type: Photo2) cells defined by the expression of cone- (*ARR3, PDE6H*) and rod-specific markers (*RHO, PDE6G, GRK1*). Both pineal types can be distinguished from parapineal photoreceptors by the expression of recoverin (*RCVRN*) and genes involved in the biosynthesis of melatonin (*TPH1/2, AANAT*) (Extended Data Fig. 8a, m, p, q; see also online atlas). Cells of the parapineal organ (types: Photo3, Photo4) express the non-visual opsin gene parietopsin and the neuropeptide gene *TAC1* (Extended Data Fig. 8a). Unlike the pineal stalk, characterized by the expression of pineal markers, parapineal ganglion, and tract cells are strongly marked by lateral habenular genes (e.g., *PPP1R14A/B/*C, *GNG2*; Extended Data Fig. 8j, m, n), confirming the long-standing hypothesis of Studnicka (1905)^39^ that these two structures are extensions of the left habenula and that they are histologically distinct from the parapineal organ.

Nearly all monoaminergic neurons, identified by the expression of monoamine transport (*SLC18A1a, SLC18A1b*) and metabolic (*TH, TPH1/2*) genes, form a unique taxon within the cell type tree (Fig. 1c; Extended Data Fig. 8a), which includes serotoninergic and dopaminergic neurons of the hindbrain, midbrain, and hypothalamus. Dopaminergic neurons of the posterior tubercle nucleus (PTN) of the hypothalamus (type: MeDopa1) co-express dopamine- and glutamate-related genes^44^ (Extended Data Fig. 8a, b) and are considered homologs of the dopaminergic neurons of the substantia nigra *pars compacta* (SNc) of amniotes^45^, an important component of basal ganglia. These cells are located next to *NTS*-producing neurons^46^ (a modulator of dopaminergic activity^47^; Fig. 4a) and express the TF *PROX1* (Fig. 1c) that is crucial for the development of dopaminergic PTN cells in zebrafish^48^.

Like in the mouse brain atlas^17^, most hypothalamic peptidergic neurons co-cluster with monoaminergic cells (Fig. 1c; Extended Data Fig. 8a). Neurons of the ventral hypothalamus and postoptic commissure nucleus (PCN) express the neuropeptide genes cholecystokinin (*CCK*)^49^ and pro-opiomelanocortins (*POMCa, POMCb*) (Fig. 4b; Extended Data Fig. 8e, g), as well as the circadian rhythm-related genes *SIX3/6a* and *PER1/2* (also expressed in the pineal complex; see online atlas). Other neuropeptides expressed in the hypothalamus are: galanin (*GAL*)^50^, somatostatins (*SSTa, SSTc*)^51^, neuropeptide Y (*NPY*)^52^, neurotensin (*NTS*), vasotocin (*VAT*), proenkephalin (*PENK*), prepronociceptin (*PNOC*), gonadotropin-releasing hormones (*GNRH1, GNRH2*), prolactin-releasing hormone (*PRLH*), and *FAM237A/B* (Extended Data Fig. 8a, c-g). Additional peptidergic neurons cluster with inhibitory neurons of the pallium/sub-pallium (Extended Data Fig. 8a); these include *GAL*^+^ neurons of the septum and preoptic area (POA) (type: DePep10) and glutamatergic neurons expressing *VAT, GNRH1*, and *NEUROD2/6a* (type: DePep9) located in the rostral paraventricular area (RPa) of POA (Fig. 4c, d; Extended Data Fig. 8a, m).

Inhibitory neurons of the telencephalon are classified into olfactory bulb (OB) and pallium/sub-pallium cell types and are all enriched for typical forebrain GABAergic markers (*GAD1/2, DLX1a, DLX1b, DLX2/3/5*; Extended Data Fig. 7l, m; Extended Data Fig. 8a). OB neurons can be recognized by: i) the conserved expression of several TFs that are characteristic of the anterior forebrain and placodes in chordates^3^ (e.g., *SP8/9, PAX6, FOXG1, ETV1*; Extended Data Fig. 8a, s), ii) the unique expression of *PRDM12* (expressed in pain-sensing nerve cells and V1 interneurons in gnathostomes^53,54^; Extended Data Fig. 8a, t), and iii) the presence of dopaminergic cells (type: TeDopa1; Extended Data Fig. 8a). *SP8/9*^+^ neurons are present also in the sub-pallium (type: TeInh4), within the medial preoptic nucleus (MPO; Fig. 4d) where they co-express *ISL1/2*, a marker of striatal projection neurons in gnathostomes (Extended Data Fig. 8a). The presence of *SP8/9*^+^-*ISL1/2*^+^ and *SP8/9*^+^-*ETV1*^+^ neurons in the sub-pallium and OB, respectively, is already known for mammals, where they originate from the lateral ganglionic eminence (LGE)^55^, suggesting that these two cell populations share the same developmental origin and migratory patterns across vertebrates.

Another important sub-pallial progenitor zone in jawed vertebrates is the medial ganglionic eminence (MGE), a developmental source of pallidal, striatal, and cortical GABAergic interneurons. The occurrence of an MGE in the cyclostome brain was proposed by Sugahara and colleagues, based on the presence of an *NKX2-1/4*^+^ domain in the sub-pallium of lamprey and hagfish embryos^7^. We identified neurons (type: TeInh5) expressing *LHX6/8* (a marker of MGE-derived cells in mammals) and showing residual expression for *NKX2-1/4a* (Fig. 4e) within two sub-pallial regions: i) the striatum, a structure located dorsal to the MPO^56^ (Fig. 4d), and ii) the putative pallidum^57^, a nucleus located ventrolaterally to the thalamic eminences (Extended Data Fig. 8j). Within the striatum, this neuron type is organized into an internal (*SSTc*^+^) and external layer (*PENK*^+^) (Fig. 4f; Extended Data Figs. 8m, n). Recursive clustering revealed the presence of subtypes that express markers of both striatal (*DRD2, PENK*) and pallidal (*TACR1, GBX1*) neurons (Fig. 4e). These results indicate that, unlike in gnathostomes, where the striatum is mostly LGE-derived and the pallidum MGE-derived, the lamprey striatum and pallidum are very similar at the transcriptomic level and are both MGE-derived.

The presence of *DLX*^+^ GABAergic neurons expressing LGE- and MGE-related markers in the pallium (Fig. 4d, h; Extended Data Fig. 8i, n, o) implies that migration from progenitor zones of the sub-pallium also occurs in lamprey. Many of these neurons express the neuropeptide genes *PENK*, and *SSTc*, which mark GABAergic interneurons types in the pallium of several gnathostome species^11,58,59^ (Fig. 4f; Extended Data Fig. 8m, n, q, r). The vasoactive intestinal peptide (*VIP;* a marker of a sub-population of cortical GABAergic interneurons in amniotes) is also present in the lamprey pallium, but, contrary to gnathostomes, it is expressed exclusively in glutamatergic neurons (Fig. 4c; Extended Data Fig. 8a, p).

The expression programs of excitatory neurons of the lamprey telencephalon are overall highly correlated to those of the corresponding cell types in mouse (Fig. 2f). This similarity is confirmed by the expression of marker genes typical of mammalian cortical glutamatergic neurons within the lamprey pallium and, partially, OB (e.g., *TBR1, EMX1/2*a, *EMX1/2*b, *RTN4R, LHX2/9, BCL11B, IKZF1/3*; Fig4h; Extended Data Fig. 7h, i; Extended Data Fig. 8a, f-h, j-l, o, q). We identified eight distinct cell types populating four different regions of the lamprey pallium: i) dorsomedial telencephalic nucleus (DMTN; type: TeExc1), ii) medial pallium (MPa; type: TeExc4), iii) sub-medial pallium (SMPa; type: TeExc3), and iv) lateral pallium (LPa; types: TeExc2, TeExc5-8) (Fig. 4g; Supplementary Table 3; see online atlas). DMTN is a relay nucleus that is innervated by tufted-like cells of the OB^60^ and is located at the interface between the pallium and OB, of which it constitutes the caudal-most portion. Like the OB the DMTN displays a layered structure with outer glutamatergic neurons, which share the same expression profile with cells of the OB glomerular layer, (e.g., *EBF1*) and inner GABAergic (*PRDM12*^+^) neurons (Extended Data Fig. 8o, p, r-u). MPa and SMPa neurons express the TFs *OTX2* and *NR2F1/6a*. They extend caudally, reaching the thalamic eminences, and are defined by the expression of *EBF1, SSTc* (TeExc4) and *C1QL3, PNOC* (TeExc3) (Fig. 4g; Extended Data Fig. 8a, h, n, r). We find that LPa neurons form a three-layered cortex with an inner GABAergic/glutamatergic layer, a middle glutamatergic layer, and an external molecular, fiber-rich layer, in accord with previous work^61^ (Fig. 4h-k). They all express several genes associated with cortical projection neurons in amniotes (e.g., *FOXP1/2/4, MEIS2, LAMP5, RORB, TCAP*; Extended Data Fig. 8a). However, contrary to what is known for amniotes, we did not observe any regional specification of gene expression patterns among these neurons (e.g., dorsal vs. ventral) that could be related to known, functionally distinct areas of the lateral pallium (e.g., somatosensory, visual, motor, olfactory)^62,63^.

## Discussion

In this study, we used extensive scRNA-seq and targeted spatial transcriptomics data to create a neural cell type atlas for a cyclostome representative: the sea lamprey (https://lampreybrain.kaessmannlab.org/). Our comparisons between filter-feeding larval (ammocoete) and parasitic adult stages suggest that the larval brain might contain sets of not yet fully differentiated cells, even though lampreys spend most of their life cycle as ammocoetes^2^. Our work thus implies that the adult stage should be used to assess the cell type diversity of lampreys and its comparison with that of other species. This notion is in agreement with the recent discovery that ammocoetes represent an evolutionarily derived and not an ancestral life stage of lampreys^64^.

Our cell type tree analyses revealed that lampreys and gnathostomes share a common fundamental cellular and molecular organization of the brain that emerged in the vertebrate stem lineage more than ∼515-645 MYA ago. This finding is in line with the shared broad brain regionalization (Fig. 1a, Extended Data Fig. 11b) and previously described patterning mechanisms across vertebrates^1^. Our comparisons of lamprey and mouse cell types revealed homologous relationships for many cell type families; that is, we identified groups of cell types partly sharing the same developmental processes and gene expression programs. These cell type families likely constituted the core of the ancestral vertebrate cell type repertoire.

However, the lamprey brain lacks key cell types present in the gnathostome brain. Notably, our work confirms the absence of oligodendrocytes and sheds new light on their origination. We found that lamprey astrocytes express several oligodendrocyte-specific genes, including master regulators and effector genes (Fig. 3b). Our observations suggest that key components of the molecular machinery of oligodendrocytes were present in astrocyte-like cells of the vertebrate ancestor, and indicate that oligodendrocytes originated from these evolutionary precursors on the gnathostome lineage (Extended Data Fig. 11b). Our work thus extends previous studies, which showed that lamprey axons seem to be physically associated with astrocytes^26^ and that key aspects of the regulatory program required for oligodendrocyte differentiation in gnathostomes are present during lamprey gliogenesis^33^. Our analyses also failed to provide evidence for the presence of granule or Purkinje cells in the rostral hindbrain, strongly supporting the notion that the mature lamprey brain lacks a proper cerebellum. We note, however, that a recent study detected expression of granule and Purkinje cell TFs in the dorsal rhombomere 1 of lamprey embryos^65^. A targeted prospective analysis of the dorsal isthmic region in the adult and developmental lamprey brain might thus reveal the presence of potential rare homologs of cerebellar cell types.

The discovery of both LGE- and MGE-derived inhibitory neurons in the lamprey telencephalon confirms that the two main GABAergic progenitor zones of the sub-pallium were already present in the common vertebrate ancestor^7,66^ (Extended Data Fig. 11b). In the mammalian sub-pallium such inhibitory neurons largely contribute to the two core basal ganglia nuclei: the striatum and pallidum, with projection neurons (LGE-derived) in the striatum projecting to the pallidum (MGE-derived). The presence of similar pathways between analogous regions was demonstrated in lamprey, implying that the main basal ganglia circuitry is shared by all vertebrates^57^. By contrast, we show that – unlike the striatum of jawed vertebrates – the lamprey striatum is mainly MGE-derived, as is its putative pallidum, suggesting that these two structures have a common developmental origin (from the MGE) and that the striata of lamprey and jawed-vertebrates are populated by non-homologous cell types. Notably, a population of LGE-derived cells (TeInh4) is located immediately ventral to the lamprey striatum, in the MPO region (Fig. 4d). However, its contribution to the basal ganglia circuitry is still unknown. Outside the lamprey sub-pallium, LGE- and MGE-derived cells also contribute to GABAergic interneurons of the OB and pallium, indicating that their migratory patterns are conserved across vertebrates (Extended Data Fig. 11b).

Within the pallium, we identified groups of cell types that are likely homologous to glutamatergic mammalian cortical neurons, supporting the hypothesis that the core cell types composing cortical/nuclear circuits across jawed vertebrates emerged in common vertebrate ancestors^63,67^. These neurons express genes that are associated with different projection modalities (e.g., input, intratelencephalic, output), but not in the same combinations as observed in jawed vertebrates^68^.

Altogether, our study provides the first global view of the cellular composition and molecular architecture of the ancestral vertebrate brain, and provides the groundwork for investigating its extensive cellular and structural diversification during vertebrate evolution.

## Acknowledgements

We thank all members of the Kaessmann group for the fruitful discussions; Elisa Panzariello, Marta Sanchez-Delgado, Nils Trost for the brain and animal illustrations, Moritz Mall for providing temporary lab-space and assistance, and Margarida Cardoso-Moreira for discussions and comments on the manuscript. Computations were performed on the Kaessmann lab server (managed by Nils Trost) and the bwForCluster from the Heidelberg University Computational Center (supported by the state of Baden-Württemberg through bwHPC and the German Research Foundation – INST 35/1134-1 FUGG). This research was supported by grants from the European Research Council (615253, OntoTransEvol), European Commission (Marie Skłodowska-Curie Actions ITN: EvoCELL), and the Tschira foundation, which funded the Illumina NextSeq machine used for sequencing. DMM and DJ were supported by the grants of National Science Foundation (IOS 1656843), National Institutes of Health/National Institute of Dental and Craniofacial Research (RDE025940), and University of Colorado, Boulder RIO Innovative Seed Grant FY21 (all to DMM). DJ was also supported by the grant from the Scientific Grant Agency of the Slovak Republic (VEGA 1/0450/21).

## Author contributions

F.L, F.H.-S., and H.K. conceived and organized the study based on H.K.’s original design. F.L, F.H.-S., and H.K wrote the manuscript with input from all authors. F.L. performed all analyses, and developed the brain atlas app. F.H.-S. established and optimized the tissue dissociation protocol, and performed all scRNA-seq and *in situ* experiments with support from A.P.O., J.S., and C.S., and guidance from M.S. F.L. and F.H.-S. annotated and interpreted the data. T.B. prepared bulk libraries with guidance from K.M. M.S. established the smFISH protocol. F.H.-S., A.P.O., D.J., D.S.C., M.L.M, and S.A.G. collected the samples. A.B.I., D.M.M., M.B., and M.C.R. provided samples. J.J.S. provided early access to genome assemblies and annotations. A.P.O., M.S., F.M., D.S.C., A.B.I., D.M.M., and D.A. provided useful feedback and discussions. H.K. supervised the study and provided funding.

## Methods

### Sea lamprey samples

Sea lamprey (*Petromyzon marinus*) samples were dissected from specimens obtained from three different sources (Supplementary Table 1). Sampled animals were euthanized by submersion in 0.1% MS-222 (Sigma, A5040-25G), unless specified otherwise, followed by decapitation according to local guidelines. Tissue samples from larvae (i.e., ammocoetes, between 90 and 130 mm in body length), juveniles (Youson stages 6-7) and adults used for bulk-tissue RNA-seq and genome annotation (see below) were collected from freshwater streams in Maine, USA, and held in large, aerated tanks with sand and freshwater until being sacrificed. All procedures were approved by the University of Colorado, Boulder Institutional Animal Care and Use Committee as described in protocol 2392. Larvae (between 70 and 120 mm in body length) used for scRNA-seq, smRNA-FISH, and Cartana experiments were collected form the River Ulla in Galicia, Spain, and kept at the Interfaculty Biomedical Research Facility (IBF) of Heidelberg University in freshwater aerated tanks with river sediment and appropriate temperature conditions (∼15°C) until used for tissue collection. All animal procedures were performed in accordance with European Union and German ethical guidelines on animal care and experimentation, and were approved by the local animal welfare authorities (Regierungspräsidium Karlsruhe). Upstream migrating mature adults used for scRNA-seq experiments were obtained from a commercial supplier (Novas Y Mar, Galicia, Spain) and were processed immediately upon their arrival to the laboratory. All procedures were approved by the Bioethics Committee of the University of Santiago de Compostela and the Xunta de Galicia Government and conformed to European Union and Spanish regulations for the care and handling of animals in research. Adult specimens used for Cartana experiments were obtained from the US Fish and Wildlife Service and Department of the Interior and were euthanized by immersion in 0.25% MS-222, followed by decapitation. All procedures were approved by the California Institute of Technology Institutional Animal Care and Use Committee (IACUC) protocol 1436.

### RNA extraction and sequencing of bulk tissue samples

In total, 63 sea lamprey tissue samples from six organs (brain, heart, liver, kidney, ovary, testis) were dissected from larval, juvenile, and adult specimens. Total RNA was extracted using different extraction protocols (Supplementary Table 1); RNA quality was inspected using the Fragment Analyzer (Advanced Analytical Technologies) and its concentration was determined using a NanoDrop (Thermo Fisher Scientific). Strand-specific RNA-seq libraries were generated using the Illumina TruSeq Stranded mRNA Library protocol. Each library was sequenced on Illumina HiSeq 2500 platforms (100nt, single-end) at the Lausanne Genomic Technologies Facility (https://www.unil.ch/gtf).

### Sea lamprey genome annotation

Bulk tissue RNA-seq reads were mapped to the sea lamprey germline genome^9^ using GSNAP^69^ (version: 2018-03-01) with the option to find known and new splice junctions in individual reads activated (--novelsplicing=1). The resulting BAM files for each stage and tissue were merged before being used for transcriptome assembly with StringTie^70^ (v1.3.4d). Each resulting GTF file was filtered for putative assembly artifacts using GffRead^71^ (v0.9.9) by discarding single-exon transcripts and multi-exon mRNAs that have any intron with a non-canonical splice site consensus (i.e., not GT-AG, GC-AG, or AT-AC). Individual annotated transcriptomes were then merged together with the already available set of annotated protein-coding genes from the germline genome study^9^ in order to obtain a non-redundant set of transcripts. Genome annotation was further refined using TransDecoder (v5.3.0; https://github.com/TransDecoder/TransDecoder) in order to identify candidate coding-regions within transcript sequences; this process involves the identification of the longest putative Open Reading Frame (ORF) within each transcript and the subsequent search of the corresponding peptides against SwissProt (https://uniprot.org) using BlastP^72^ (v2.5.0+) and Pfam (https://pfam.xfam.org) using HMMER^73^ (v3.2). Annotation quality was assessed by comparing the number of reads mapping to exonic, intronic, and intergenic regions of the genome (Extended Data Fig. 1). Annotation completeness was also estimated using BUSCO^74^ (v3) by comparing the set of translated longest CDS from each transcript against a set of metazoan-conserved single-copy orthologs from OrthoDB^75^ (Supplementary Table 6).

### Orthology assignment

Homology information for the set of annotated genes was retrieved by applying the OrthoFinder^76^ (v2.3.11) pipeline against a group of selected chordates: vase tunicate (*Ciona intestinalis*), inshore hagfish (*Eptatretus burgeri*), Australian ghostshark (*Callorhinchus milii*), spotted gar (*Lepisosteus oculatus*), zebrafish (*Danio rerio*), West Indian Ocean coelacanth (*Latimeria chalumnae*), Western clawed frog (*Xenopus tropicalis*), red junglefowl (*Gallus gallus*), house mouse (*Mus musculus*), and human (*Homo sapiens*). By reconstructing a complete set of rooted gene trees among the analyzed species, this tool allows to establish all orthology relationships among all genes, and to infer duplication events and cross reference them to the corresponding nodes on the gene and species trees. Proteomes were downloaded from Ensembl (remaining species; v97) databases and used for a BlastP Best Reciprocal Hit (BRH) analysis; in order to avoid redundancies in the blast results, only the peptides coming from the longest isoform within each gene were used. Rooted gene trees from the inferred orthogroups – i.e., groups of genes descended from a single gene in the Last Common Ancestor (LCA) – were obtained using Multiple Sequence Alignments (MSA; MAFFT^77^ v7.455) with IQ-TREE^78^ (v1.6.12; 1,000 bootstrap replicates) and STRIDE^79^. Orthology relationships can be explored in our online atlas.

### Cell dissociation and single cell RNA-seq data generation

Larval and adult heads were air dissected and brains were placed in 1x HBSS (Life Technologies, 14185052) for cleaning and removal of the meninges. Once cleaned, brains were further treated as a whole sample or, for the second set of experiments, divided in regions (telencephalon, diencephalon, mesencephalon and rhombencephalon). Brain tissue was dissociated using the Papain Dissociation System (Worthington, LK003150), according to the manufacturer’s protocol, with the following modifications: the tissue was incubated in papain solution (volume adjusted for tissue size, 100 – 300 µl) at 28 °C for 15 min under constant agitation. Then, the tissue was gently and collected by centrifugation for 1 min at 300g. This step was followed by a second incubation in fresh papain solution and a final trituration, performed as described above. The dissociated cells were spun down at 300g for 5 min and resuspended in the inhibitor solution (prepared following the Papain Dissociation System specifications). The suspension was filtered using a 40 µM falcon strainer (Sigma-Aldrich, CLS431750-50EA) and, immediately after, a discontinuous density gradient was performed. The cells were then resuspended in Leibovitz’s L-15 Medium (Life Technologies, 21083027) reaching a final volume between 50 and 100 µl, depending on the original tissue size. Cells were examined for viability and counted using a trypan blue staining and a Neubauer counting chamber (Assistent).

After ensuring a cell viability greater than 90% and a concentration equal or higher than 300 cells per µl, cell suspensions (∼15,000 cells per reaction) were loaded onto the Chromium system (10x Genomics). cDNA amplification and scRNA-seq libraries were constructed using Single-Cell 3′ Gel Bead and Library v2 (for larvae) and v3 kits (for adults), following the instructions of the manufacturer. For one larval whole brain we additionally produced a library using the v3 kit (SN352), in order to confirm observed (biological) differences between the larval and adult datasets; that is, to rule out that technical differences explain the observed differences (Supplementary Fig. 4b, c). cDNA libraries were amplified using 12-13 PCR cycles and quantified on a Qubit Fluorometer (Thermo Fisher Scientific). Average fragment size was determined on a Fragment Analyzer (Agilent). Libraries were sequenced using the NextSeq 500/550 High Output Kit v2.5 on the Illumina NextSeq 550 system (28 cycles for Read 1, 56 cycles for Read 2, 8 cycles for i7 index and 0 cycles for i5 index).

### scRNA-seq data processing

scRNA-seq reads were mapped to the reference genome^9^ with our extended annotation (see above), and Unique Molecule Identifier (UMI) count matrices were produced using CellRanger v3.0.2 (10x genomics). Cell-containing droplets were obtained from the CellRanger calling algorithm and validated by checking: i) the cumulative distribution of UMIs; ii) the distribution of UMIs coming from mitochondrial genes; iii) the distribution of the proportion of UMIs coming from intronic regions. Putative multiplets (i.e., droplets containing more than one cell) were identified using DoubletFinder^80^ and Scrublet^81^; droplets labeled as multiplets by any of the two methods were removed from the count matrices.

The obtained count matrices were analyzed using Seurat v3.1.5^82^ and pre-processed by keeping only genes expressed in at least five cells and by removing cells containing less than 200 UMIs and more than 5% (ammocoete) or 10% (adult) mitochondrial UMIs. Raw UMI counts were then normalized using the SCTransform method^83^ and the top 3,000 Highly Variable Genes (HVGs) across all cells were used for subsequent analyses. Principal Component Analysis (PCA) was applied to the normalized HVG matrices, and the resulting 75 most significant PCs were used for building a Shared Nearest Neighbor (SNN) graph that was then clustered using the Louvain method with different resolution values (0.5-10). Differential expression analysis was run in order to find potential marker genes from all clusters across all resolution values (Wilcoxon Rank Sum Test: logFC ≥ 0.25; min.pct = 0.1; Bonferroni-adjusted p-value < 0.01). The PCA-transformed matrices were finally embedded in two-dimensional space using Uniform Manifold Approximation and Projection (UMAP) and t-distributed Stochastic Neighbor Embedding (t-SNE) dimensionality reduction techniques.

The clustered cells were further manually inspected in order to identify and then remove spurious clusters (i.e., clusters composed by damaged/stressed cells or multiplets/empty droplets that escaped the previous filtering steps). Cell types/states were annotated on top of the clusters obtained using the highest resolution value (10); a putative phenotype/function was assigned to each cluster by allocating marker genes to any of the following Gene Ontology^84^ (GO) categories: transcription (co-)factor, neurotransmitter metabolism, neurotransmitter transport, neurotransmitter receptor, neuropeptide^85^, neuropeptide receptor^85^, immune response, erythrocyte differentiation, blood vessel development, neurogenesis, gliogenesis. Annotated clusters that were contiguous on the UMAP and t-SNE embeddings were manually inspected and joined together if they were showing similar expression patterns among their respective marker genes. Additional functional information was added by comparing the annotated clusters to published vertebrate neural single-cell datasets^17,86^.

Datasets coming from different samples were integrated using integrative non-Negative Matrix Factorization (iNMF) as implemented in LIGER v0.5.0^87^. Datasets were integrated at two levels: i) integration of replicates coming from the same brain region (i.e., telencephalon, diencephalon, mesencephalon, rhombencephalon and whole brain) and stage (i.e., ammocoete, adult); ii) integration, within each stage, of all replicates together in the same dataset encompassing all sampled regions. Each integrated dataset was then imported to Seurat to perform SNN graph construction, clustering, DE analysis, 2D-embedding and cluster annotation as described above.

We noticed that the number of UMIs per cell was sensibly and consistently lower for the larval dataset (produced using Chromium kit v2) compared to the adult one (produced using Chromium kit v3; Supplementary Tables 1, 2). In order to establish whether this difference reflected an actual biological property of the two stages, we downsampled the adult count matrices to 50% and assessed its impact on cluster resolution (Extended Data Fig. 4a-c). In addition, we produced a larval dataset using the v3 kit and compared its number of UMIs per cell to the larval v2 and adult v3 datasets (Extended Data Fig. 4b, c) (see also data generation section above).

### Lamprey-mouse comparisons

In order to find cross-vertebrate similarities and differences in neural cell types, the adult integrated brain atlas was compared against a published juvenile mouse nervous system atlas^17^. The two datasets were first compared via a correlation-based approach. That is, the raw UMI count matrices were extracted from both species datasets and orthology information for the corresponding gene IDs was added; orthology relationships between mouse and lamprey were obtained from the OrthoFinder analysis (see above; Supplementary Table 7). The UMI counts coming from paralogs in the respective species were summed (“meta-gene” method^19^) and the species-specific gene IDs were replaced by numeric indexes (1..*n*, where *n* is the number of orthology groups between the mouse and lamprey) shared by the two species. The new “meta-gene” count matrices were then normalized using SCTransform, filtered for HVGs, and averaged across all annotated clusters. Expression levels were finally transformed to Specificity Indexes (SI) using the method of Tosches and colleagues^11^ and used for Pearson correlation analyses. Dendrograms relating cell-type families between lamprey and mouse were constructed using the pvclust^88^ R package with complete hierarchical clustering and 1,000 replicates.

In addition, the two datasets were compared using the Self Assembling Manifold mapping (SAMap; v0.2.3) algorithm^18^, a method that enables mapping single-cell transcriptomic atlases between phylogenetically distant species. A gene-gene bipartite graph with cross species edges connecting homologous gene pairs was constructed by performing reciprocal BlastP searches between the two proteomes of the two species. The graph was used in a second step to project the two datasets into a joint, lower-dimensional manifold representation, where expression correlation between homologous genes was iteratively used to update the homology graph connecting the two atlases. After the analysis was run, a mapping score (ranging from 0 to 1) was computed among all possible cross-species cluster pairs.

### *In situ* sequencing

Whole brains (adults) and heads (larvae) were embedded in OCT mounting medium and then flash-frozen by laying them on isopentane, previously cooled on liquid nitrogen. Adult tissues were rinsed with ice cold PBS before being frozen. Tissues were cryosectioned in 10 µm coronal and sagittal sections and stored at −80 °C until further use. Sections were processed for *in situ* sequencing using the High Sensitivity Library Preparation Kit from CARTANA AB (10x Genomics). The method and data processing are described by Ke and colleagues^10^. Processing of sections was done following CARTANA’s protocol with minor modifications. In brief, sections on SuperFrost Plus glass slides (Thermo Fisher Scientific) were air dried for 5 min. Afterwards, sections were fixed by 3.7% (v/v) paraformaldehyde in UltraPure distilled water (DNase/RNase-Free, Thermo Fisher Scientific, 10977035) for 7 min and washed in PBS (Thermo Fisher Scientific, 70011036; diluted in UltraPure distilled water), followed by 0.1 N HCl treatment for 5 min and a wash with PBS. The sections were then dehydrated with ethanol and air dried before covering them with SecureSeal hybridization chambers (Grace Bio-Labs, 10910000). All subsequent steps, including probe hybridization and ligation, amplification, fluorescent labeling and quality control imaging, followed manufacturer’s specifications. Finally, mounted sections were shipped to CARTANA’s facility (Solna, Sweden) for *in situ* sequencing.

### Single molecule RNA-FISH

Larval whole heads were snap frozen and cryosectioned (horizontal sections) as described above. This time, however, the sections were collected on coverslips (22 mm x 22 mm) previously pretreated with a silanization solution (0.3% (v/v) bind-silane (GE Healthcare Life Sciences, 17-1330-01), 0.1% (v/v) acetic acid and 99.6% (v/v) ethanol).

To reduce tissue autofluorescence, sections were embedded in polyacrylamide gel, RNAs were anchored to the gel by LabelX treatment, and cellular proteins and lipids were cleared as previously described^89,90^, with modifications. LabelX solution was prepared by reacting Label-IT (Mirus Bio) with Acryloyl X – SE (Thermo Fischer Scientific) as described by Chen and colleagues^90^. Specifically, sections were air dried for 15-20 min and fixed in 3.7% paraformaldehyde in PBS for 10-15 min, followed by a 2 min incubation in 4% SDS in PBS and washes with PBS. Fixed sections were then incubated in 70% ethanol at 4°C for at least 16h. Next, sections on coverslips were washed twice with PBS, once with 1x MOPS pH 7.7 (Sigma-Aldrich, M9381) and incubated with LabelX (diluted to a concentration of 0.006 mg/mL in 1xMOPS) at room temperature for 4 hours, followed by two PBS washes. To anchor LabelX-modified RNAs, sections were embedded in thin 4% polyacrylamide (PA) gels. First, coverslips were washed for 2 min with a PA solution, consisting of 4% (v/v) of 19:1 acrylamide/bis-acrylamide (Sigma-Aldrich, A9926-5), 60 mM Tris·HCl pH 8, and 0.3 M NaCl. Coverslips were then washed for 2 min with the PA solution supplemented with ammonium persulfate (Sigma-Aldrich, 7727-54-0) and TEMED (Sigma-Aldrich, T7024) at final concentrations of 0.03% (w/v) and 0.15% (v/v), respectively. To cast the gel, 75 µl of the PA solution (supplemented with the polymerizing agents), was added to glass slides previously treated with Repel Silane (GE Healthcare Life Sciences, 17-1332-01) and washed with ethanol. Each coverslip was then layered on top of a slide, with one drop of PA solution, ensuring that a thin PA layer forms between the slide and the coverslip. The gel was allowed to cast at room temperature for 1.5 h. Coverslips and slides were gently separated leaving coverslips with sections embedded into the PA gel. Coverslips were then washed with digestion buffer consisting of 0.8 M guanidine-HCl, 50 mM Tris·HCl pH 8, 1 mM EDTA, and 0.5% (v/v) Triton X-100. Coverslips were incubated with digestion buffer supplemented with 8 U/ml of proteinase K (Sigma-Aldrich, P2308) at 37 °C for 2 – 3 h.

After background reduction, sections were hybridized with HuluFISH probes, designed and developed by PixelBiotech. The hybridization protocol followed the manufacturer’s recommendations. Briefly, coverslips were washed twice with HuluWash buffer (PixelBiotech GmbH) and incubated in 50 µl of probe solution, consisting of each probe diluted in hybridization buffer at a concentration of 1:100. Coverslips were incubated at 37 °C for 12 h, inside a light-protected humidified chamber. Afterwards, coverslips were washed 4 times with HuluWash buffer. Each wash lasted 10 min and was done at room temperature. The last wash was supplemented with Hoechst 33342 (Thermo Fisher Scientific, H3569). Coverslips were then mounted in 2 drops of Prolong Diamond mounting medium (Thermo Fisher Scientific, P36961). The mounted sections were allowed to cure at room temperature for 24 hours.

All sections were imaged on a Leica TCS-SP5, a confocal laser scanning microscope controlled by the Leica Application Suite (LAS). All images shown are the projection of mosaics built by stitching individual z-stacks. Each z-stack consisted of individual images (50 images for *SSPOa, VAT, GNRH1a*, 15 images for *ZFP704*) taken by setting a range of 10-15 µm and a step size below 0.8 µm. Images were captured with a 63x immersion oil objective and sequentially excited by a 405 nm Diode laser (for the Hoechst 33342 staining), followed by the laser required for each probe (561 nm DPSS laser for *SSPOa, ZFP704* and *VAT*; and 633 nm HeNe laser for *GNRH1a*). Projections of the z-stacks were performed in Fiji 2^91^ by using the average intensity projection. Further processing (only when required) involved contrast enhancing (saturated pixels between 0.1 and 0.3%) and background subtraction for noise reduction (rolling ball with a radius of 50 pixels).

## Data availability

Raw and processed bulk and single-cell RNA-seq data have been deposited to ArrayExpress with the accession numbers E-MTAB-11085 (bulk) and E-MTAB-11087 (single cell) (https://www.ebi.ac.uk/arrayexpress/). Additional data are available as supplementary information or upon request. Information about gene expression, cell type annotation, and gene orthology relationships across species can be visualized using the online atlas (https://lampreybrain.kaessmannlab.org/).

## Code availability

All code underlying the published atlas is available on GitHub (https://github.com/f-lamanna/LampreyBrainAtlas/) together with detailed instructions about its usage. Additional code is available upon request.

---

**Supplementary Table 1**

Lists of specimens and samples used in this study.

**Supplementary Table 2**

scRNA-seq sequencing statistics.

**Supplementary Table 3**

Lists of larval and adult detected cell types and their putative location.

**Supplementary Table 4**

Lists of *in situ* marker genes used in this study (ISS and smFISH).

**Supplementary Table 5**

Table of SAMap mapping scores for all lamprey and mouse cell types.

**Supplementary Table 6**

Results of the BUSCO analysis on the lamprey genome annotation.

**Supplementary Table 7**

Lamprey and mouse orthologs obtained with OrthoFinder.

**Supplementary Table 8**

List of all lamprey gene names used in this manuscript with their respective gene IDs (as reported by our custom annotation).

**Supplementary Data 1**

Lamprey genome custom annotation files.

**Supplementary Data 2**

*In situ* images produced in this study.

## Extended Data Figures

**Extended Data Fig. 1.**
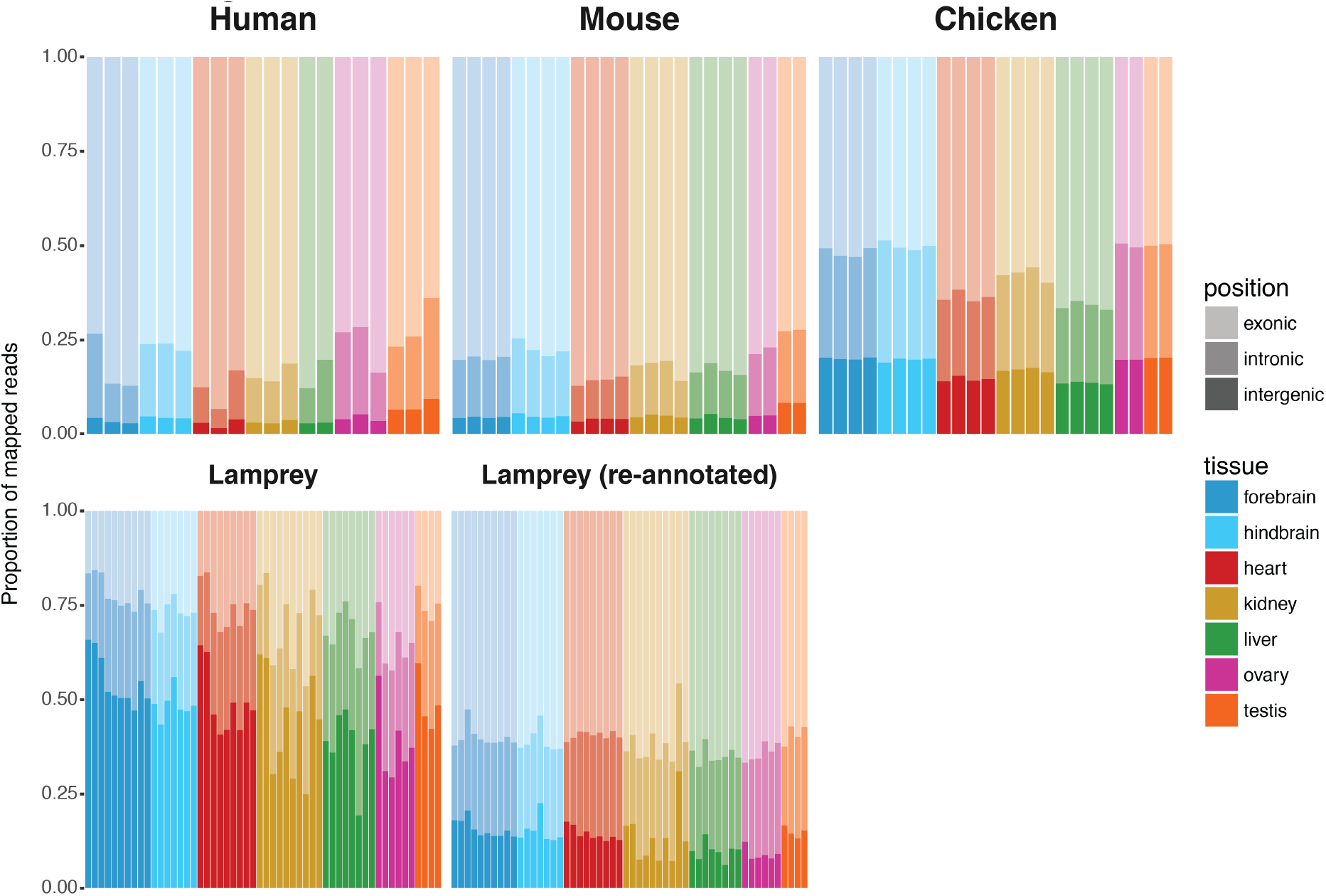
Assessment of lamprey genome annotation quality. Barplots comparing the proportion of reads mapping to exonic, intronic, and intergenic regions between the lamprey genome annotation produced in this study (re-annotated) and the published annotations of lamprey^9^, chicken, mouse, and human.

**Extended Data Fig. 2.**
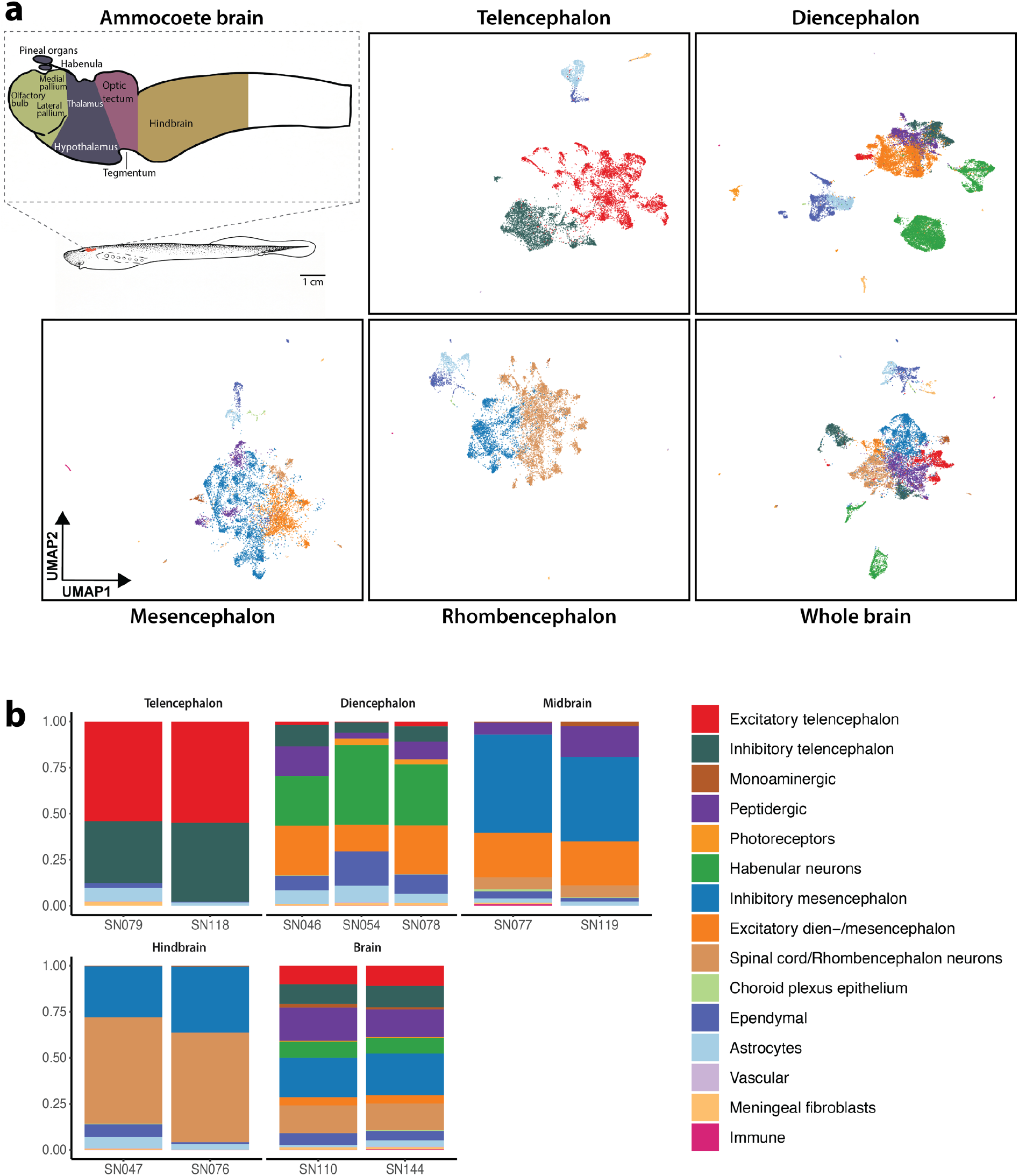
Larval (ammocoete) brain dataset. **a**, Schematic of the sea lamprey larval brain showing the different regions dissected for this study and UMAP projections for each brain region. Additional information is available in the interactive atlas (https://lampreybrain.kaessmannlab.org/ammocoete.html). **b**, Barplots showing the proportions of each cell type group (as reported in a and Fig. 1b, c) for each sample.

**Extended Data Fig. 3.**
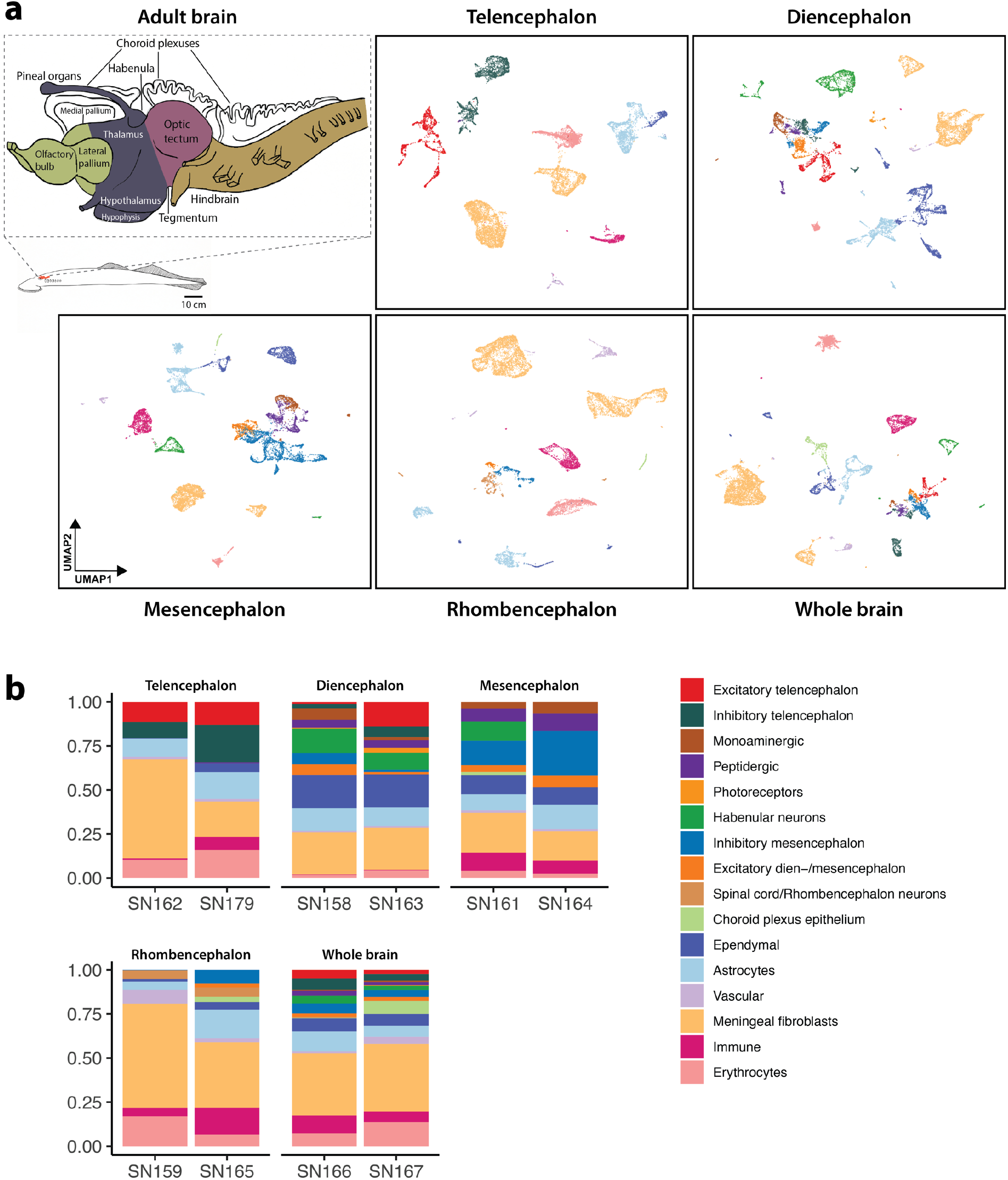
Adult brain dataset. **a**, Schematic of the sea lamprey adult brain showing the different regions dissected in this study and UMAP projections for each brain region. Additional information available in the interactive atlas (https://lampreybrain.kaessmannlab.org/adult.html). **b**, Barplots showing the proportions of each cell type group (as reported in a and Fig. 1b, c) for each sample.

**Extended Data Fig. 4.**
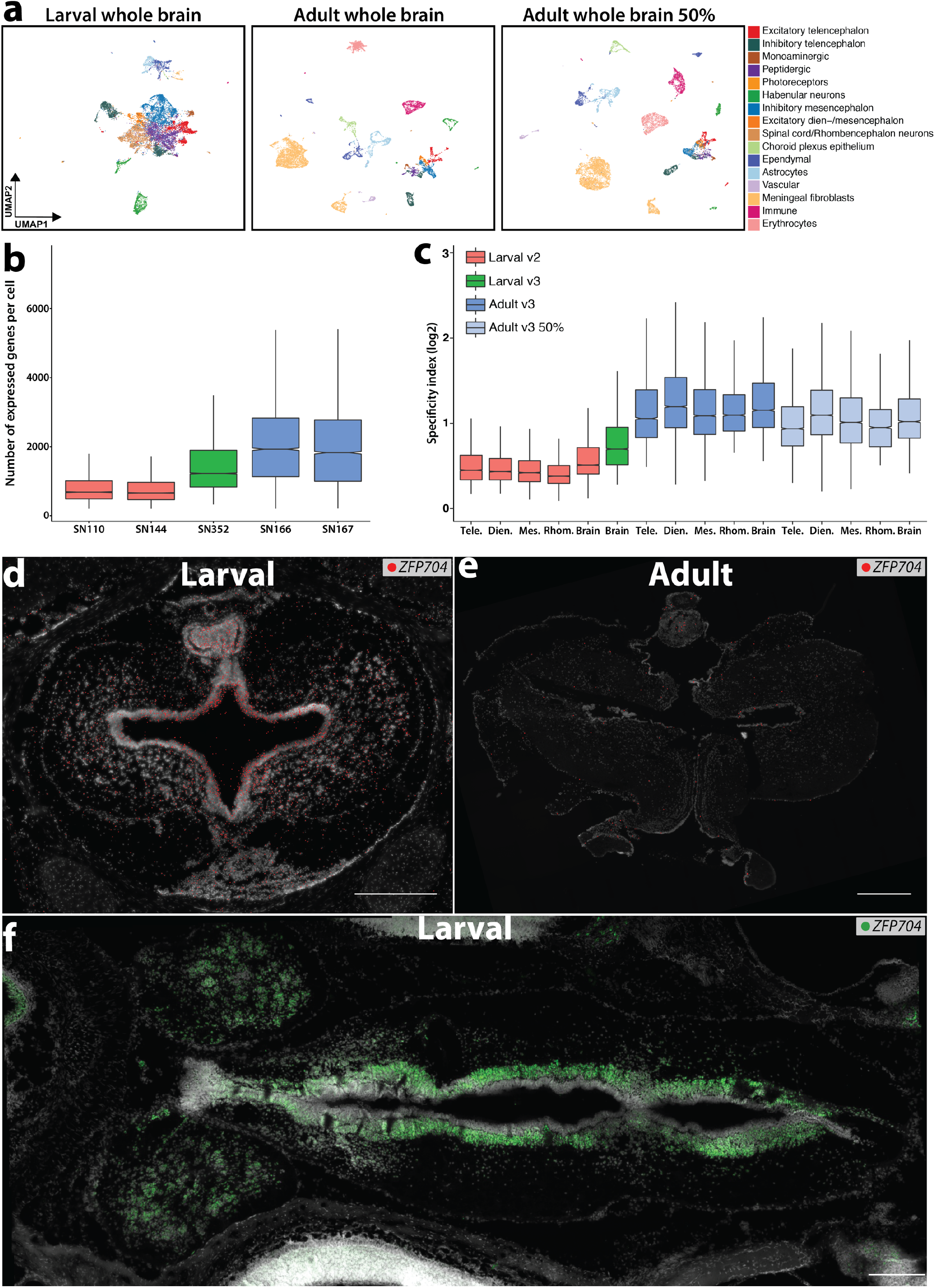
Differences between larval and adult datasets. **a**, UMAP of 13,301 and 10,557 cells from the larval (left) and adult (middle, right) whole brain datasets, respectively. Right panel: dataset downsampled to 50% of its original number of UMIs per cell in order to account for differences in the Chromium kit versions used for the larval (v2) and adult (v3) samples; see Methods. **b**, Distribution of the number of expressed genes per cell for each whole brain sample. **c**, Distributions of specificity index scores for each brain region (neurons only) across the two stages for both Chromium kit versions (larval) and for v3 version with all UMI counts and 50% of the original number of UMIs per cell (adult). **d, e**, ISS maps showing the expression of *ZFP704* on coronal sections of the larval (d) and adult (e) telencephalon (see Extended Data Fig. 12 for the ISS dissection schemes). **f**, Horizontal section (anterior end to the left) of the larval brain showing the periventricular expression of *ZFP704* (smFISH) in neurons along the whole brain. Scale bars, 500µm.

**Extended Data Fig. 5.**
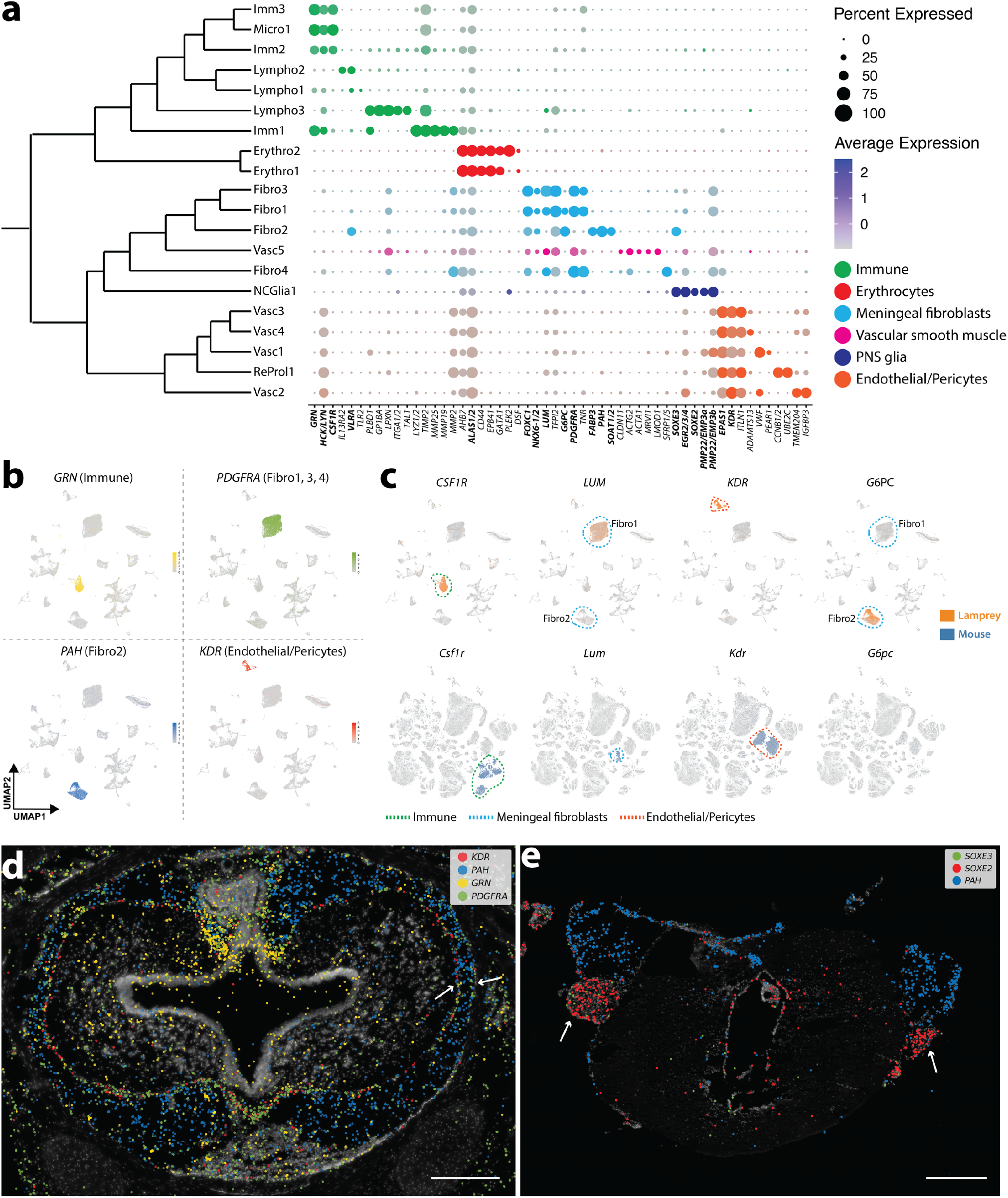
Hematopoietic, vascular, and PNS cells. **a**, Subtree from the dendrogram of Fig. 1c displaying the expression of selected marker genes for each cell type (gene names mentioned in main text are highlighted in bold). **b**, Expression of marker genes for immune (*GRN*), meningeal (*PDGFRA, PAH*), and vascular (*KDR*) cells. **c**, Expression of lamprey markers and their mouse orthologs in the respective brain atlases (UMAPs). **d**, Coronal section of the larval telencephalon showing the spatial expression of the genes shown in b (same color code). Arrows mark leptomeningeal layers. **e**, Coronal section of the adult isthmic region (mesencephalon/rhombencephalon) showing the expression of *SOXE1* and *SOXE2* within cranial nerve roots (white arrows). See Extended Data Fig. 12 for ISS section schemes; scale bars, 500µm.

**Extended Data Fig. 6.**
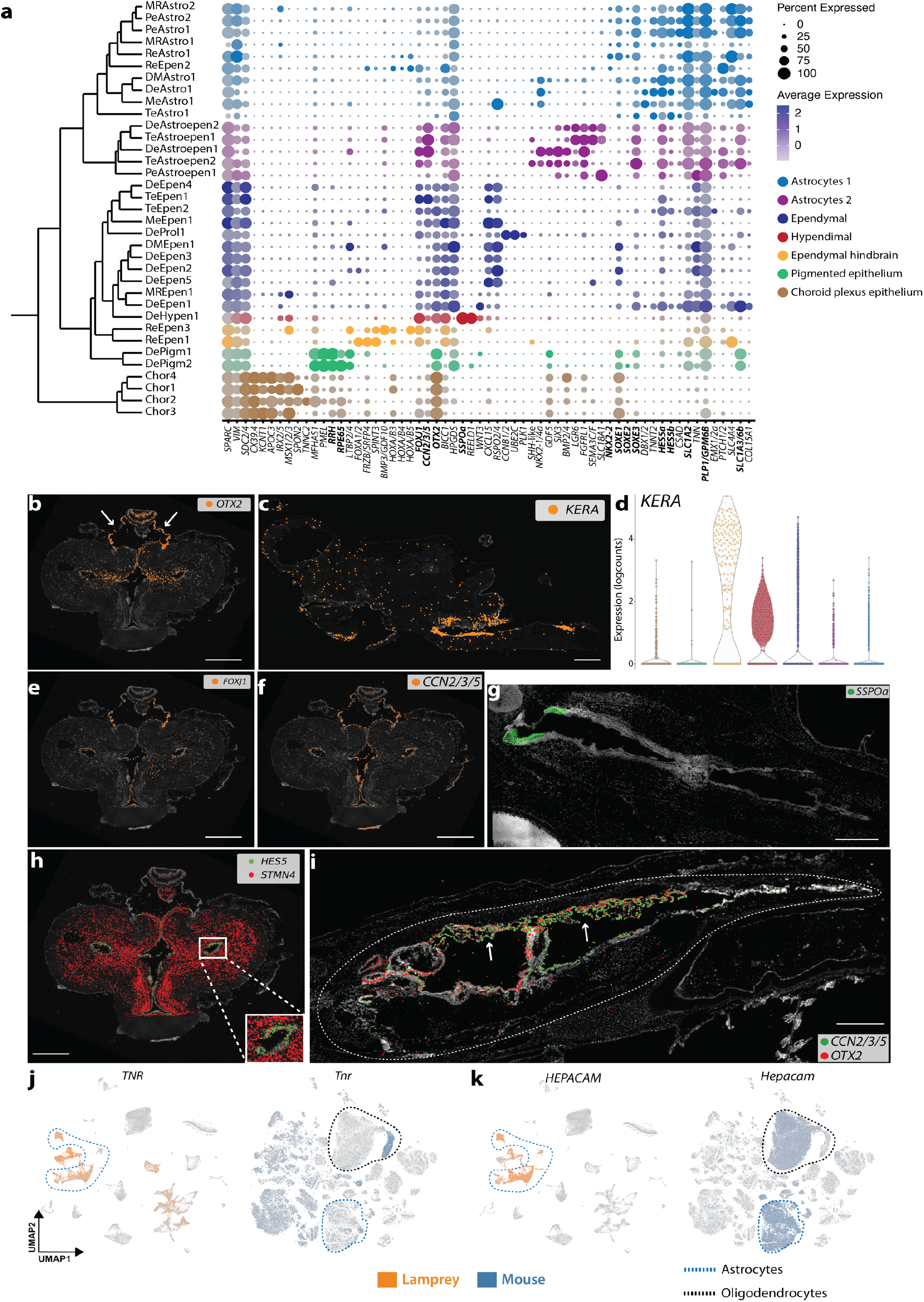
**a**, Subtree from the dendrogram of Fig. 1c displaying the expression of selected marker genes for each cell type (gene names mentioned in main text are highlighted in bold). **b**, Expression of *OTX2* within the adult telencephalon. White arrows point to choroid plexus. **c**, Expression of *KERA* within the adult brain (sagittal section; anterior end to the left), showing its concentration in the hindbrain. **d**, Violin plot displaying *KERA* expression among astroependymal cell types; color code as in a. **e, f**, Expression of *FOXJ1* (e) and *CCN1-5* (f) within the adult telencephalon. **g**, Horizontal section of the larval brain (anterior end to the upper left corner) showing the expression of *SSPO* (smFISH) around the rostral end of the third ventricle (sub-commissural organ). **h**, Expression of *STMN4* (neurons) and *HES5* (astrocytes) within the adult telencephalon showing the periventricular localization of lamprey astrocytes. **i**, Sagittal section (anterior end to the left) of a larval head (brain enclosed within white dashed line) showing the expression of *CCN1-5* and *OTX2*. White arrows point to choroid plexuses. **j, k**, Expression of *TNR* (j), *HEPACAM* (k), and their mouse orthologs on the respective brain atlases. See Extended Data Fig. 12 for ISS section schemes; scale bars, 500µm.

**Extended Data Fig. 7.**
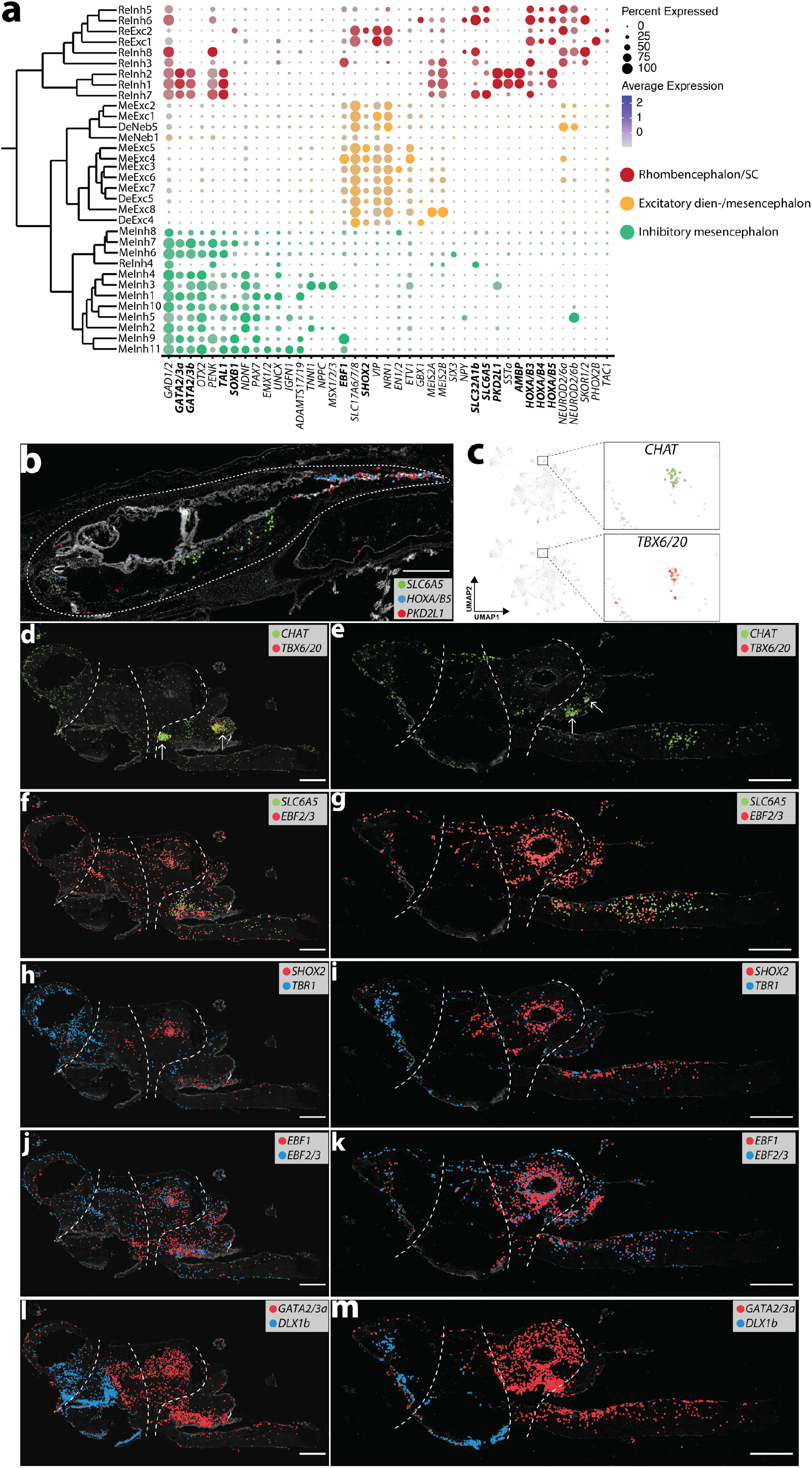
Posterior forebrain, midbrain, and hindbrain neurons. **a**, Subtree from the dendrogram of Fig. 1c displaying the expression of selected marker genes for each cell type (gene names mentioned in main text are highlighted in bold). SC, spinal cord. **b**, Sagittal section (anterior end to the left) of a larval head (white dashed line outlines the brain) showing the expression of *SLC6A5, HOXA/B5*, and *PKD2L1*. Dashed lines separate the main brain regions. **c**, UMAP projection of a larval hindbrain dataset showing the expression of *CHAT* and *TBX6/20*. **d-l** Sagittal sections (anterior end to the left) of the adult brain showing the expression of selected marker genes. Arrows in d and e indicate the putative location of cranial nerve nuclei. See Extended Data Fig. 12 for the ISS section schemes; scale bars, 500µm.

**Extended Data Fig. 8.**
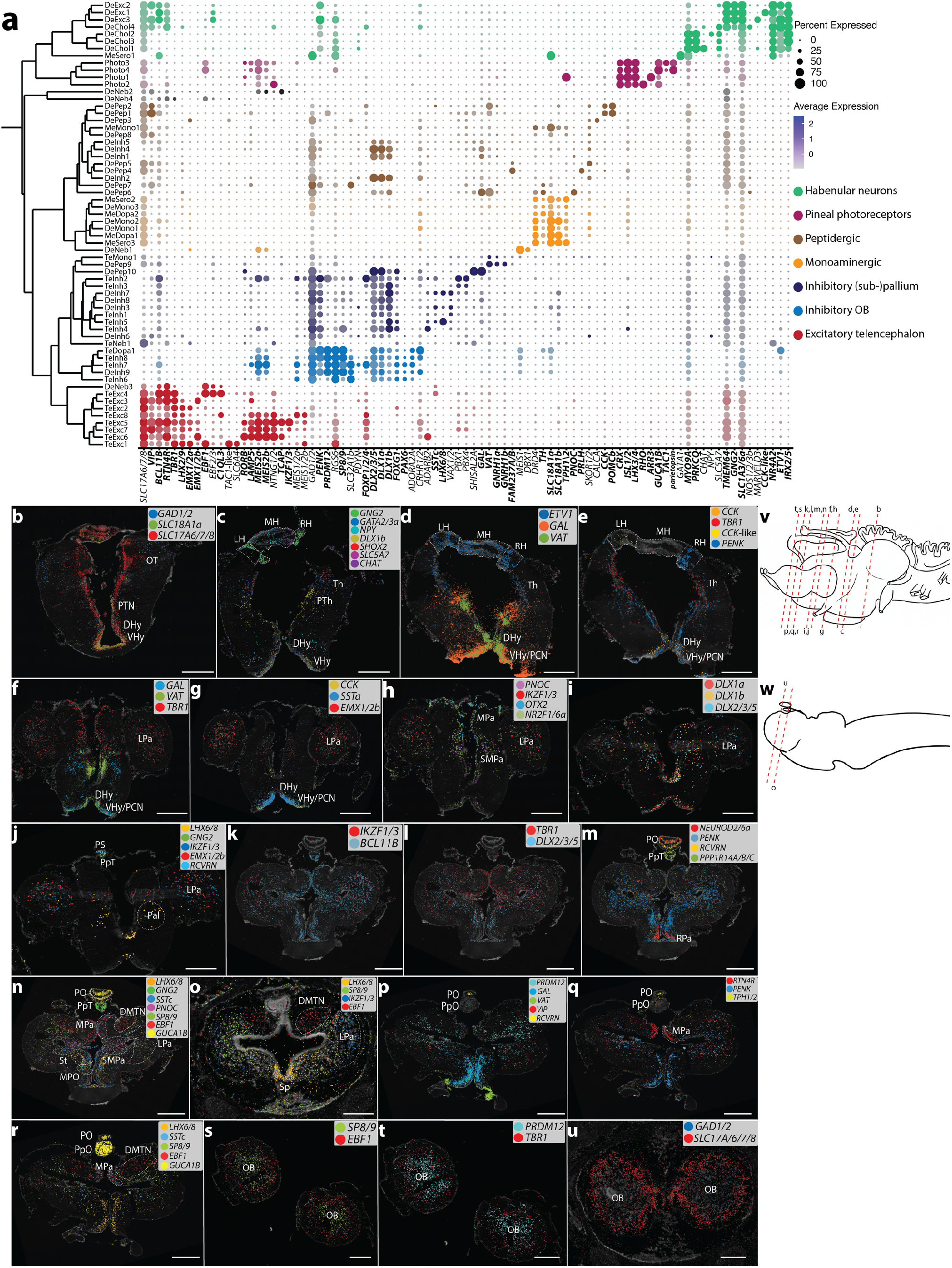
Anterior forebrain and epithalamic neurons. **a**, Subtree from the dendrogram of Fig. 1c displaying the expression of marker genes for each cell type (gene names mentioned in main text are highlighted in bold). **b-u**, ISS maps of selected neuronal marker genes across various adult and larval (o, u) brain coronal sections. DHy, dorsal hypothalamus; DMTN, dorsomedial telencephalic nucleus; LH, left habenula; LPa, lateral pallium; MH, medial habenula; MPa, medial pallium; MPO, medial preoptic nucleus; OB, olfactory bulb; OT, optic tectum; Pal, pallidum; PCN, postoptic commissure nucleus; PO, pineal organ; PpO, parapineal organ; PpT, parapineal tract; PS, pineal stalk; PTh, pre-thalamus; PTN, posterior tubercle nucleus; RH, right habenula; RPa, rostral paraventricular area; SMPa, sub-medial pallium; Sp, septum; St, striatum; Th, thalamus; VHy, ventral hypothalamus. Scale bars, 500µm.

**Extended Data Fig. 9.**
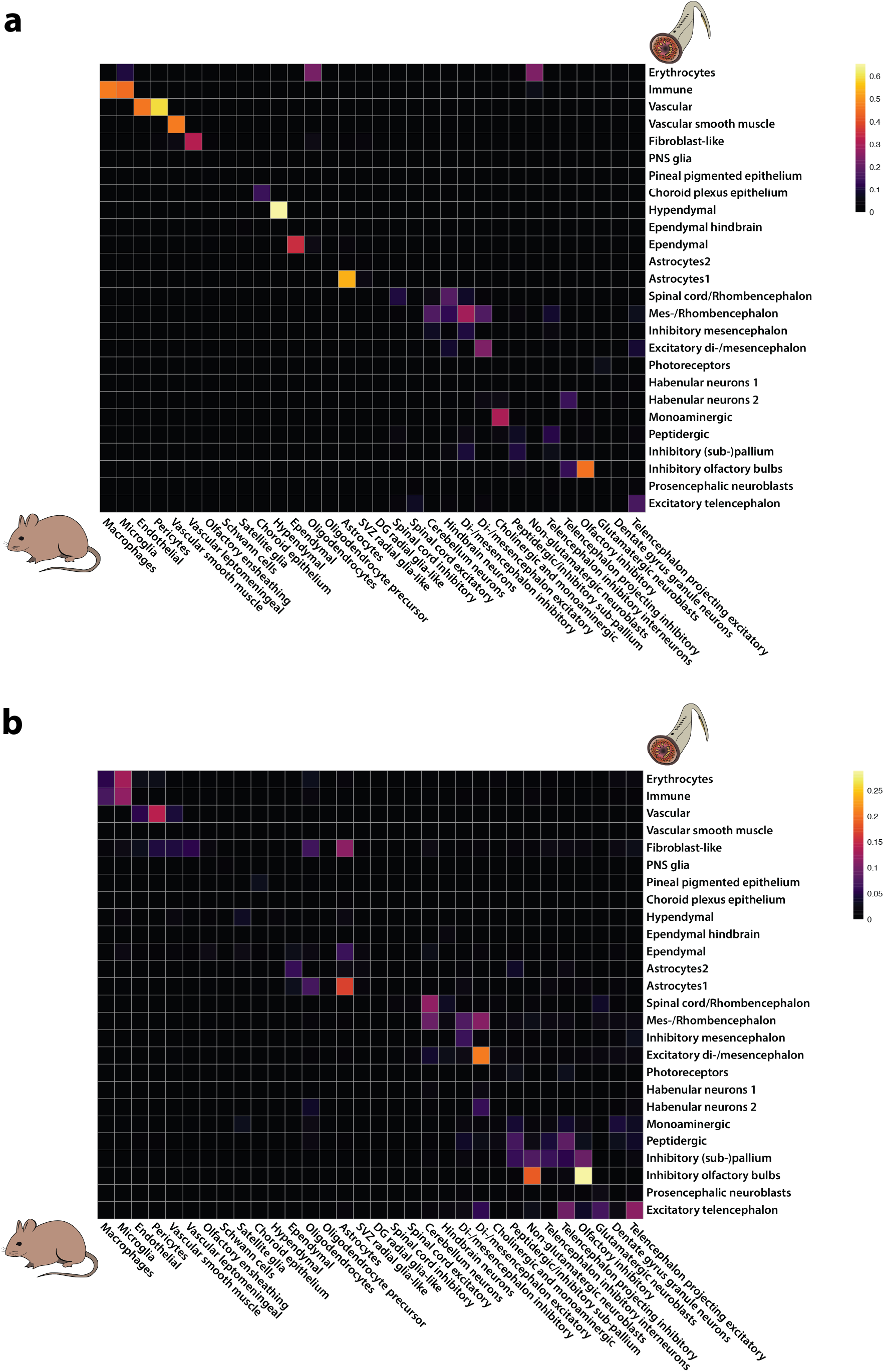
SAMap scores for all cell type groups. **a, b**, Heatmaps of SAMap mapping scores for all groups of non-neuronal and neuronal cell types between mouse and lamprey, including all genes (a) and TF genes only (b).

**Extended Data Fig. 10.**
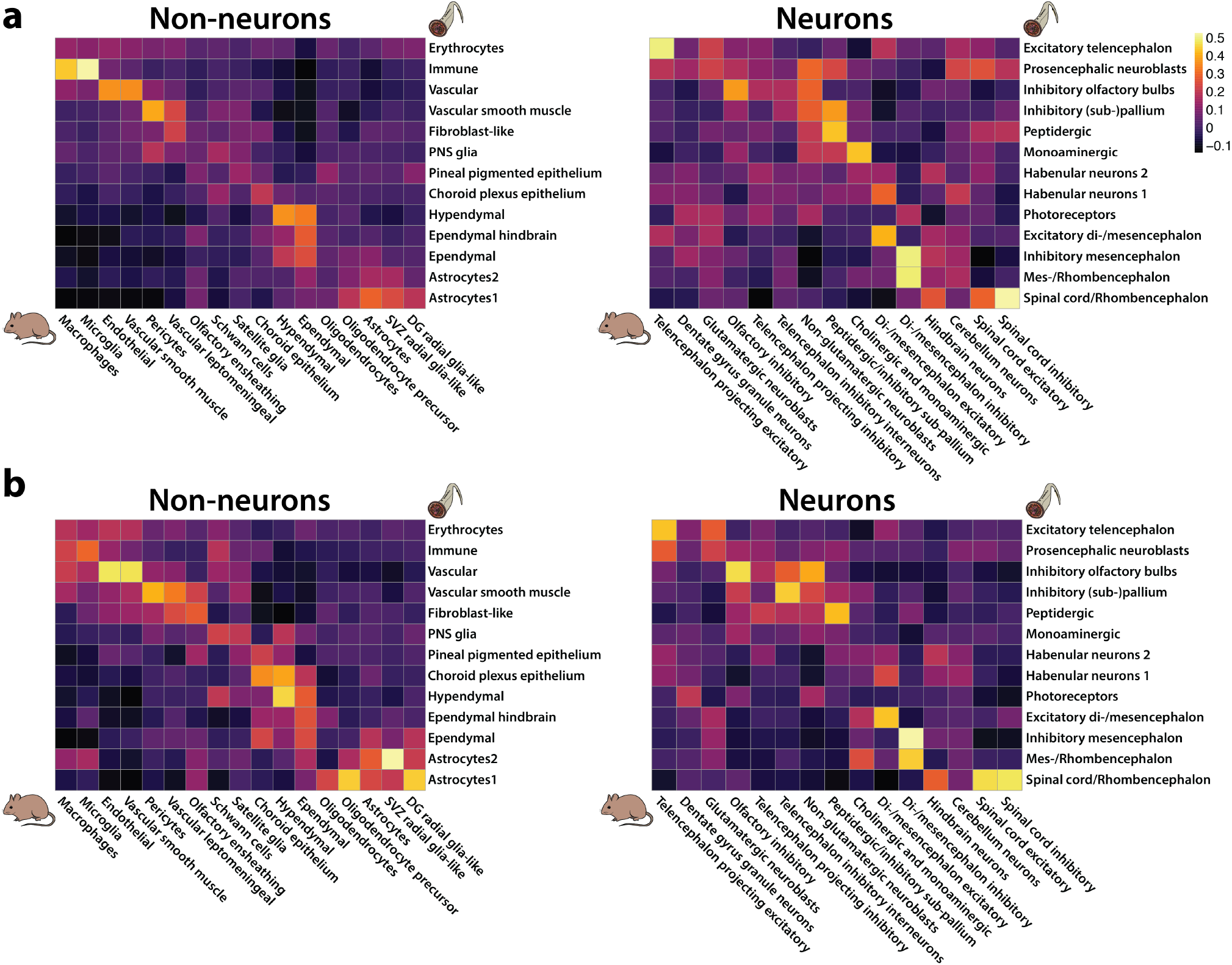
Correlations between cell type groups. **a, b**, Heatmaps showing Pearson’s correlation coefficients of specificity indexes of lamprey and mouse cell type groups for all orthologous genes (a) and for TFs only (b).

**Extended Data Fig. 11.**
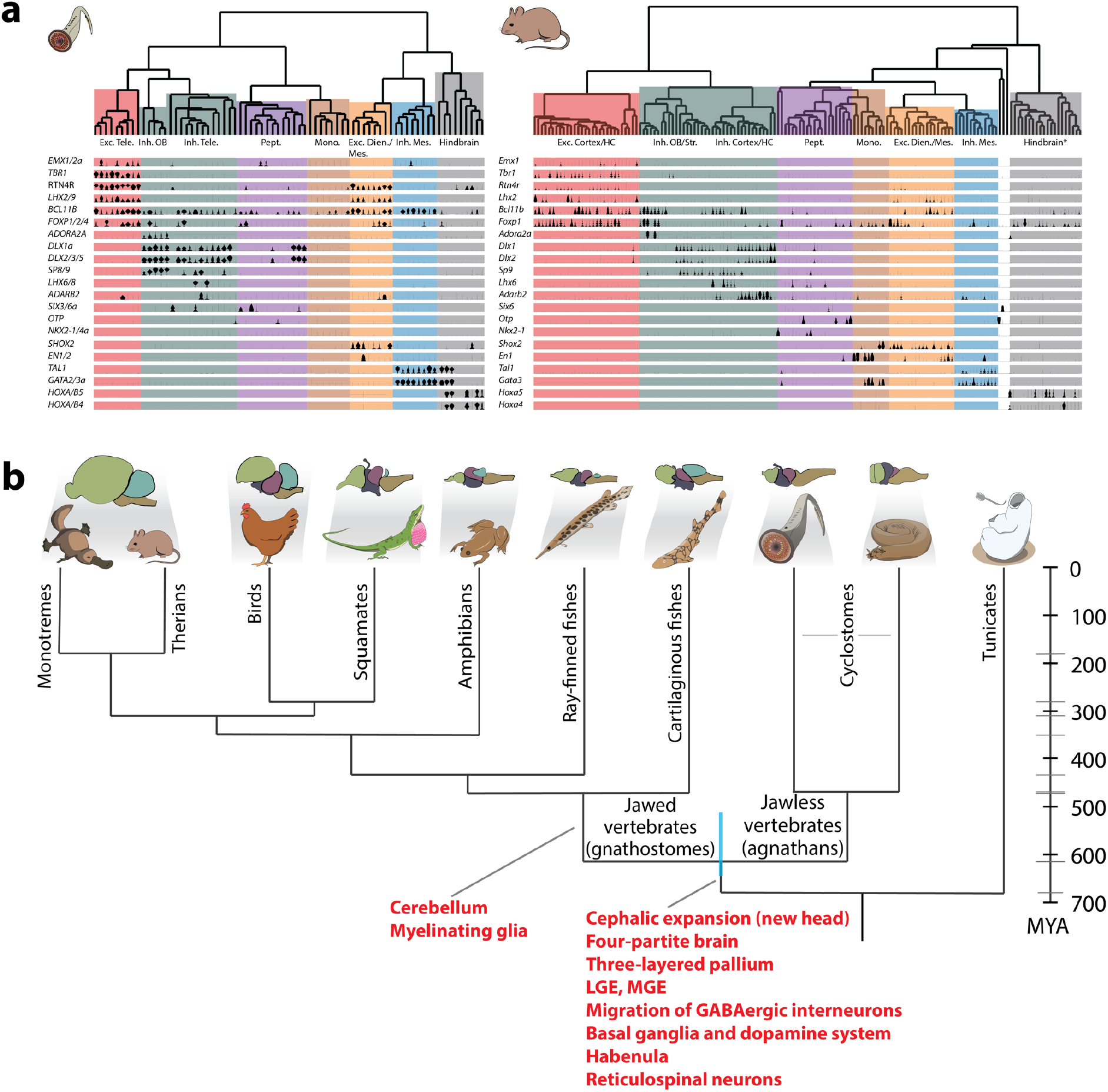
**a**, Upper panel: lamprey and mouse dendrograms of selected homologous neuronal families obtained based on correlations of expression levels of TF genes only. *Cerebellum excluded. Lower panel: violin plots showing the expression of selected TF genes for each cell type. Color code as in Fig. 1c. **b**, Vertebrate phylogenetic tree as in Fig. 1a showing key brain innovations (in red) as indicated, shown, or confirmed by our study.

**Extended Data Fig. 12.**
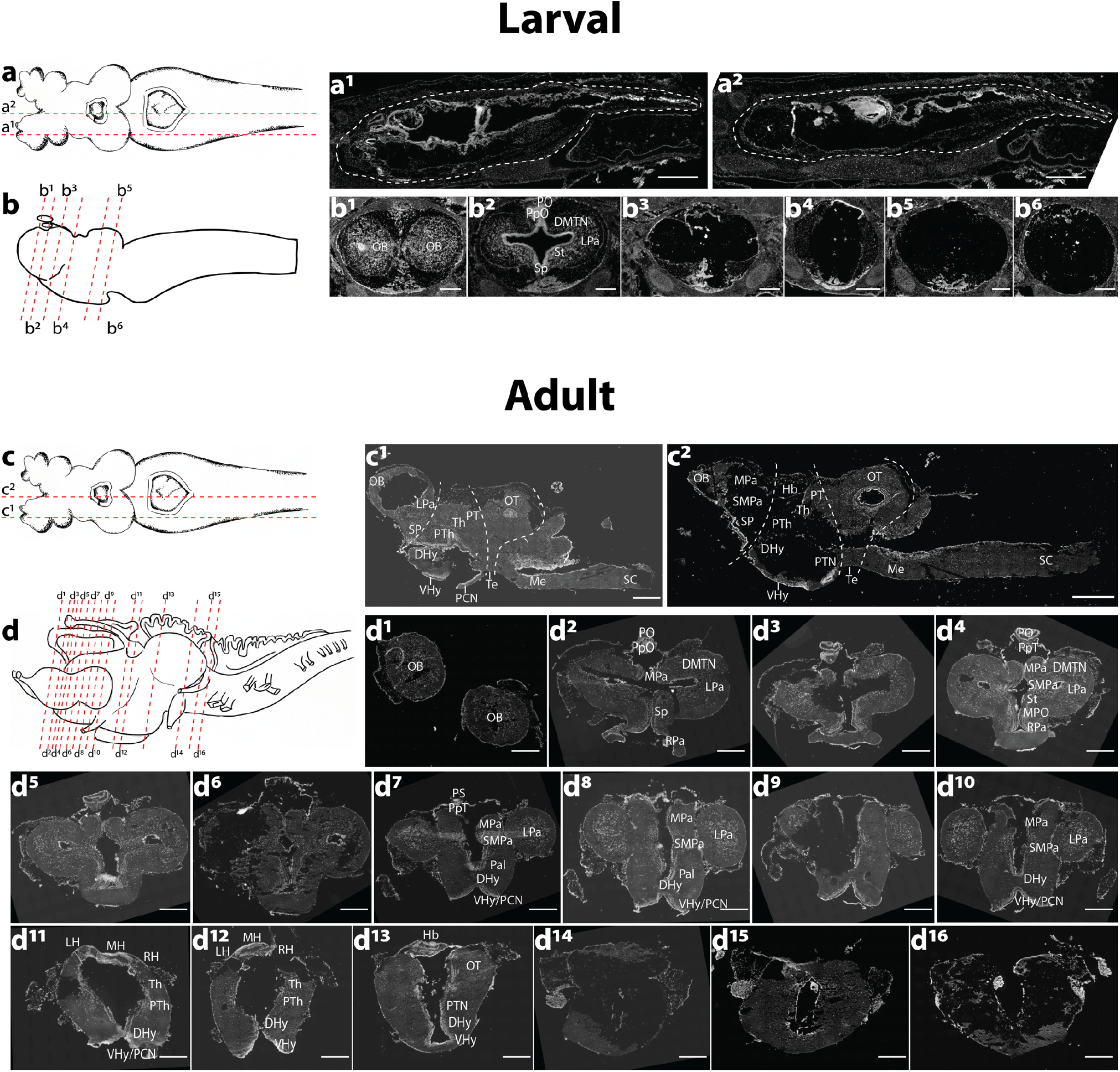
ISS dissection schemes. **a-d**, ISS dissection schemes (DAPI) of sagittal (a, c) and coronal (b, d) sections of lamprey larval and adult brains. Dashed lines on adult sagittal sections separate the main brain regions. DHy, dorsal hypothalamus; DMTN, dorsomedial telencephalic nucleus; Hb, habenula; LH, left habenula; LPa, lateral pallium; Me, medulla; MH, medial habenula; MPa, medial pallium; MPO, medial preoptic nucleus; OB, olfactory bulb; OT, optic tectum; Pal, pallidum; PCN, postoptic commissure nucleus; PO, pineal organ; PpO, parapineal organ; PpT, parapineal tract; PS, pineal stalk; PT, pre-tectum; PTh, pre-thalamus; PTN, posterior tubercle nucleus; RH, right habenula; RPa, rostral paraventricular area; SC, spinal cord; SMPa, sub-medial pallium; Sp, septum; St, striatum; Te, tegmentum; Th, thalamus; VHy, ventral hypothalamus. Scale bars, 500µm.

