## Supplementary figures and images for "Reconstructing the ancestral vertebrate brain using a lamprey neural cell type atlas"

### Hulu_sspo.jpg

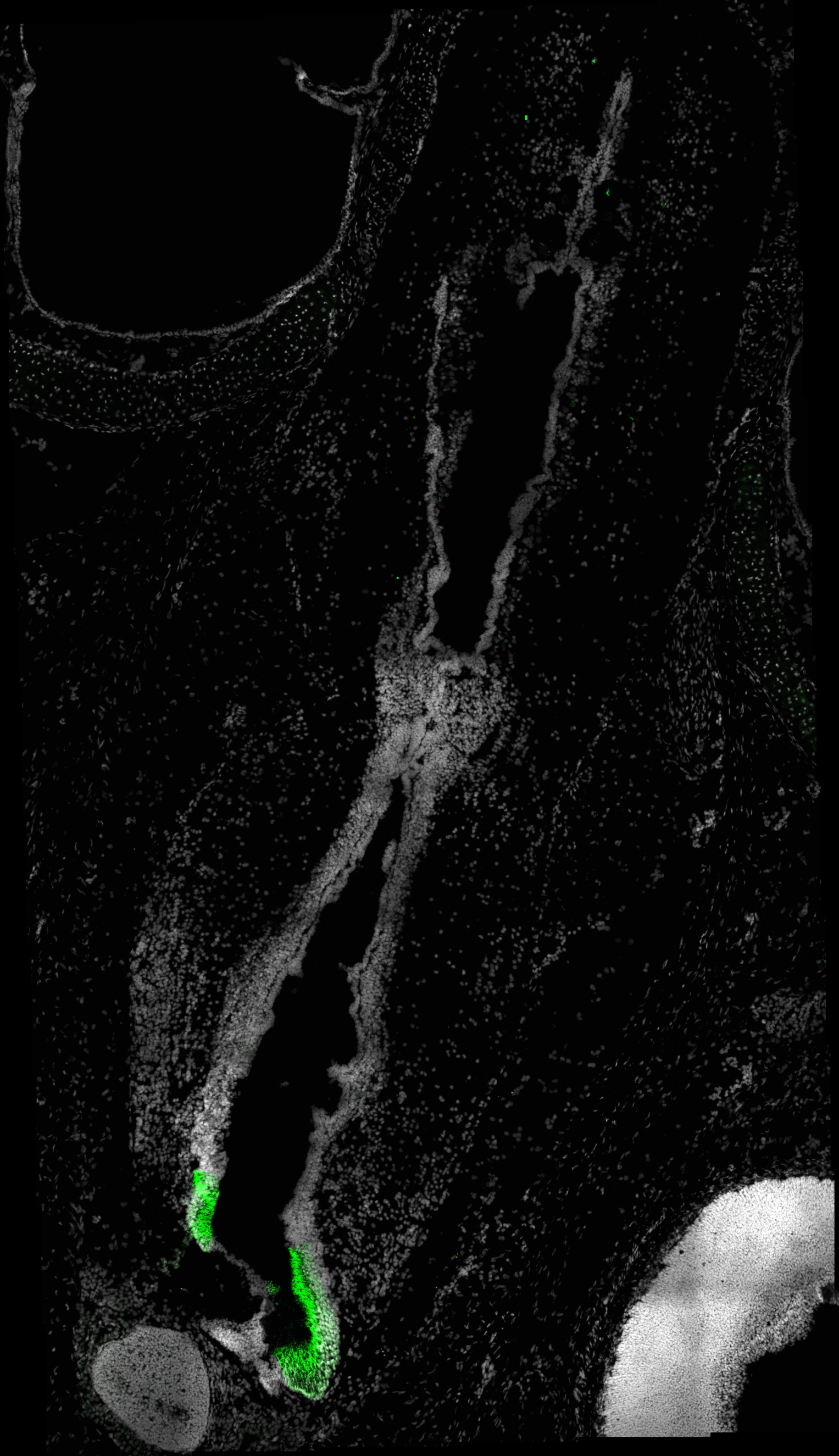

### Hulu_vat_gnrh1a.jpg

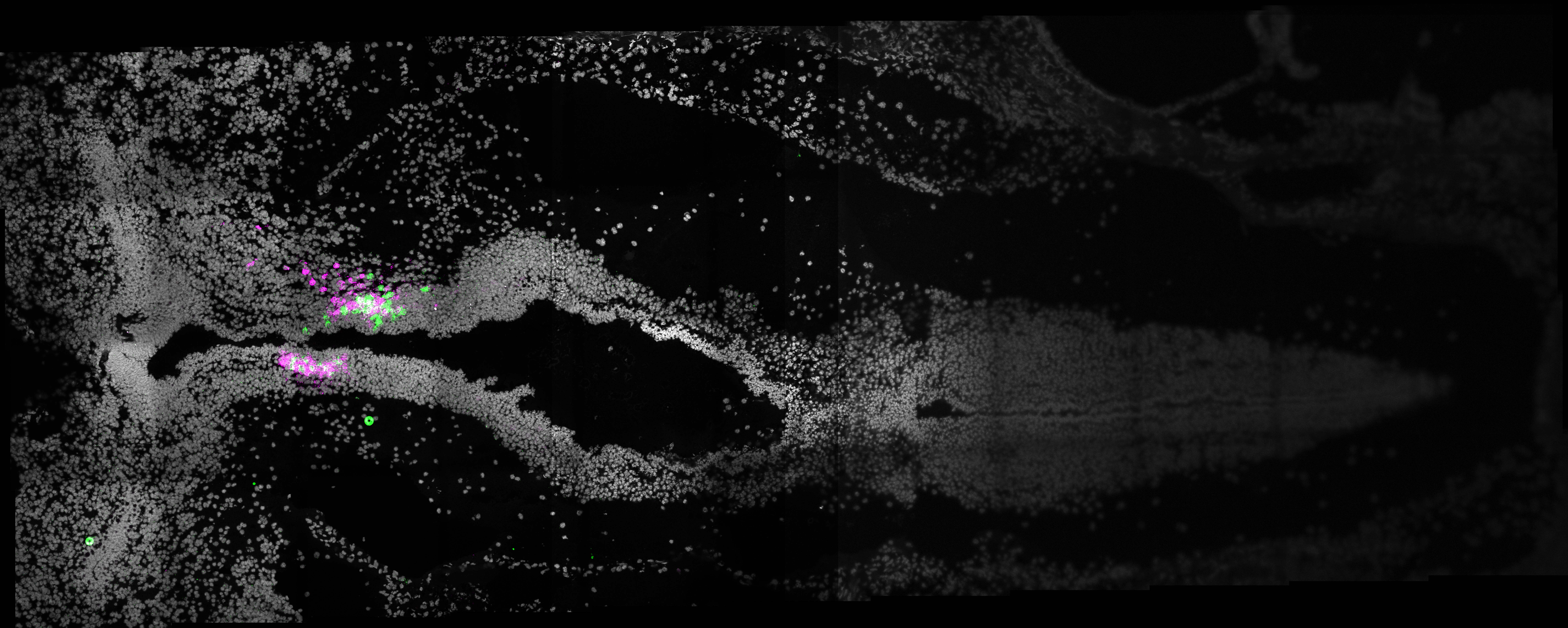

### Hulu_zfp704.jpg

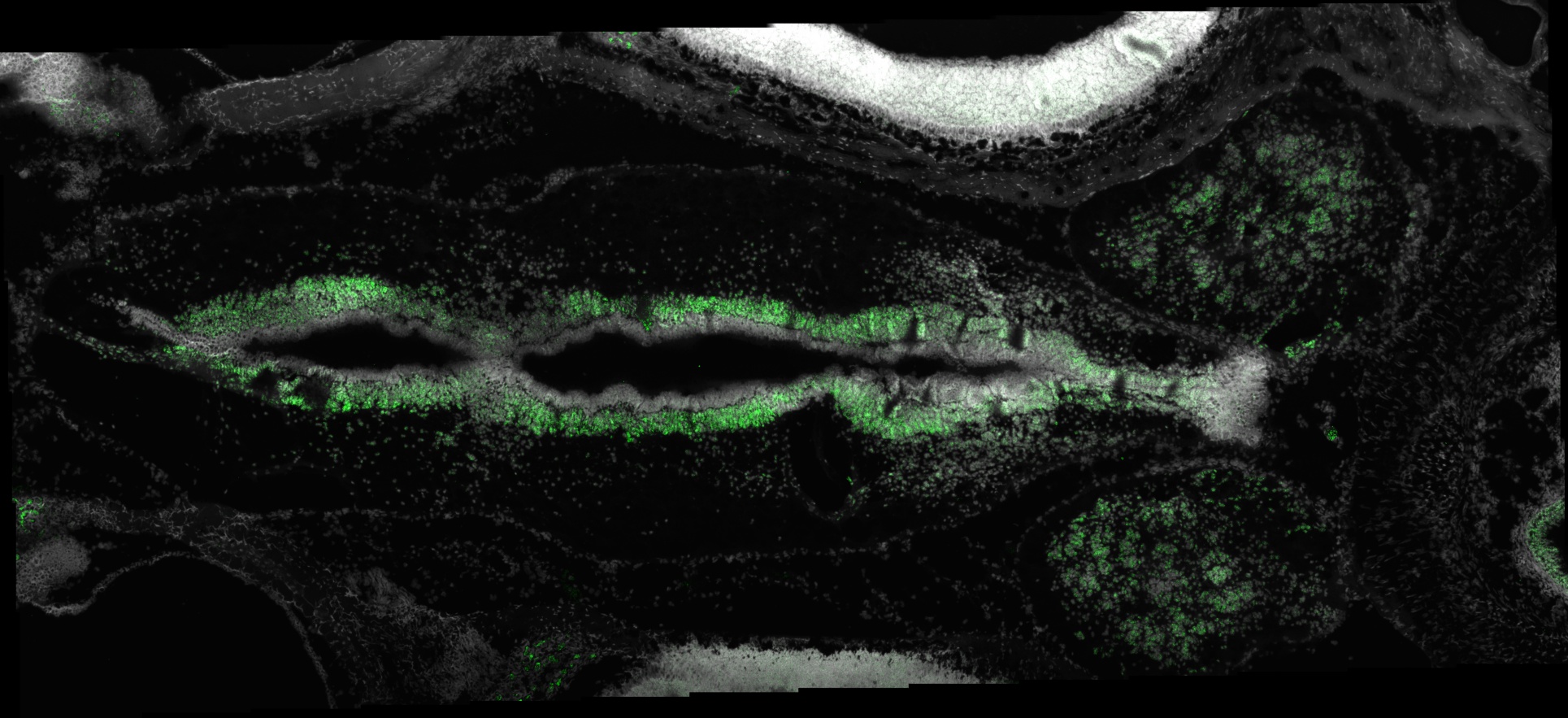

### MSTRG.149.jpg

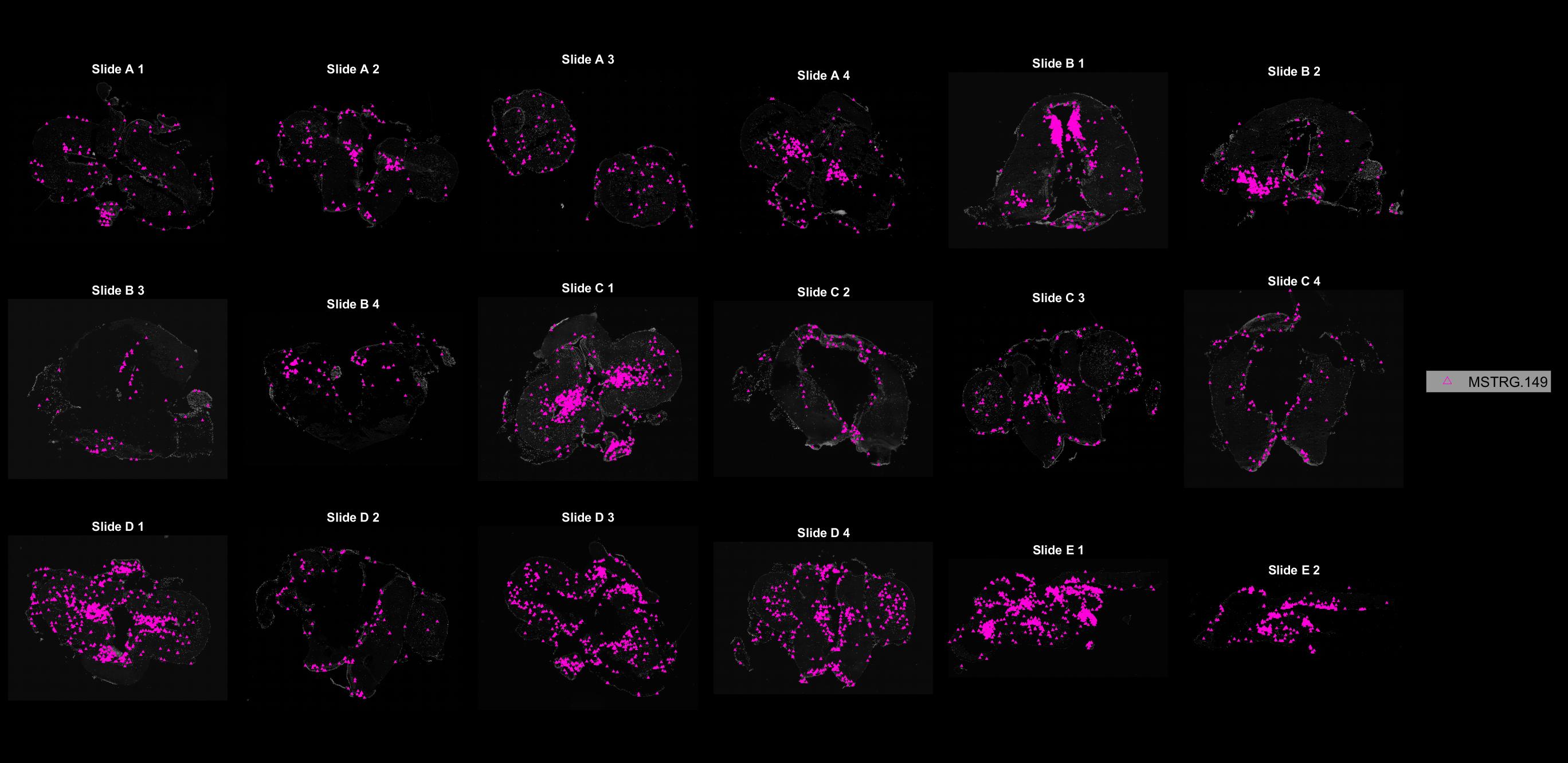

### MSTRG.322.jpg

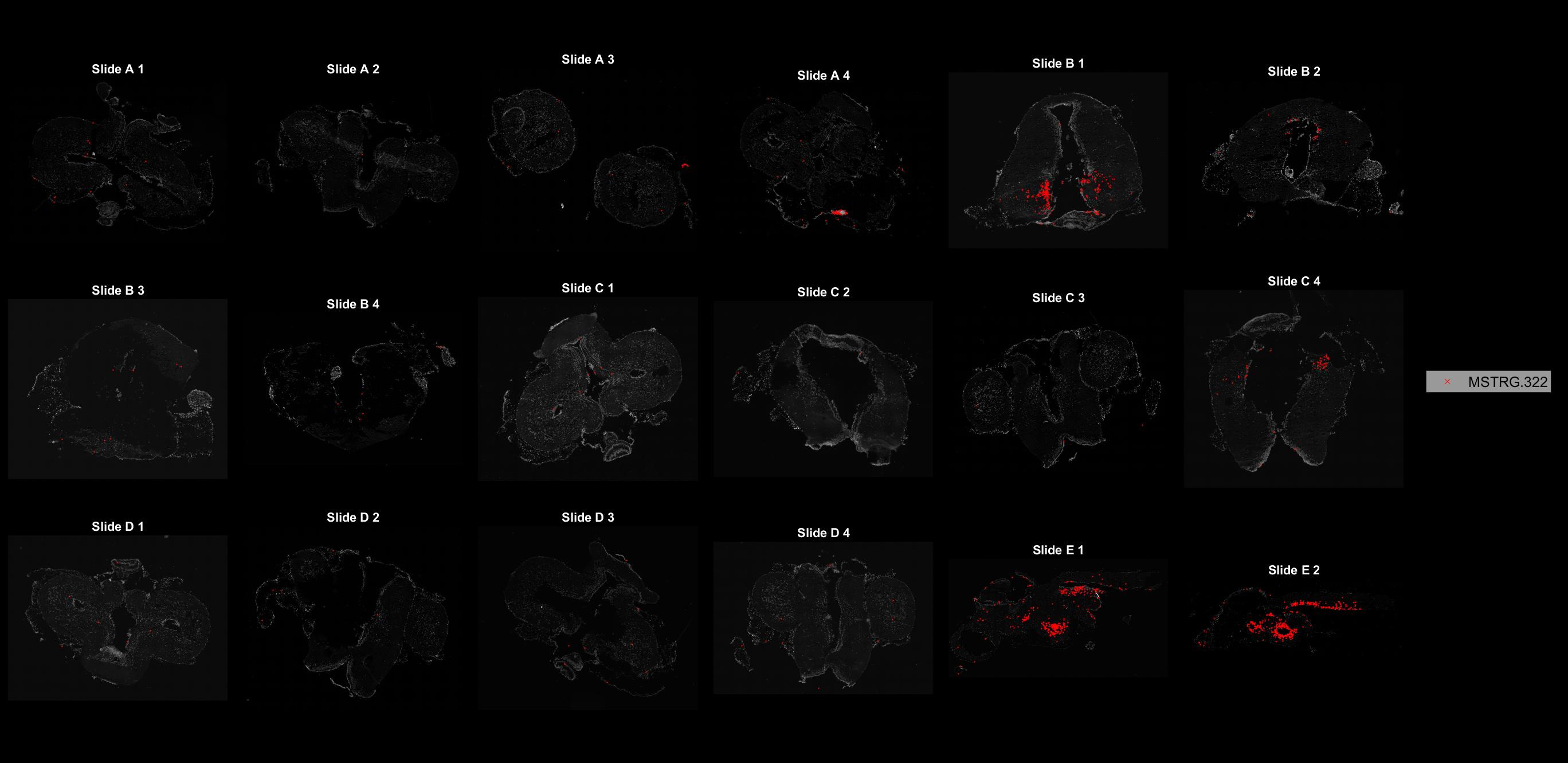

### MSTRG.419.jpg

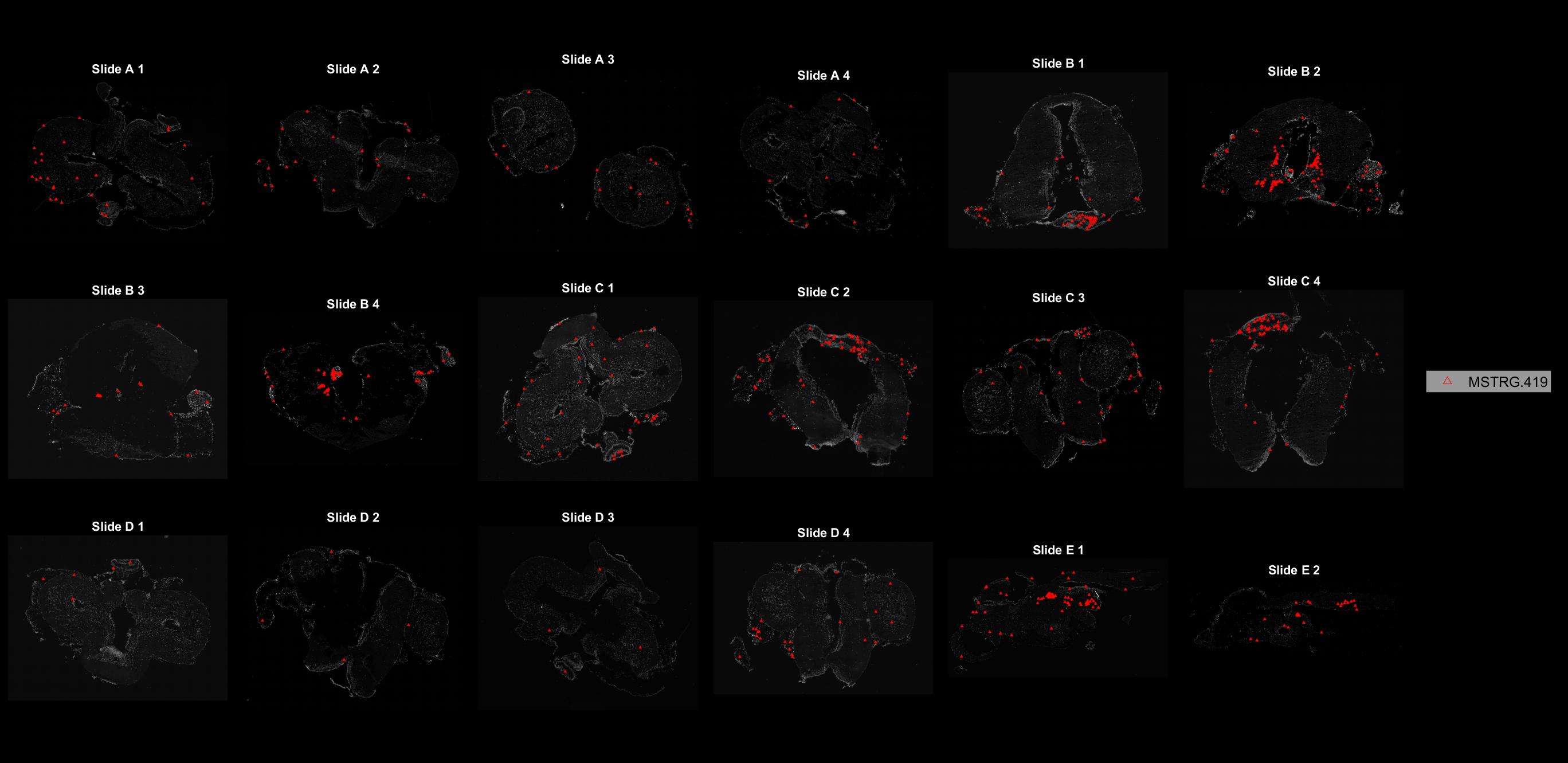

### MSTRG.616.jpg

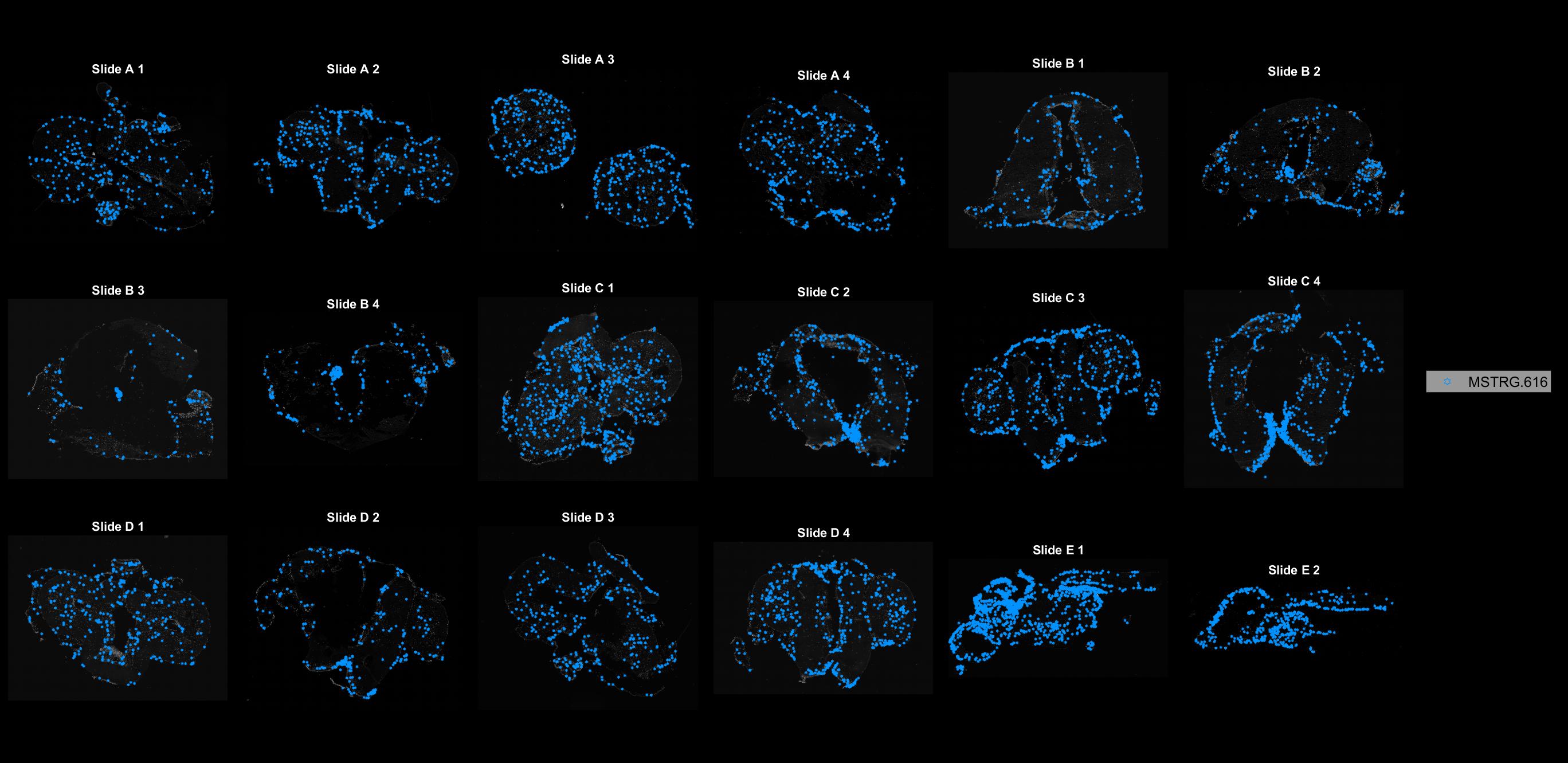

### MSTRG.621.jpg

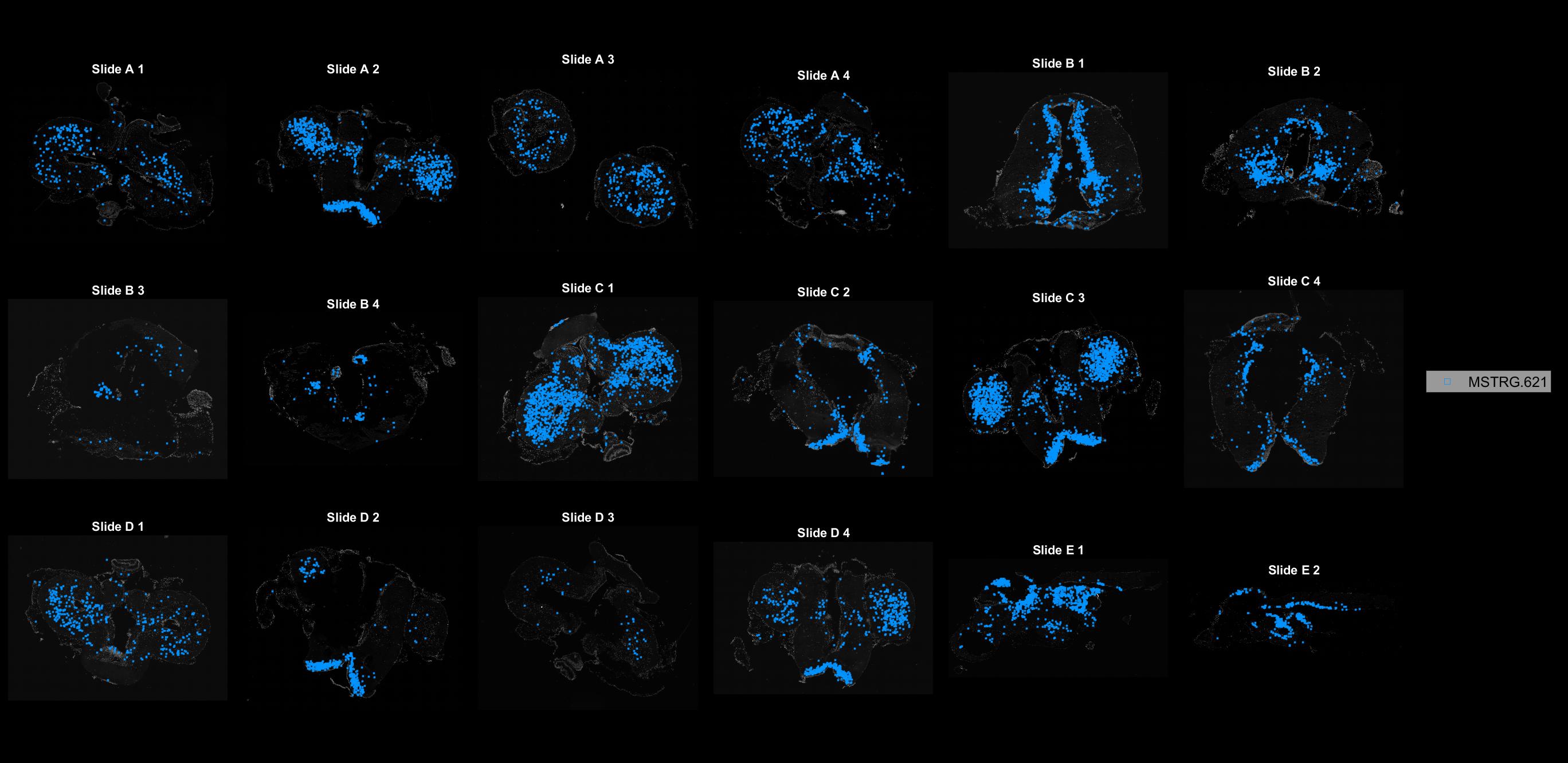

### MSTRG.850.jpg

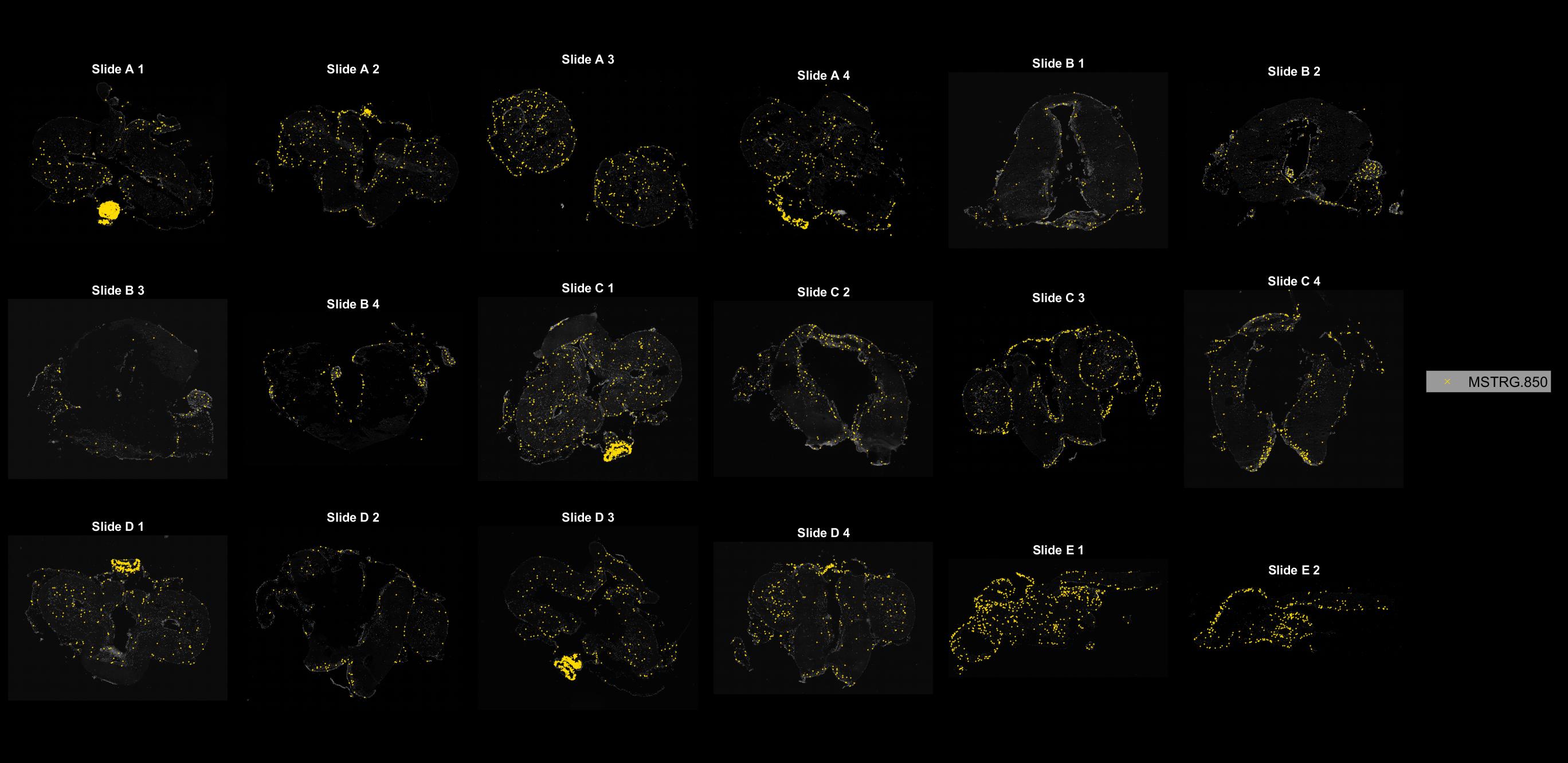

### MSTRG.857.jpg

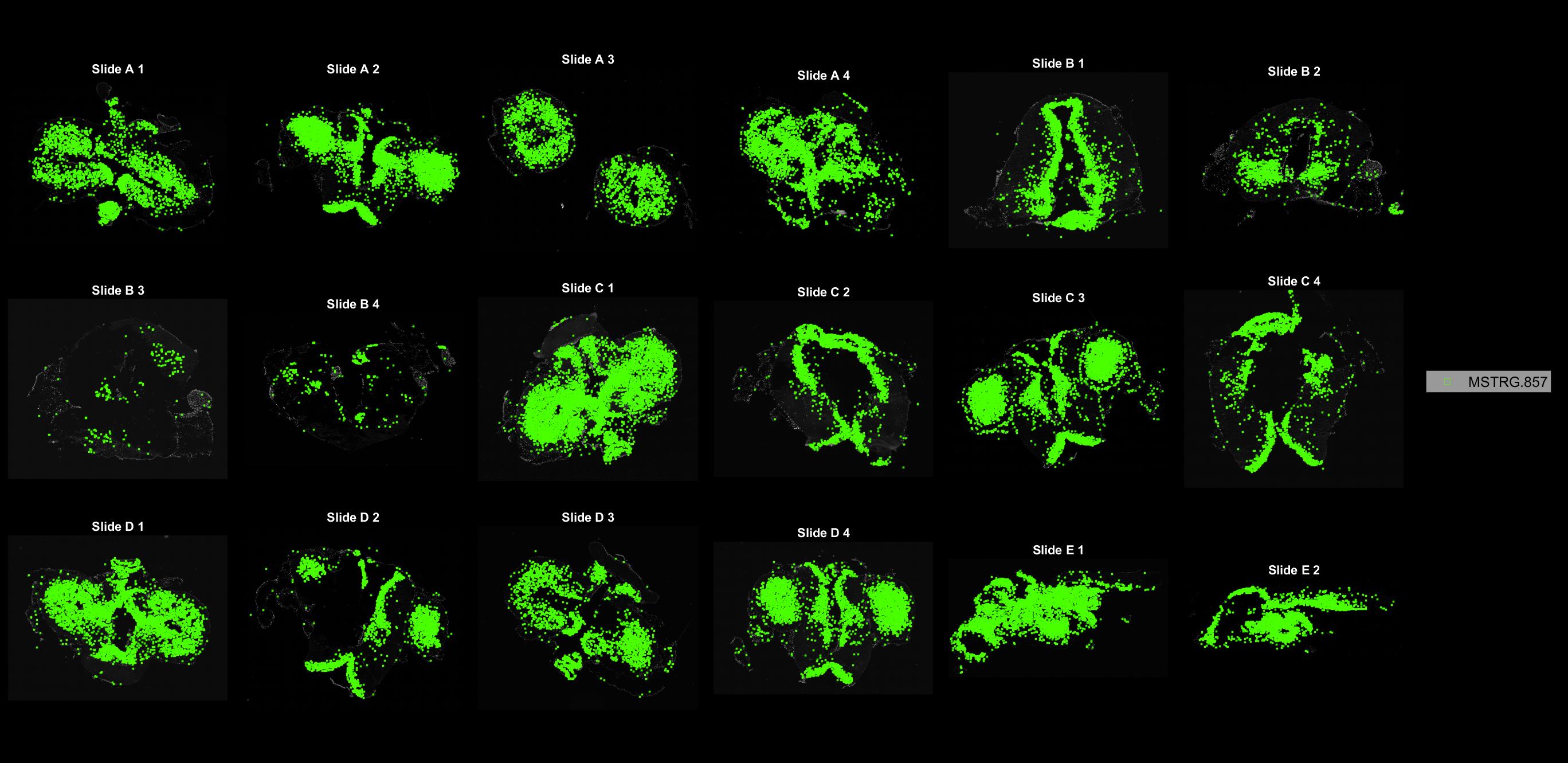

### MSTRG.911.jpg

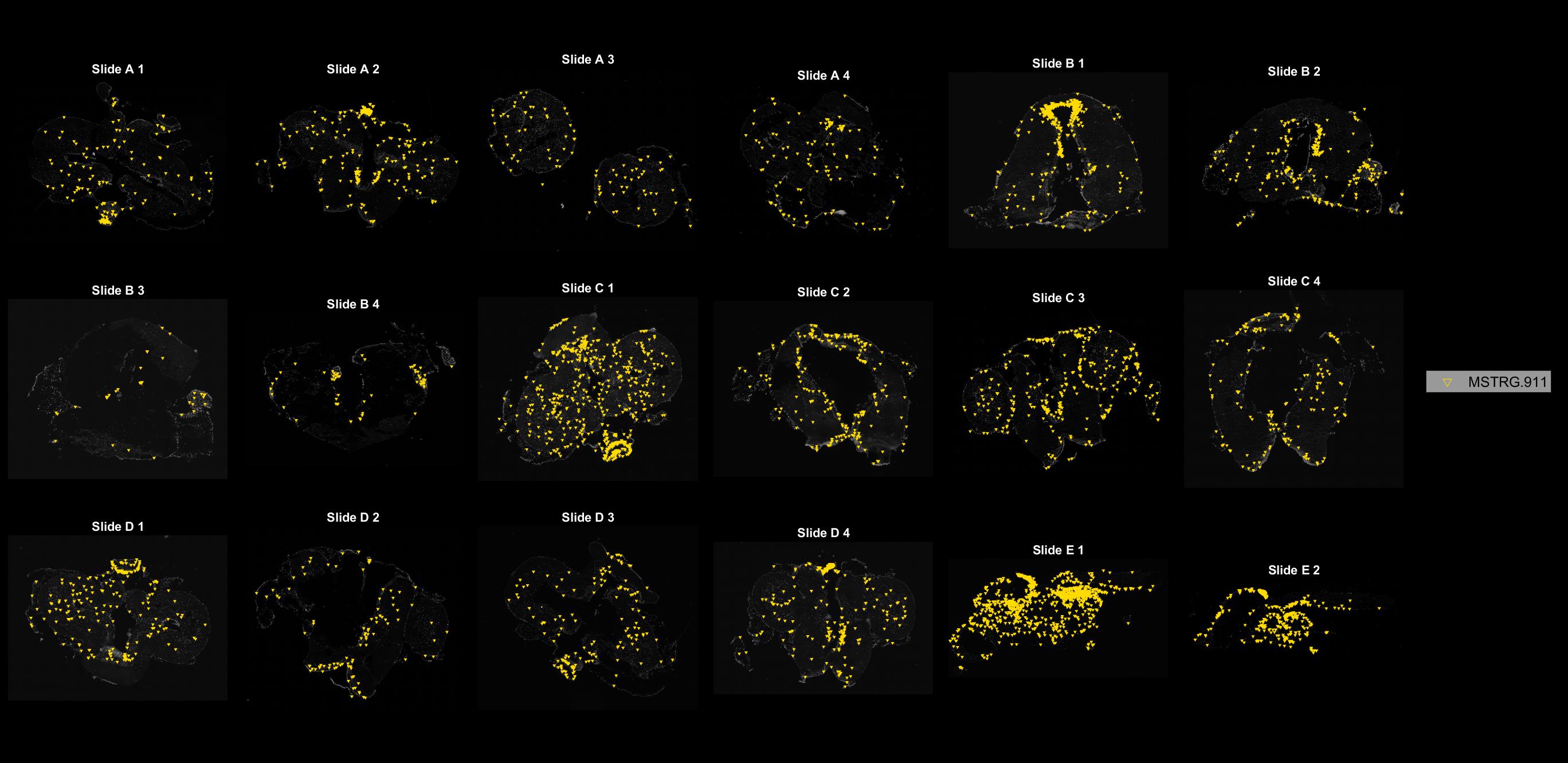

### MSTRG.939.jpg

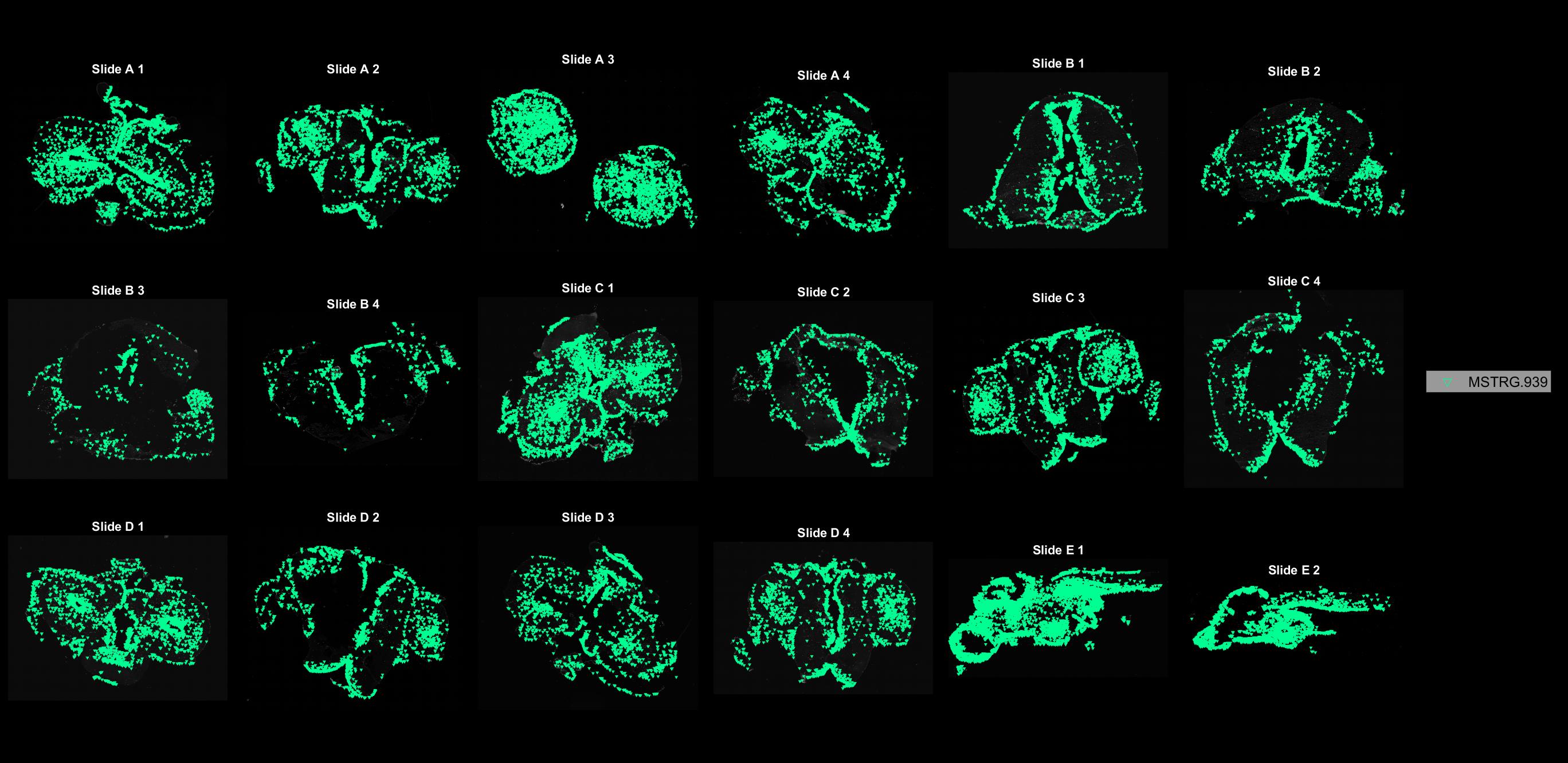

### MSTRG.1234.jpg

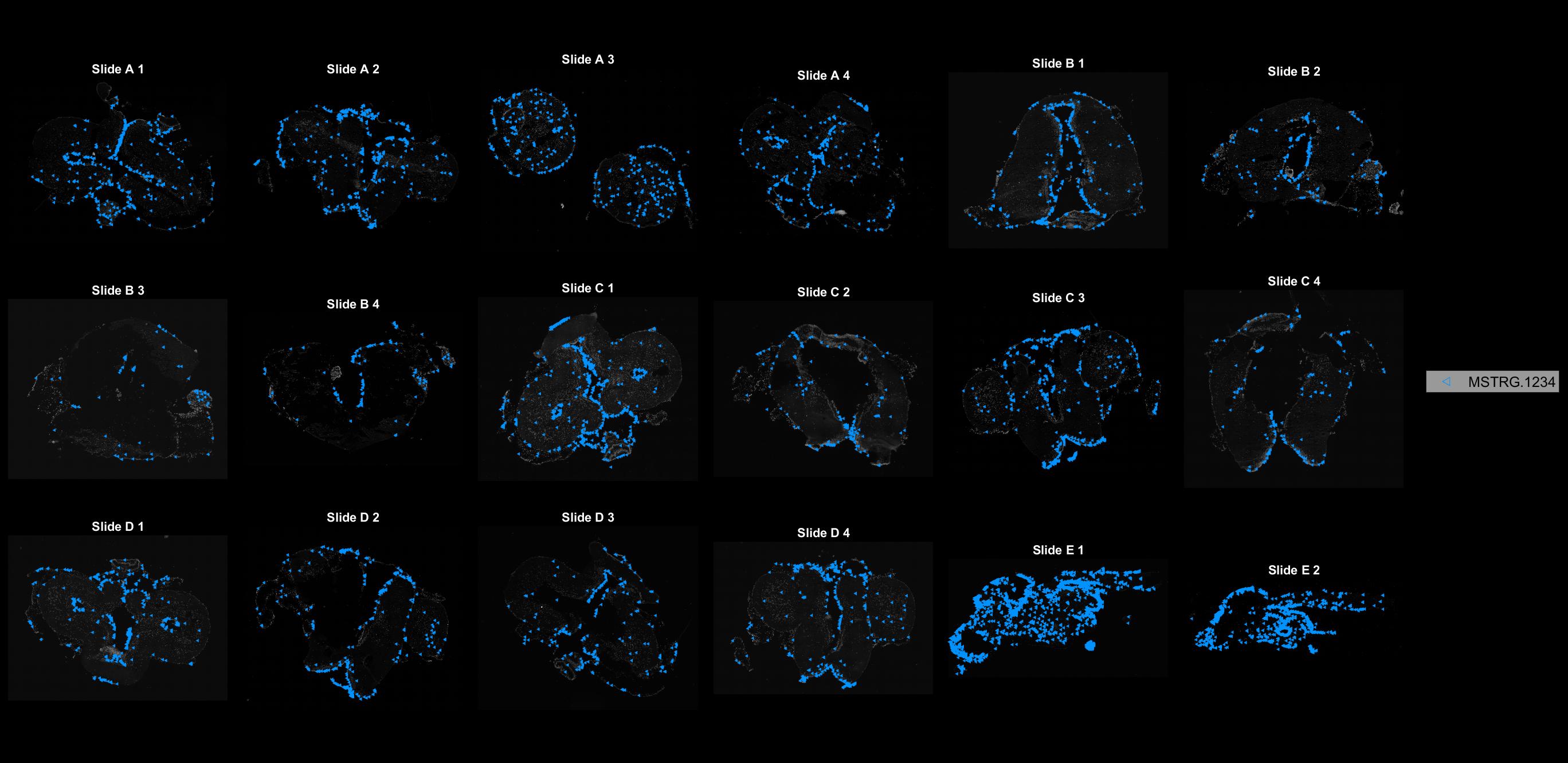

### MSTRG.1257.jpg

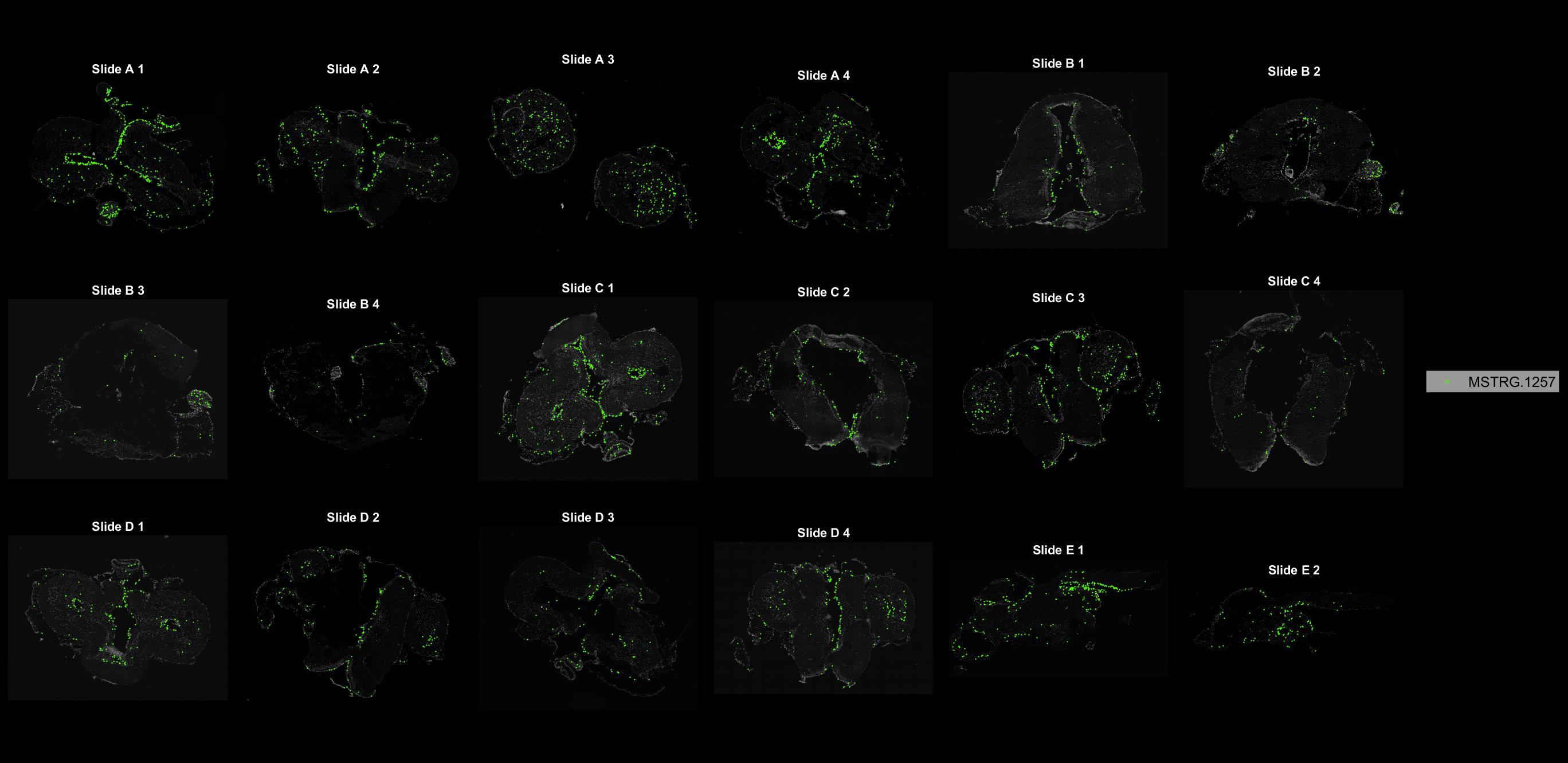

### MSTRG.1268.jpg

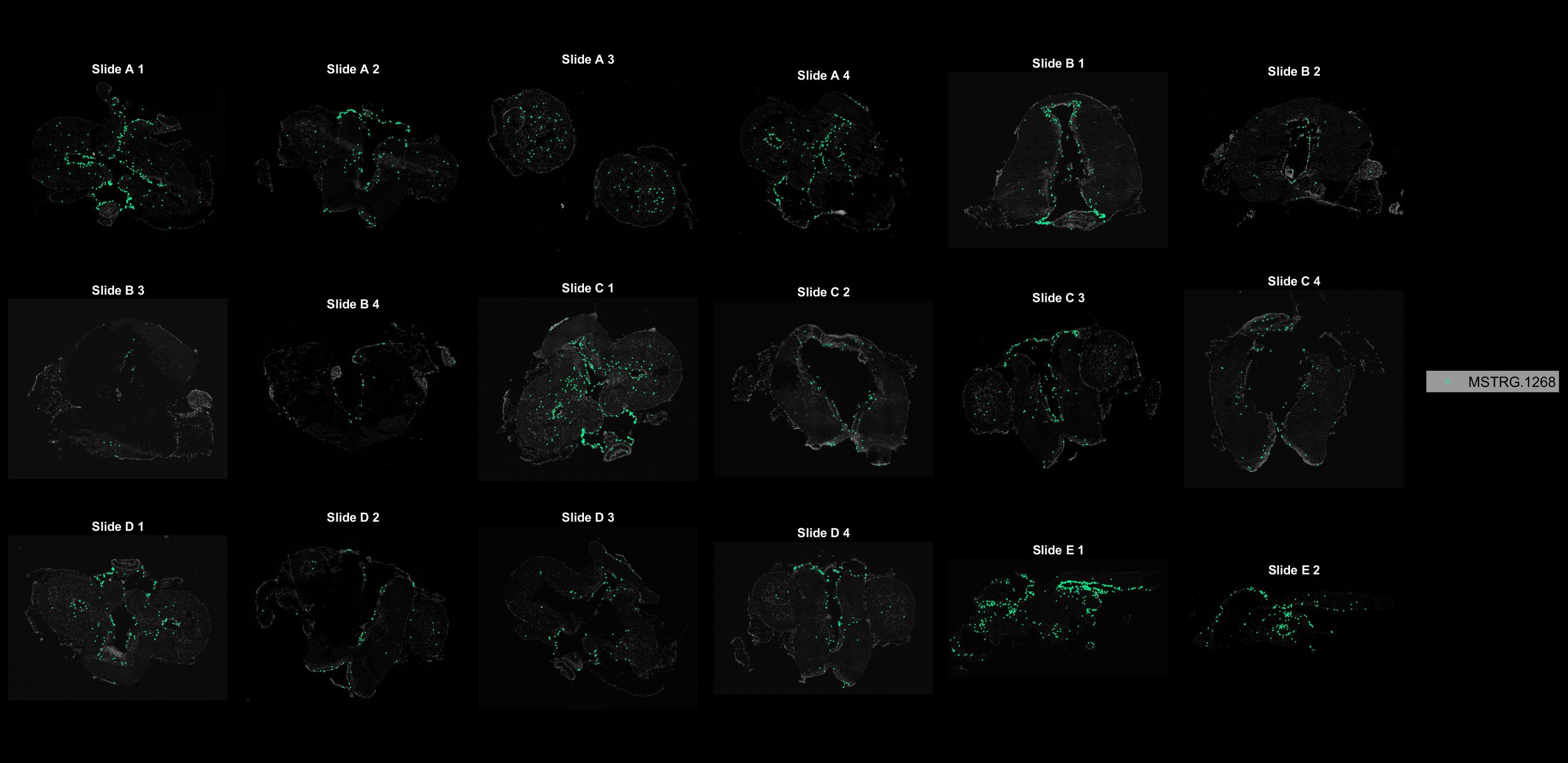

### MSTRG.1319.jpg

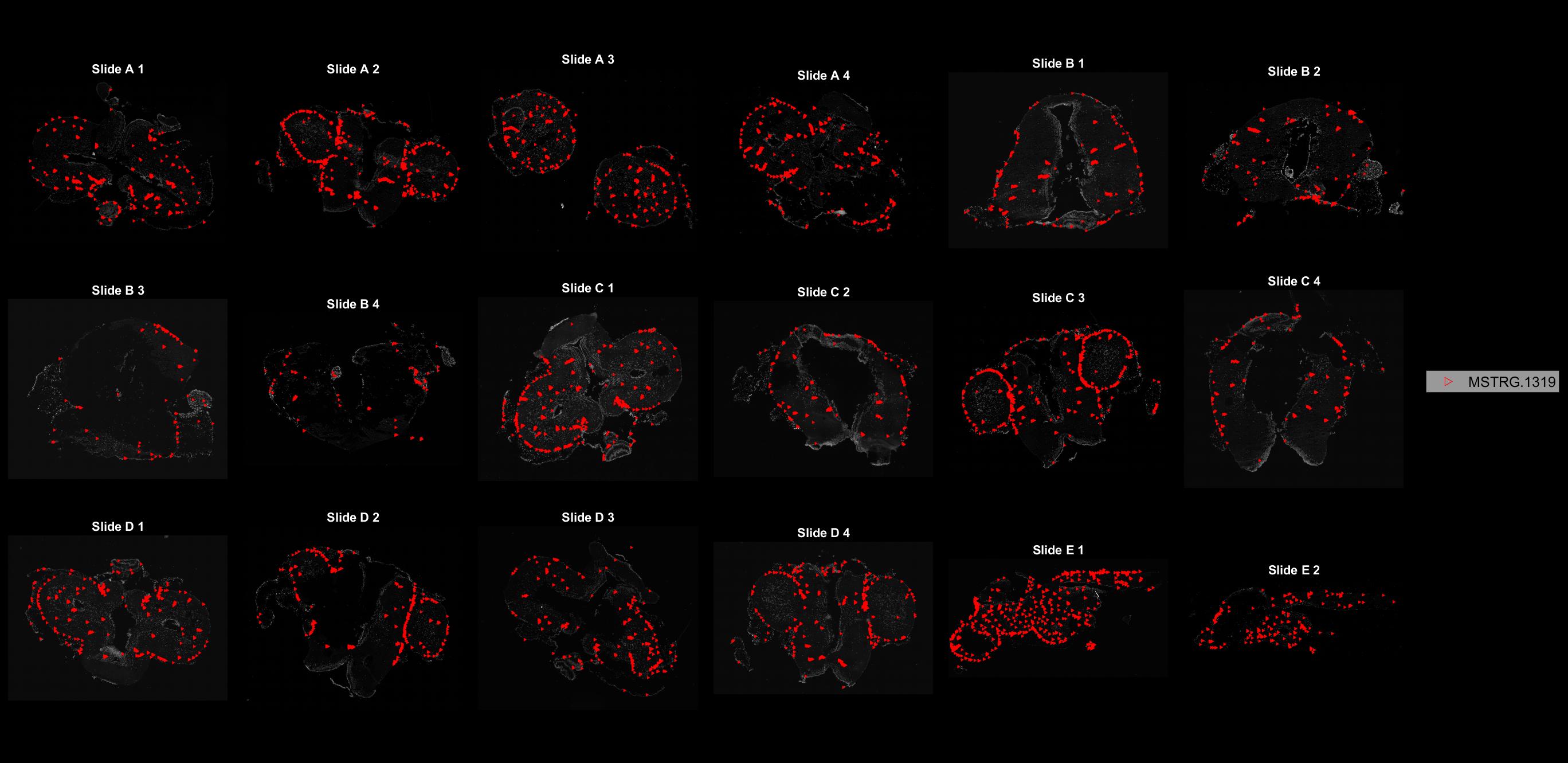

### MSTRG.1633.jpg

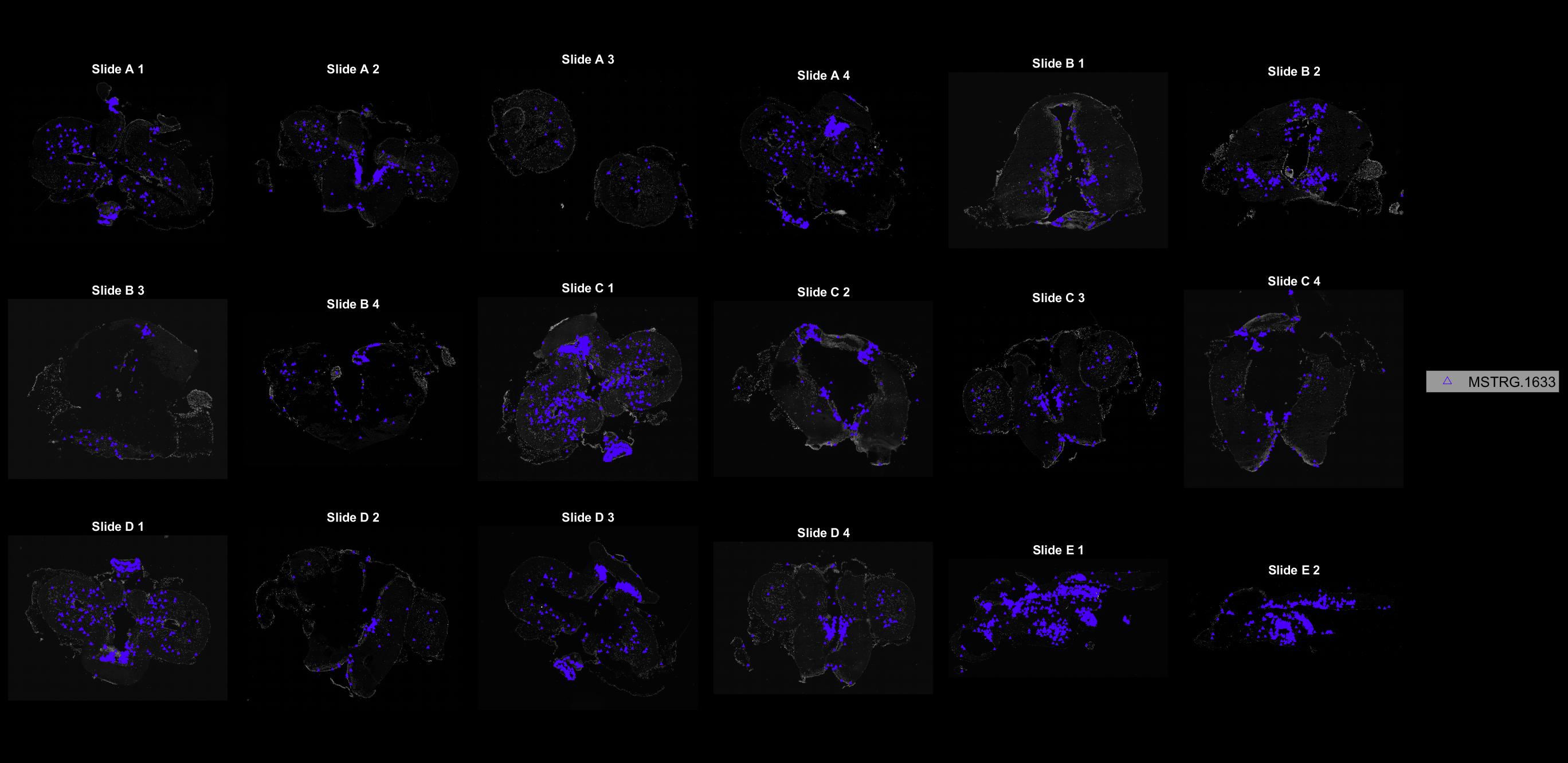

### MSTRG.1871.jpg

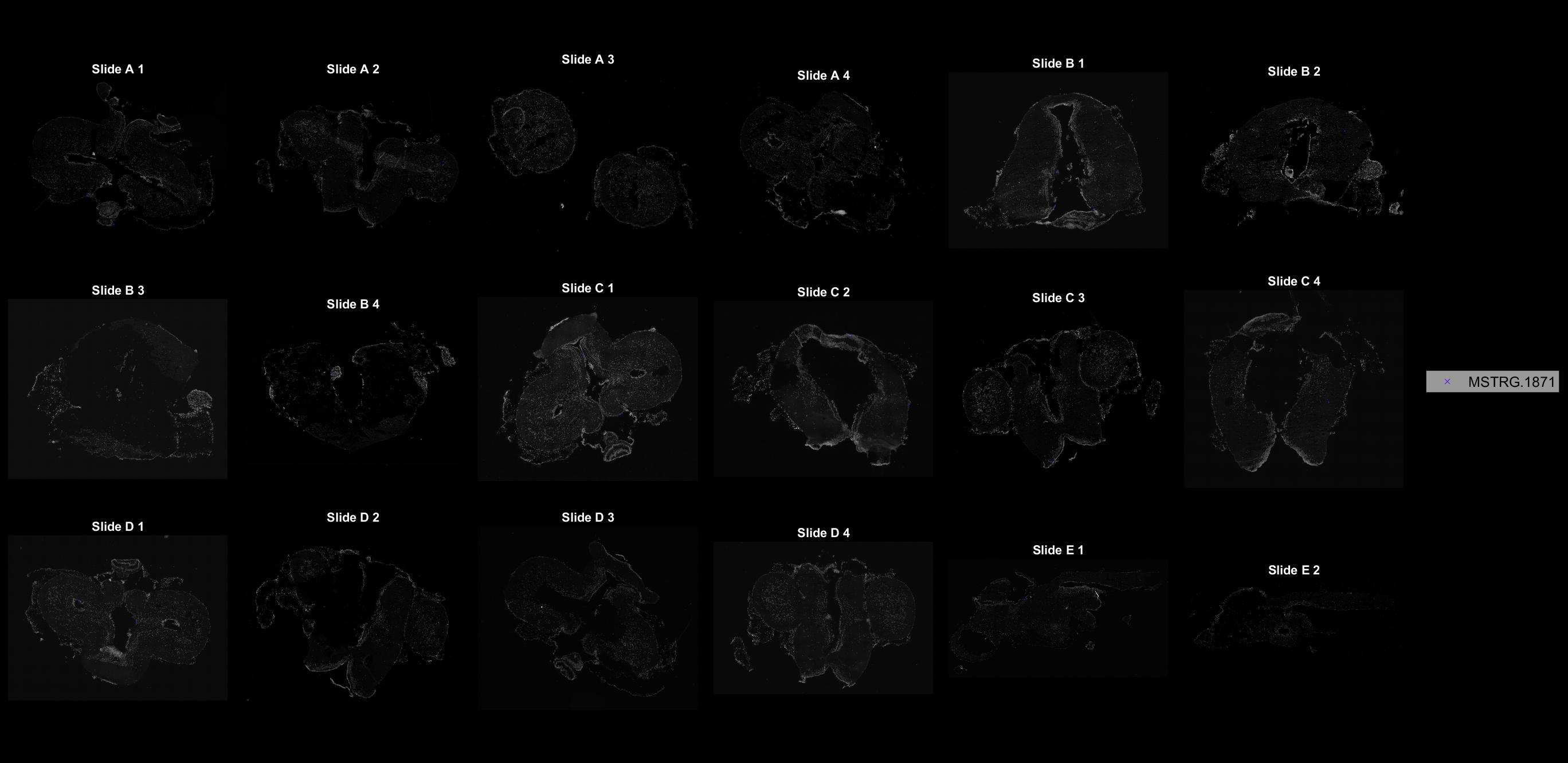

### MSTRG.2004.jpg

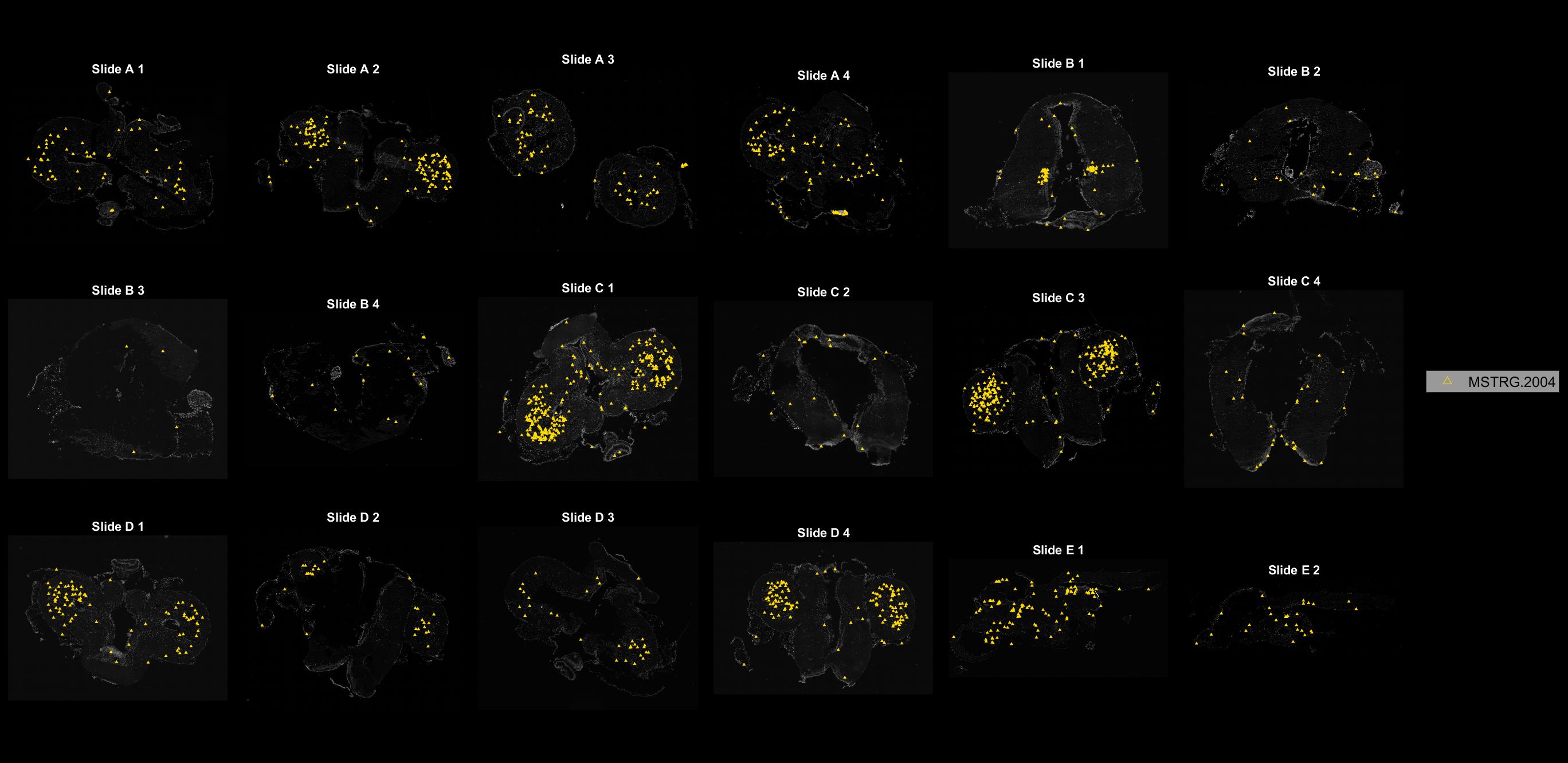

### MSTRG.2012.jpg

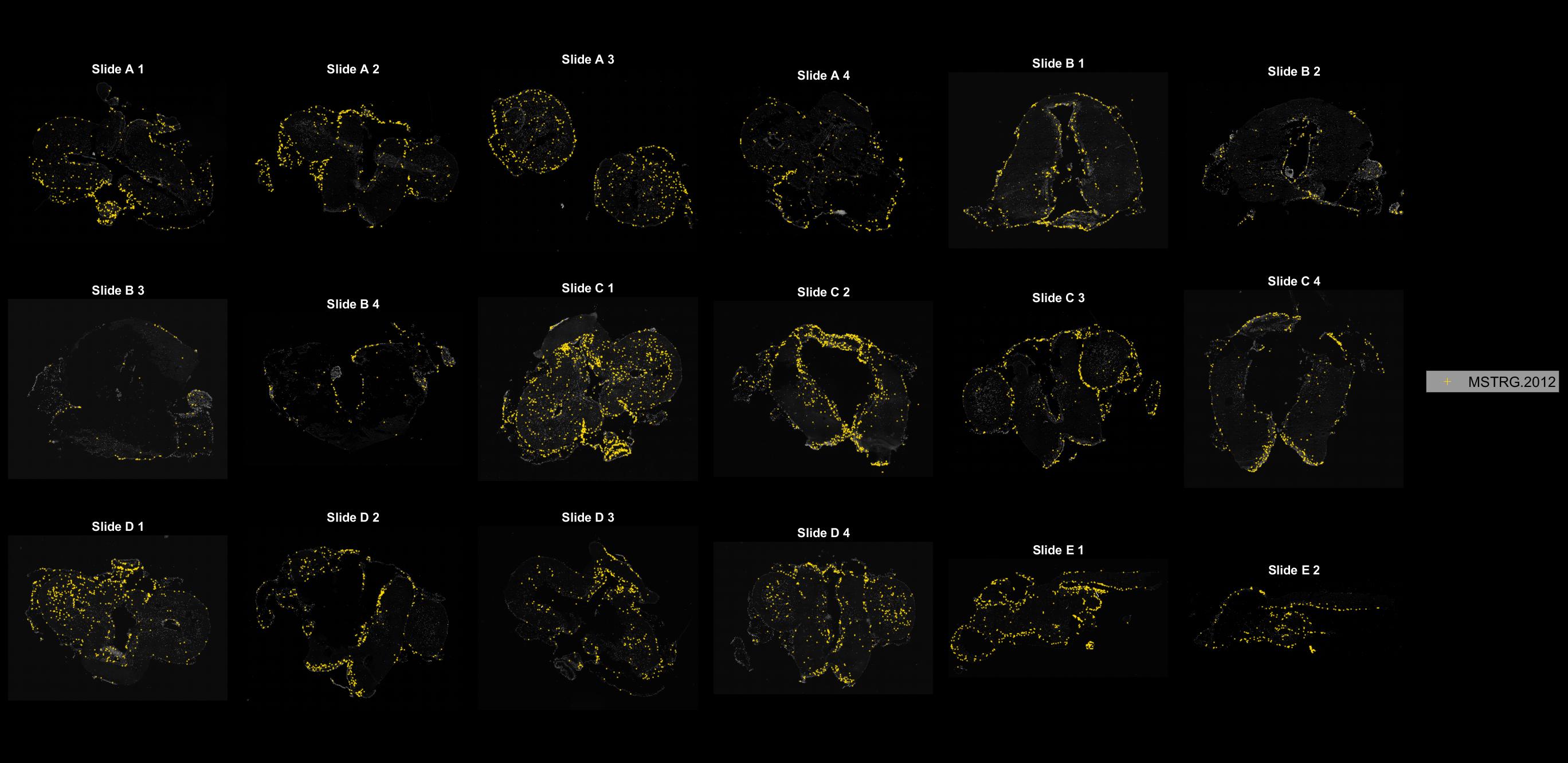

### MSTRG.2032.jpg

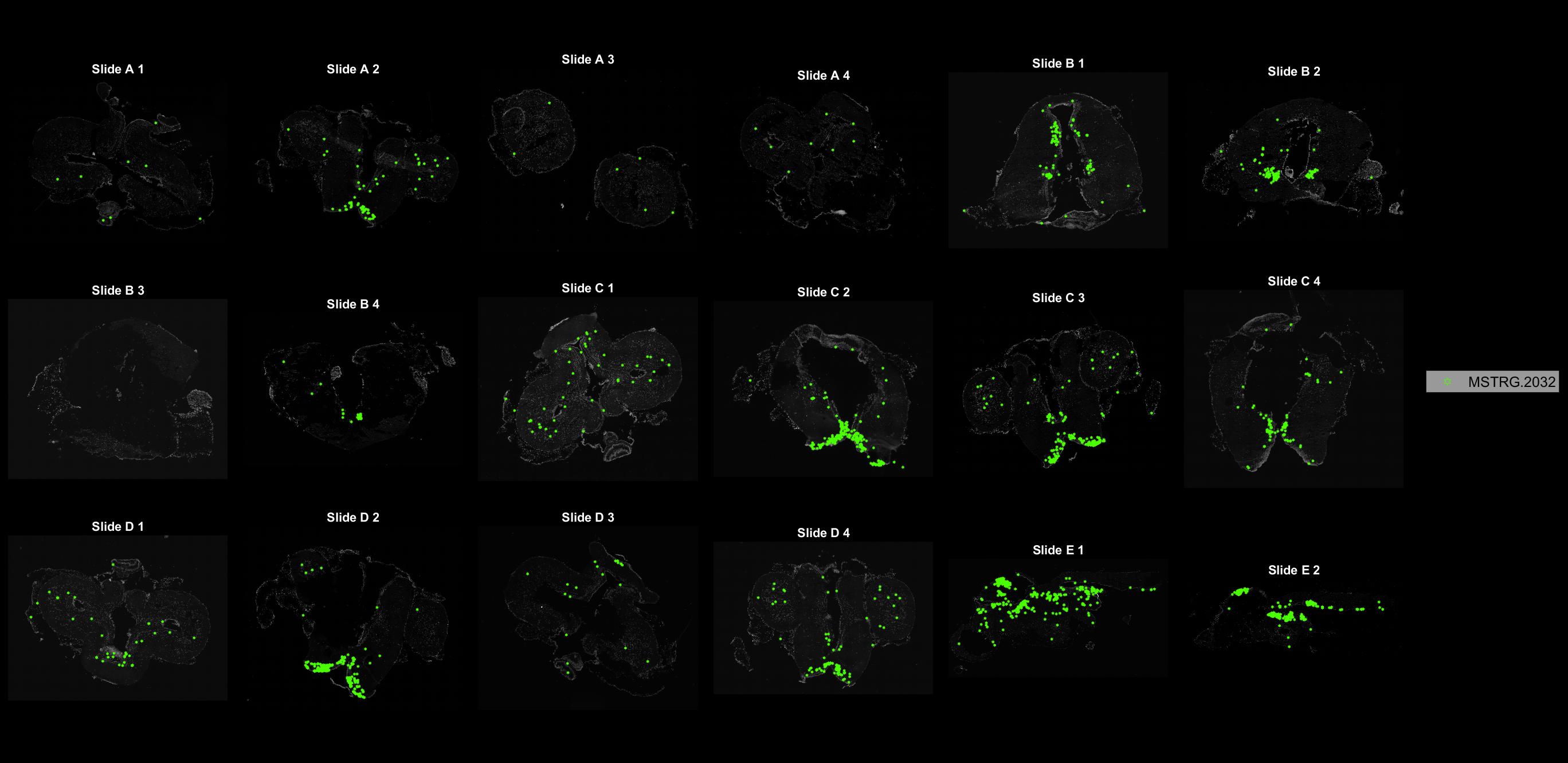

### MSTRG.2258.jpg

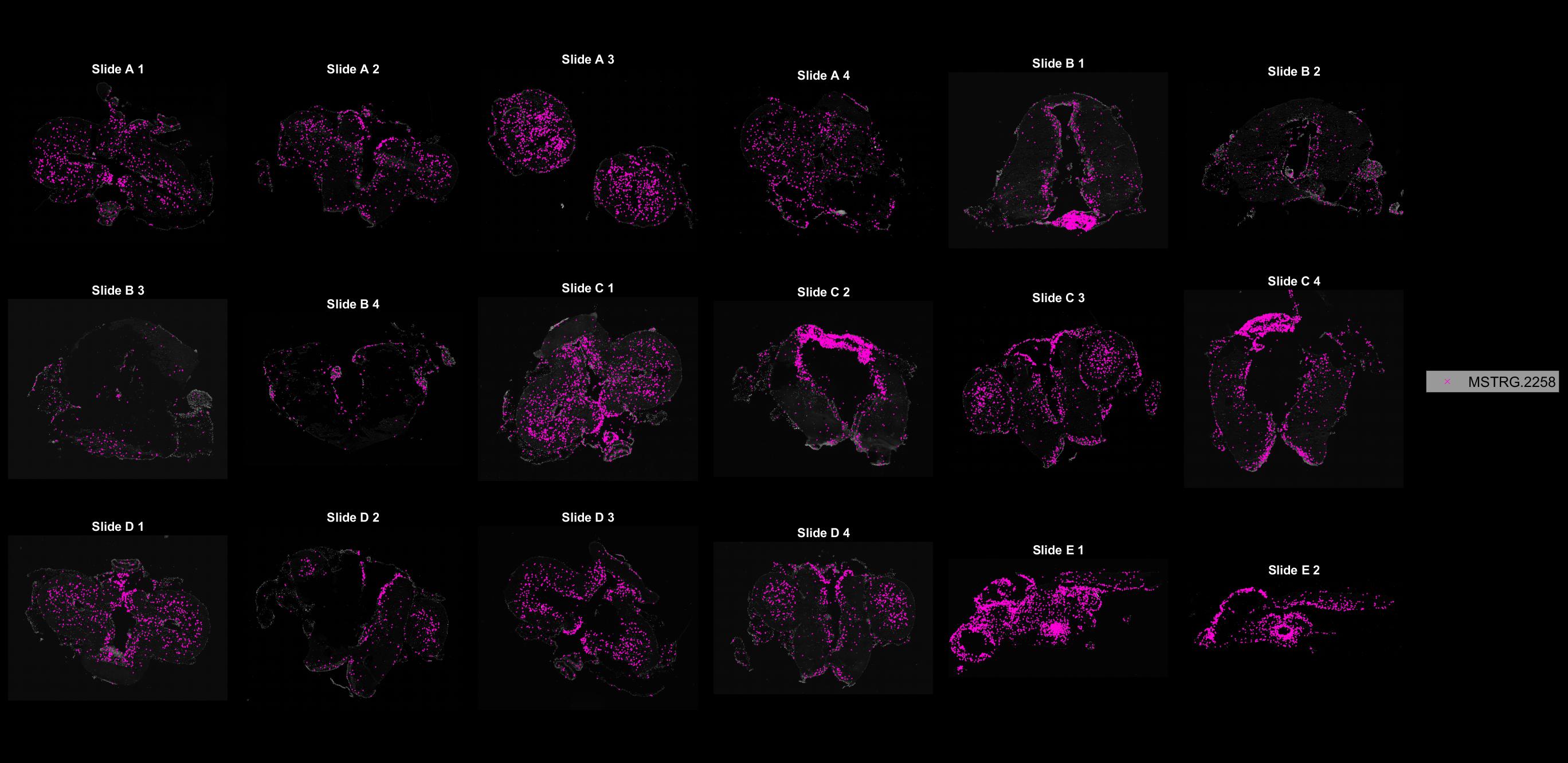

### MSTRG.2263.jpg

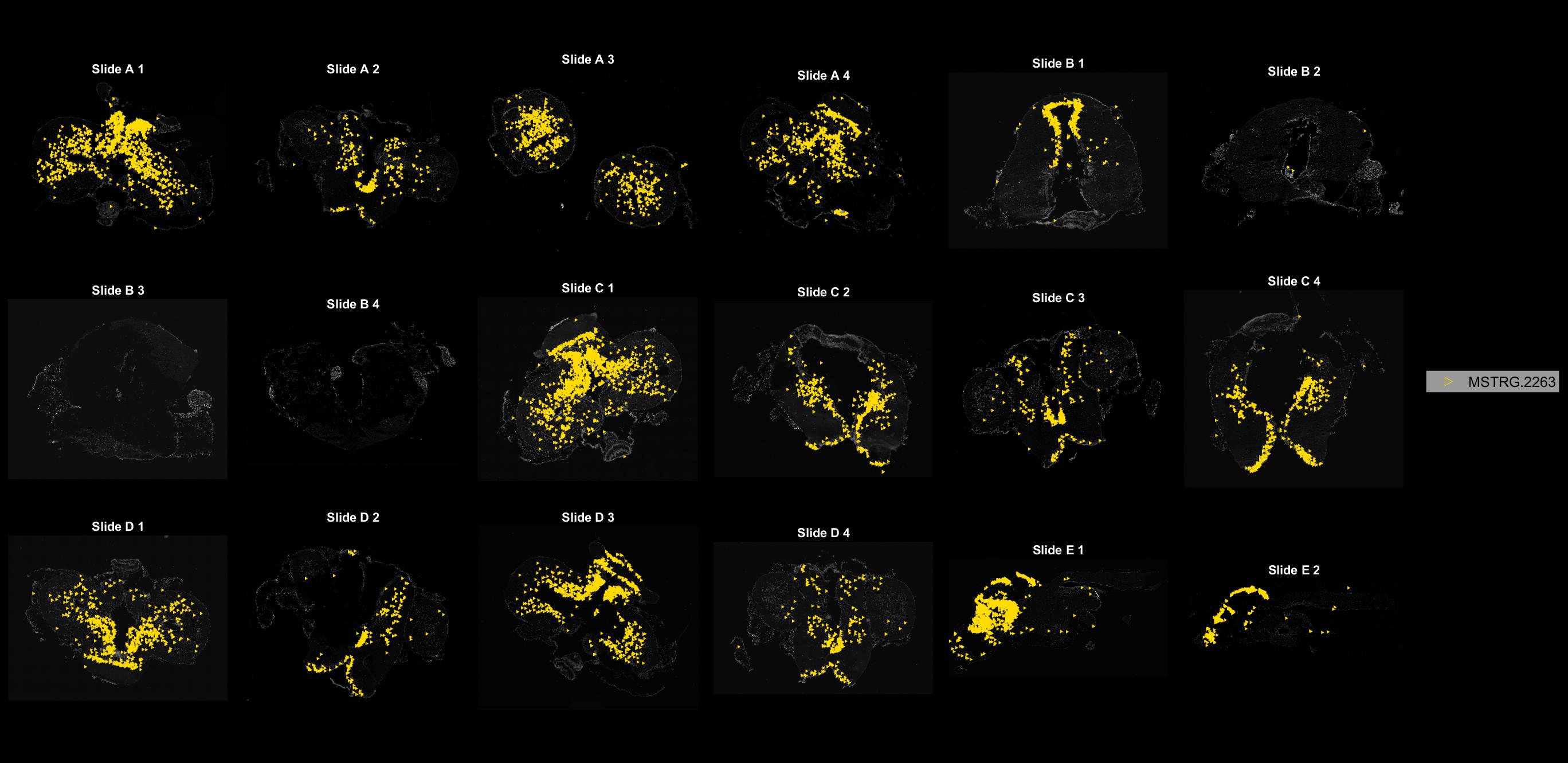

### MSTRG.2984.jpg

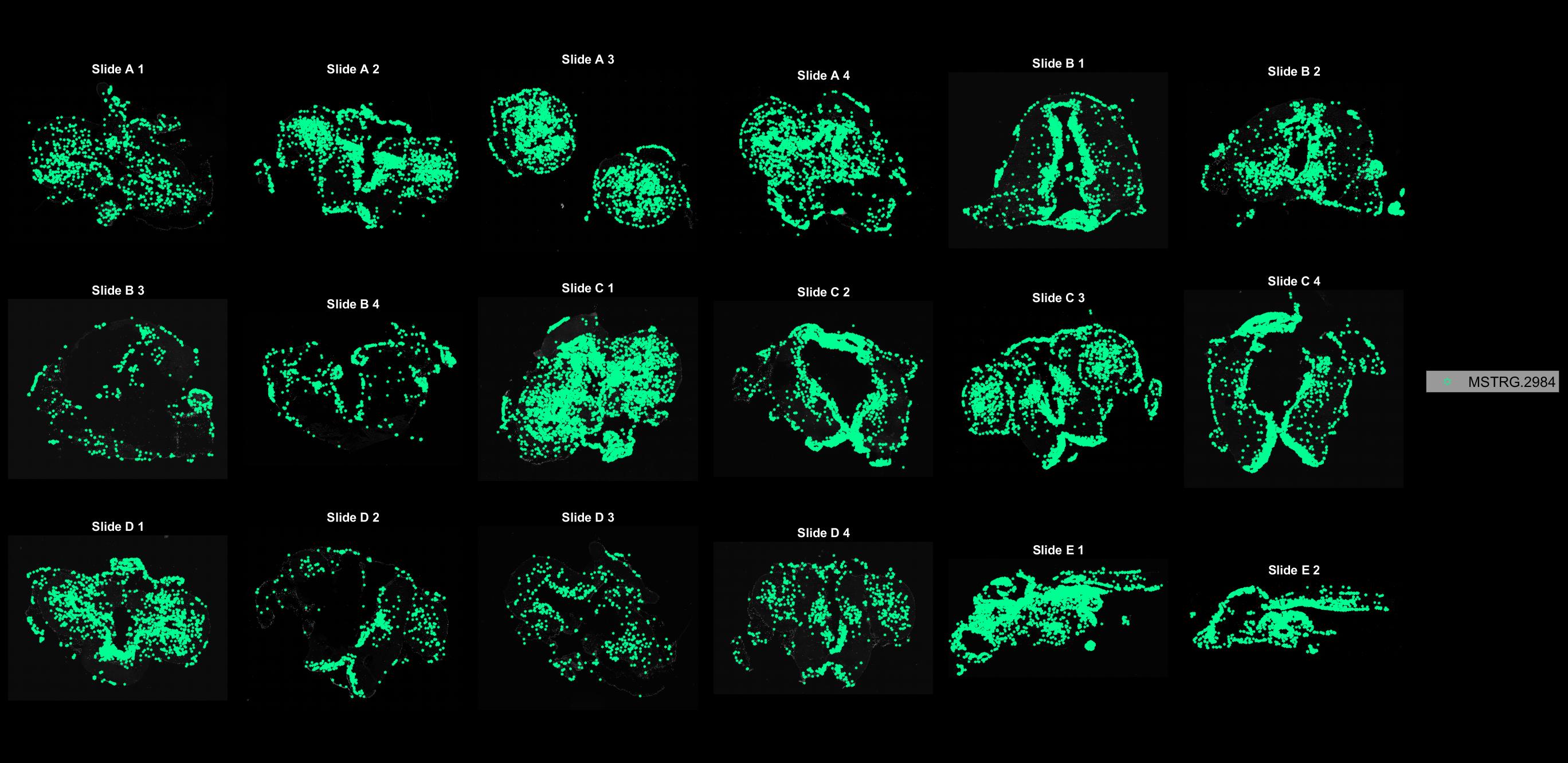

### MSTRG.3380.jpg

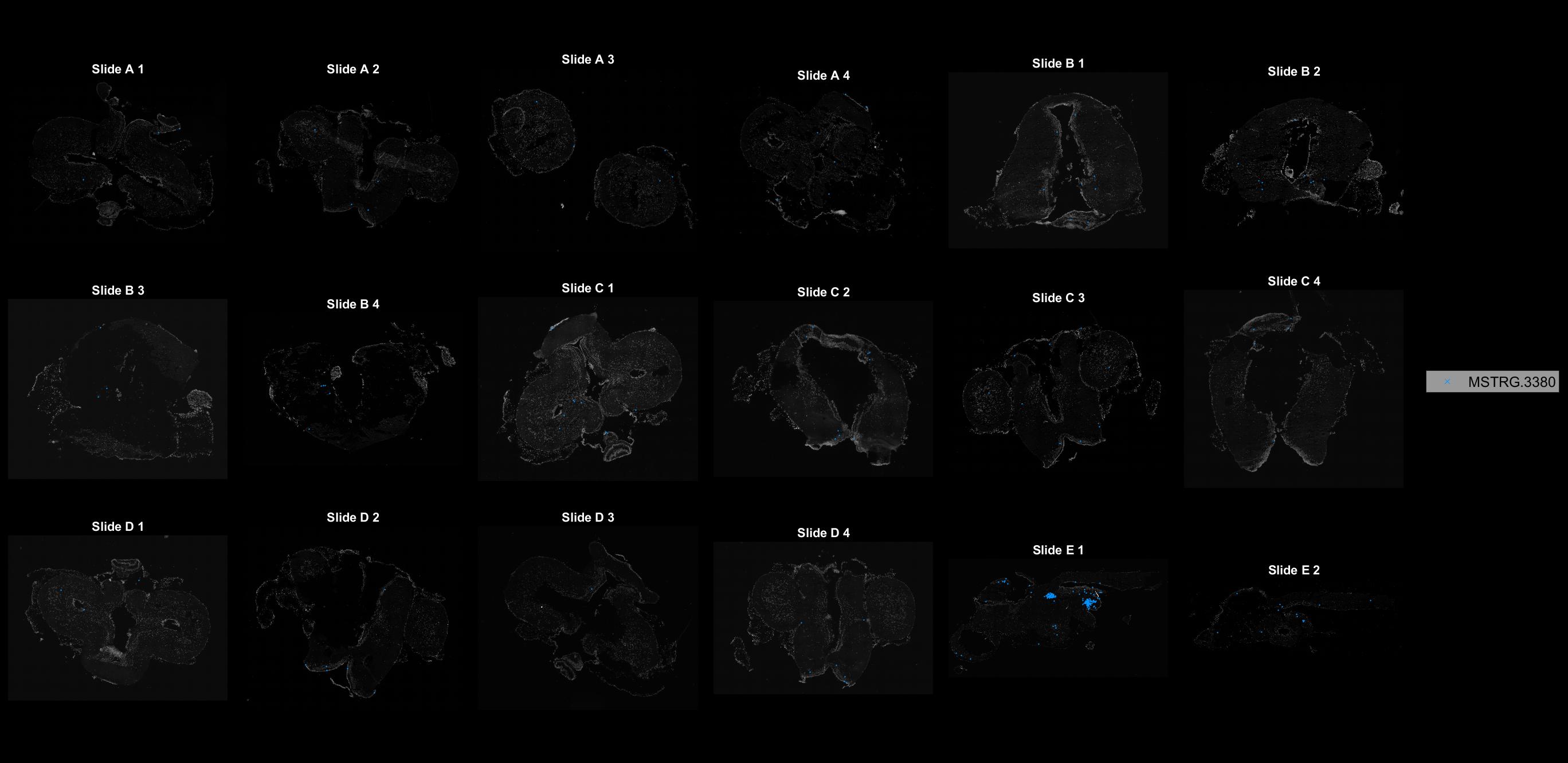

### MSTRG.3757.jpg

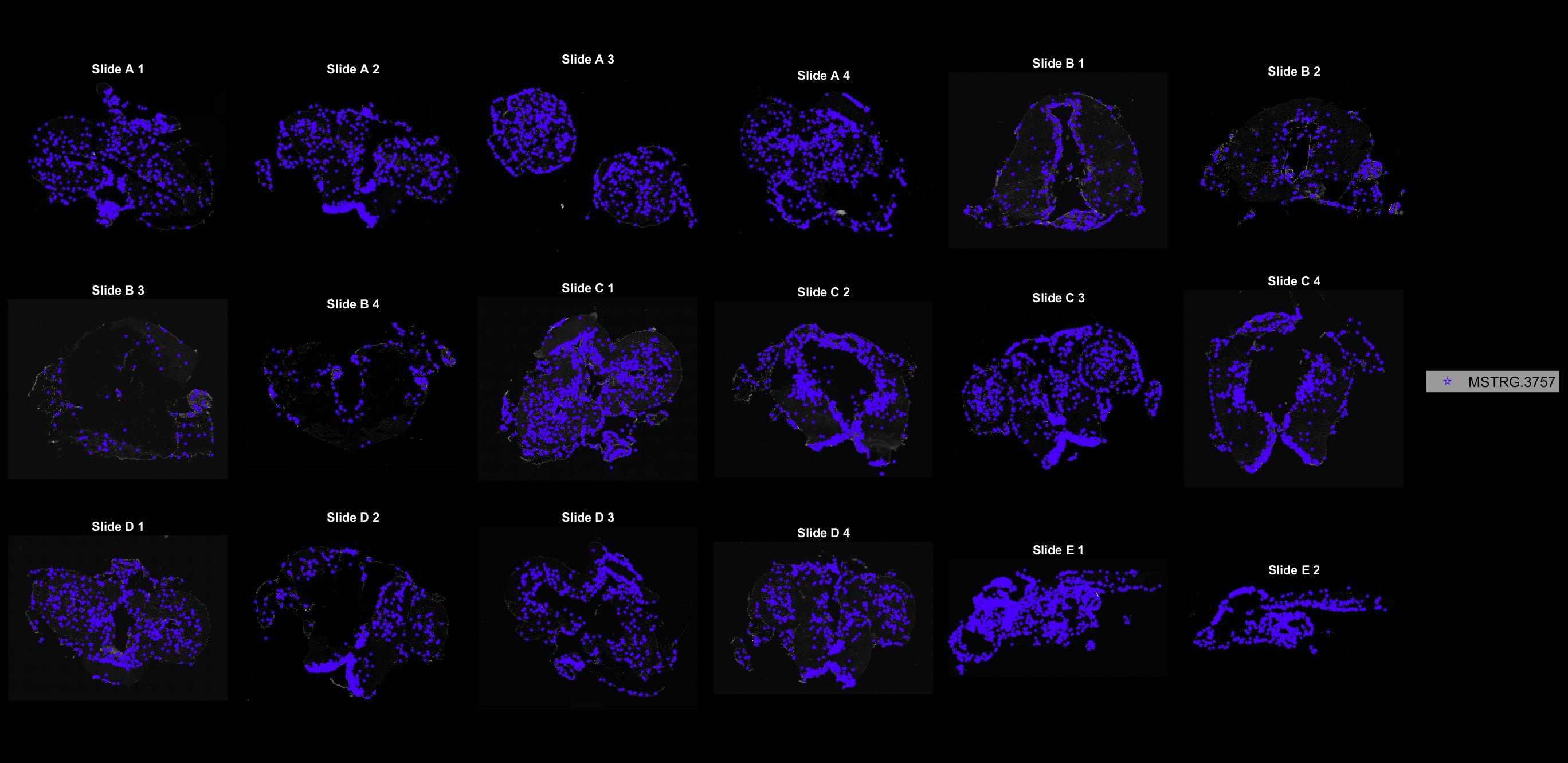

### MSTRG.3884.jpg

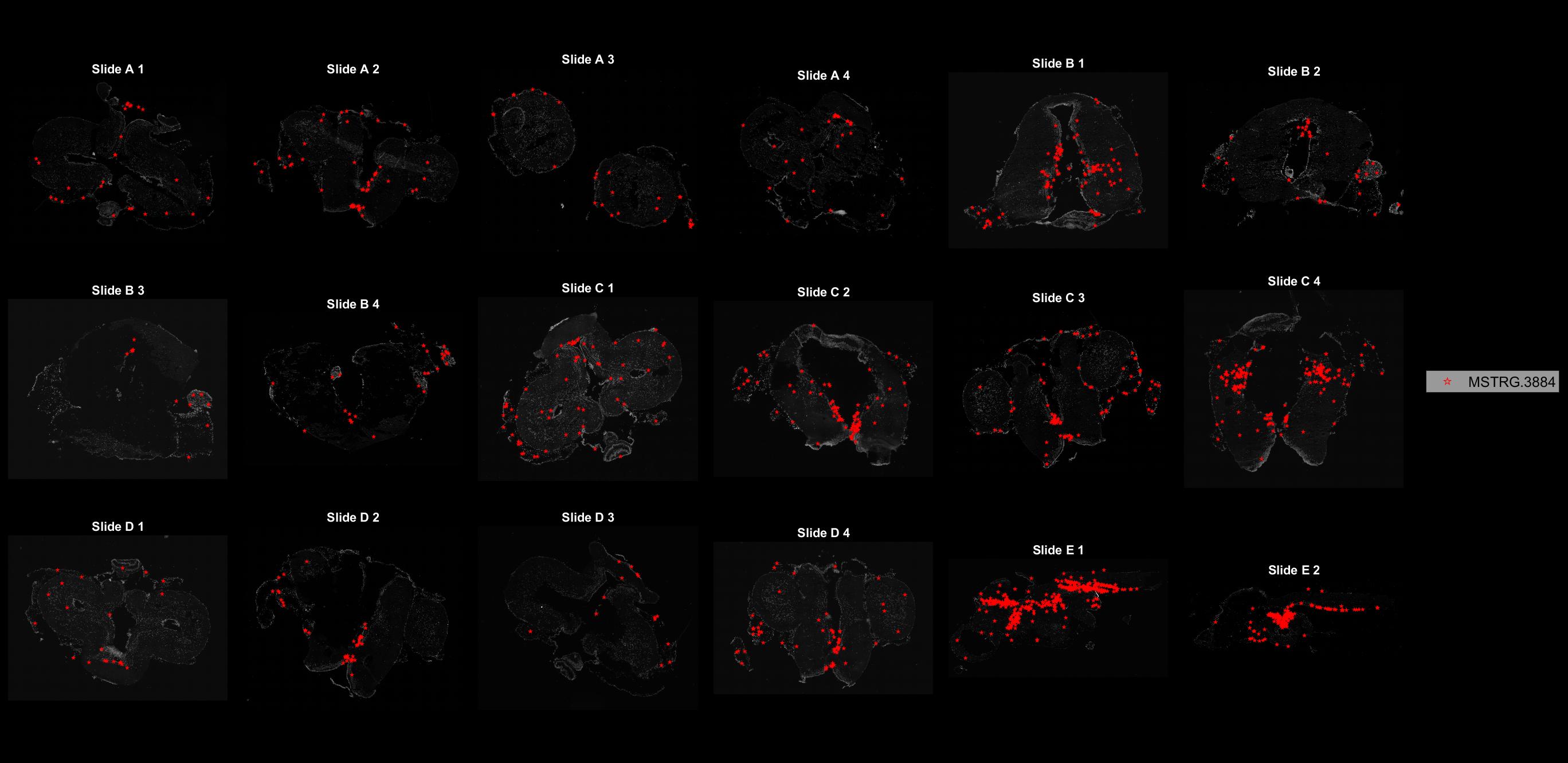

### MSTRG.3946.jpg

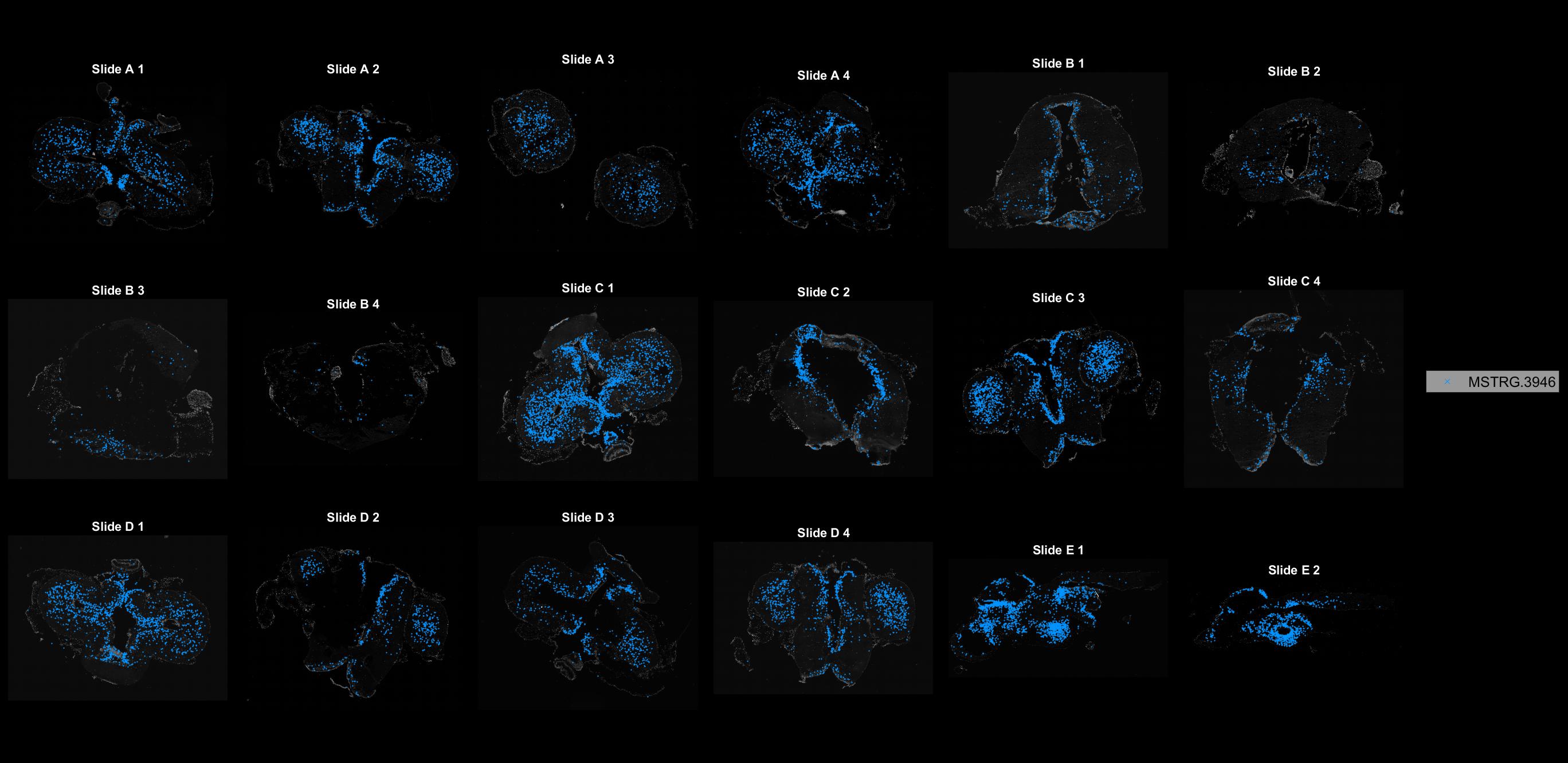

### MSTRG.4061.jpg

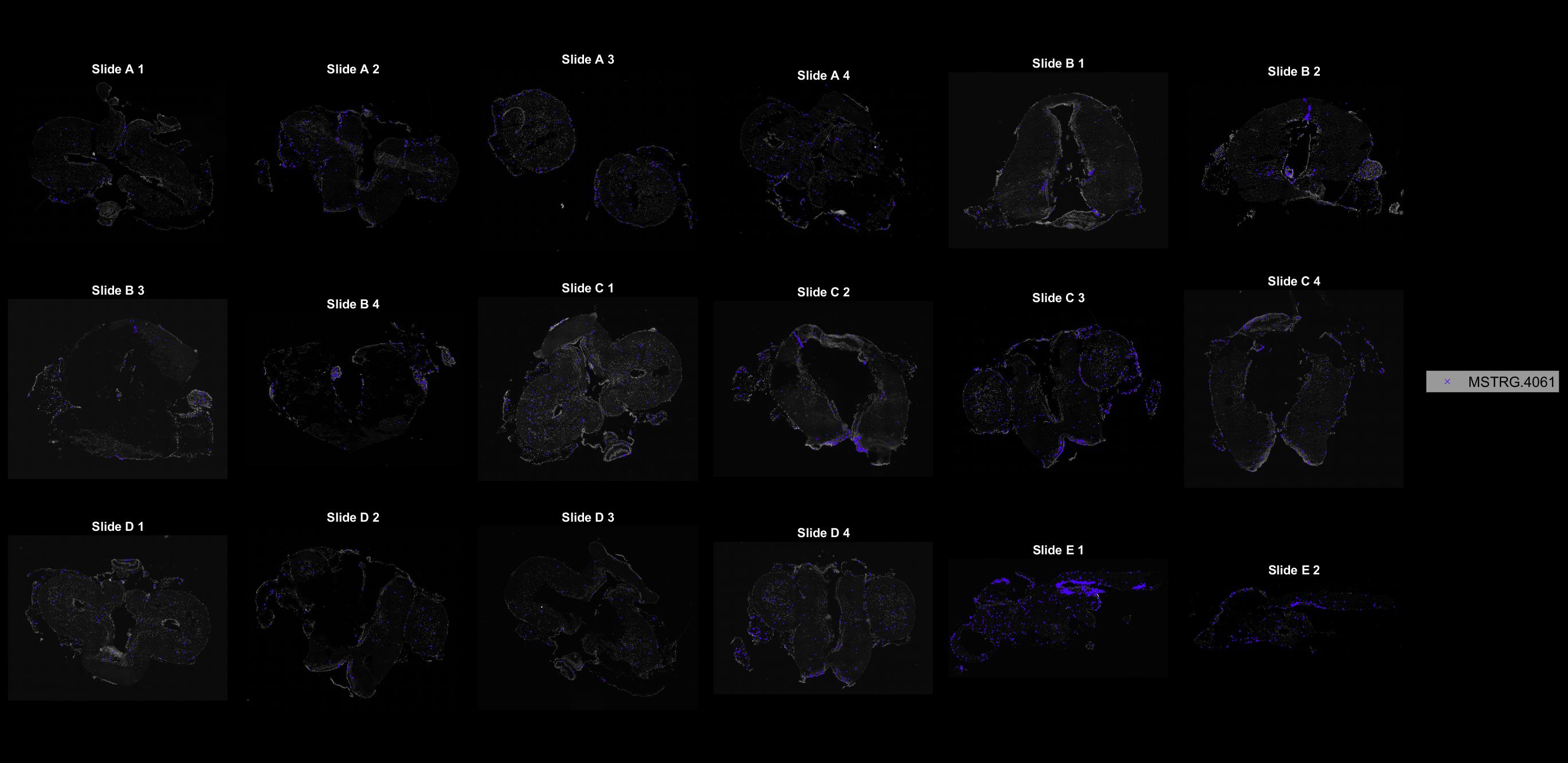

### MSTRG.4350.jpg

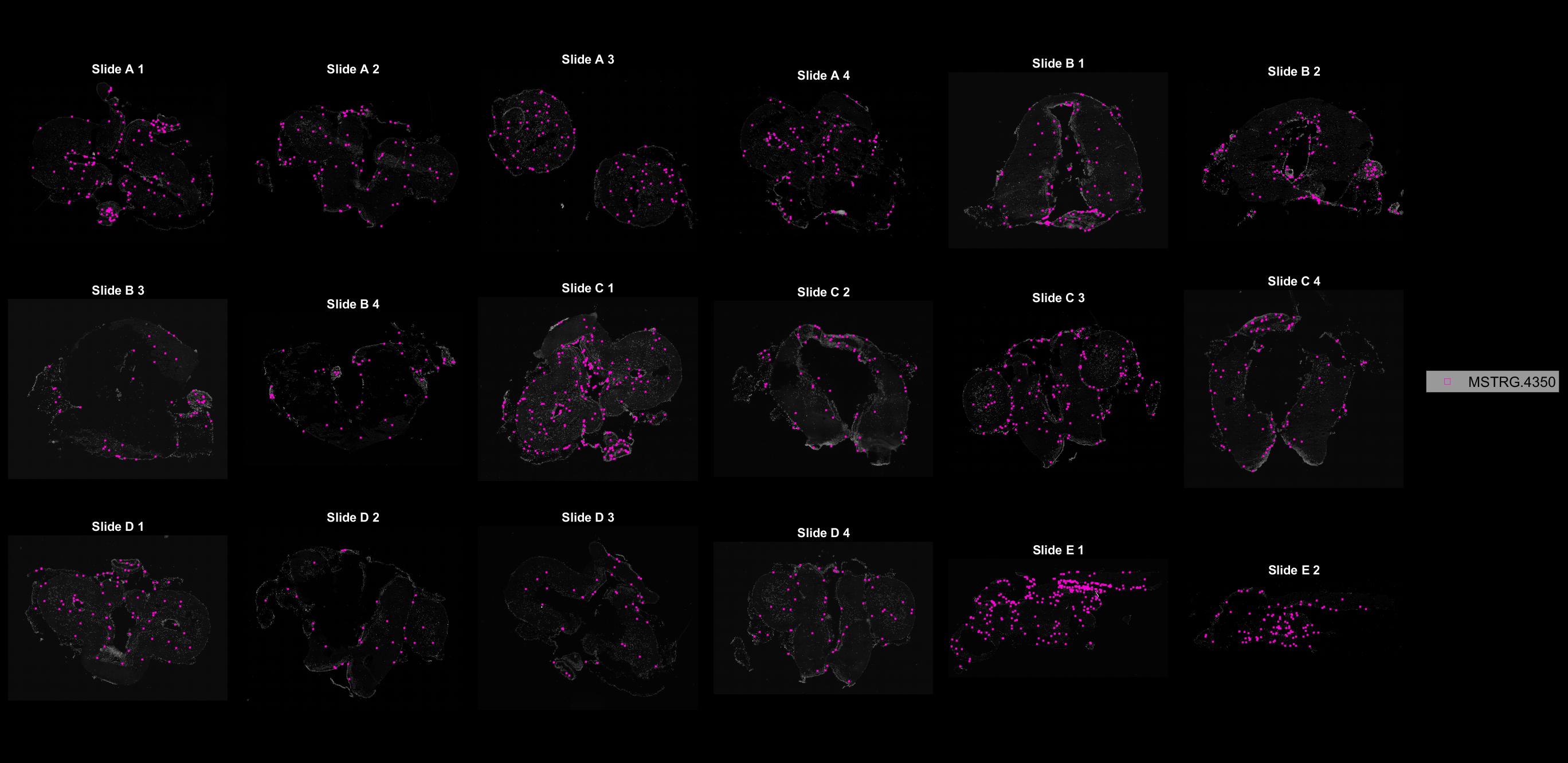

### MSTRG.4456.jpg

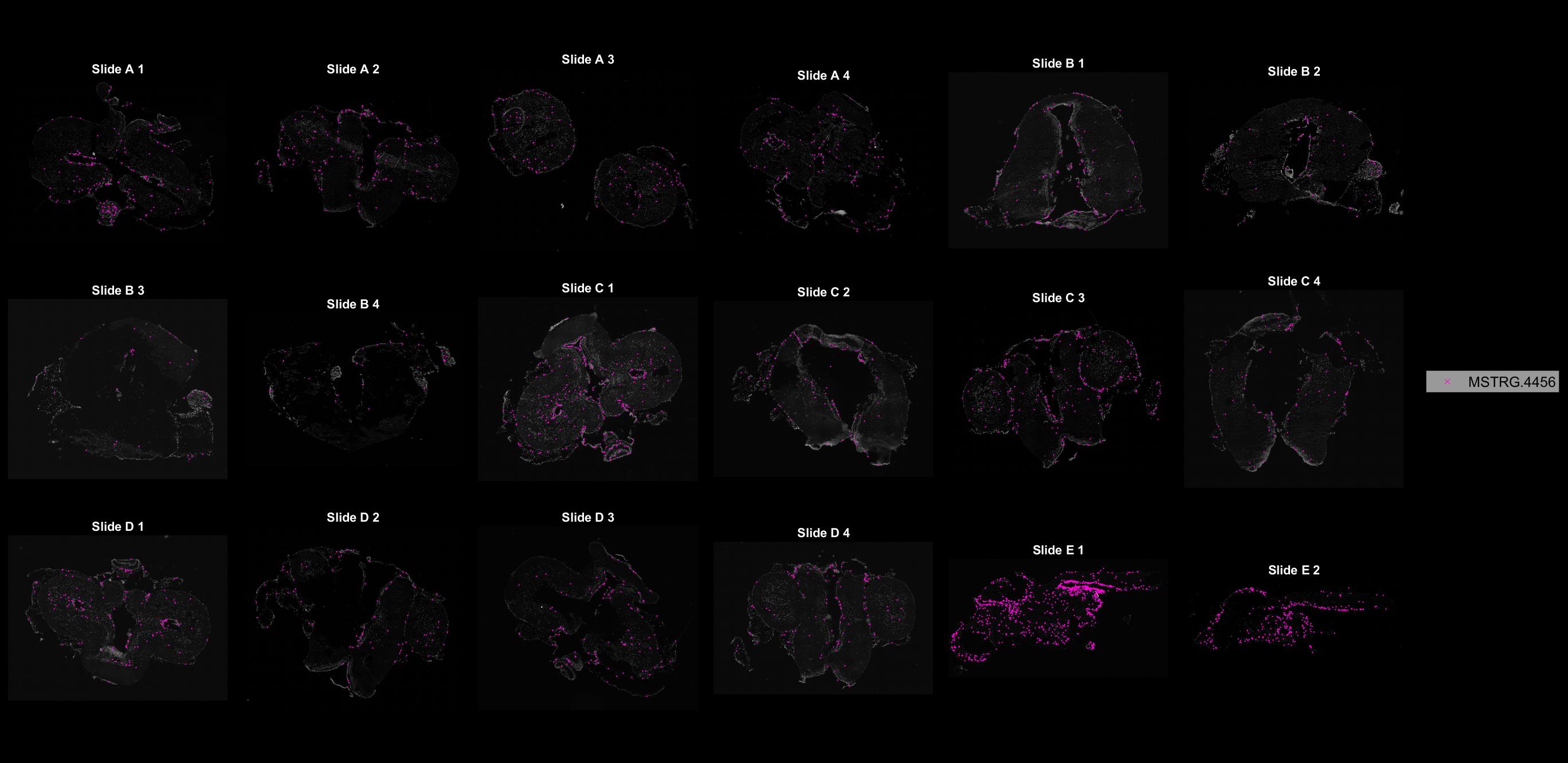

### MSTRG.4578.jpg

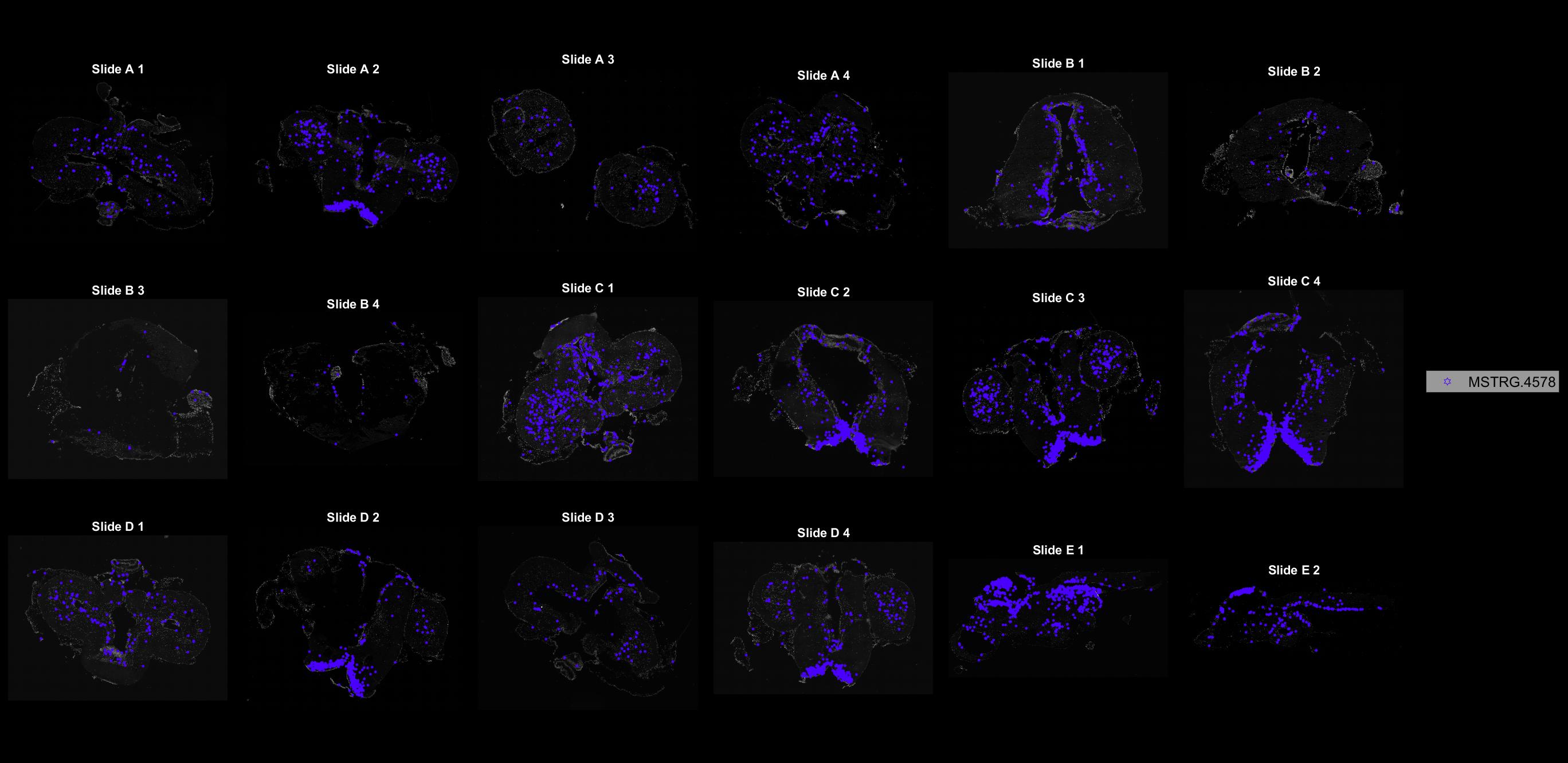

### MSTRG.4669.jpg

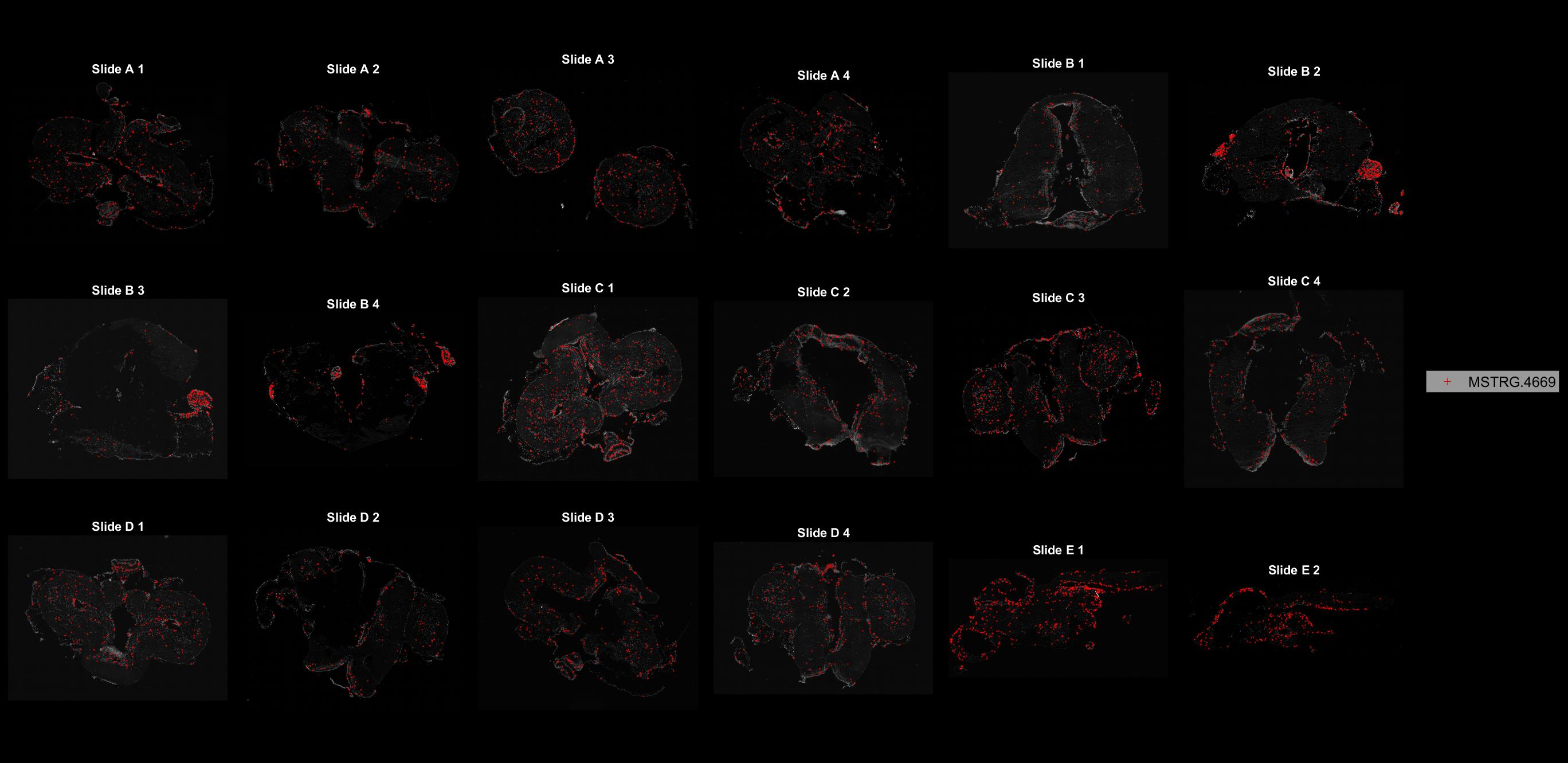

### MSTRG.4895.jpg

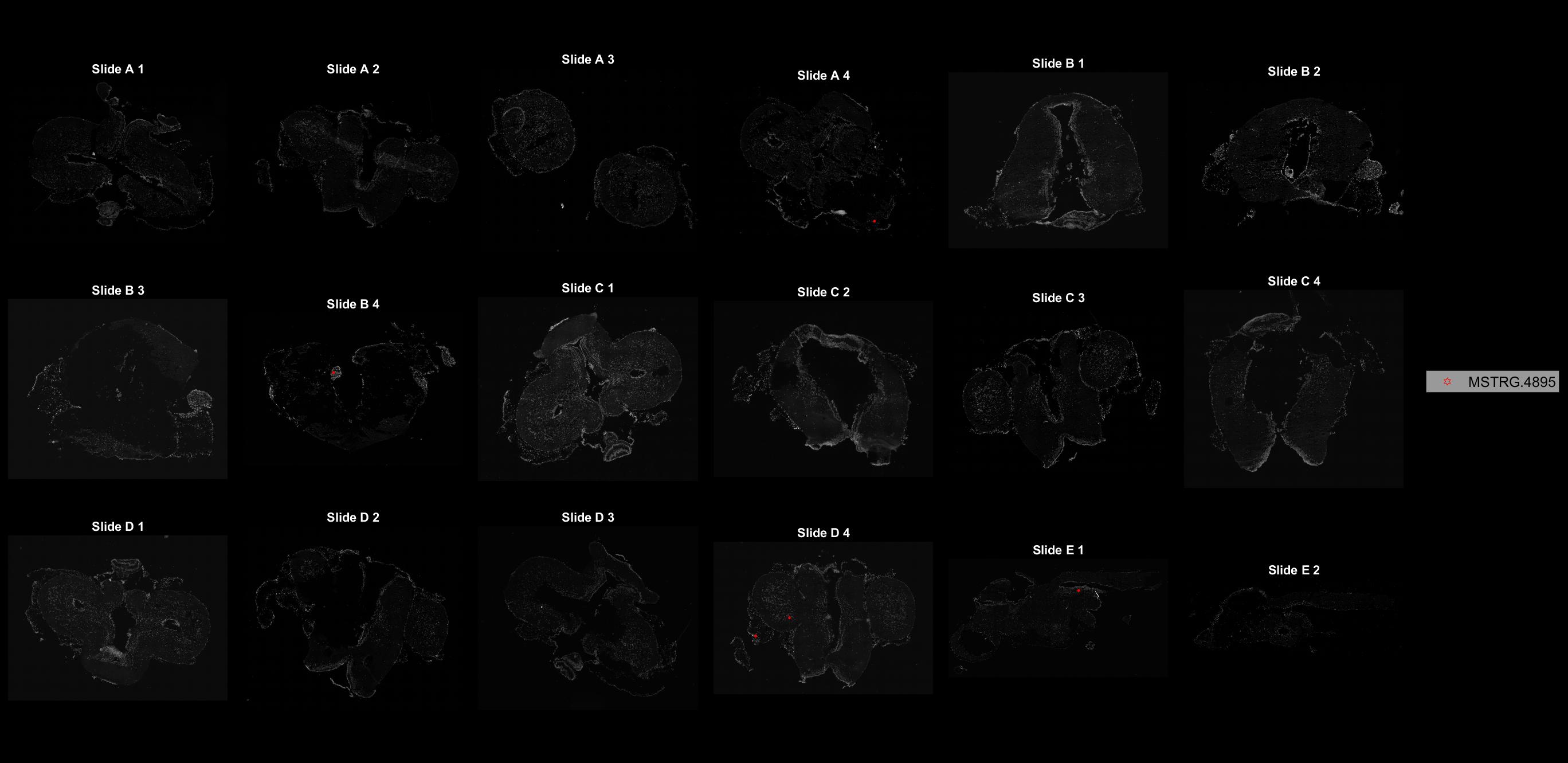

### MSTRG.4985.jpg

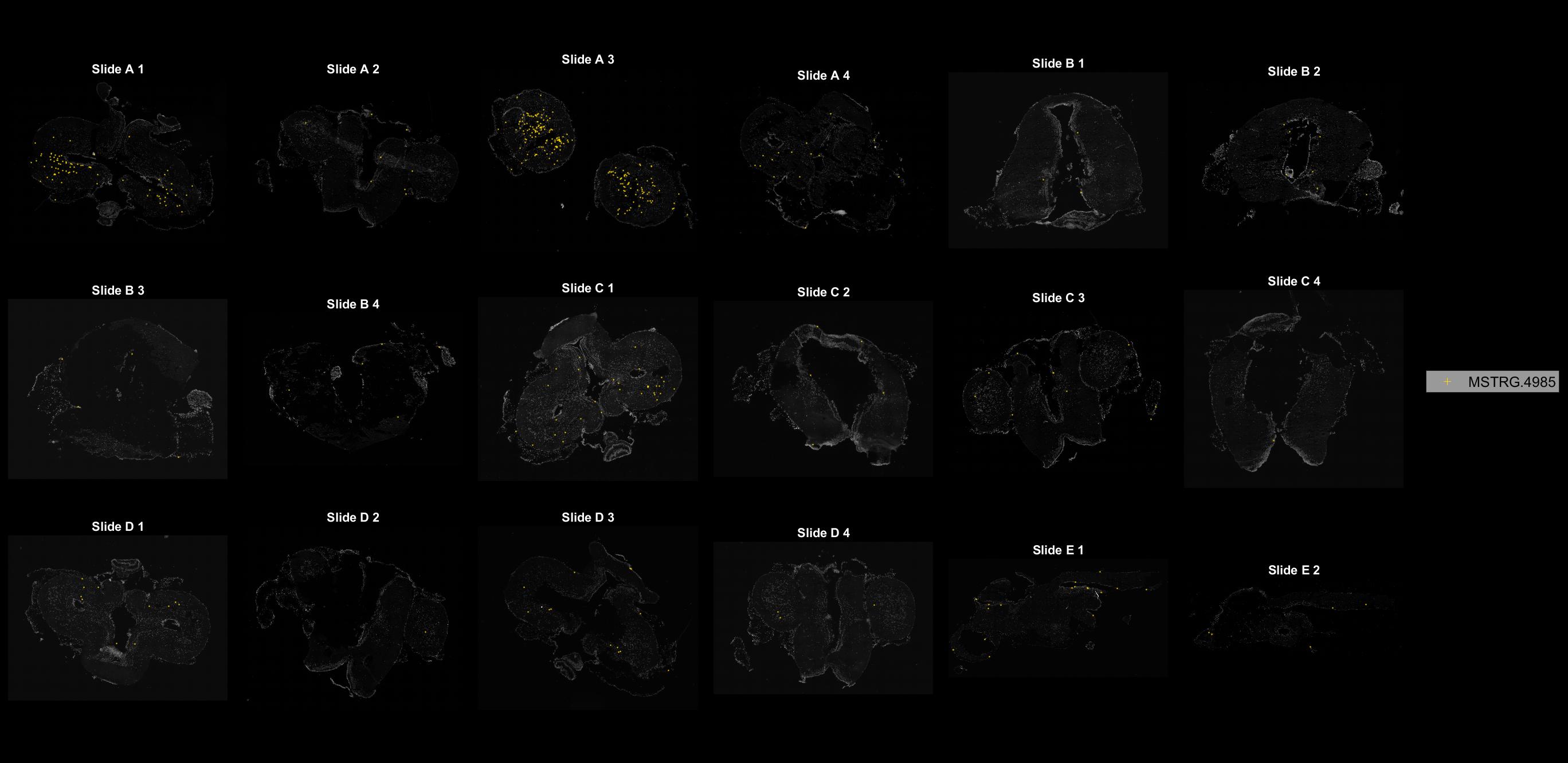

### MSTRG.4994.jpg

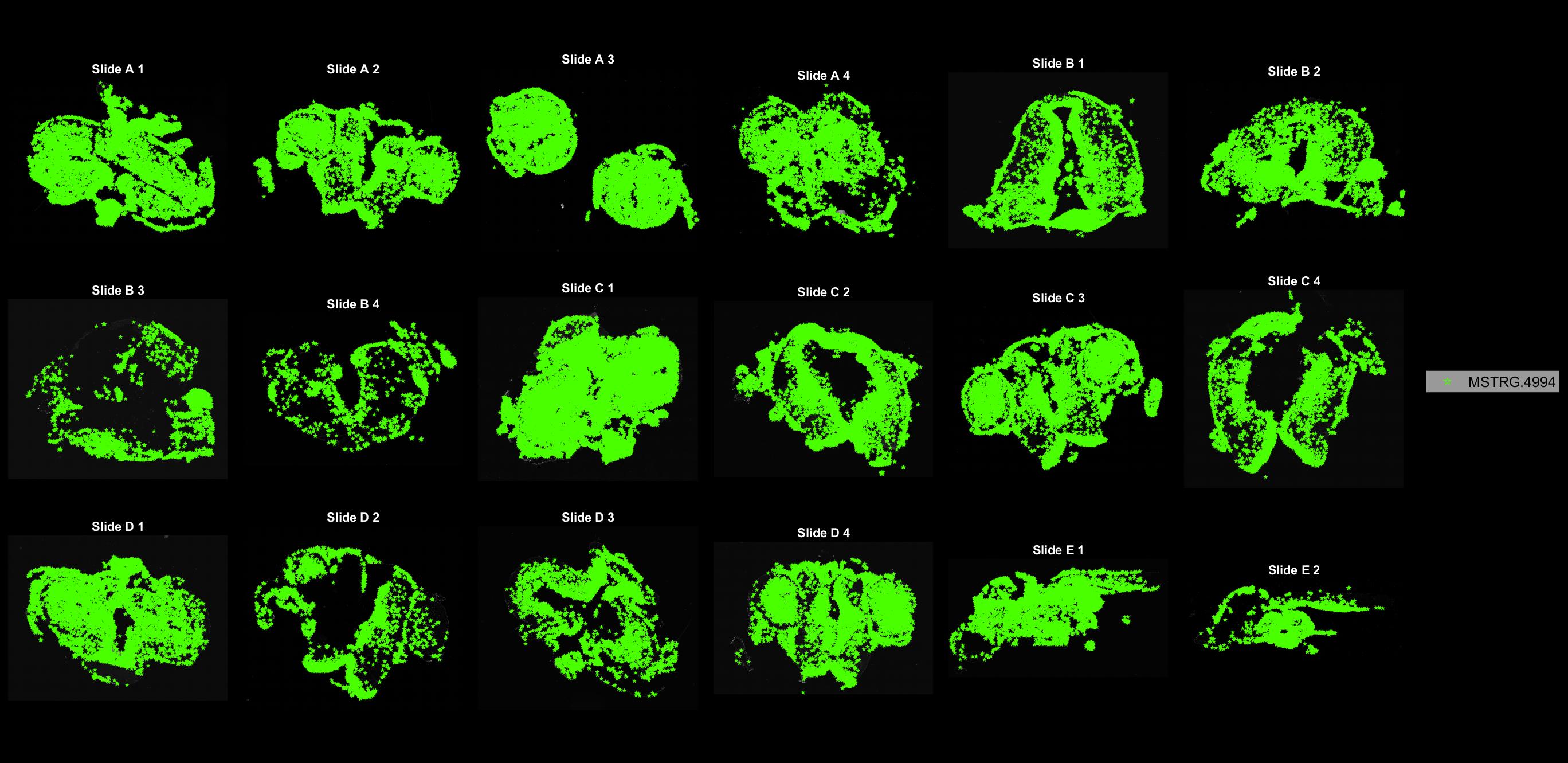

### MSTRG.5128.jpg

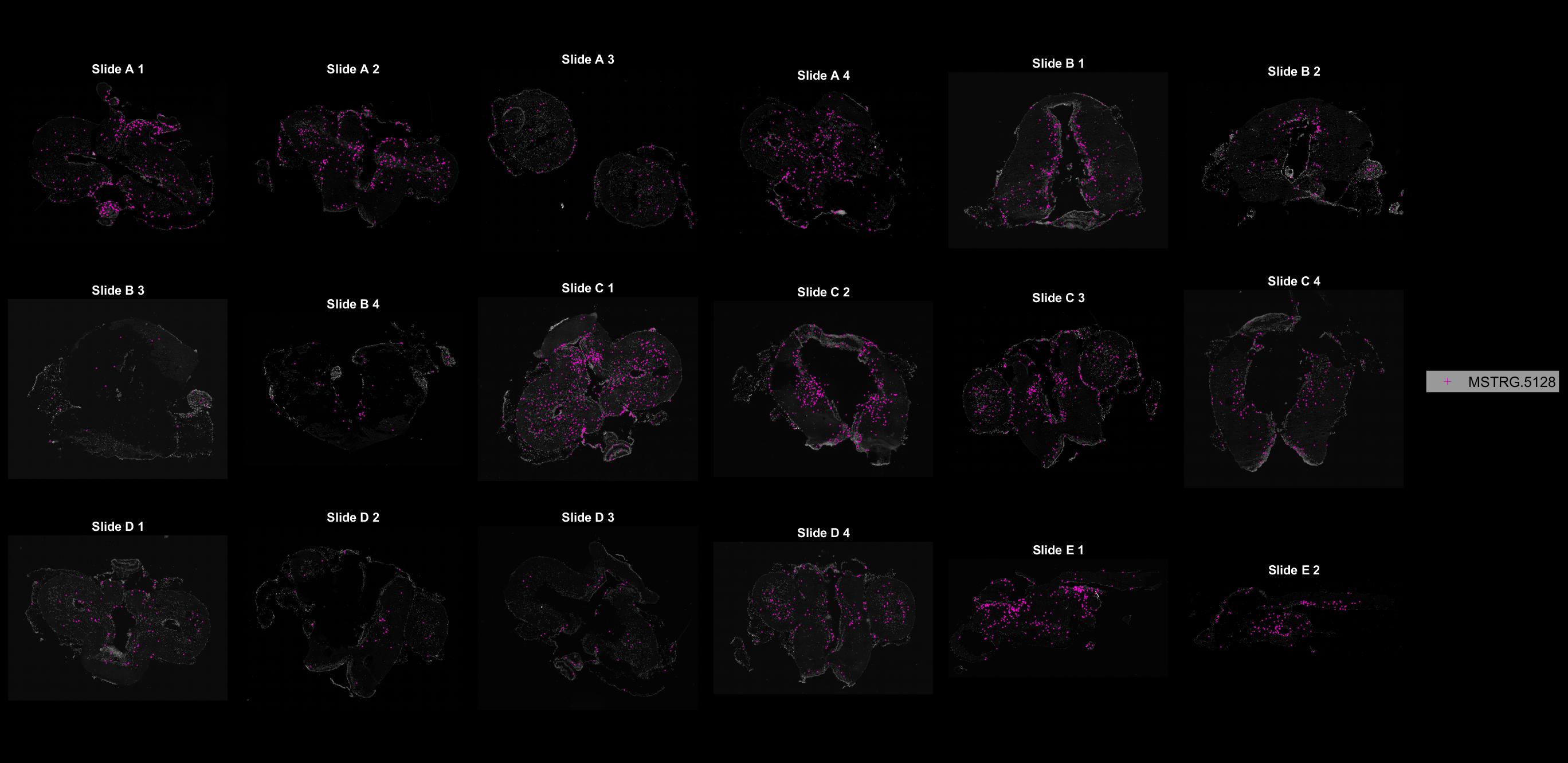

### MSTRG.5329.jpg

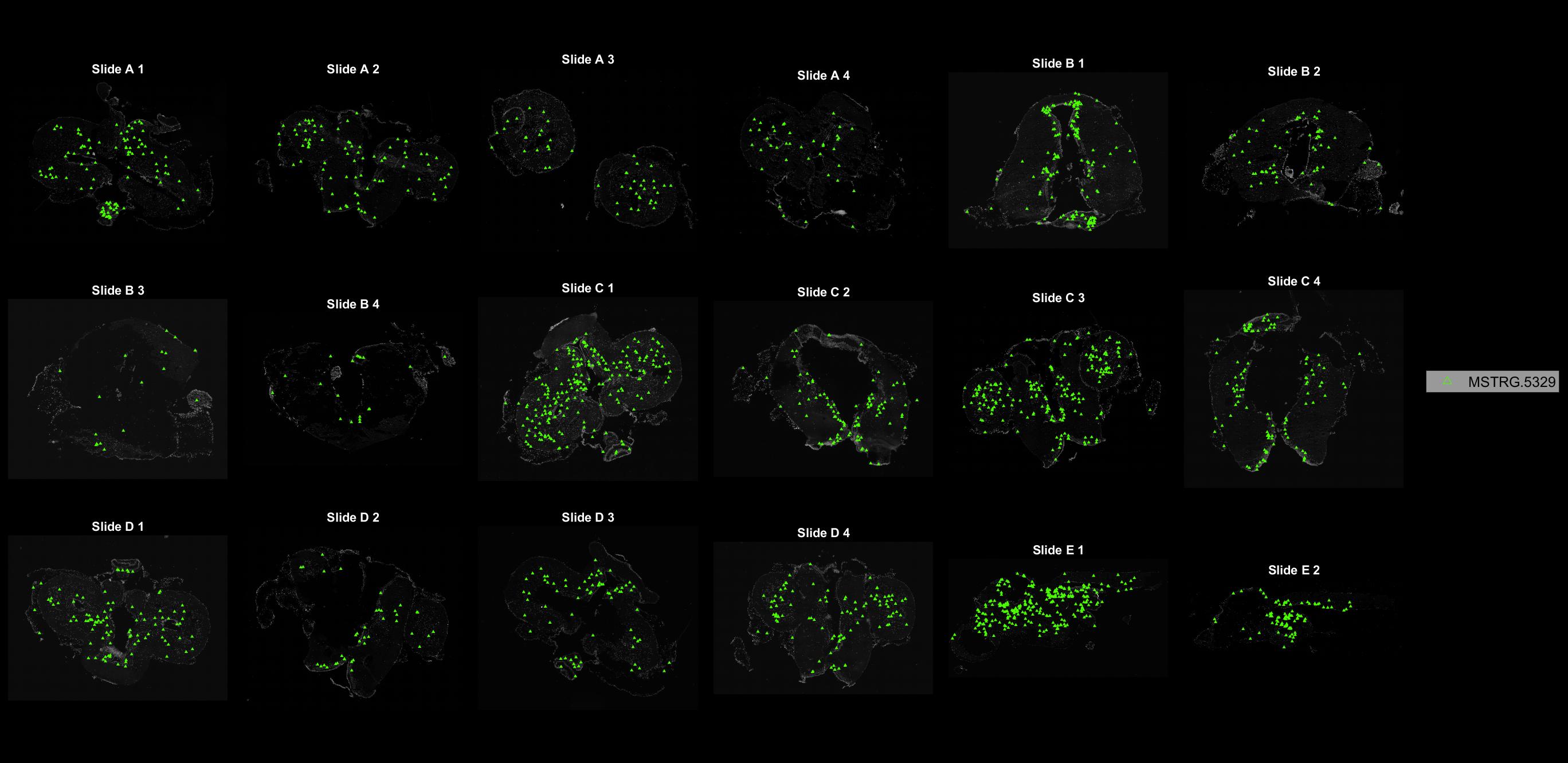

### MSTRG.5332.jpg

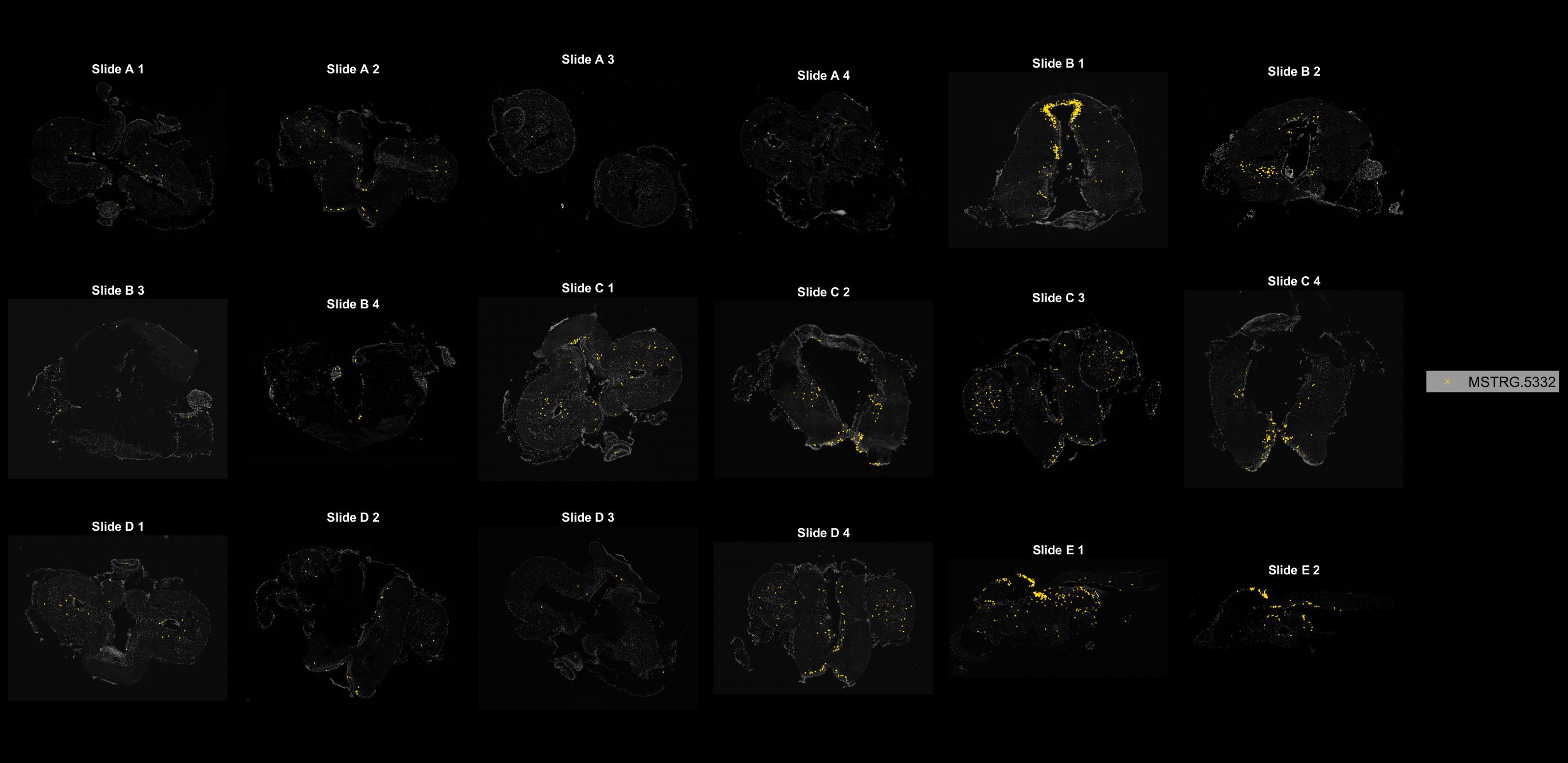

### MSTRG.5367.jpg

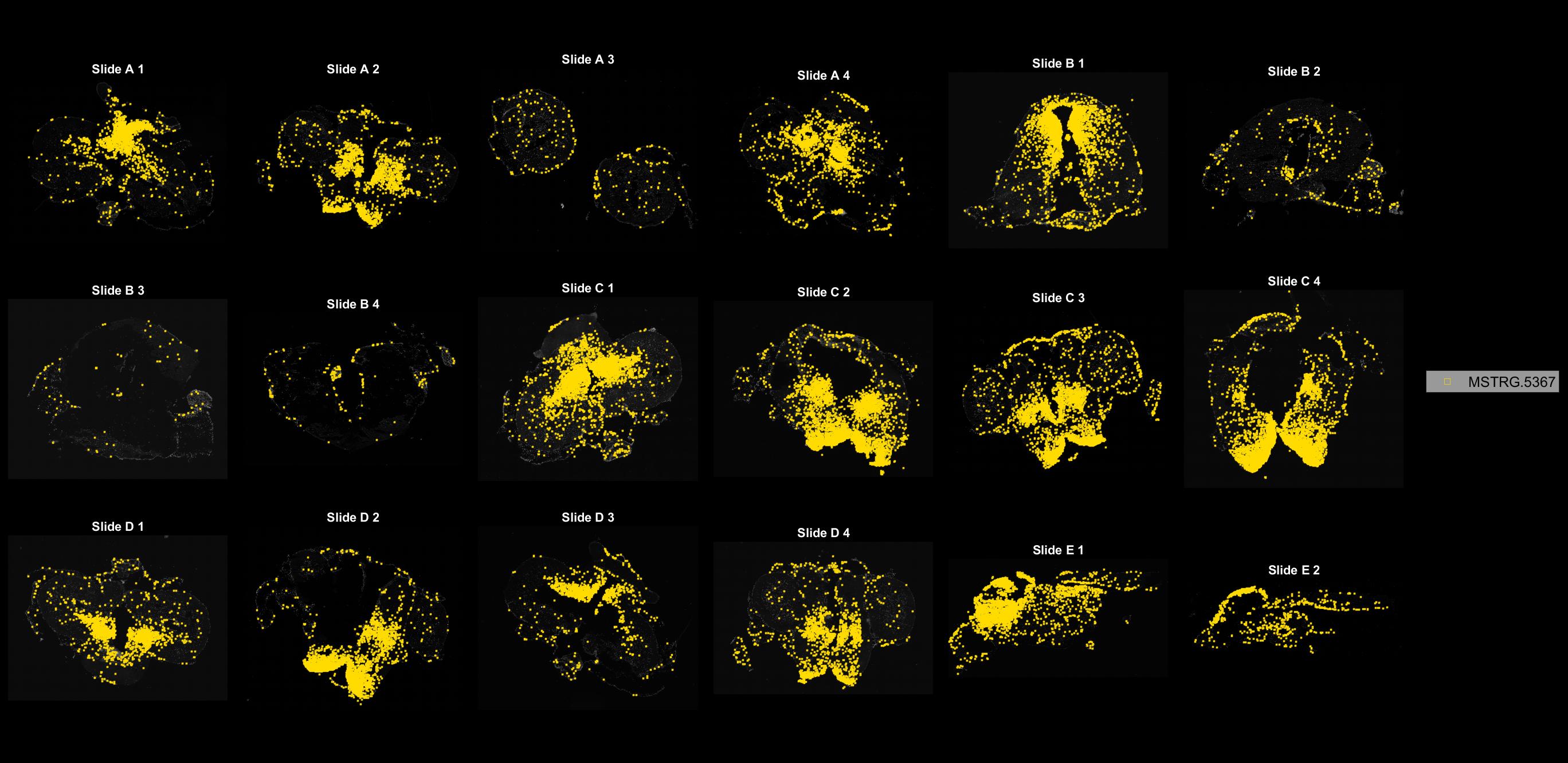

### MSTRG.5559.jpg

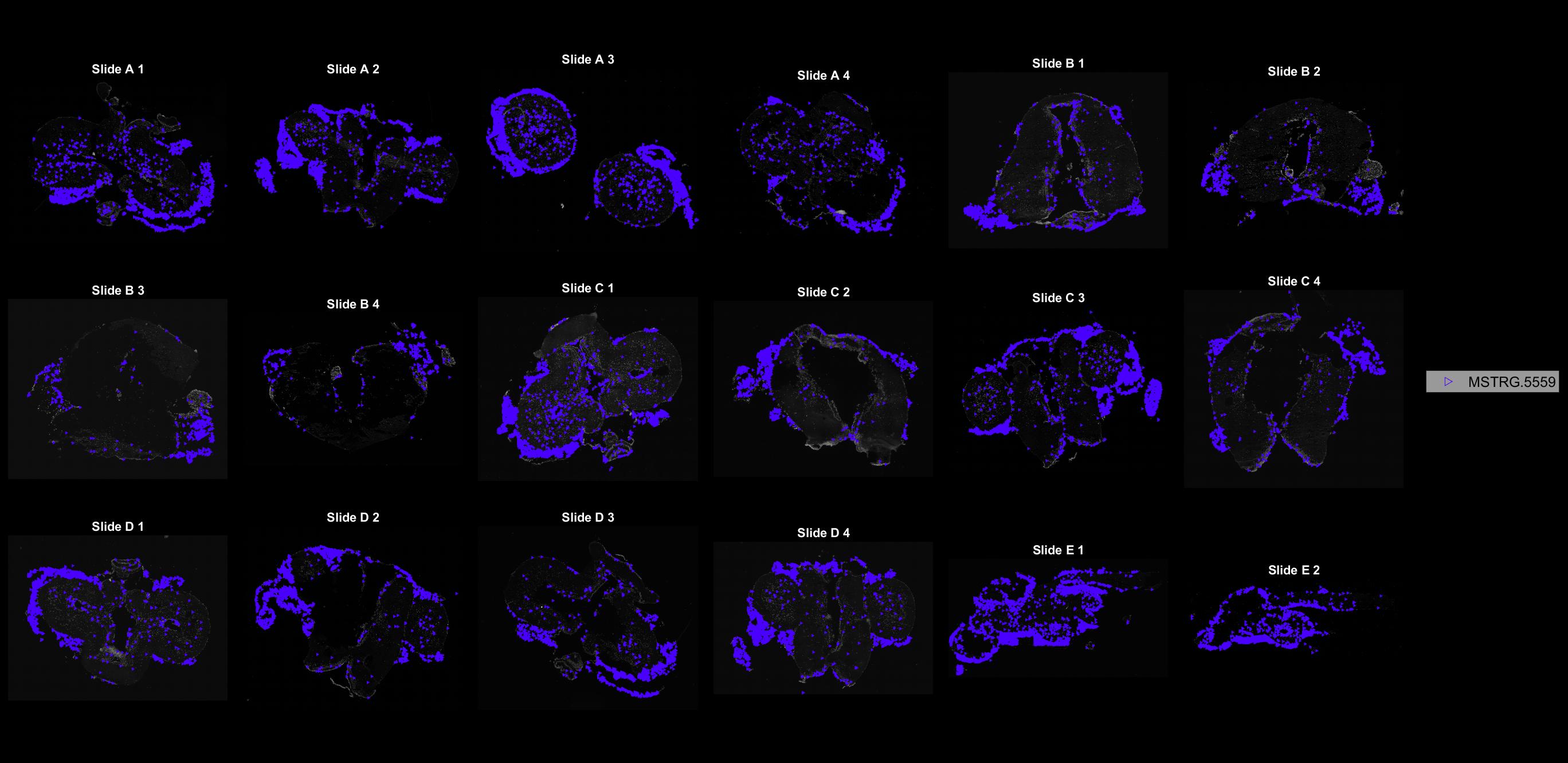

### MSTRG.5560.jpg

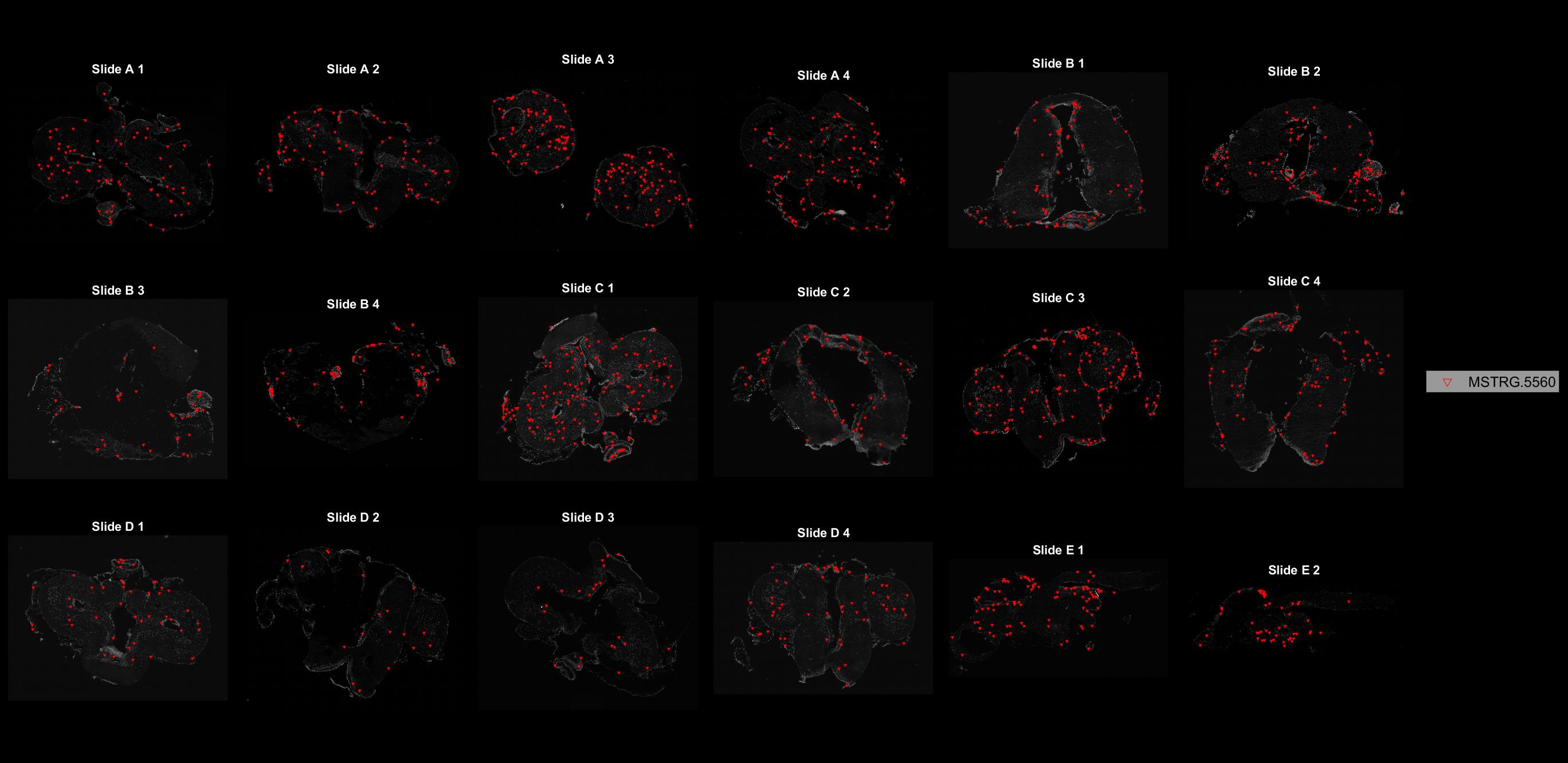

### MSTRG.5689.jpg

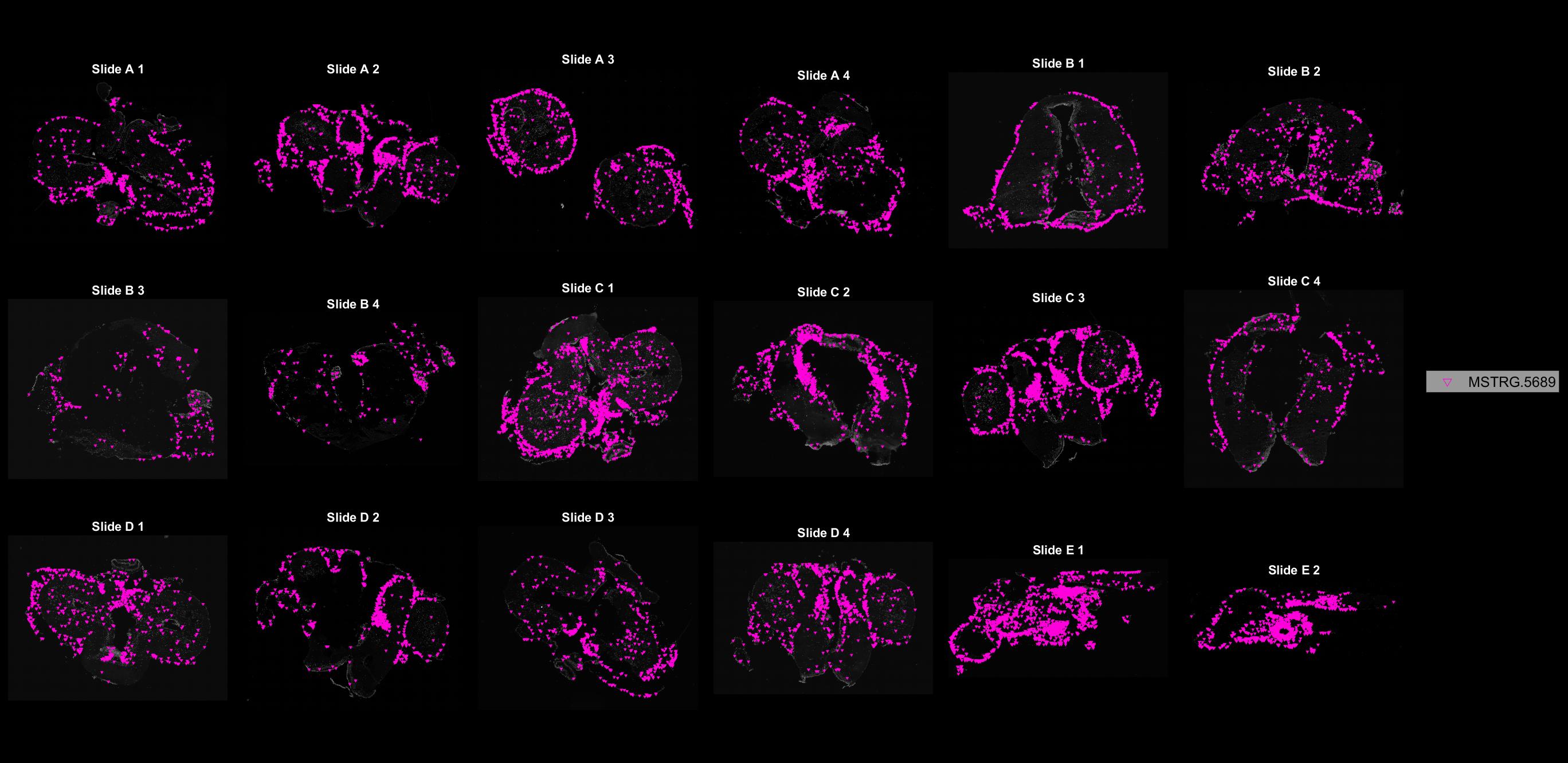

### MSTRG.5690.jpg

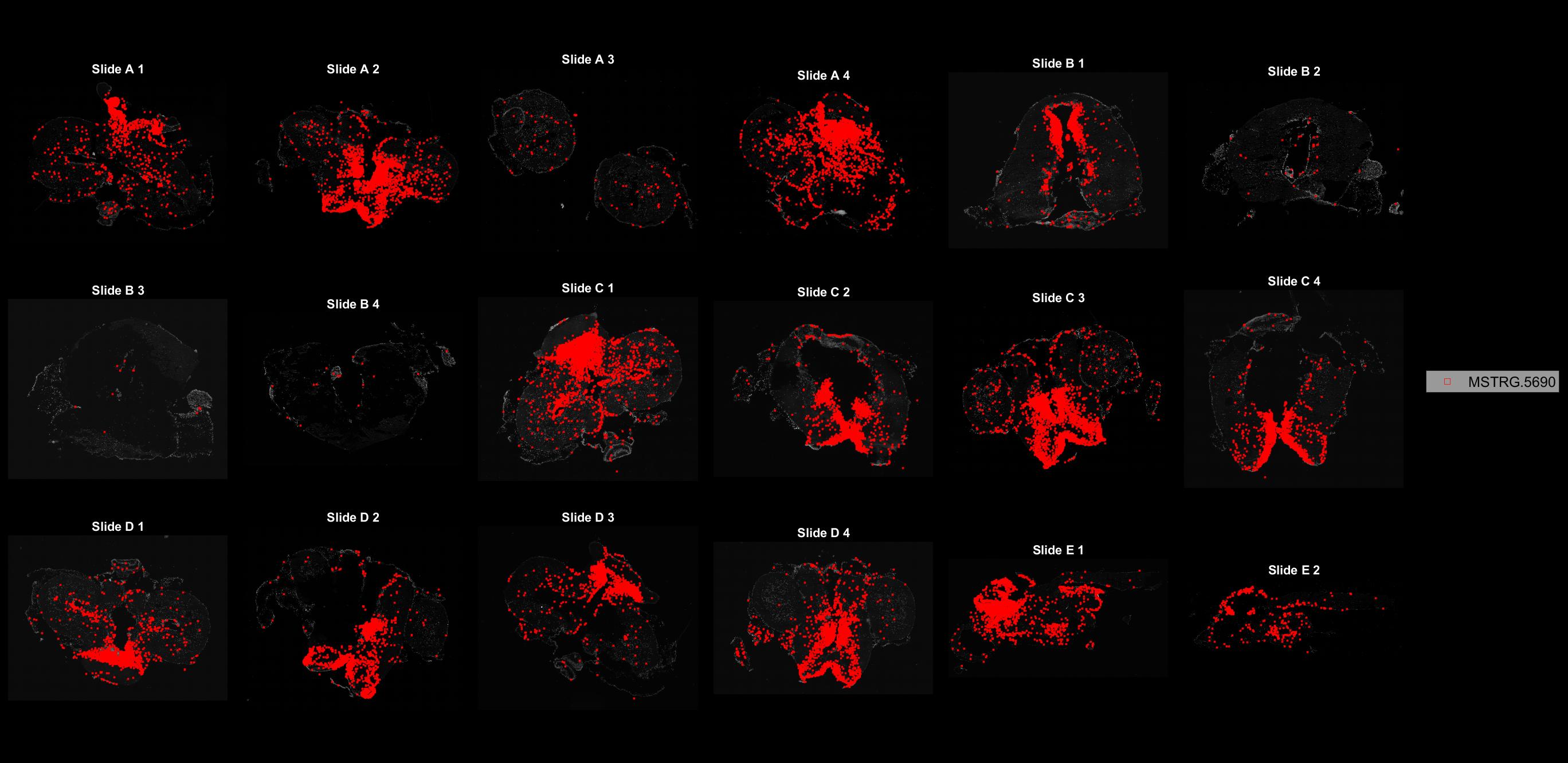

### MSTRG.5760.jpg

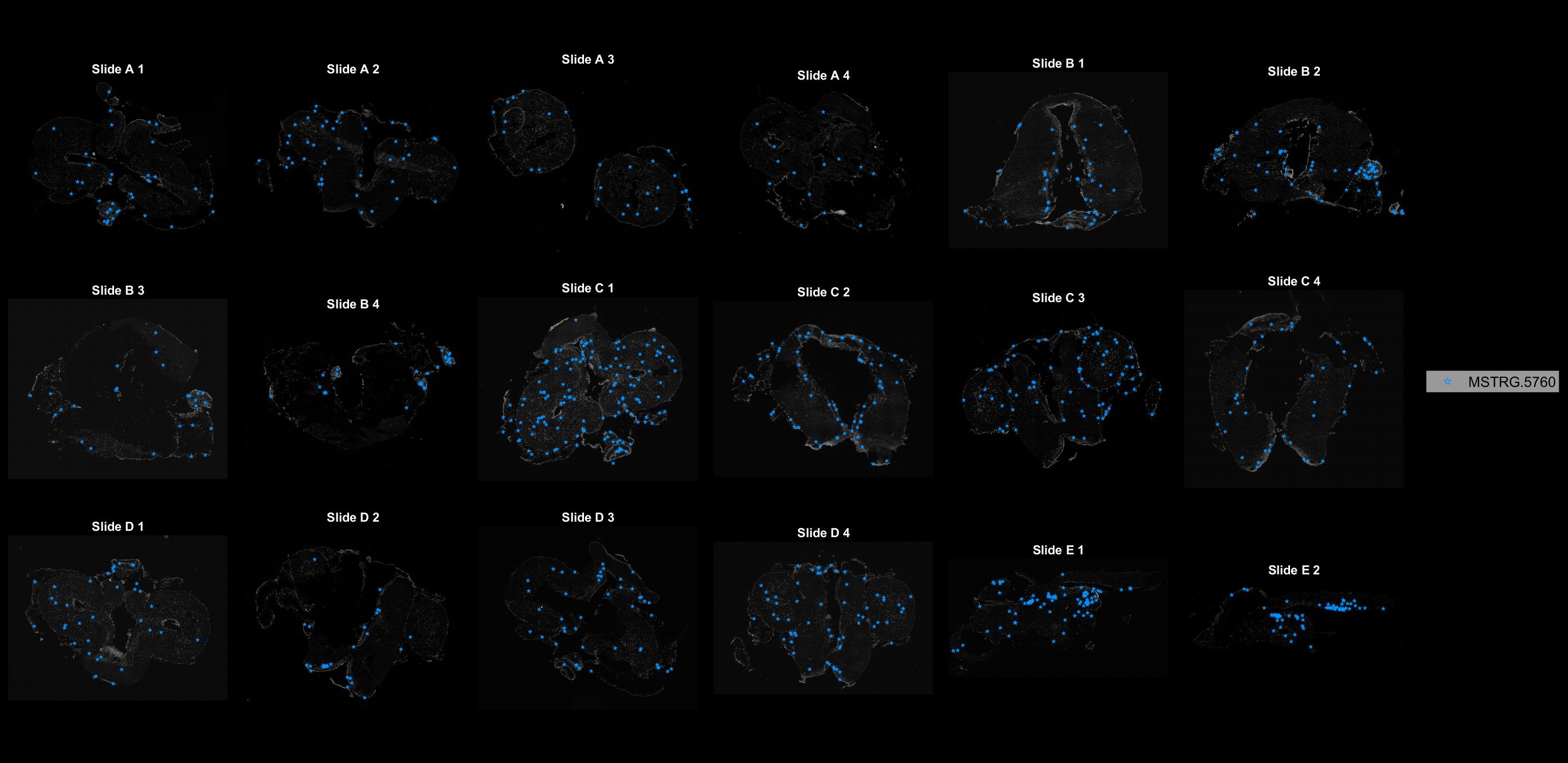

### MSTRG.5777.jpg

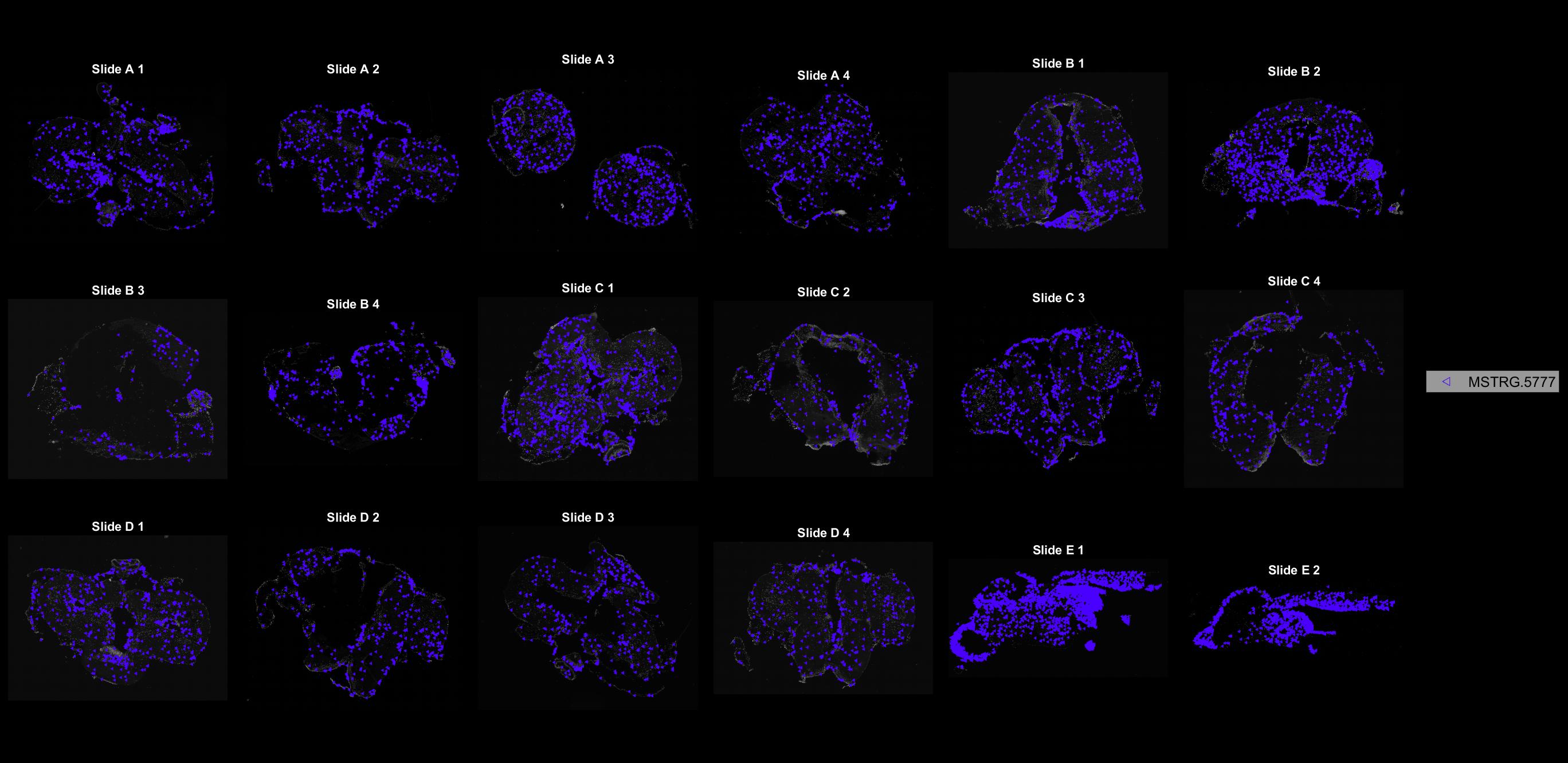

### MSTRG.5979.jpg

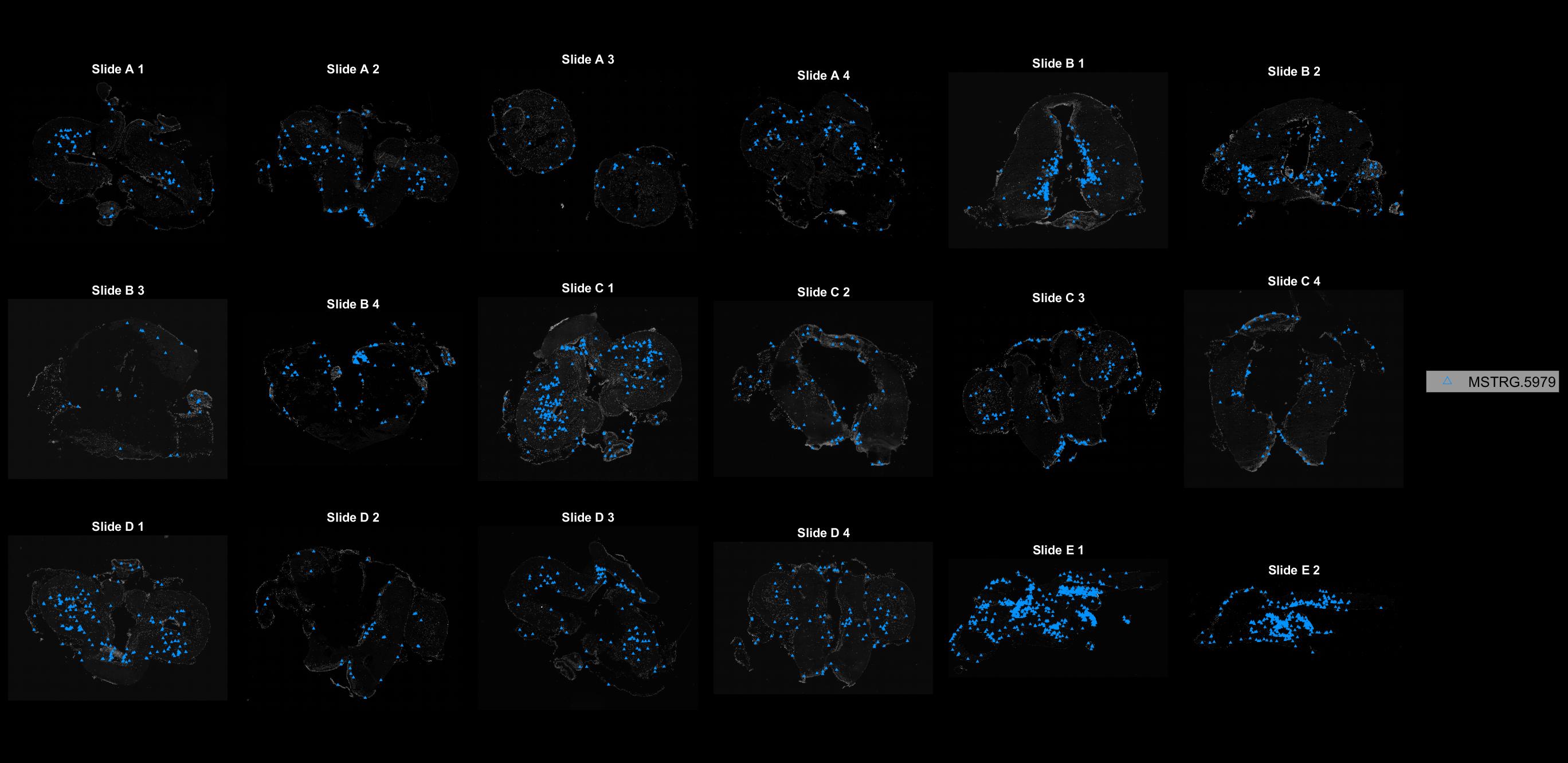

### MSTRG.6042.jpg

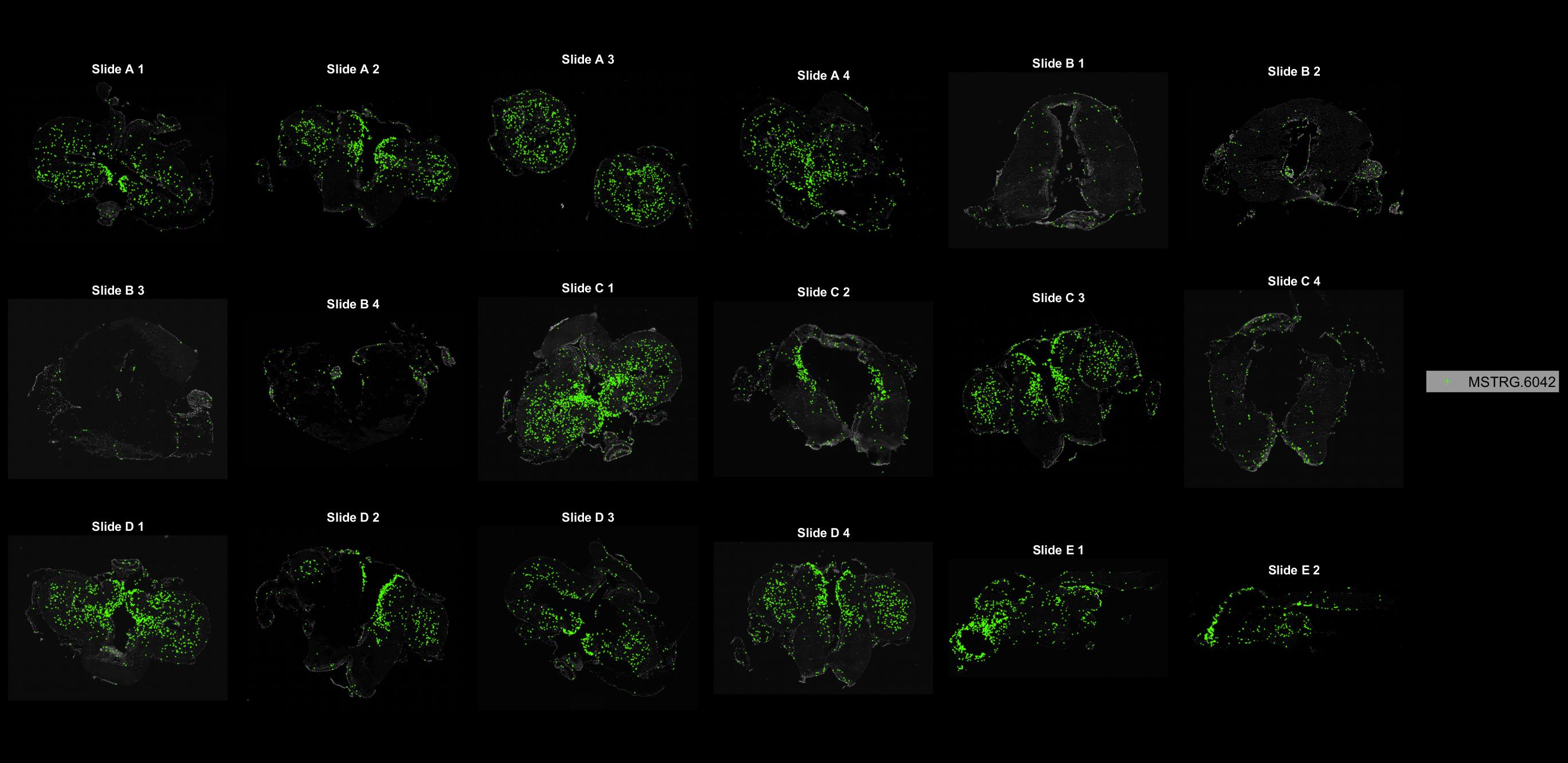

### MSTRG.6070.jpg

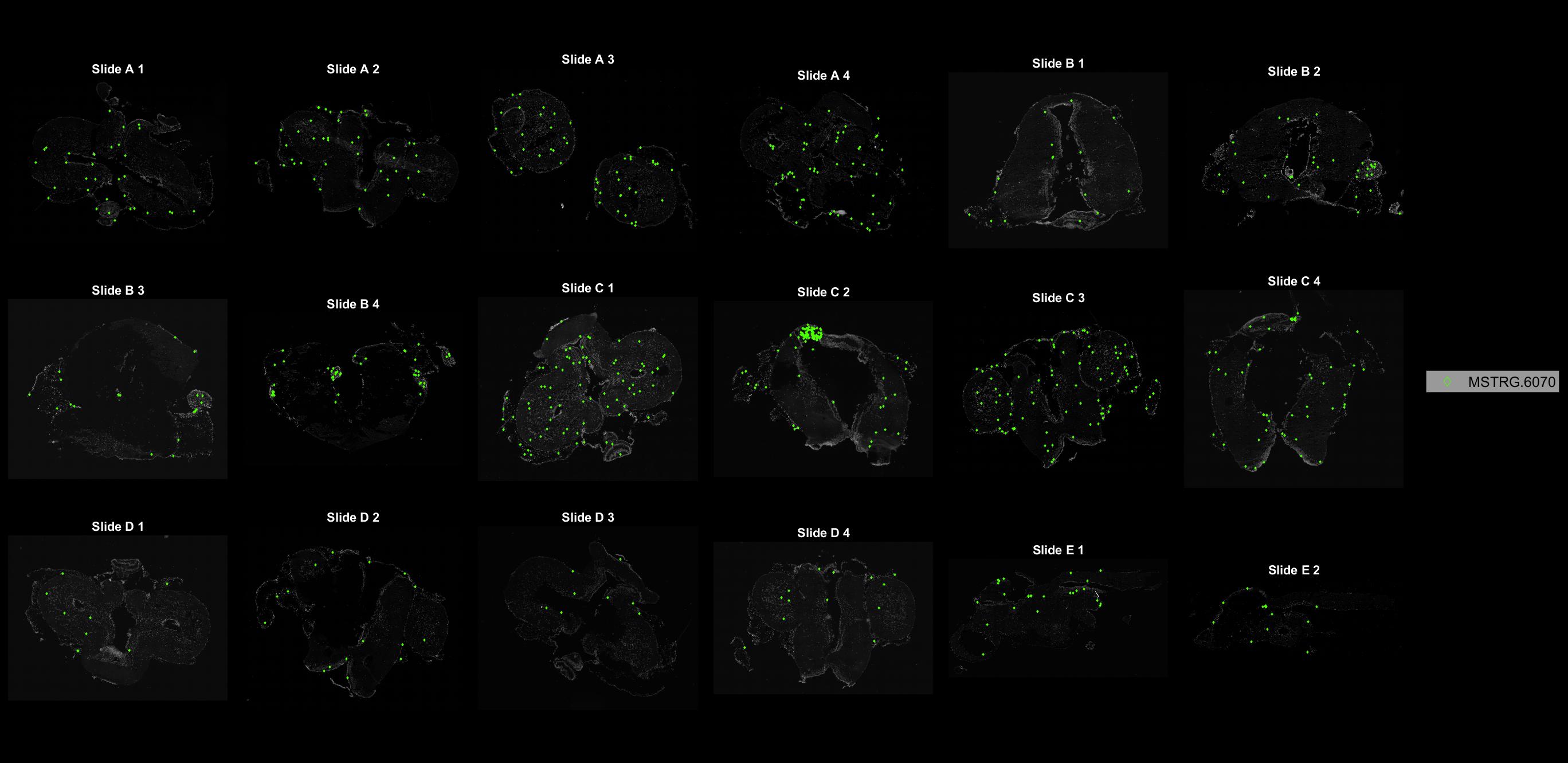

### MSTRG.6108.jpg

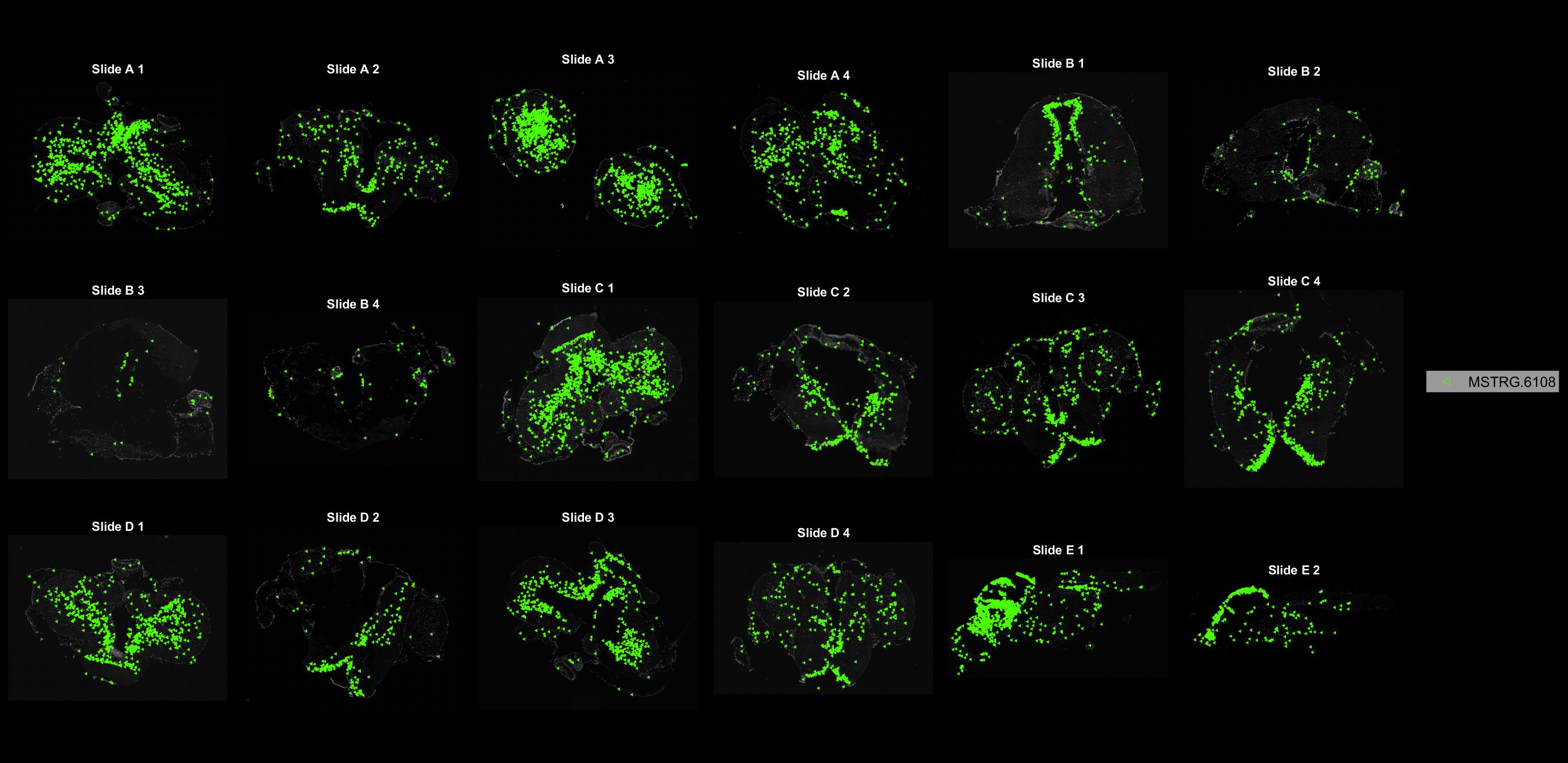

### MSTRG.6109.jpg

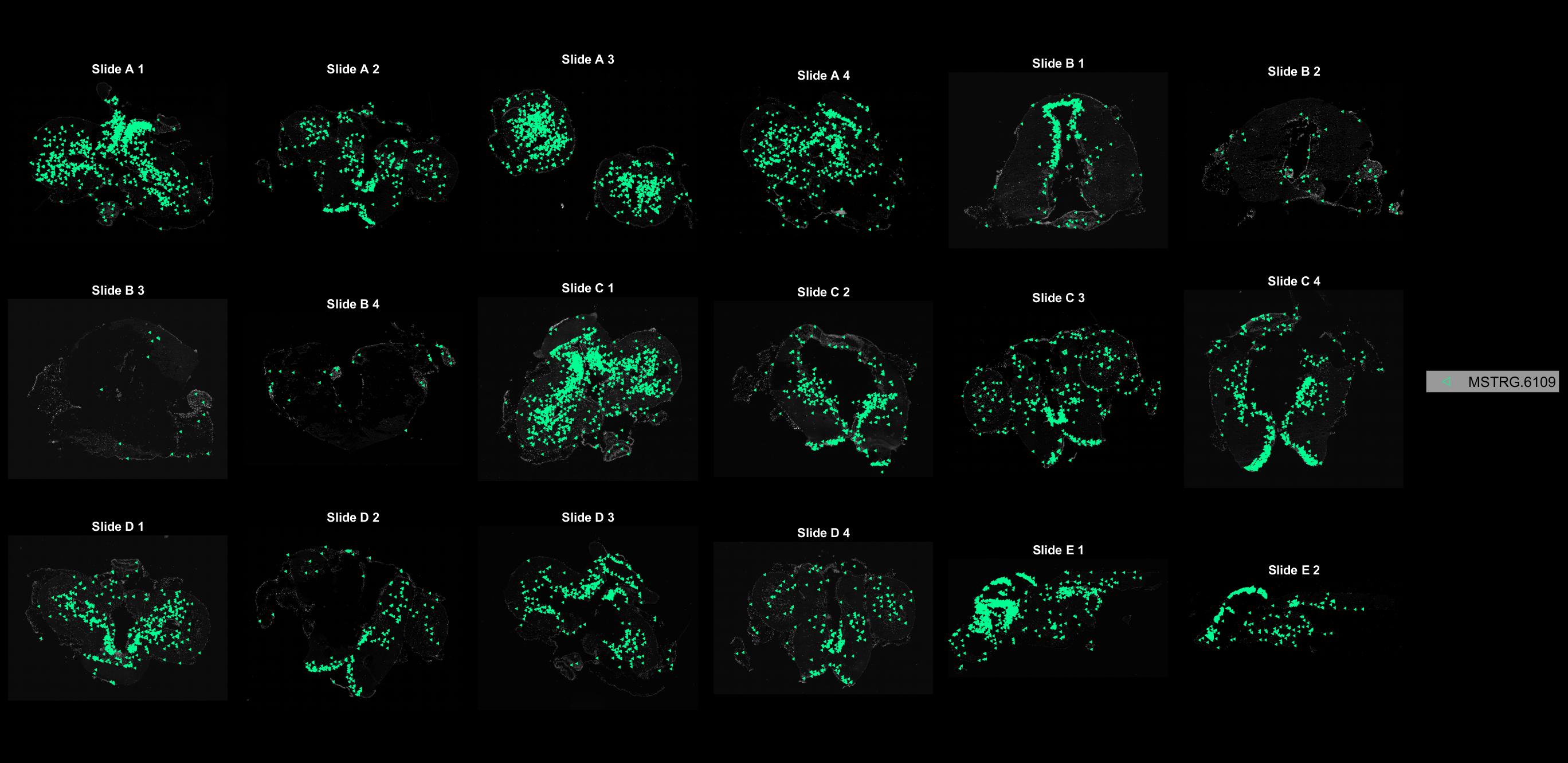

### MSTRG.6115.jpg

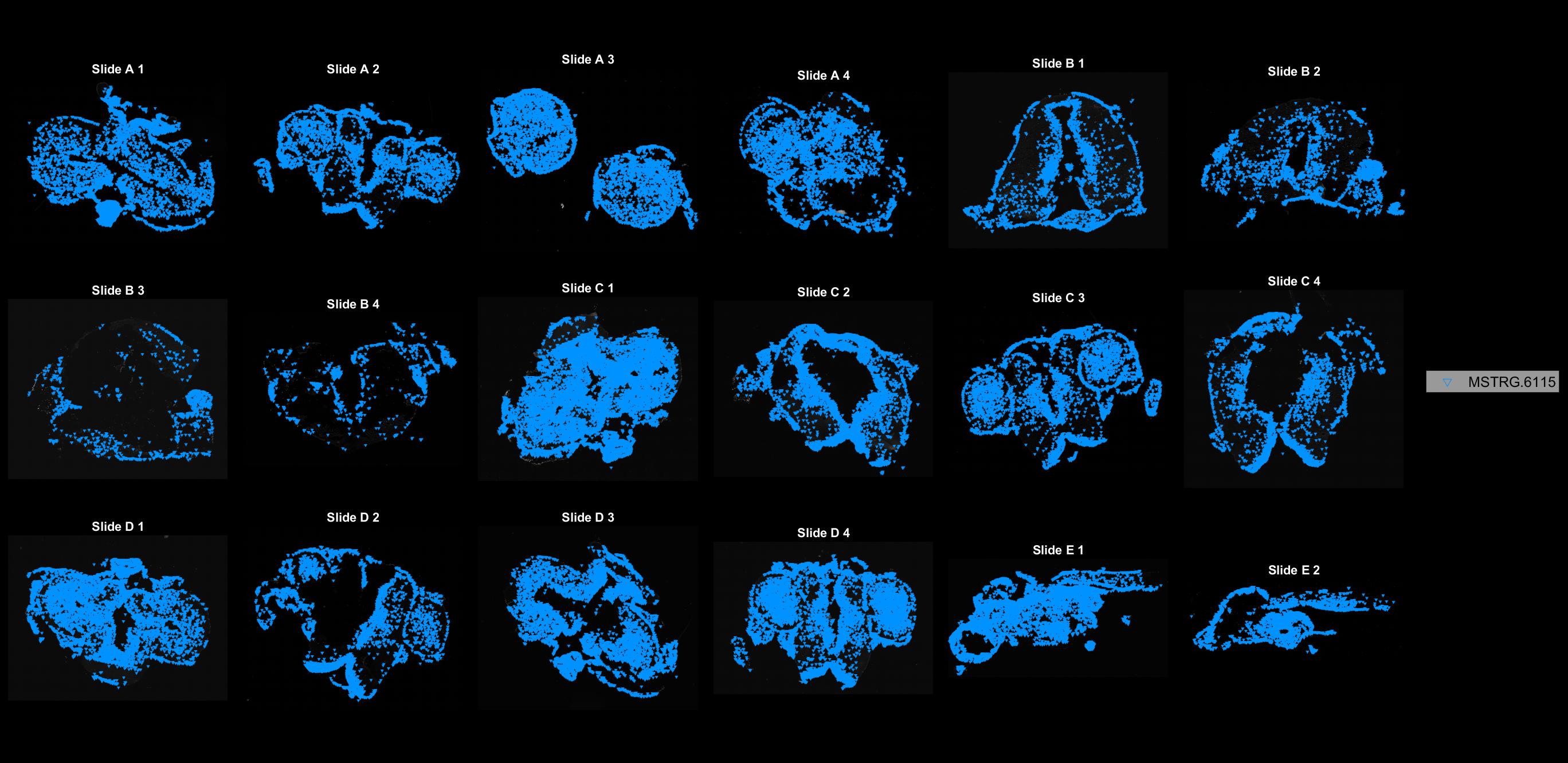

### MSTRG.6365.jpg

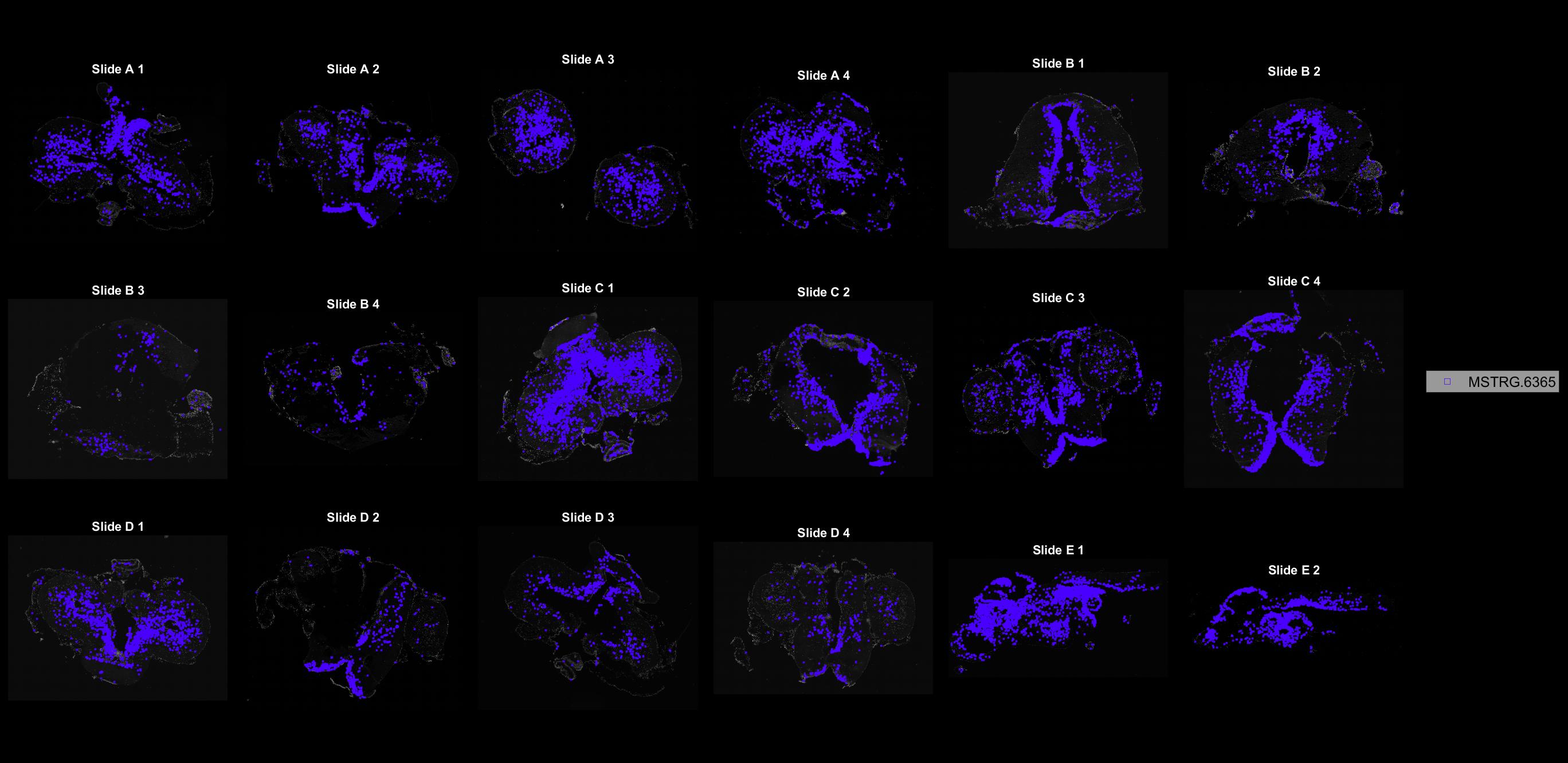

### MSTRG.6410.jpg

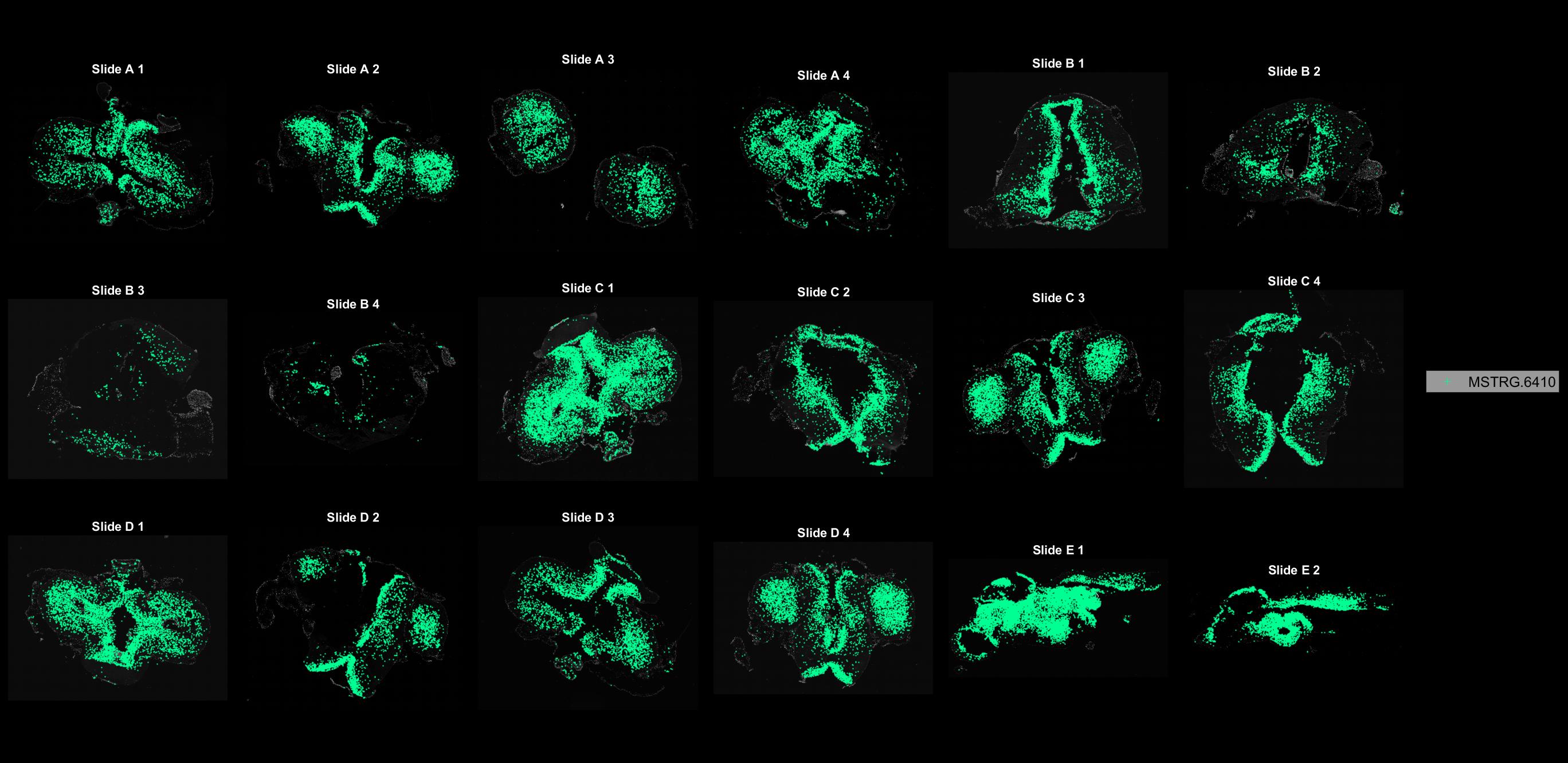

### MSTRG.6549.jpg

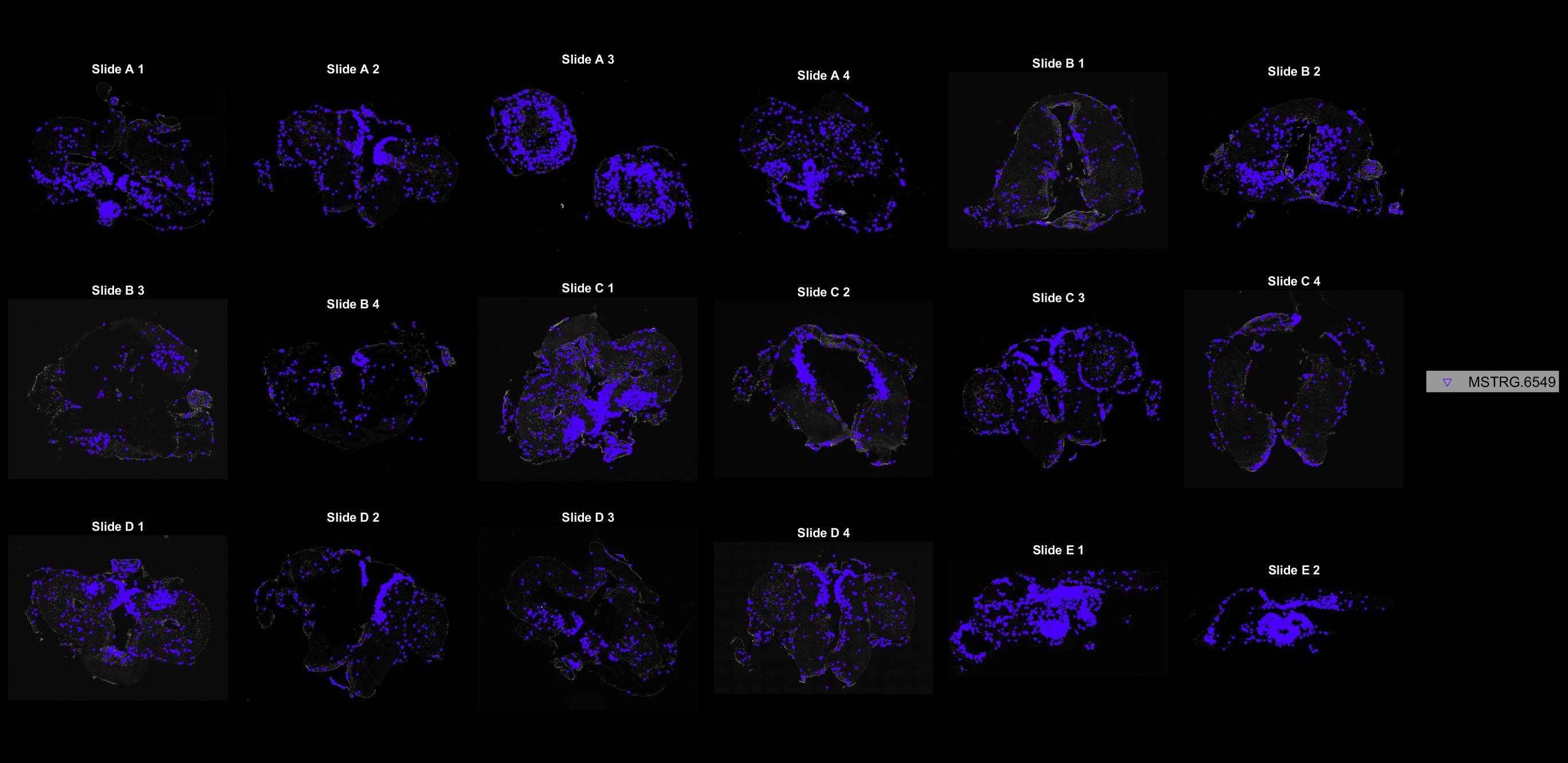

### MSTRG.6582.jpg

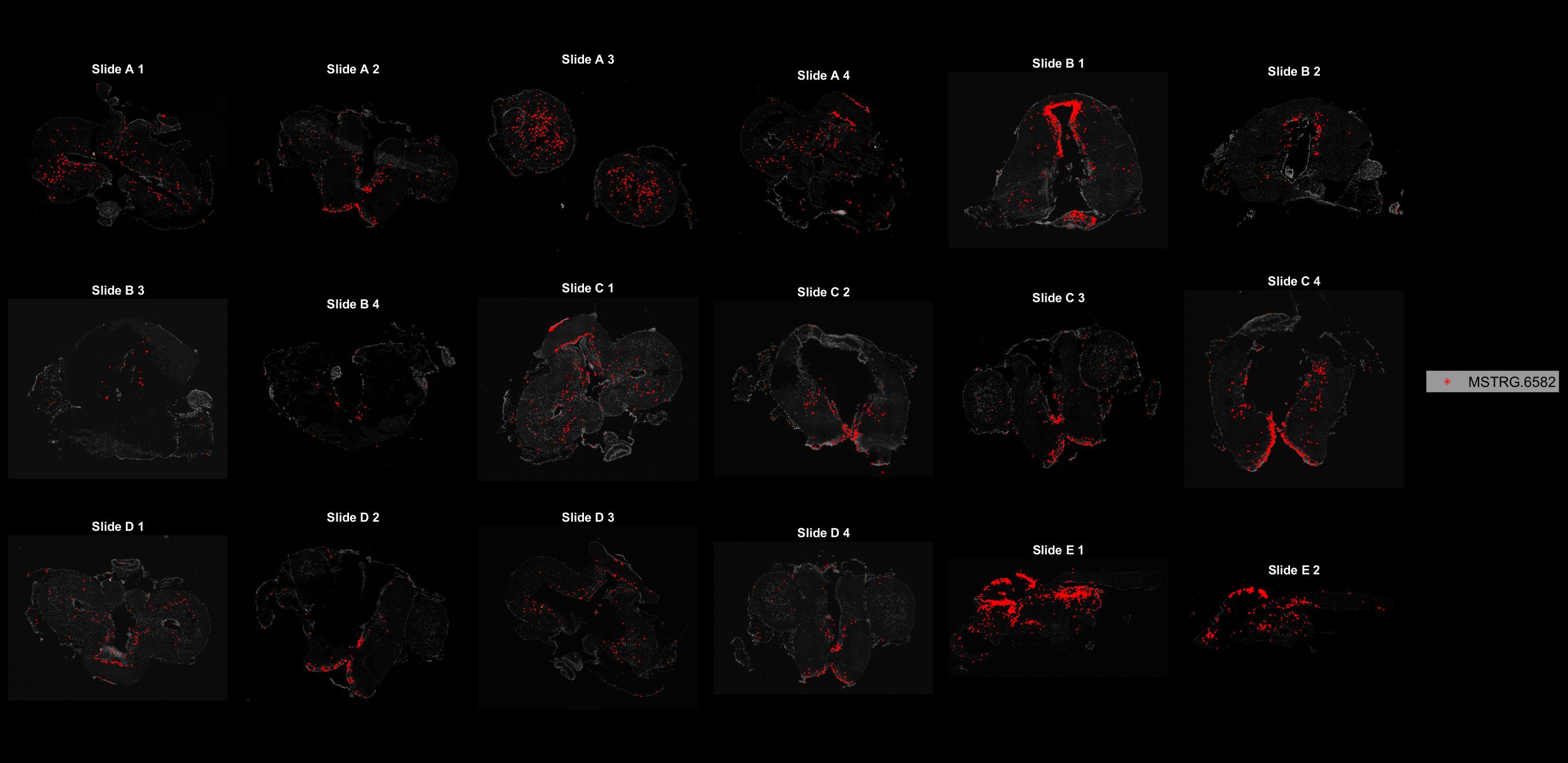

### MSTRG.6588.jpg

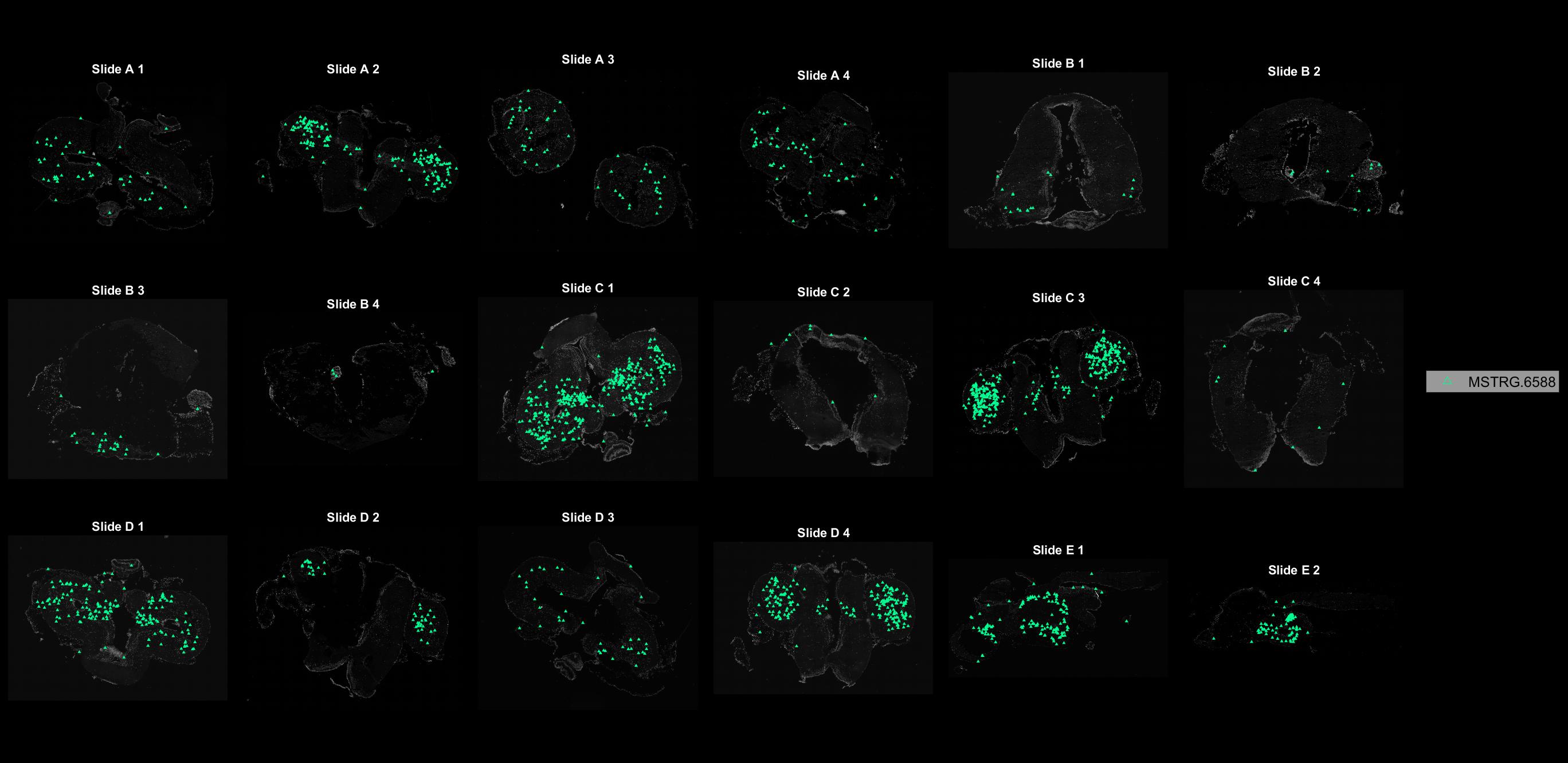

### MSTRG.6637.jpg

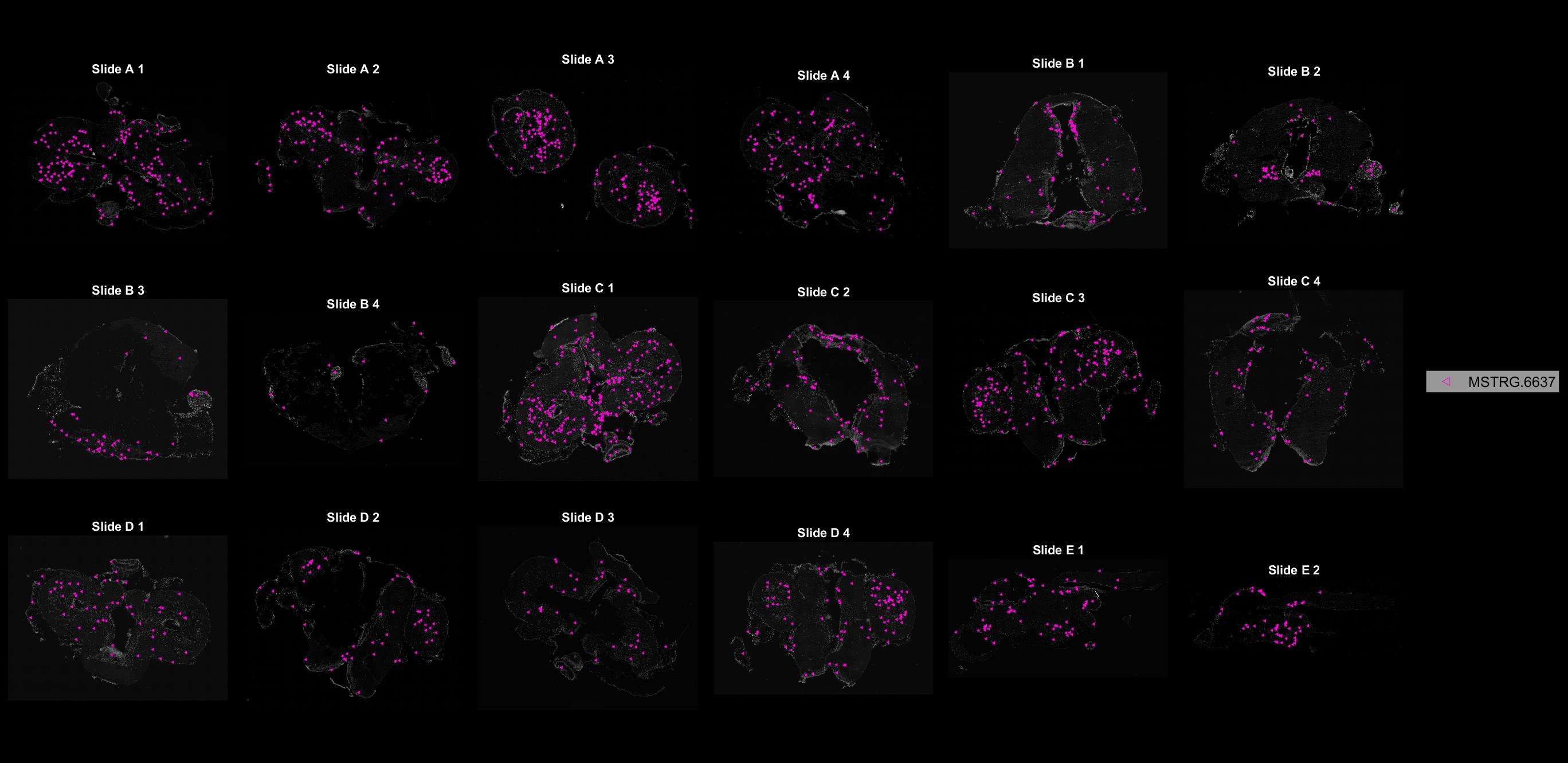

### MSTRG.6758.jpg

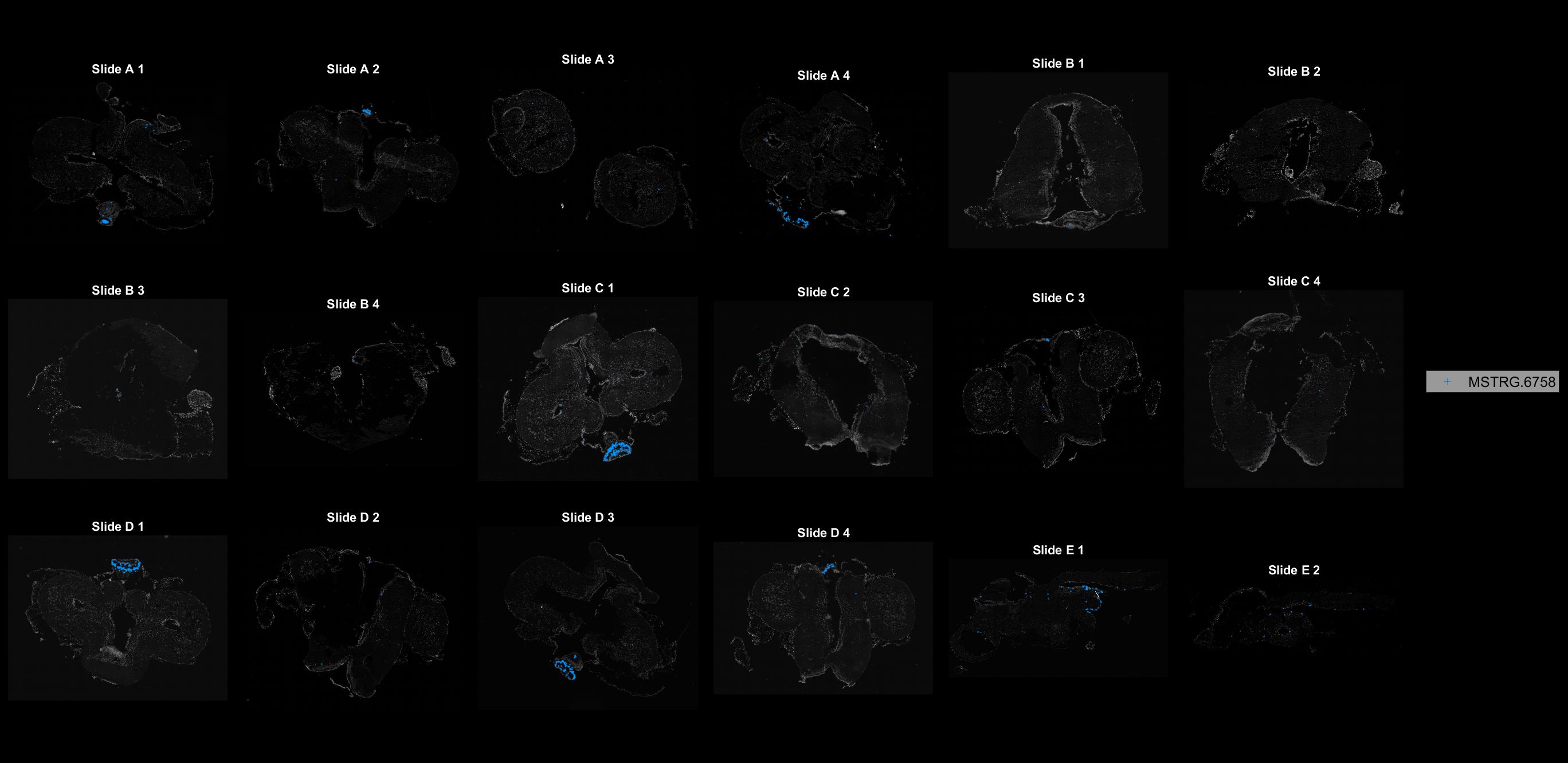

### MSTRG.6835.jpg

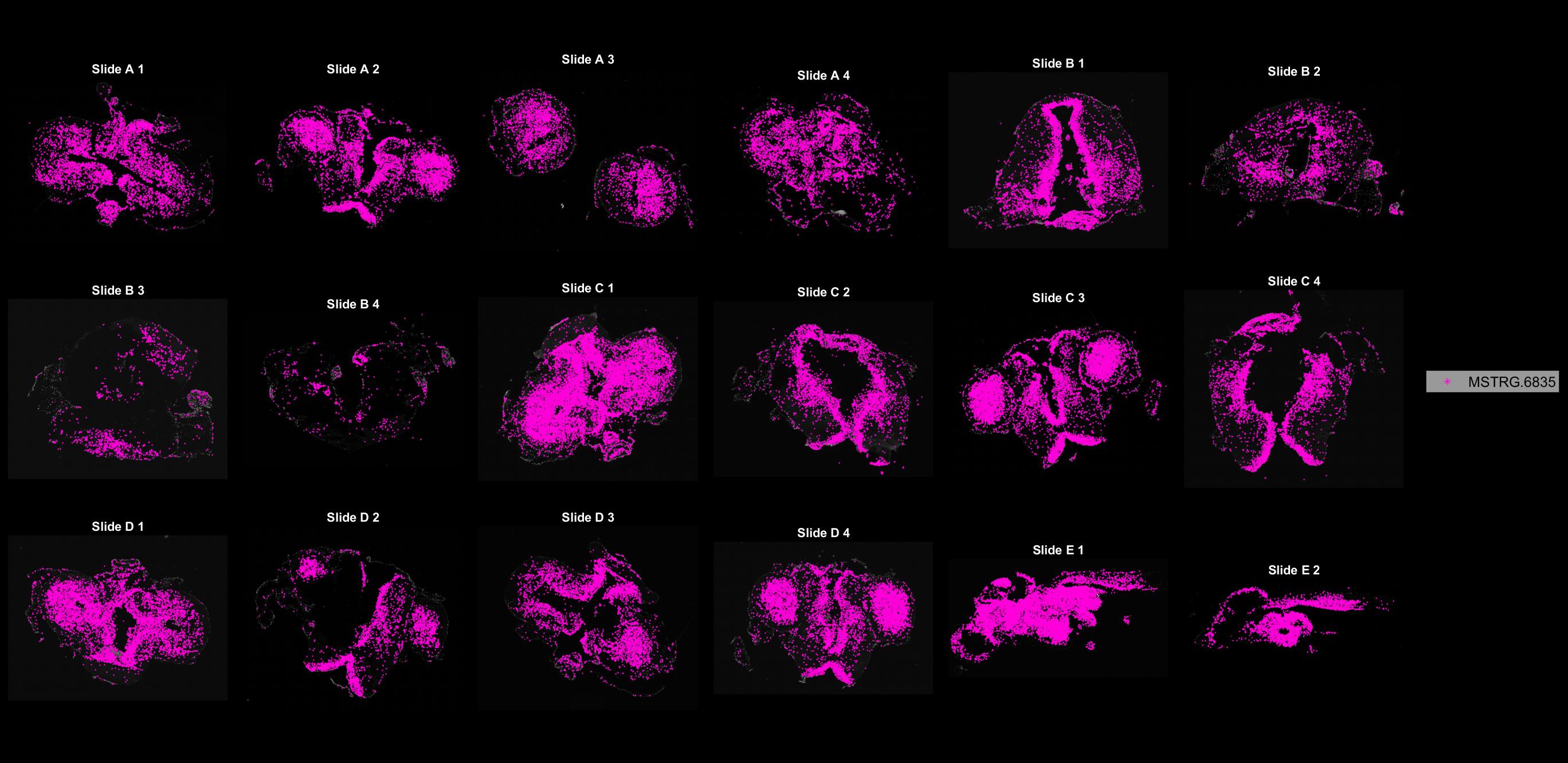

### MSTRG.6852.jpg

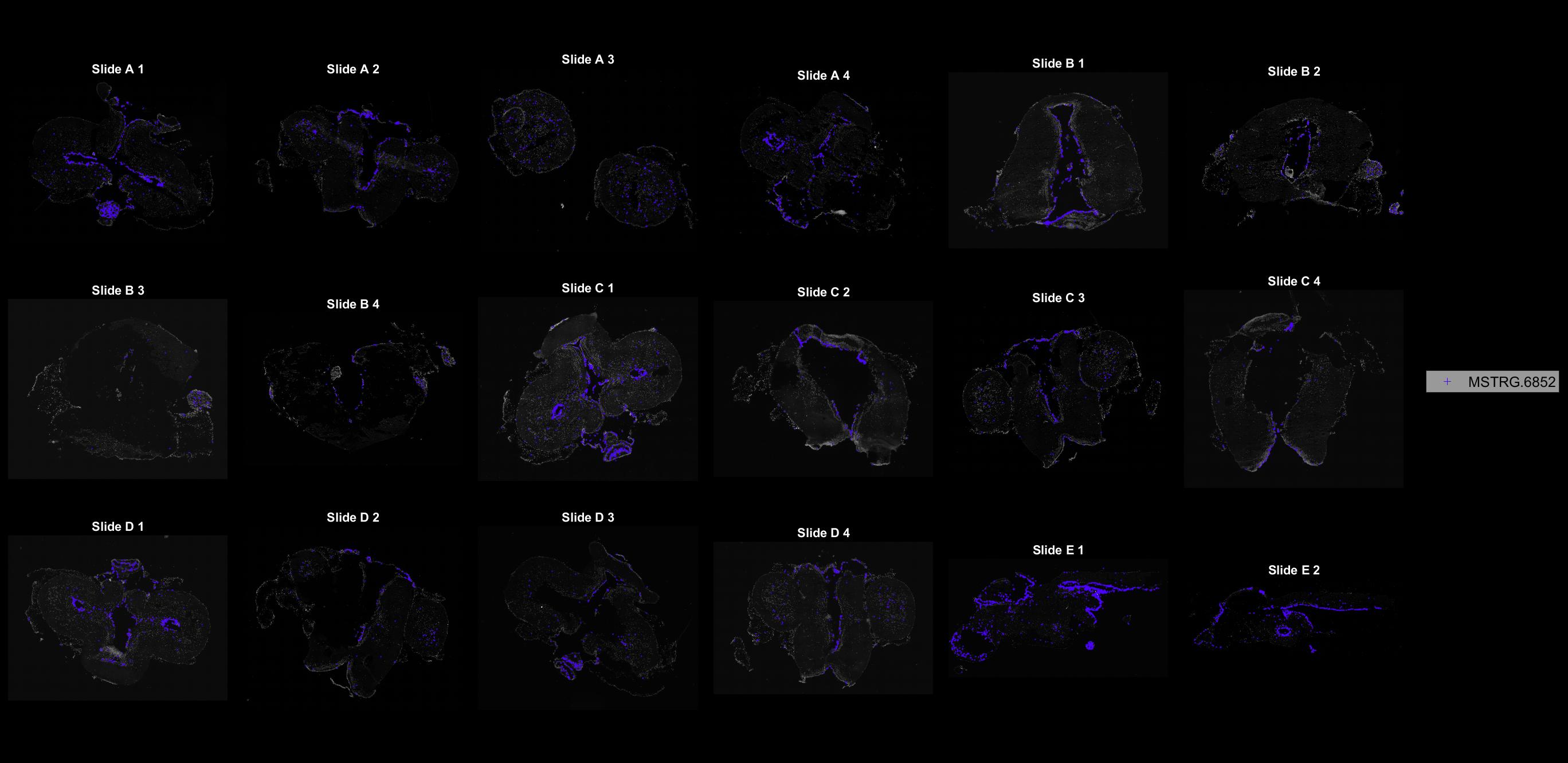

### MSTRG.7030.jpg

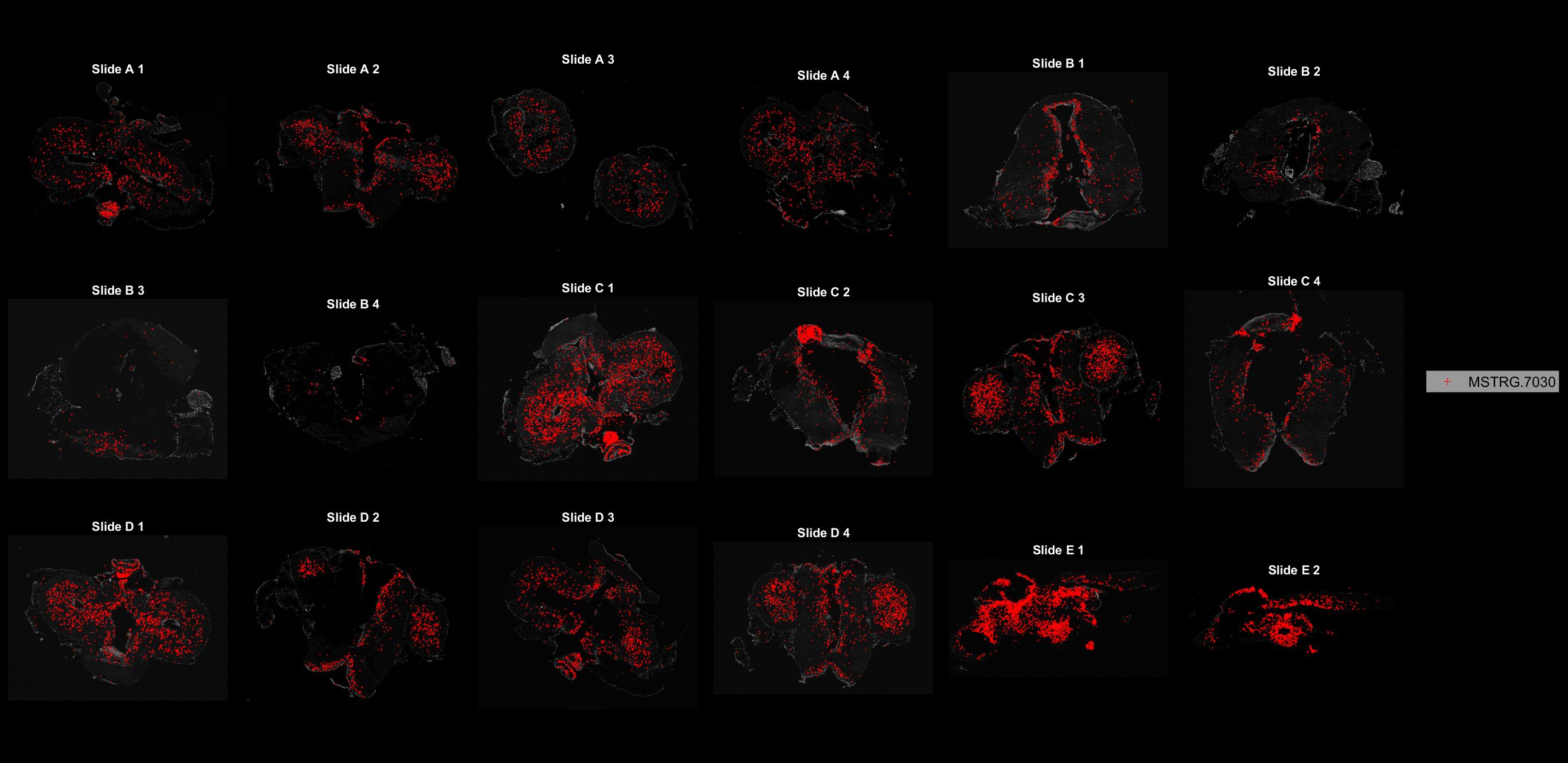

### MSTRG.7309.jpg

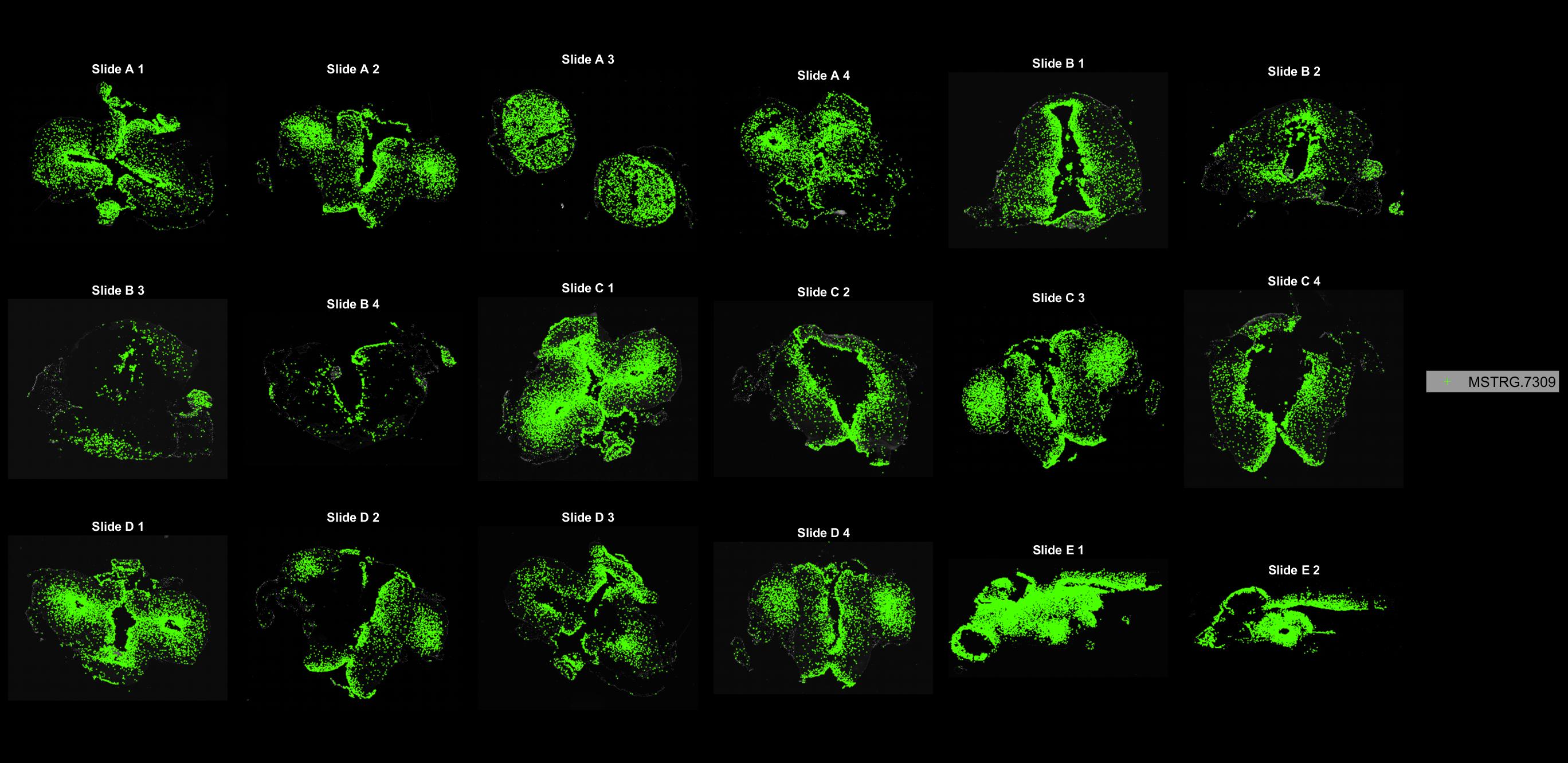

### MSTRG.7780.jpg

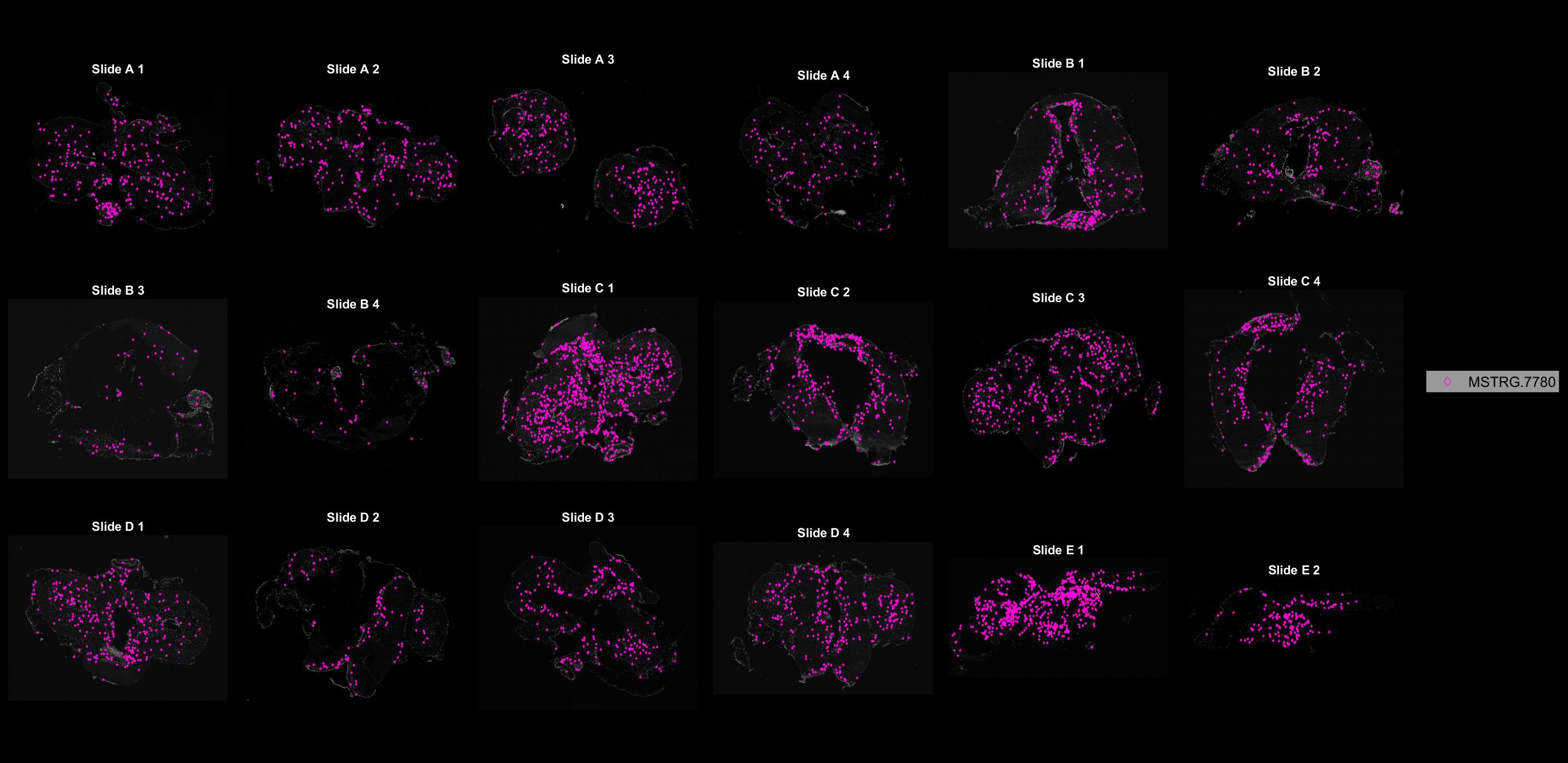

### MSTRG.7934.jpg

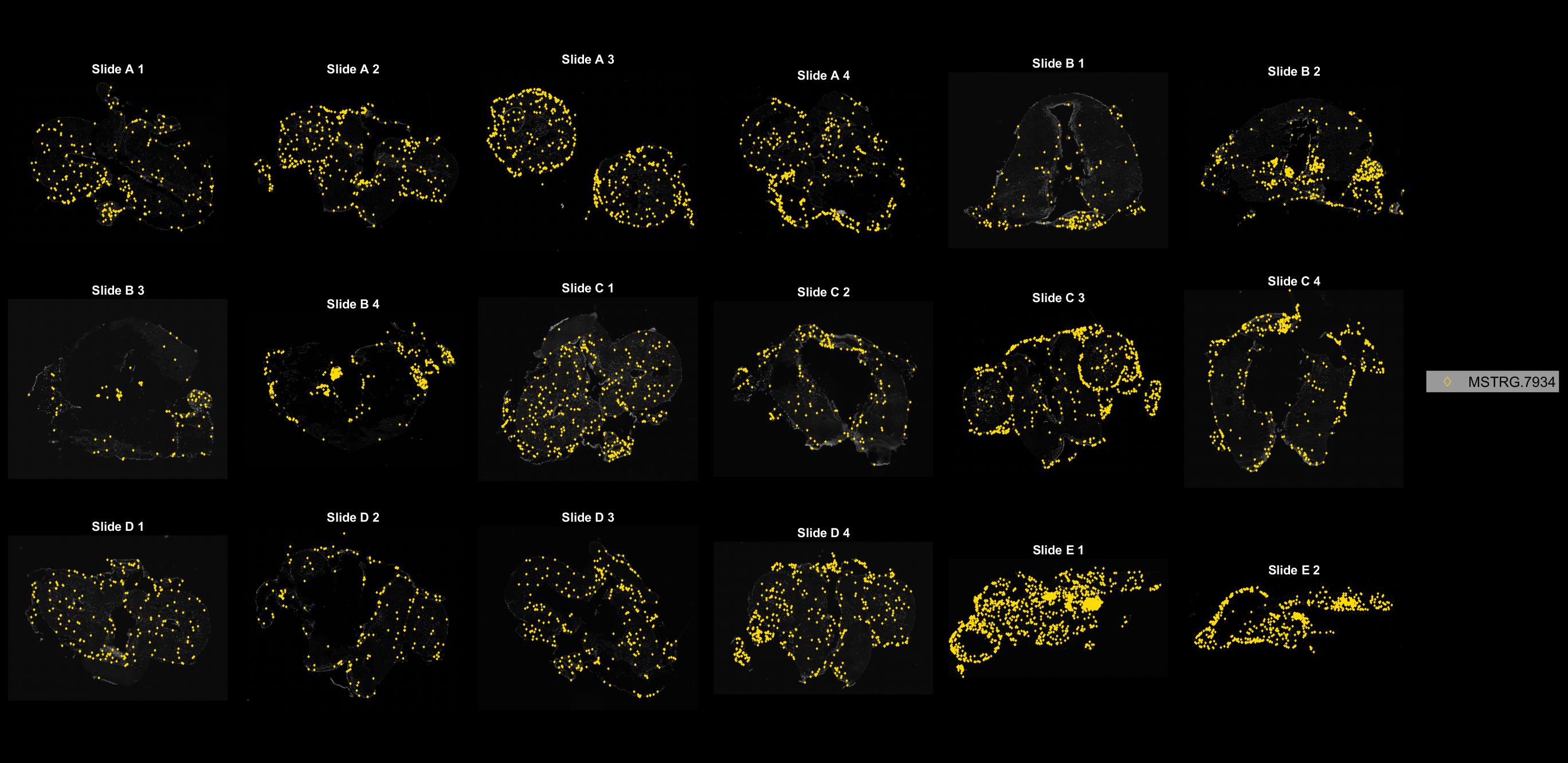

### MSTRG.8275.jpg

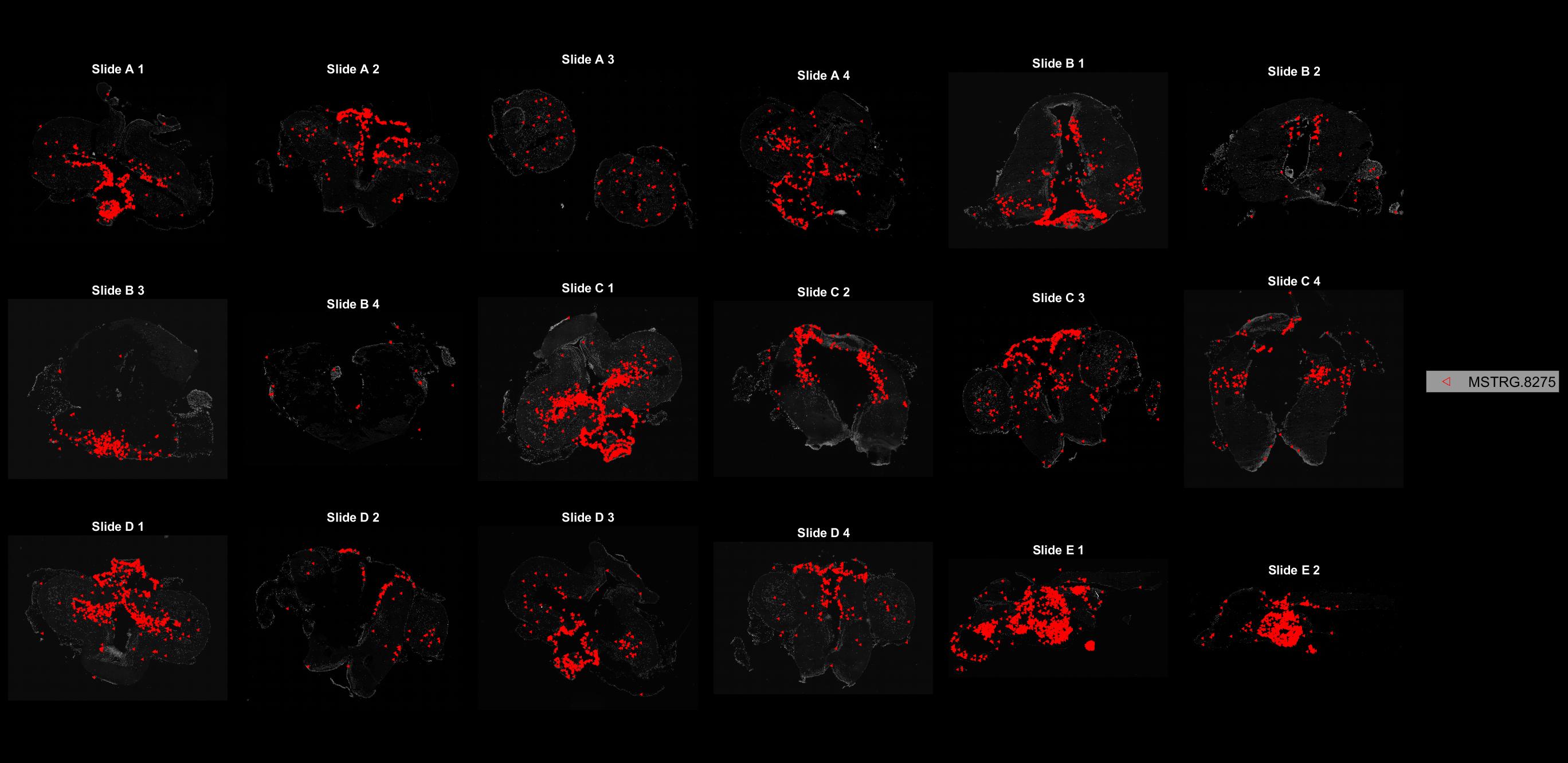

### MSTRG.8398.jpg

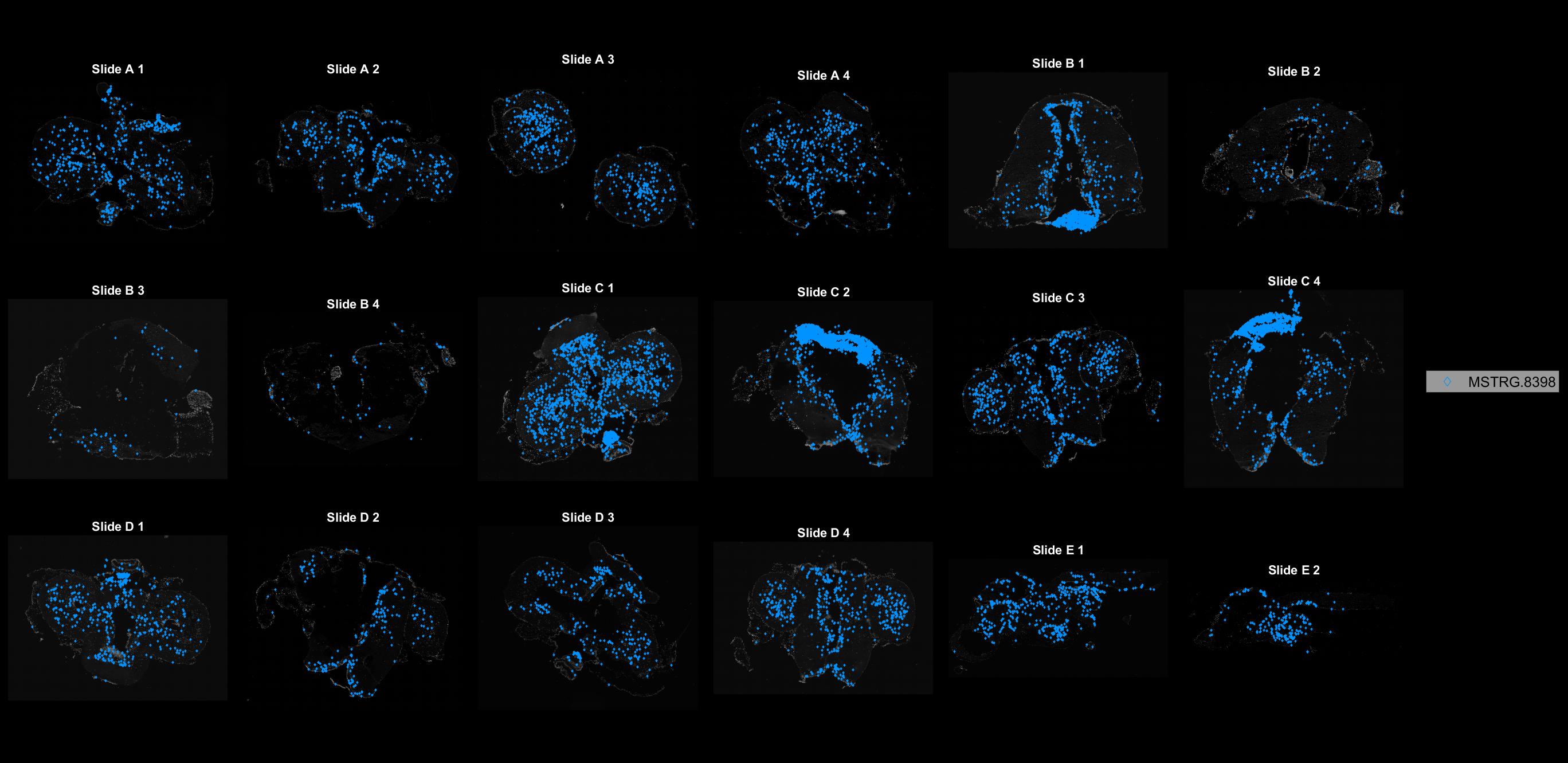

### MSTRG.8489.jpg

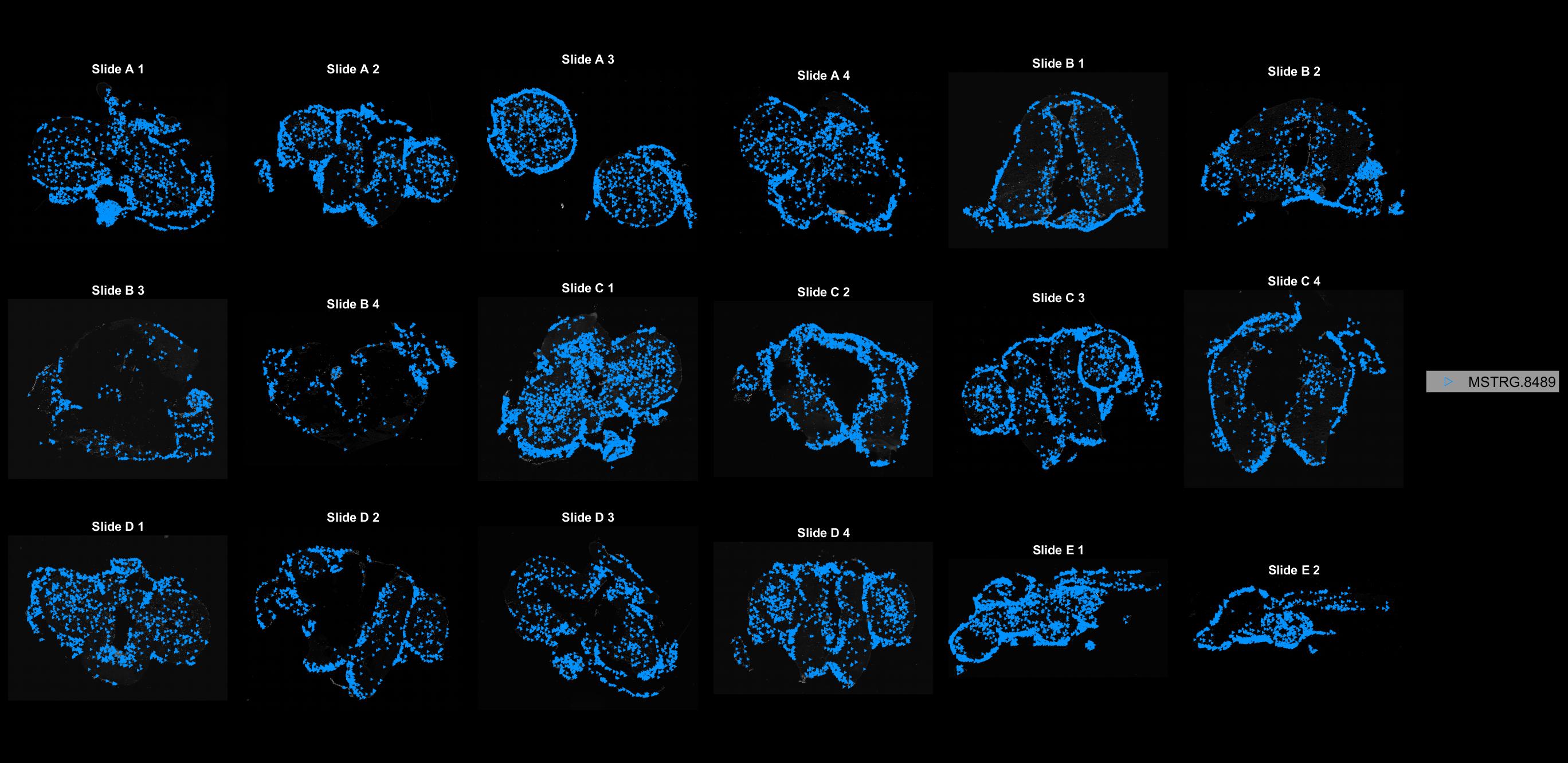

### MSTRG.8560.jpg

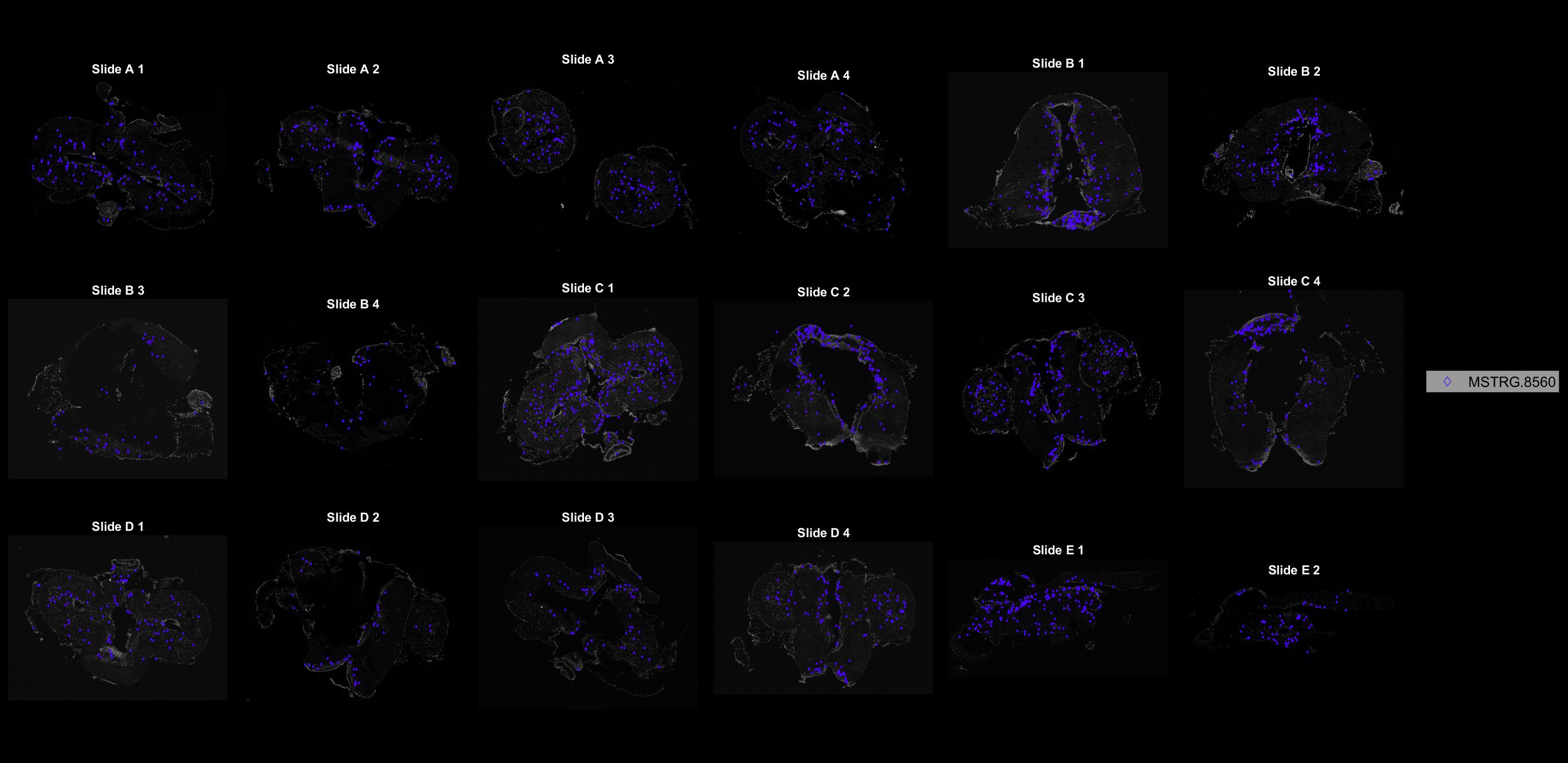

### MSTRG.8862.jpg

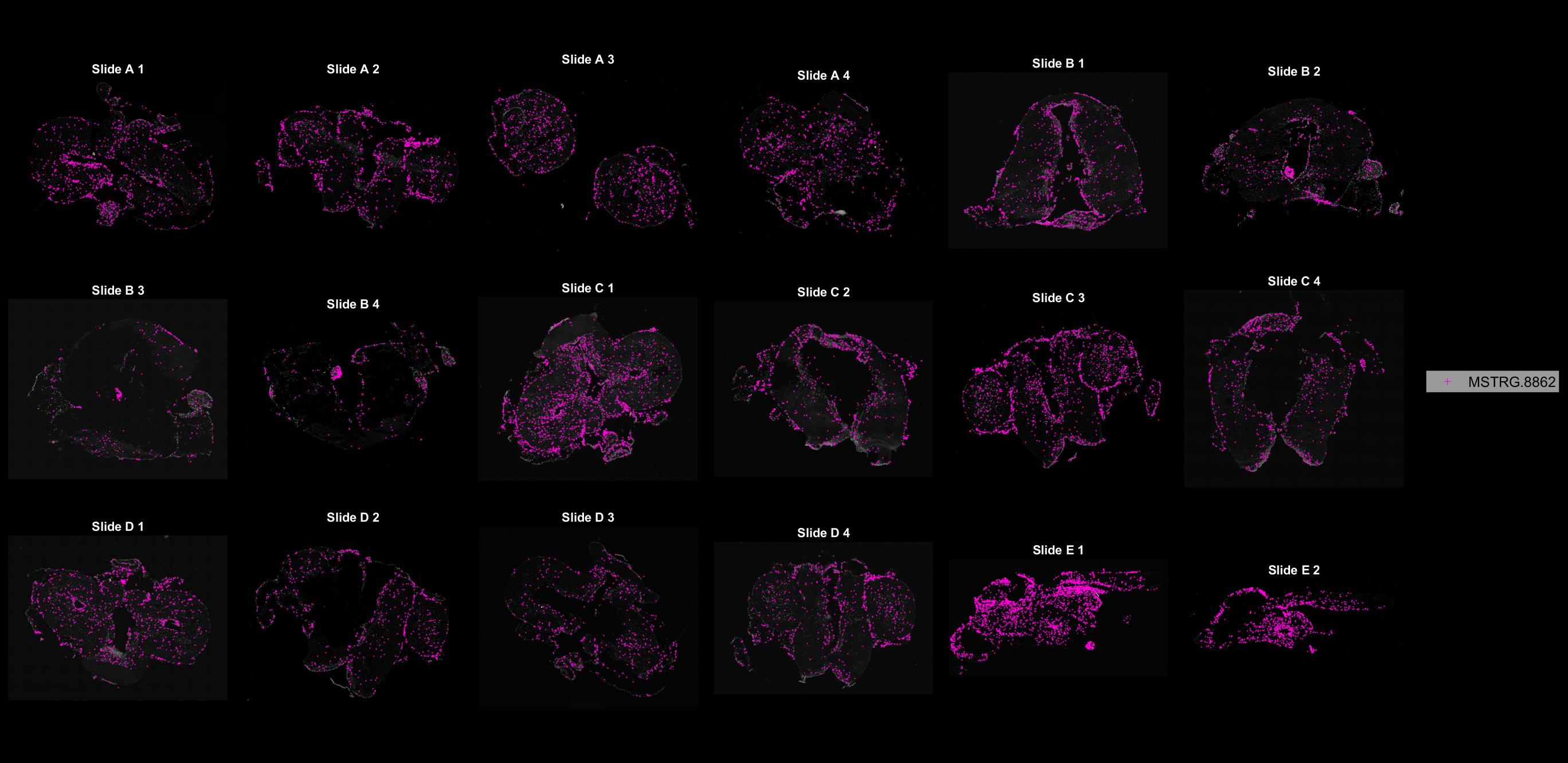

### MSTRG.9098.jpg

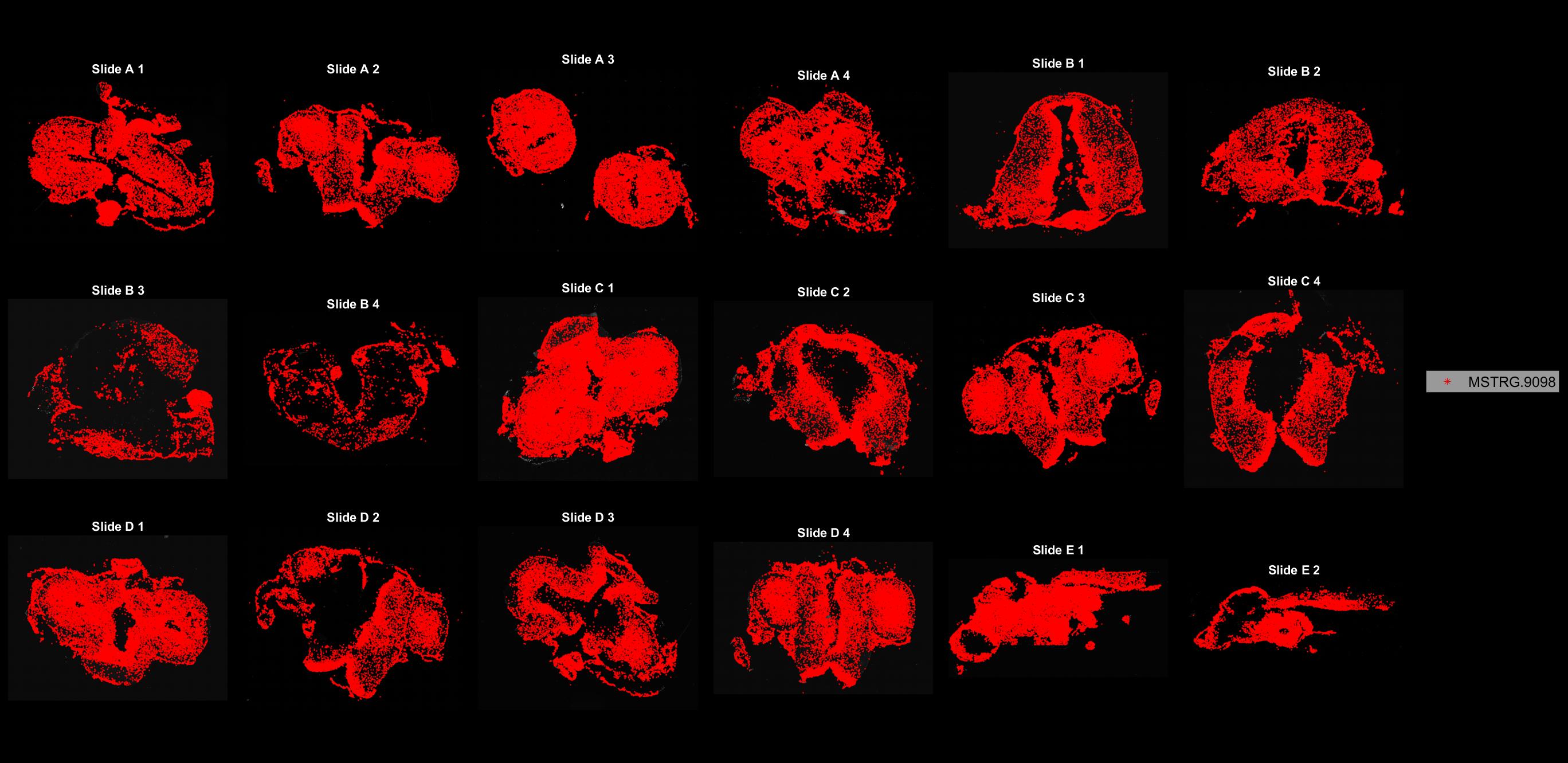

### MSTRG.9450.jpg

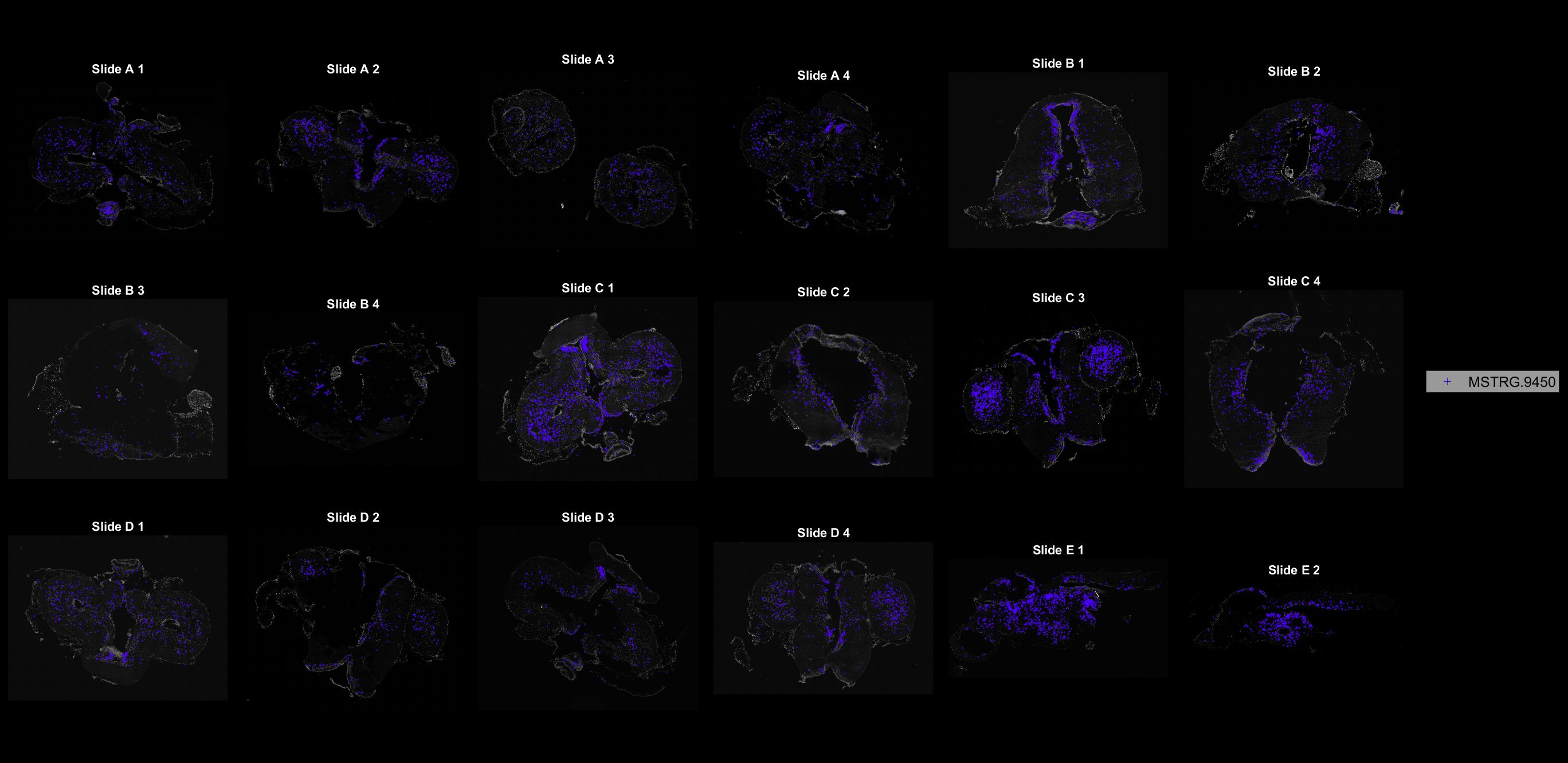

### MSTRG.9797.jpg

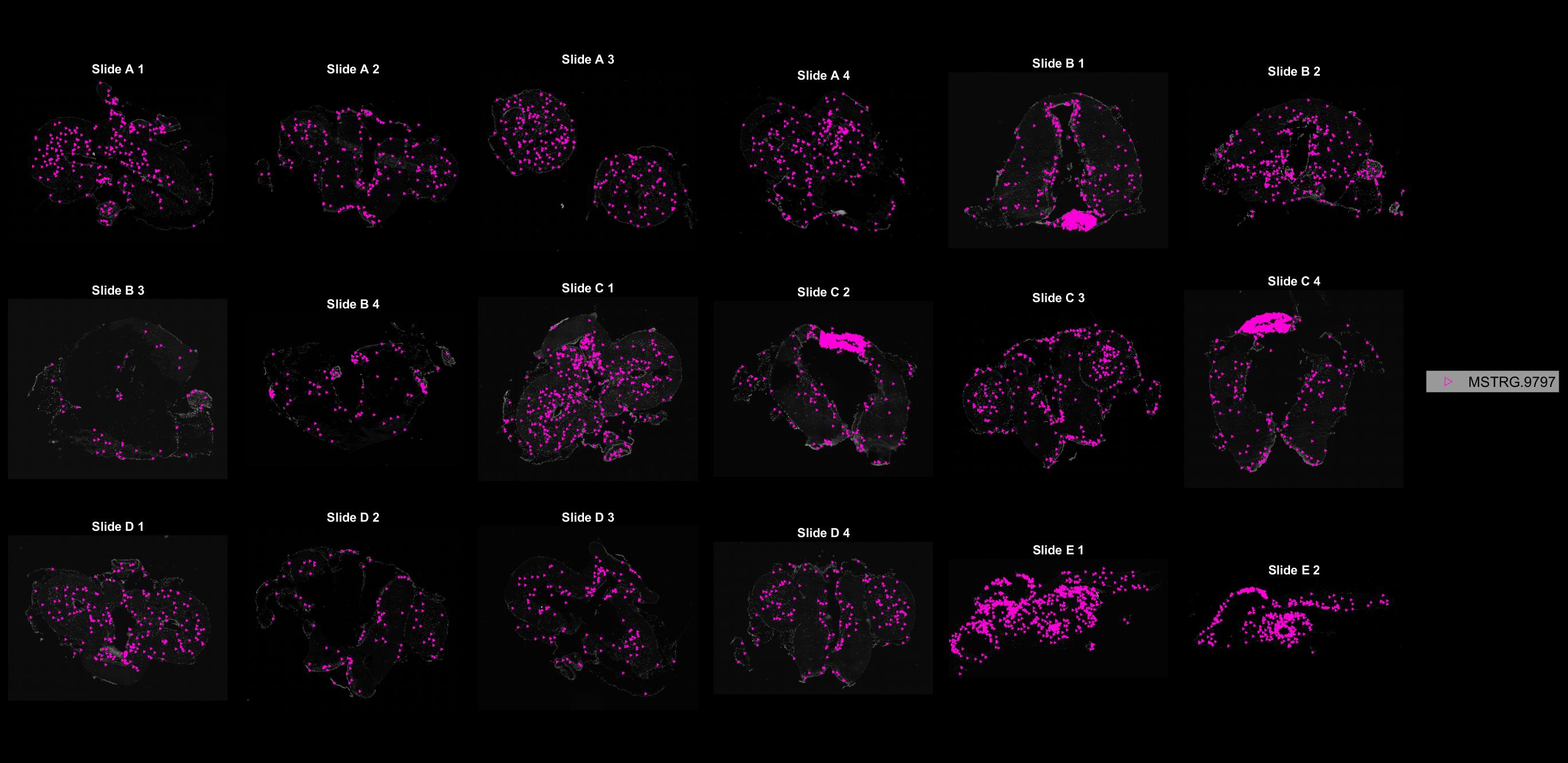

### MSTRG.9839.jpg

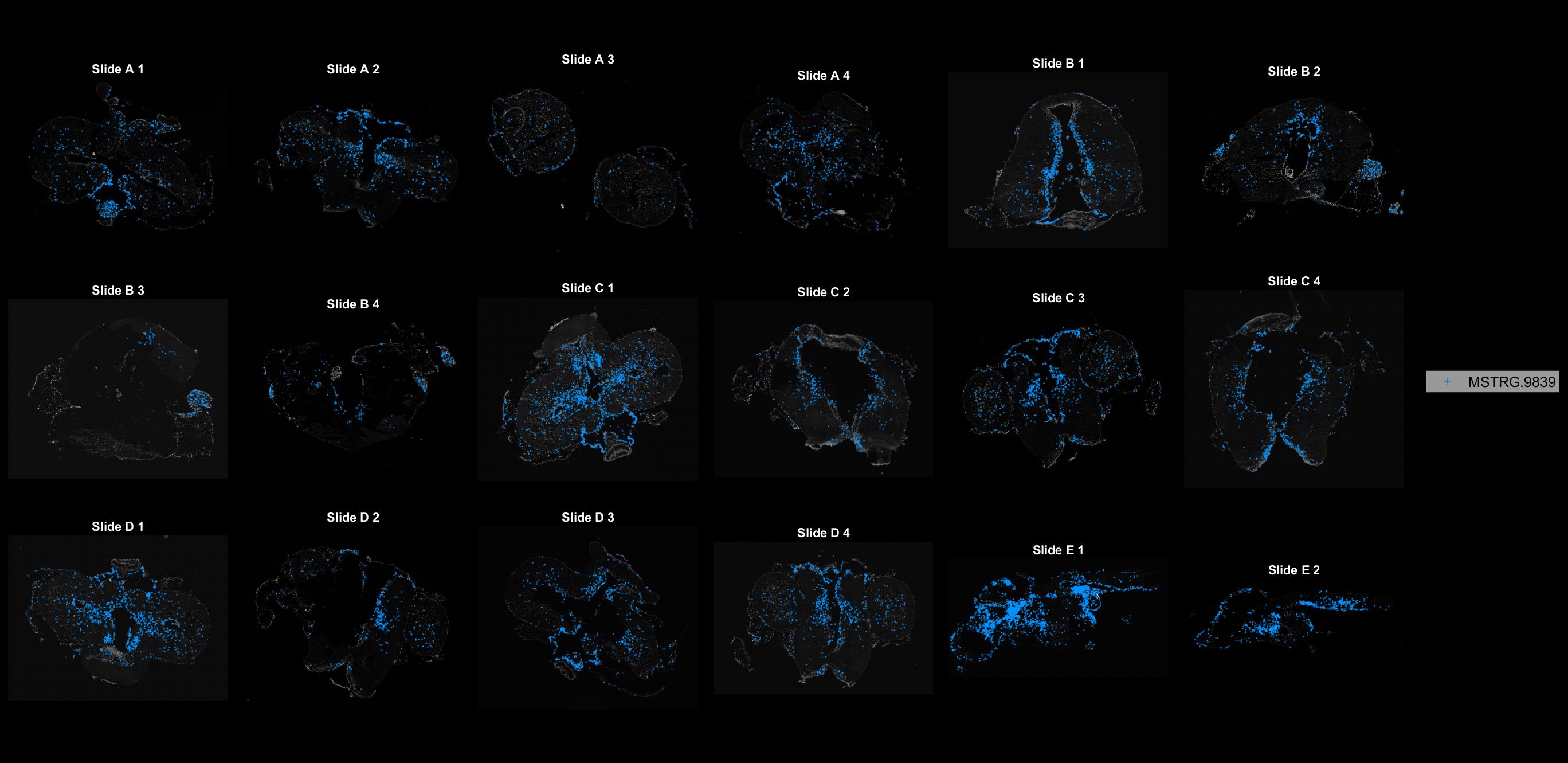

### MSTRG.10404.jpg

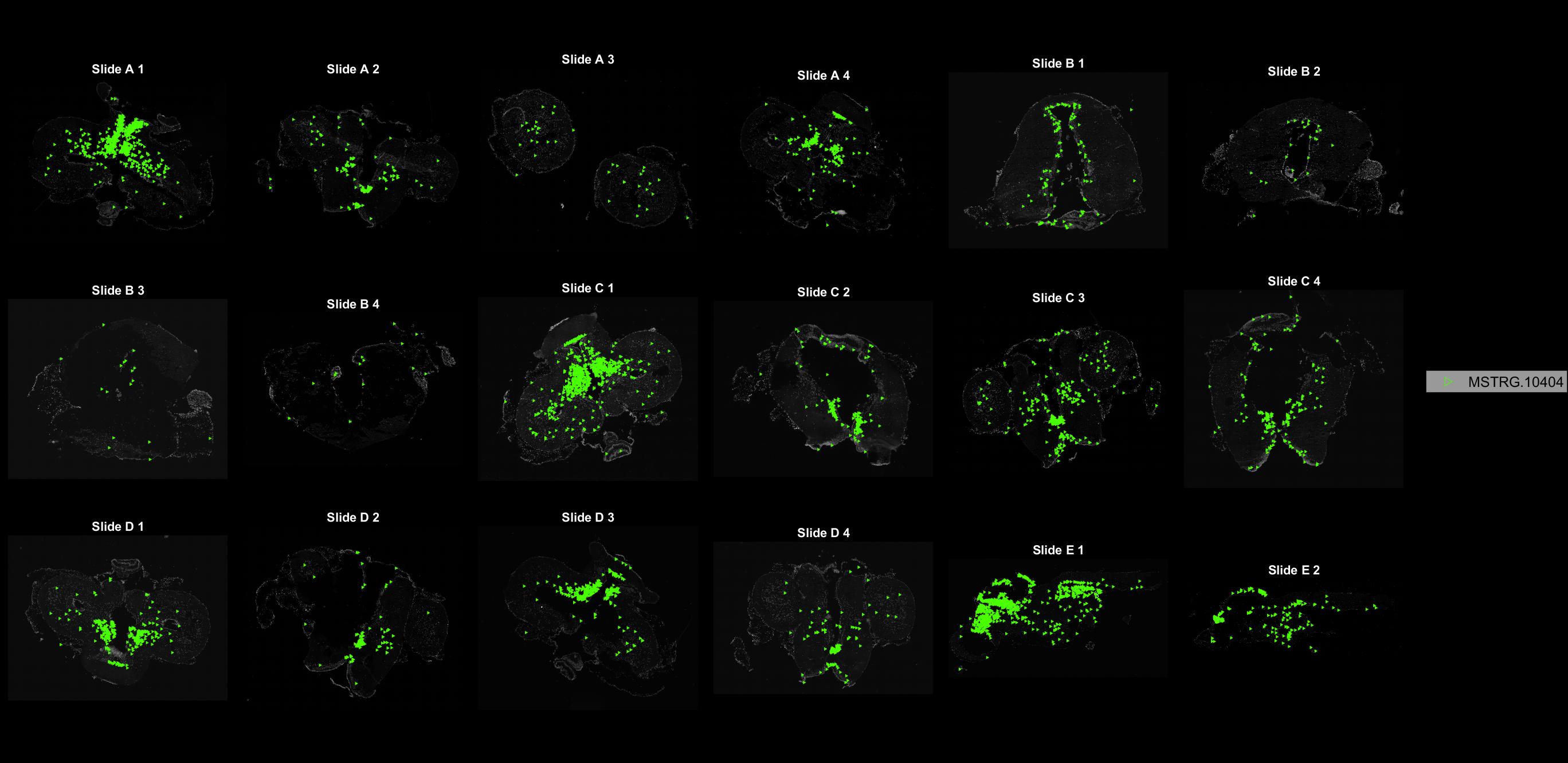

### MSTRG.10634.jpg

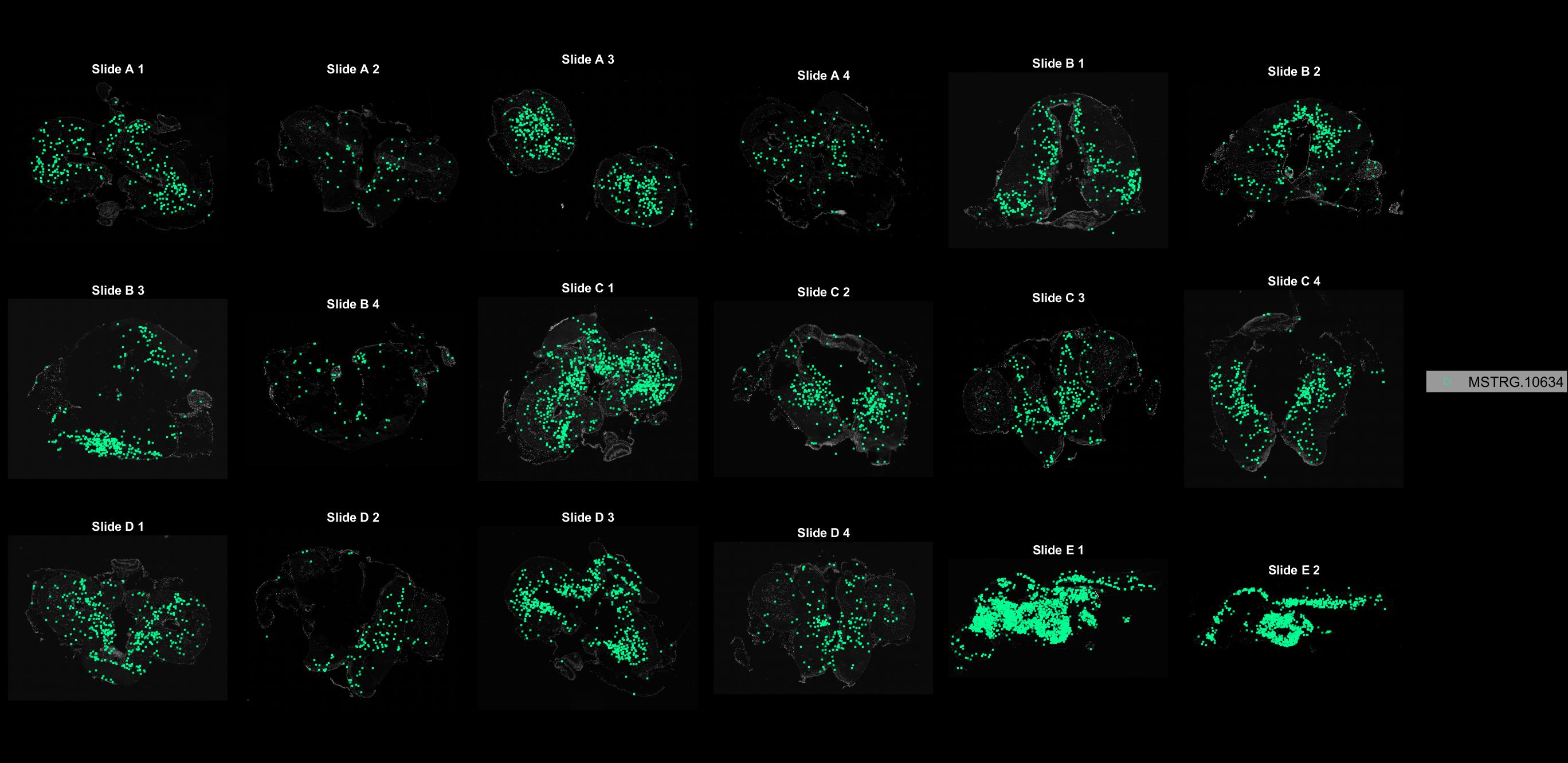

### MSTRG.11404.jpg

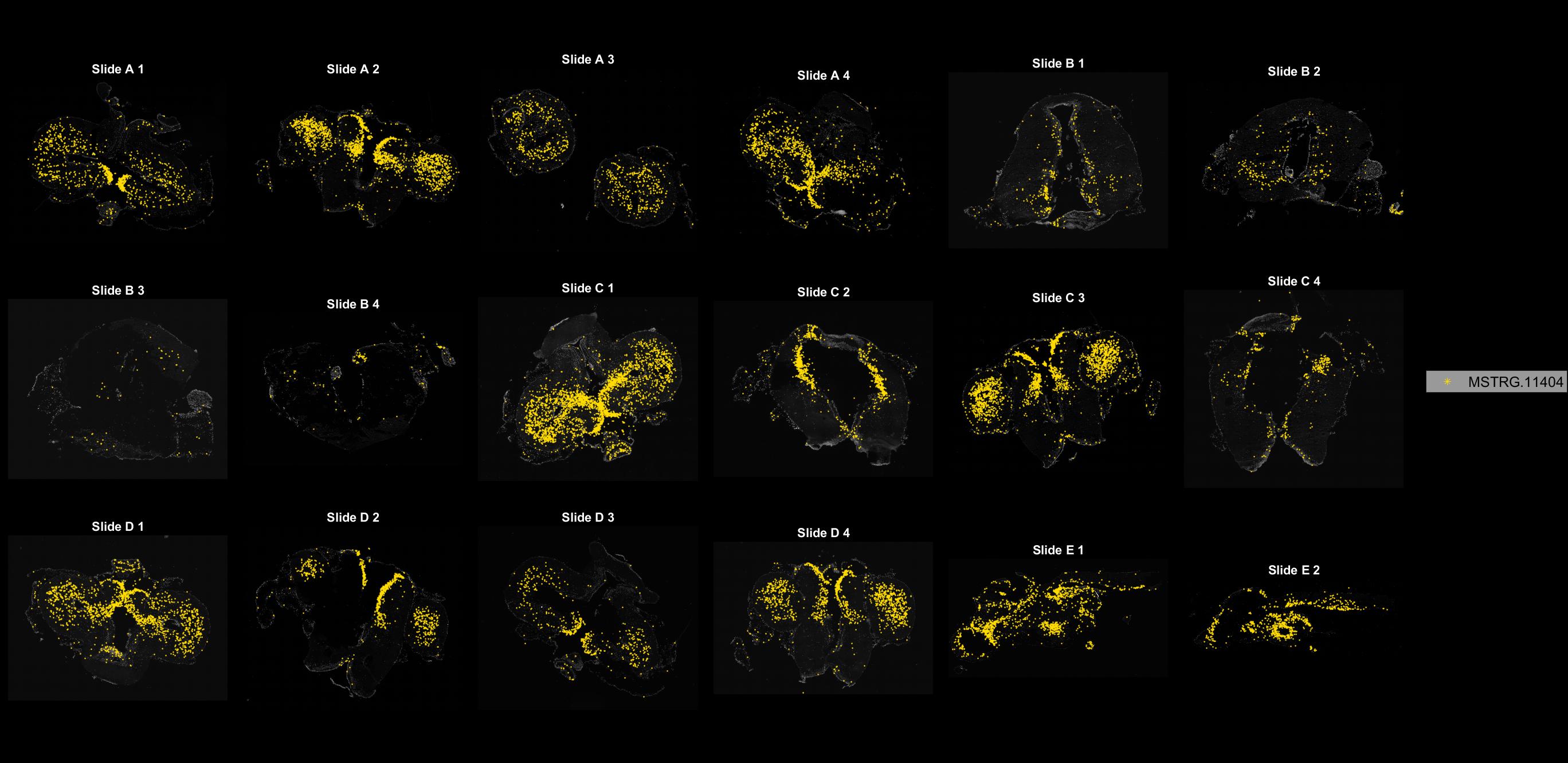

### MSTRG.11808.jpg

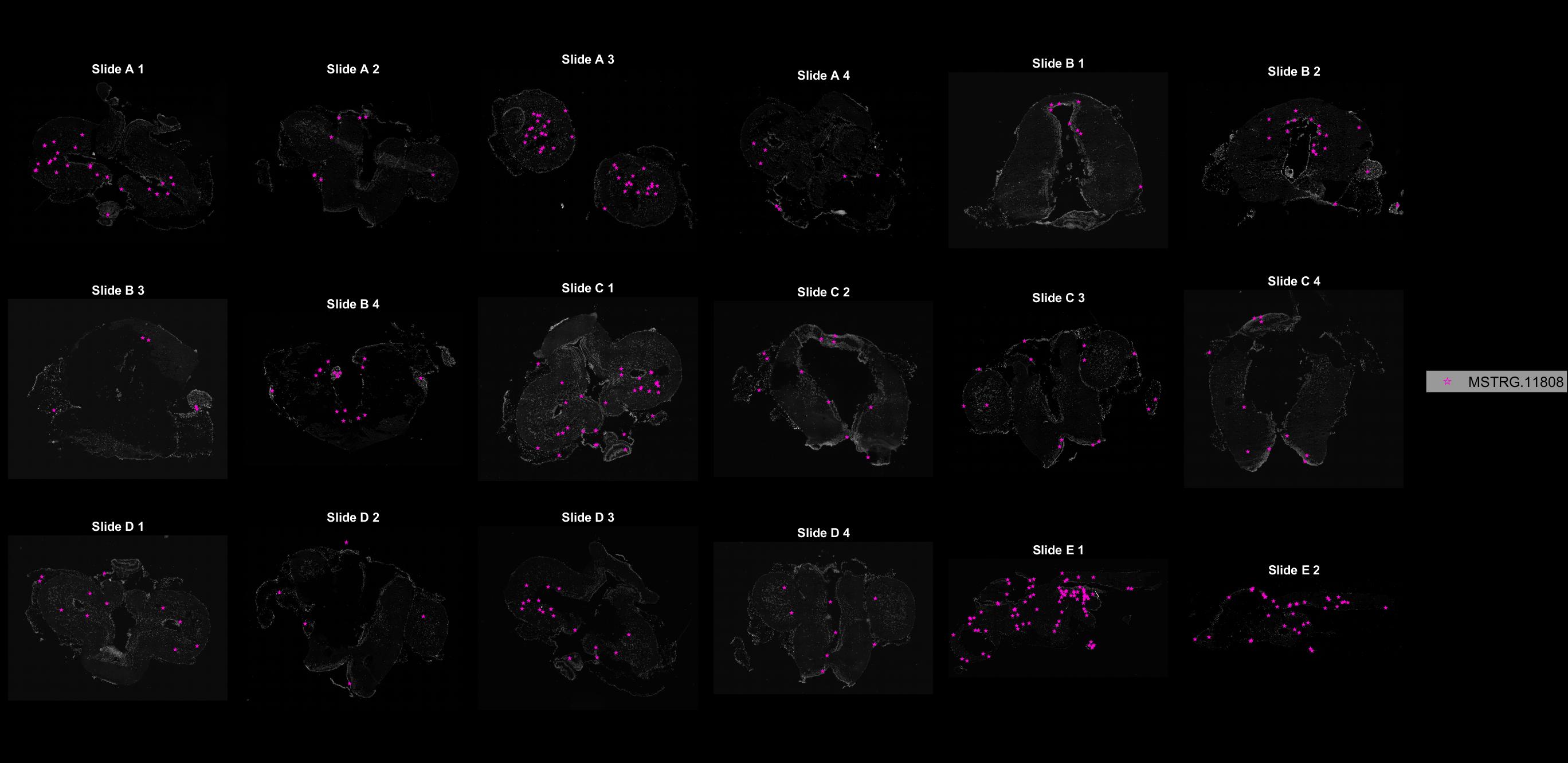

### MSTRG.11994.jpg

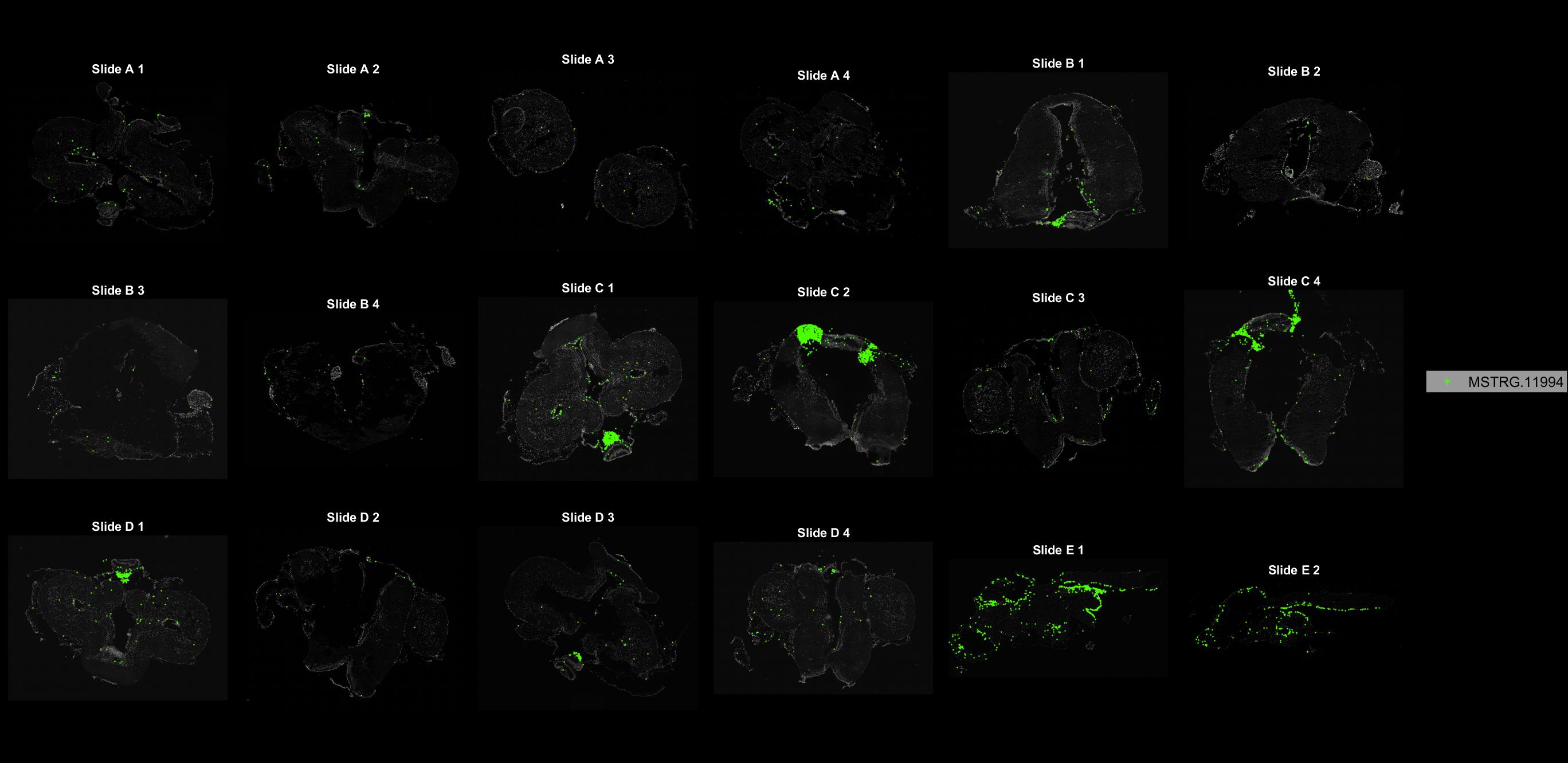

### MSTRG.12094.jpg

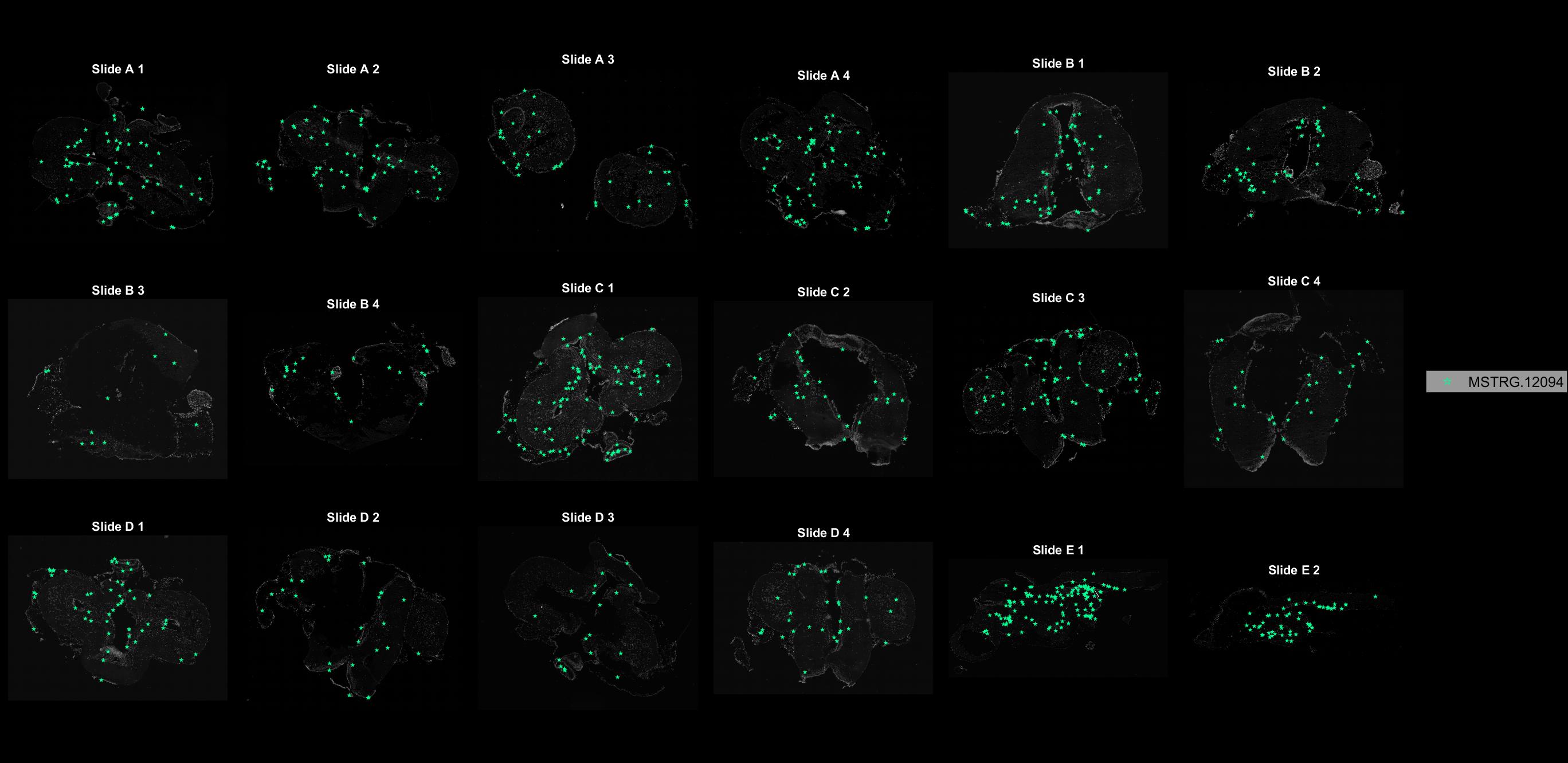

### MSTRG.12608.jpg

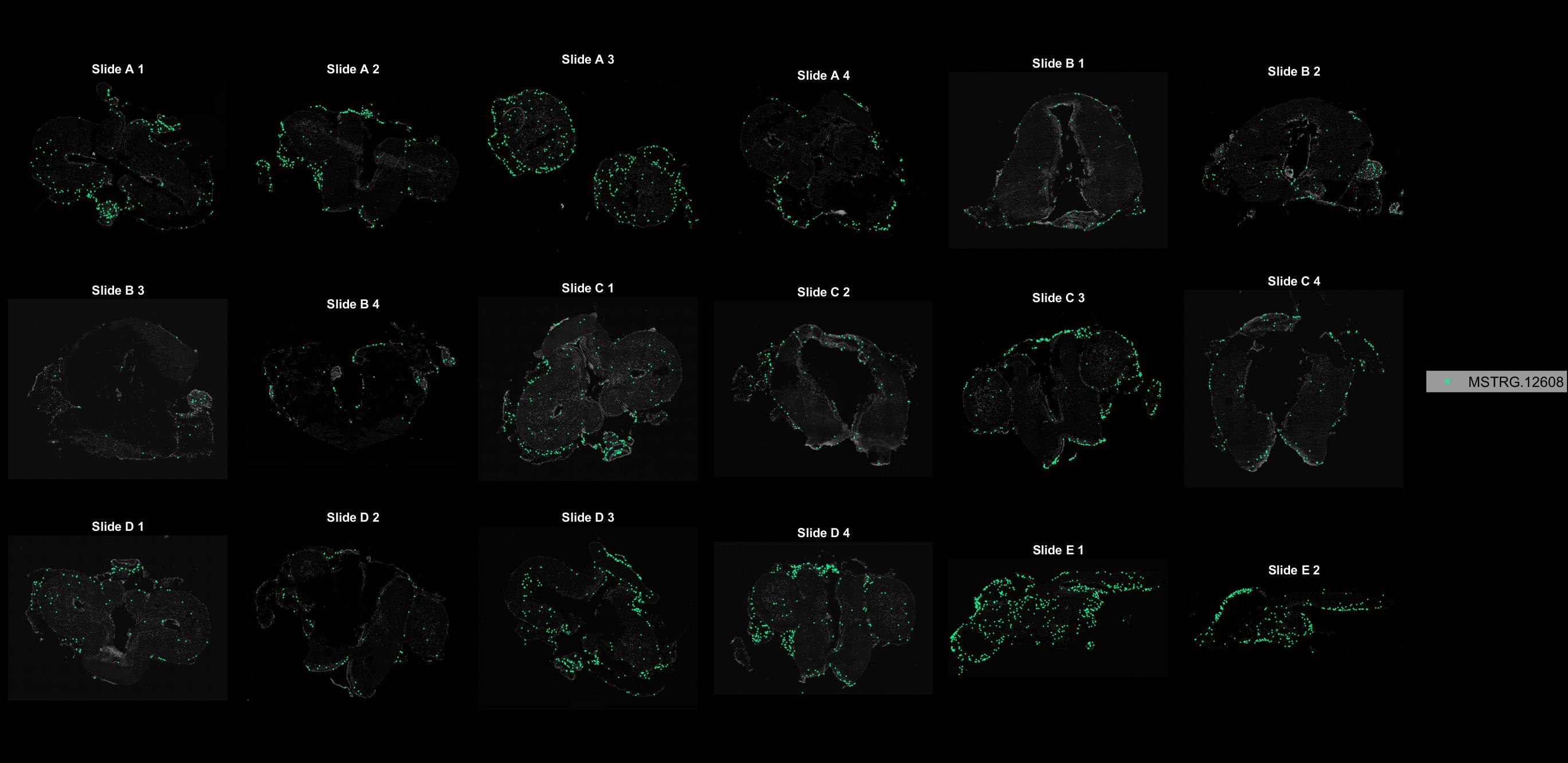

### MSTRG.13621.jpg

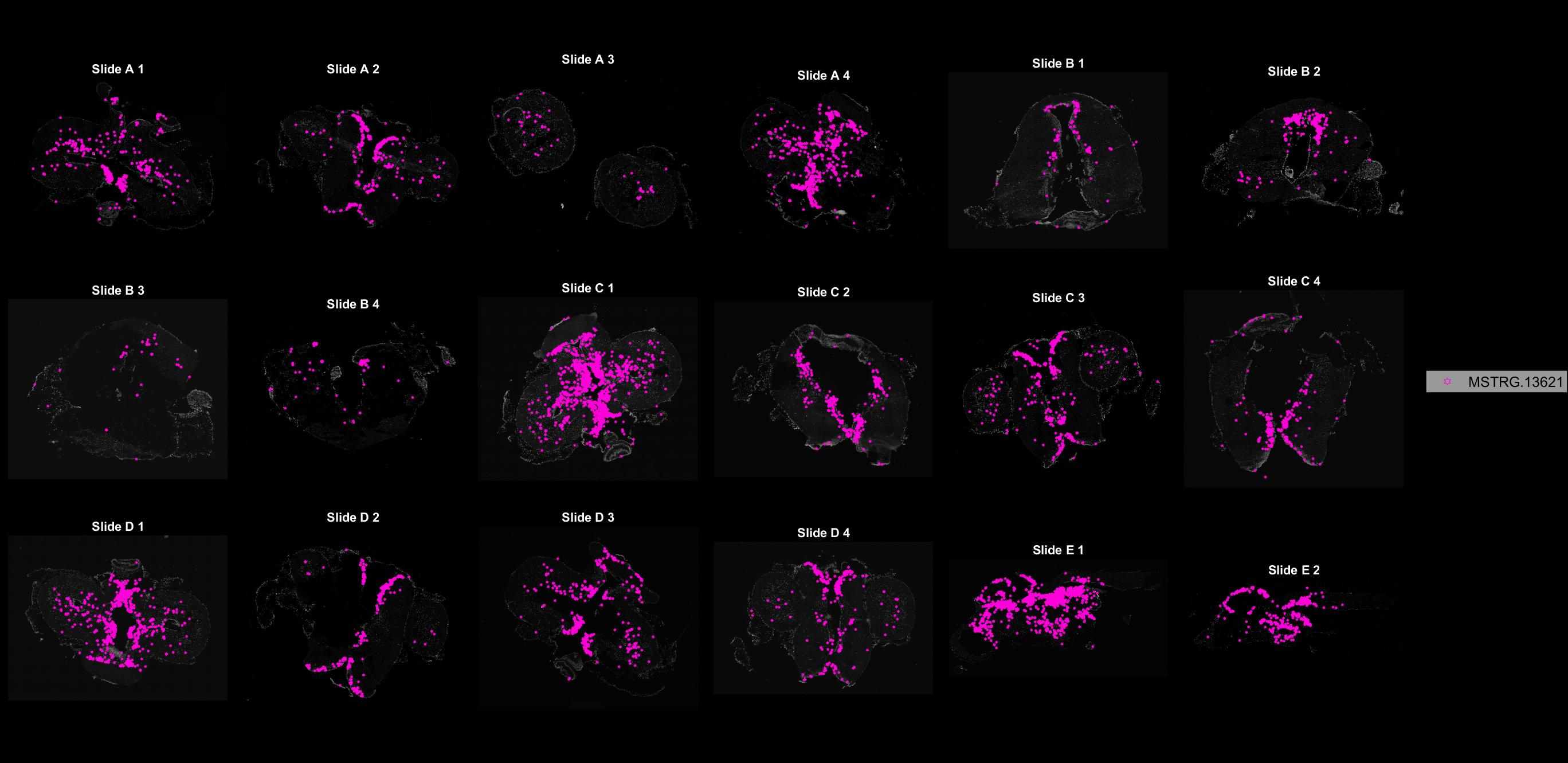

### MSTRG.13735.jpg

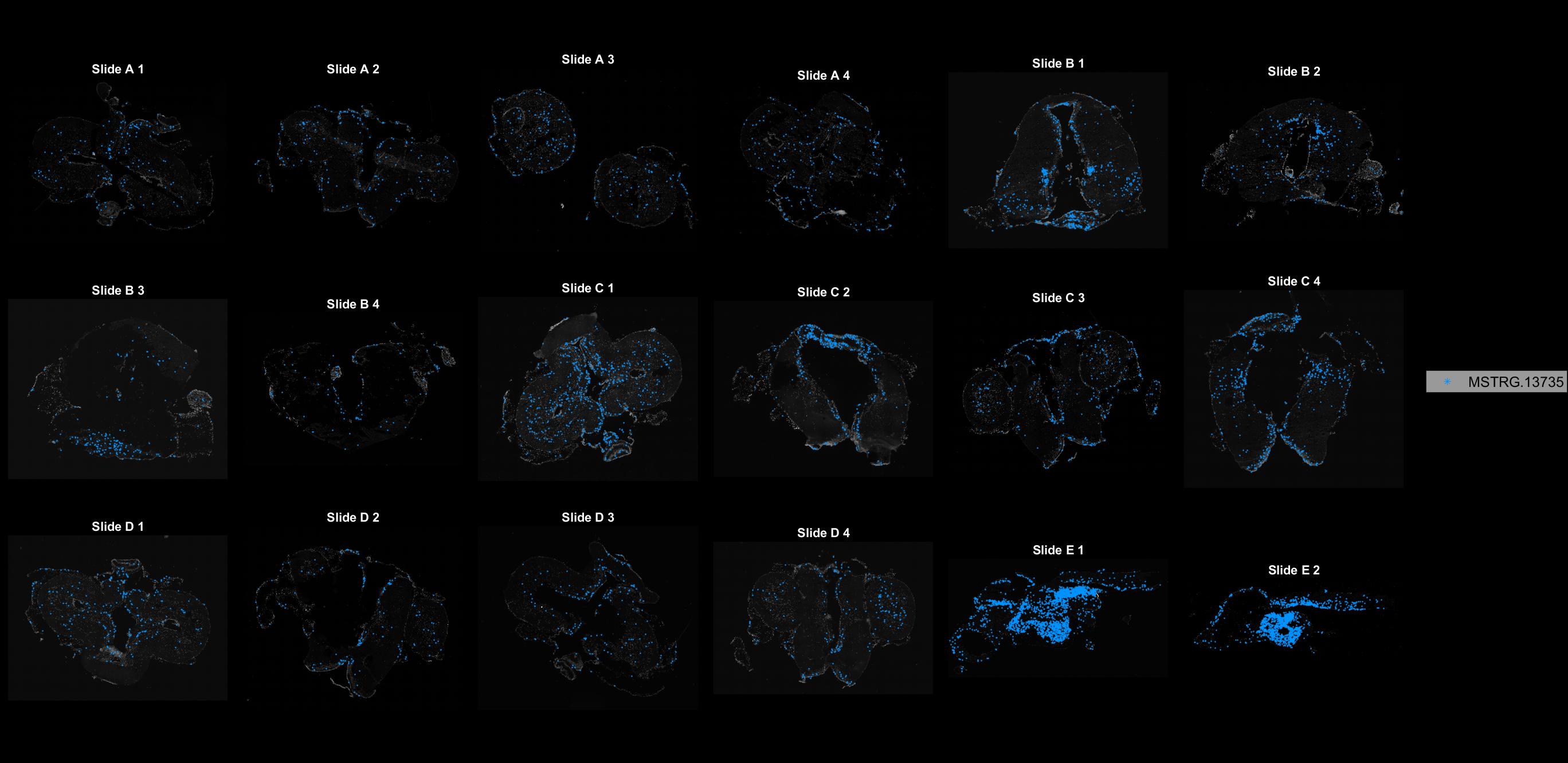

### MSTRG.14371.jpg

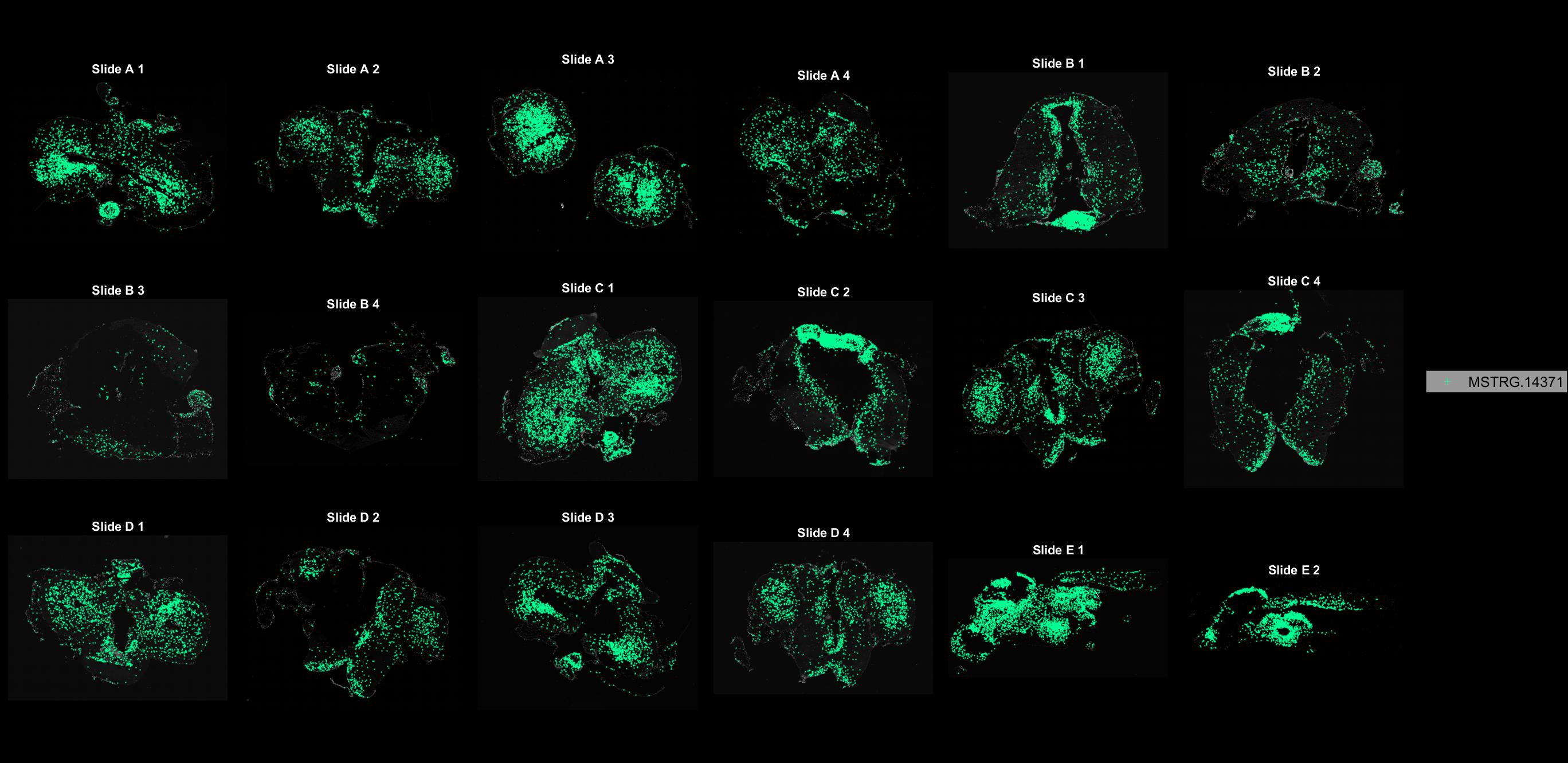

### MSTRG.14454.jpg

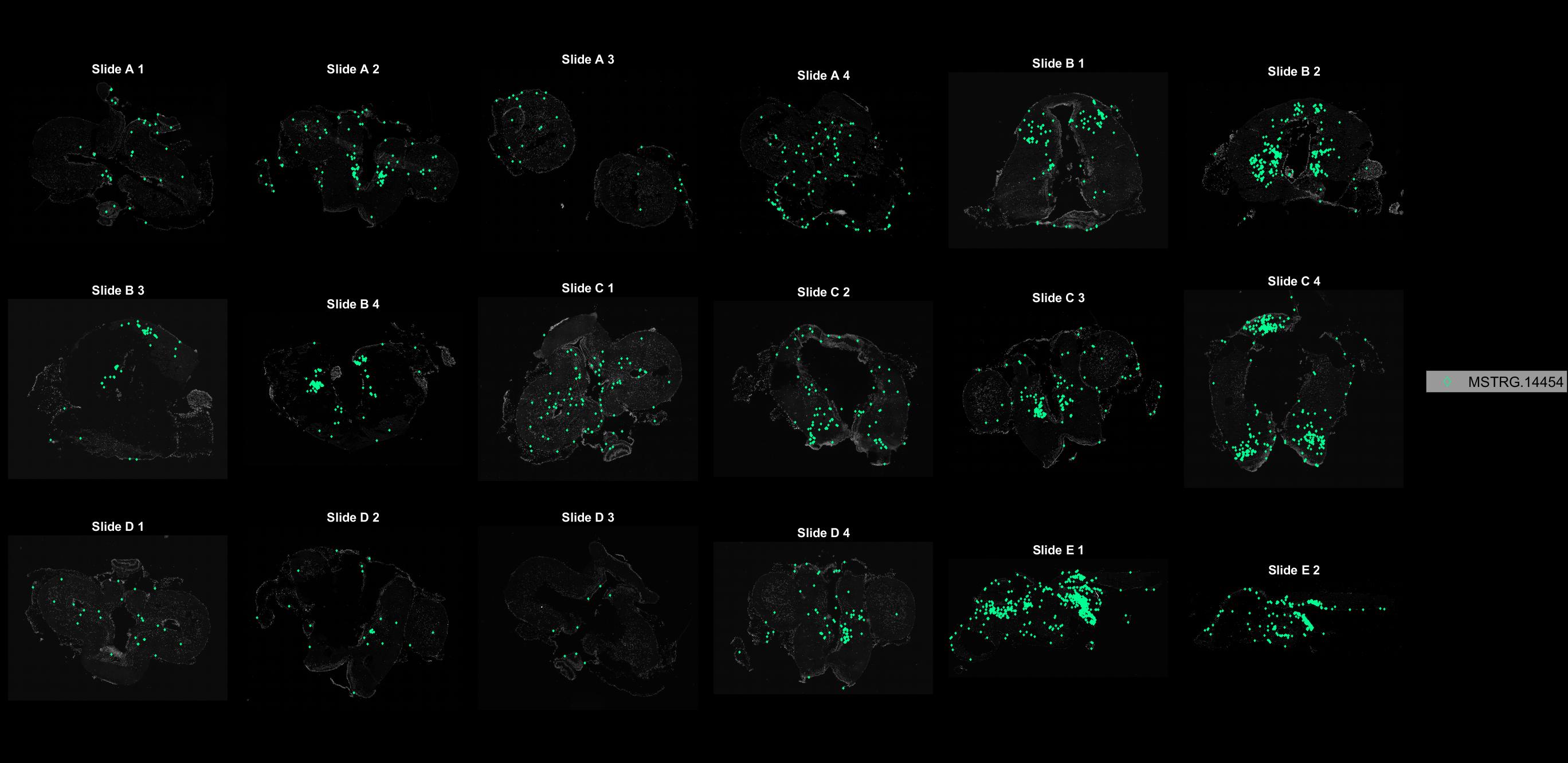

### MSTRG.16163.jpg

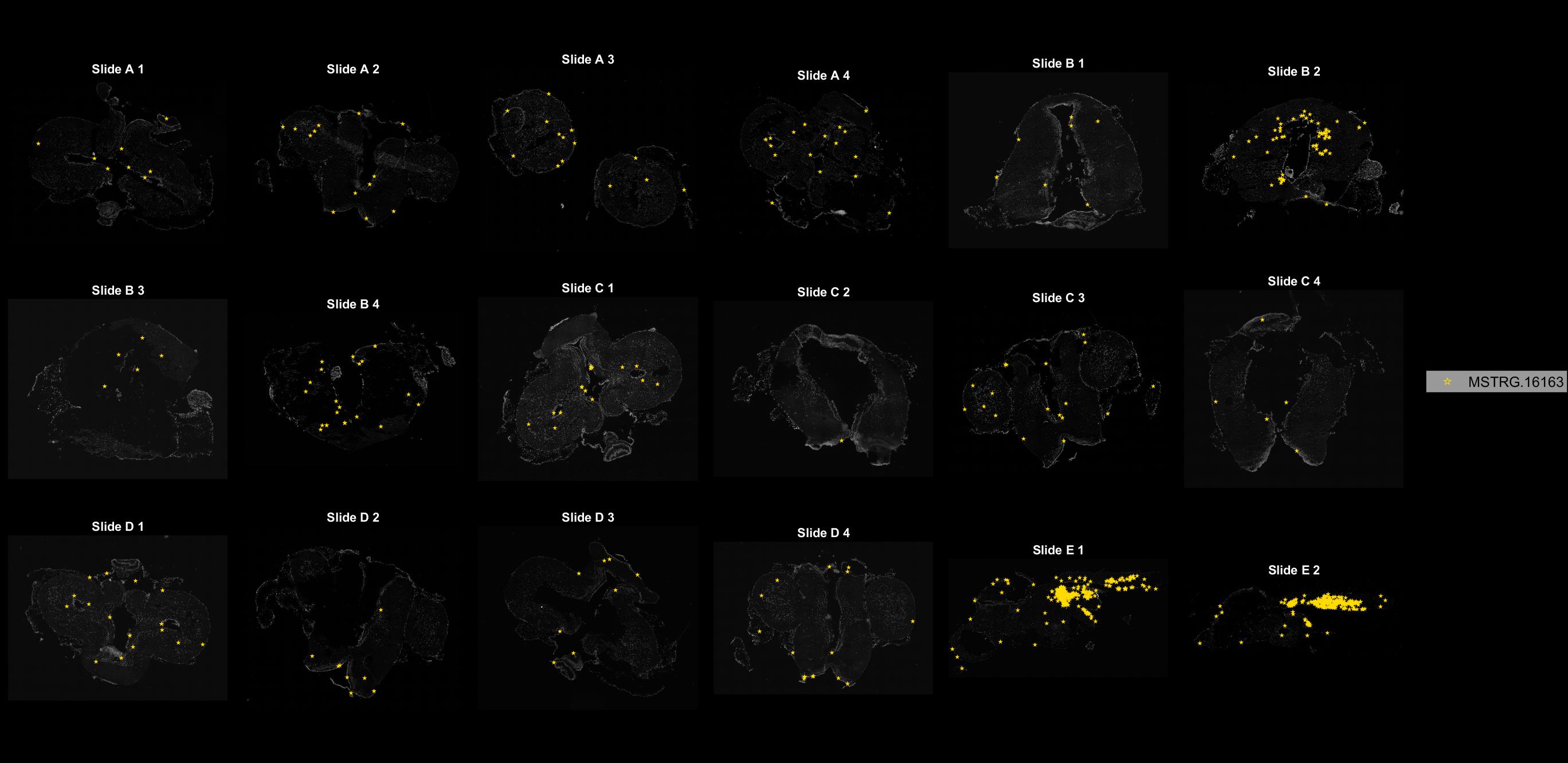

### MSTRG.16433.jpg

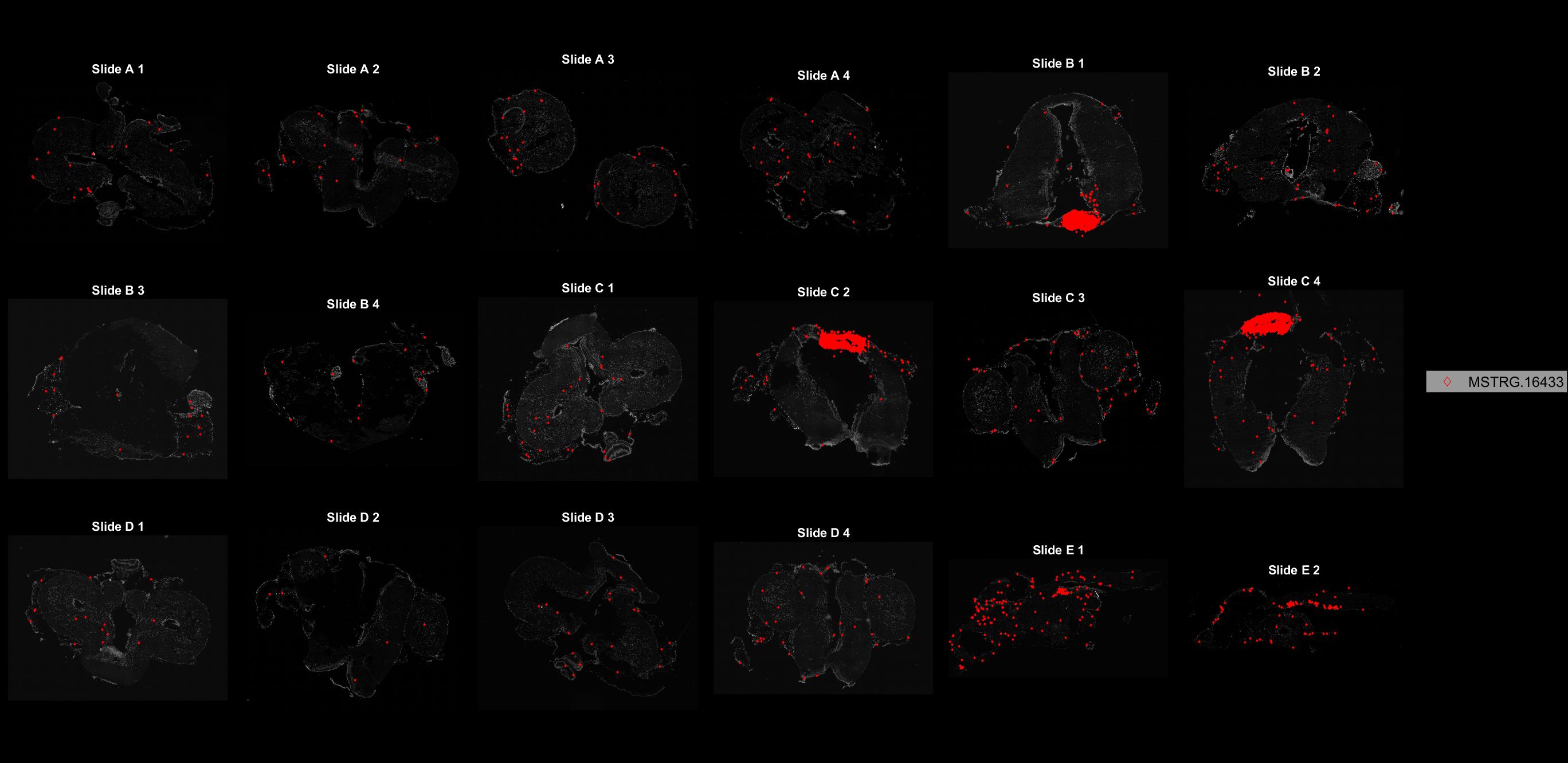

### MSTRG.16491.jpg

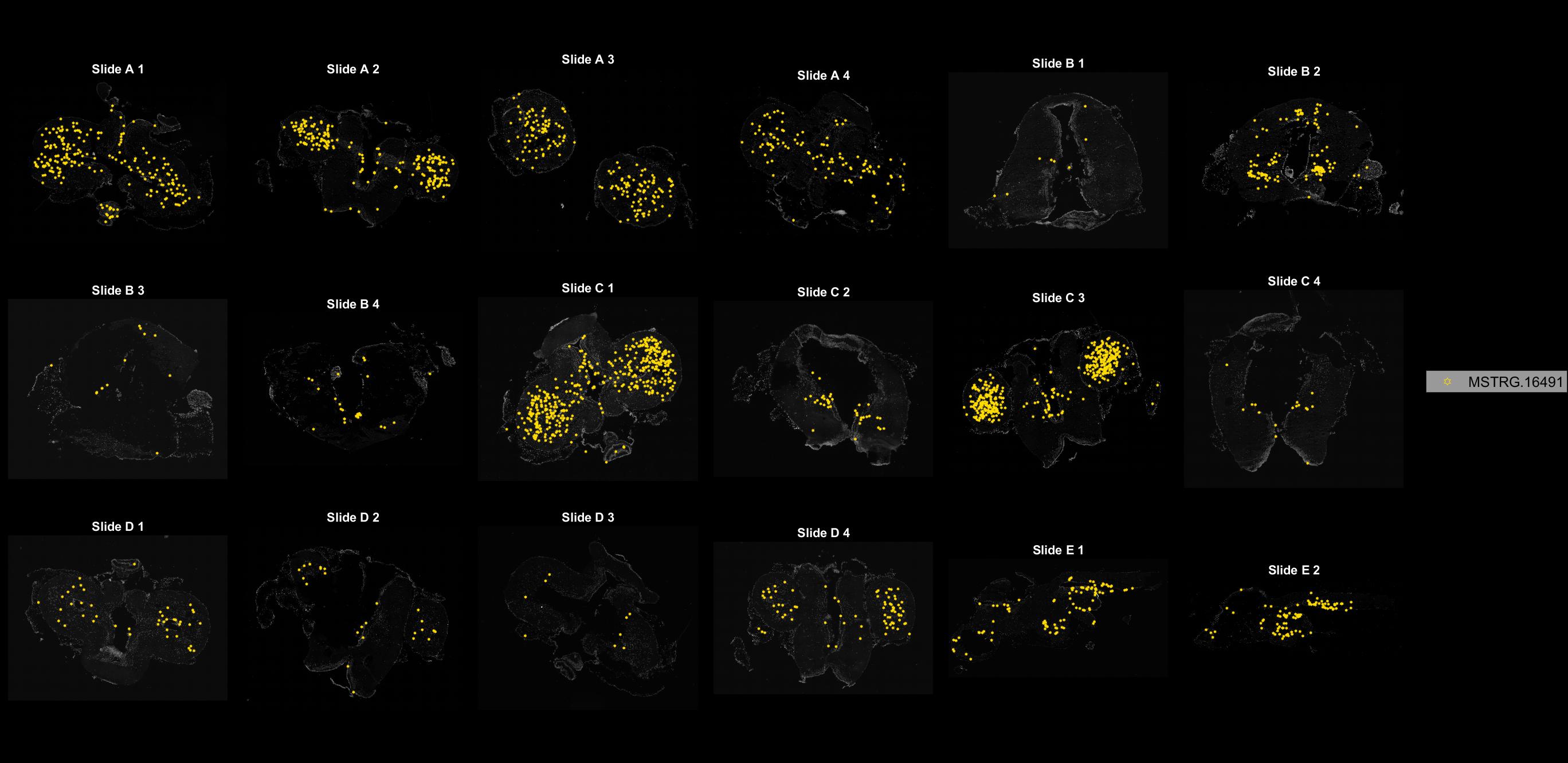

### MSTRG.17344.jpg

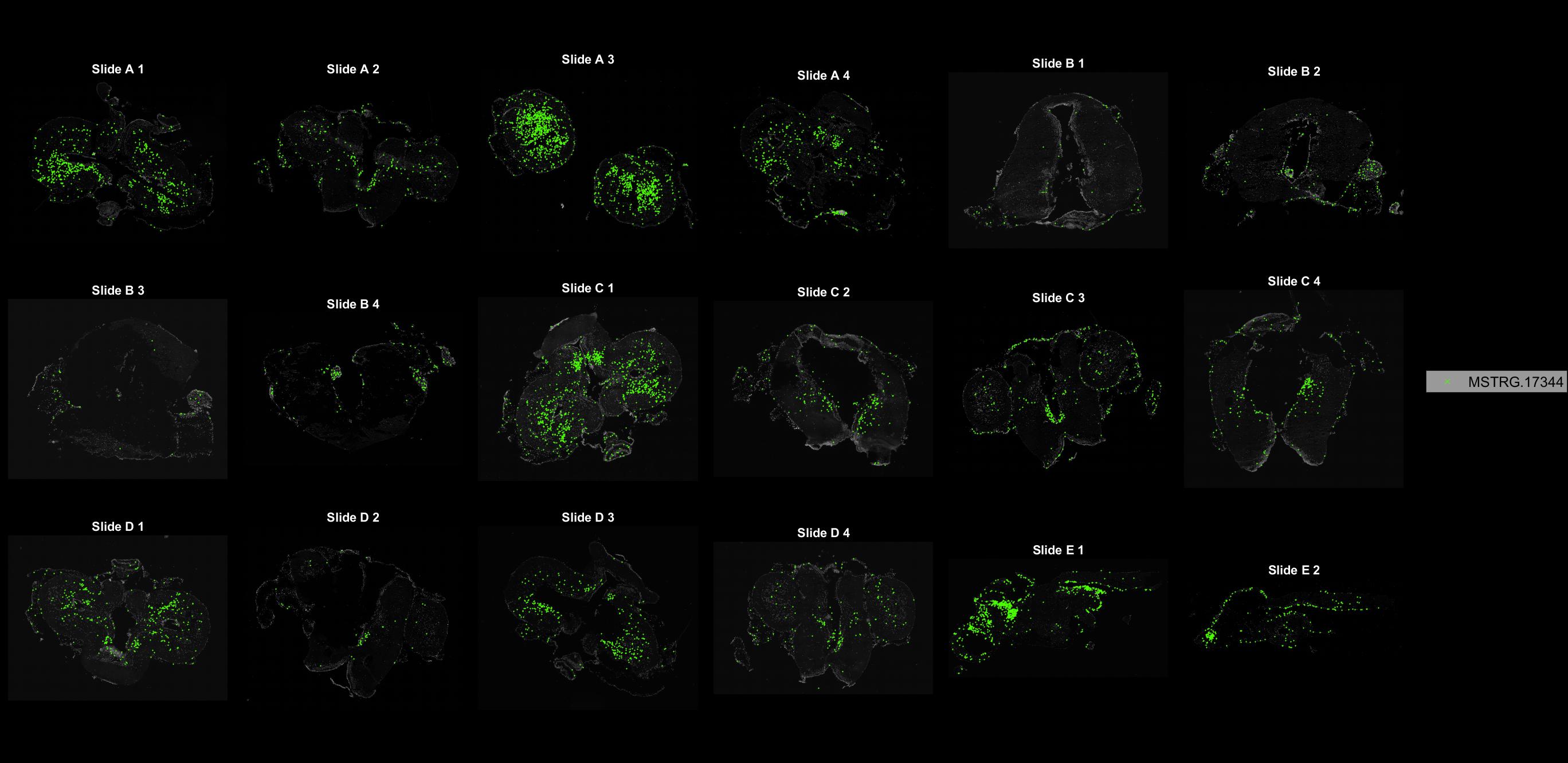

### MSTRG.17402.jpg

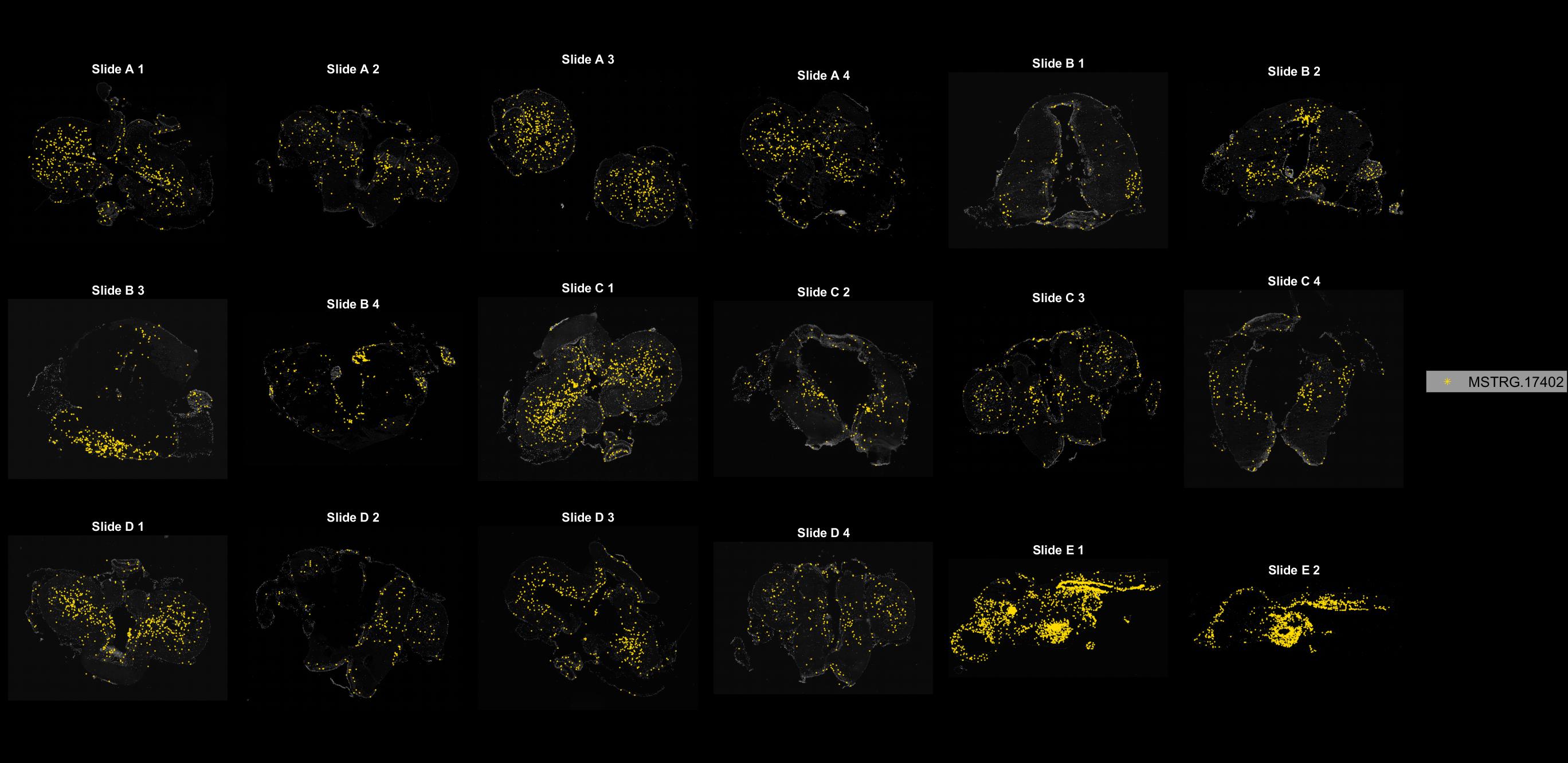

### MSTRG.17441.jpg

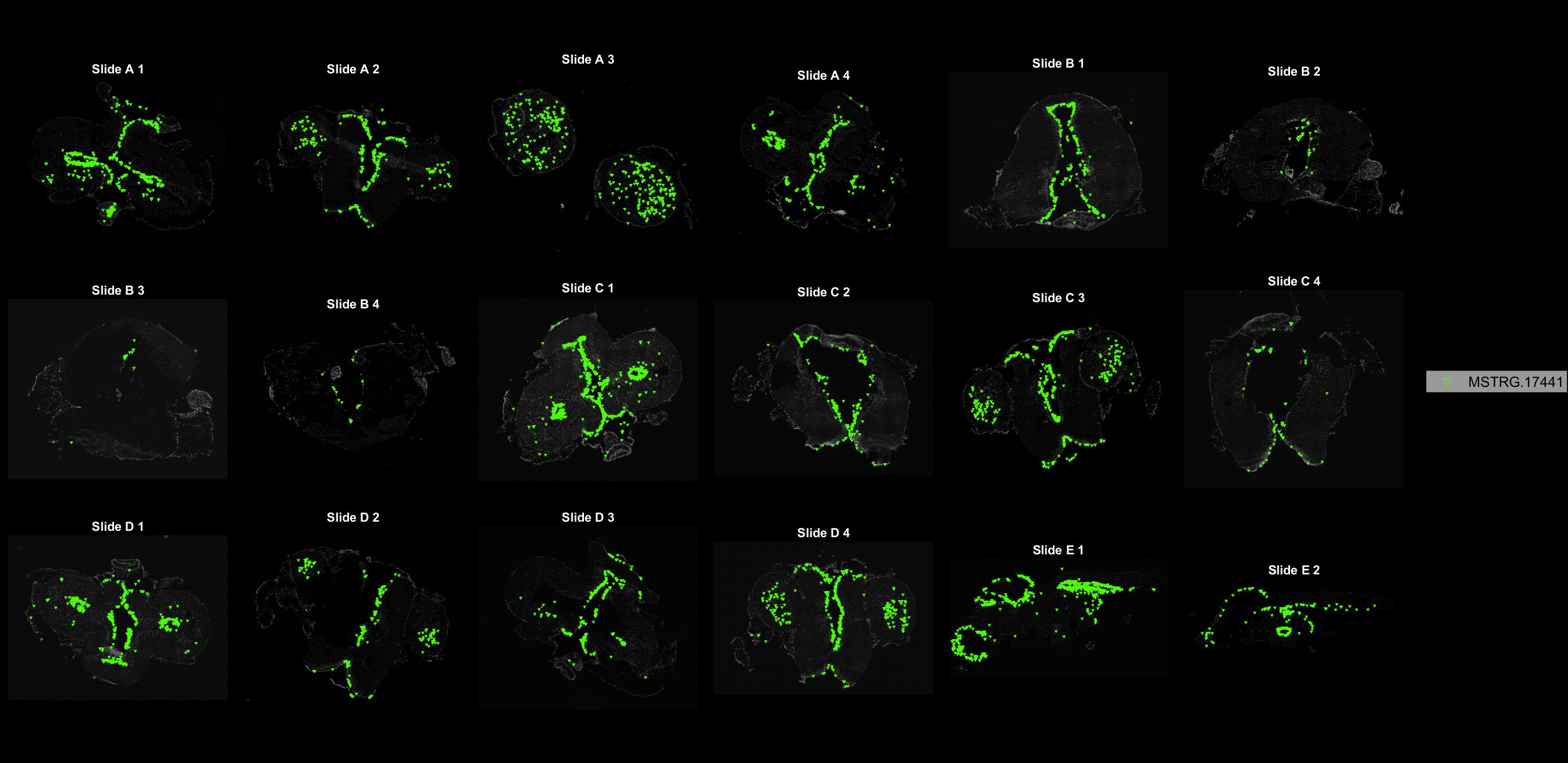

### MSTRG.18084.jpg

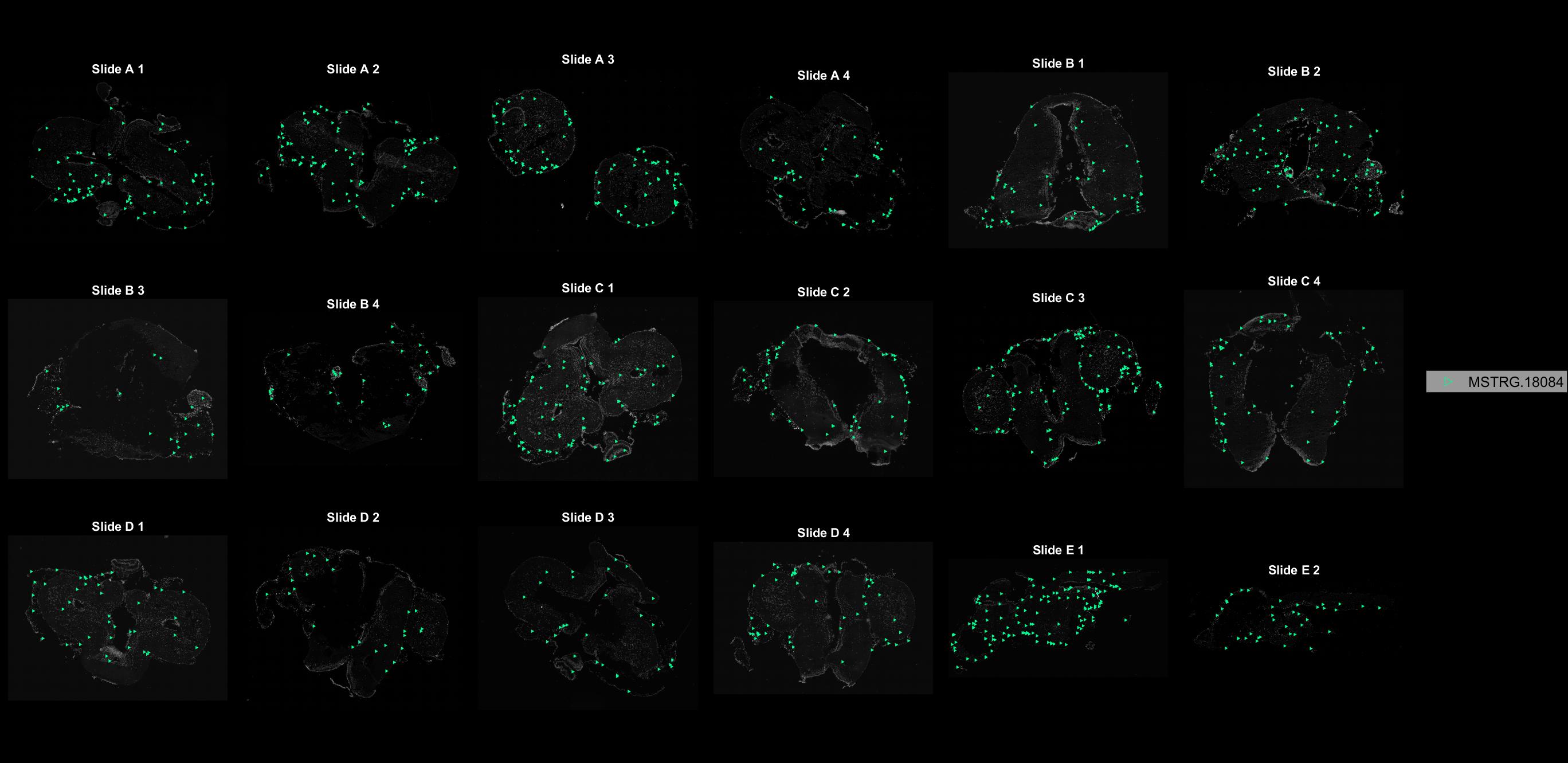

### MSTRG.18138.jpg

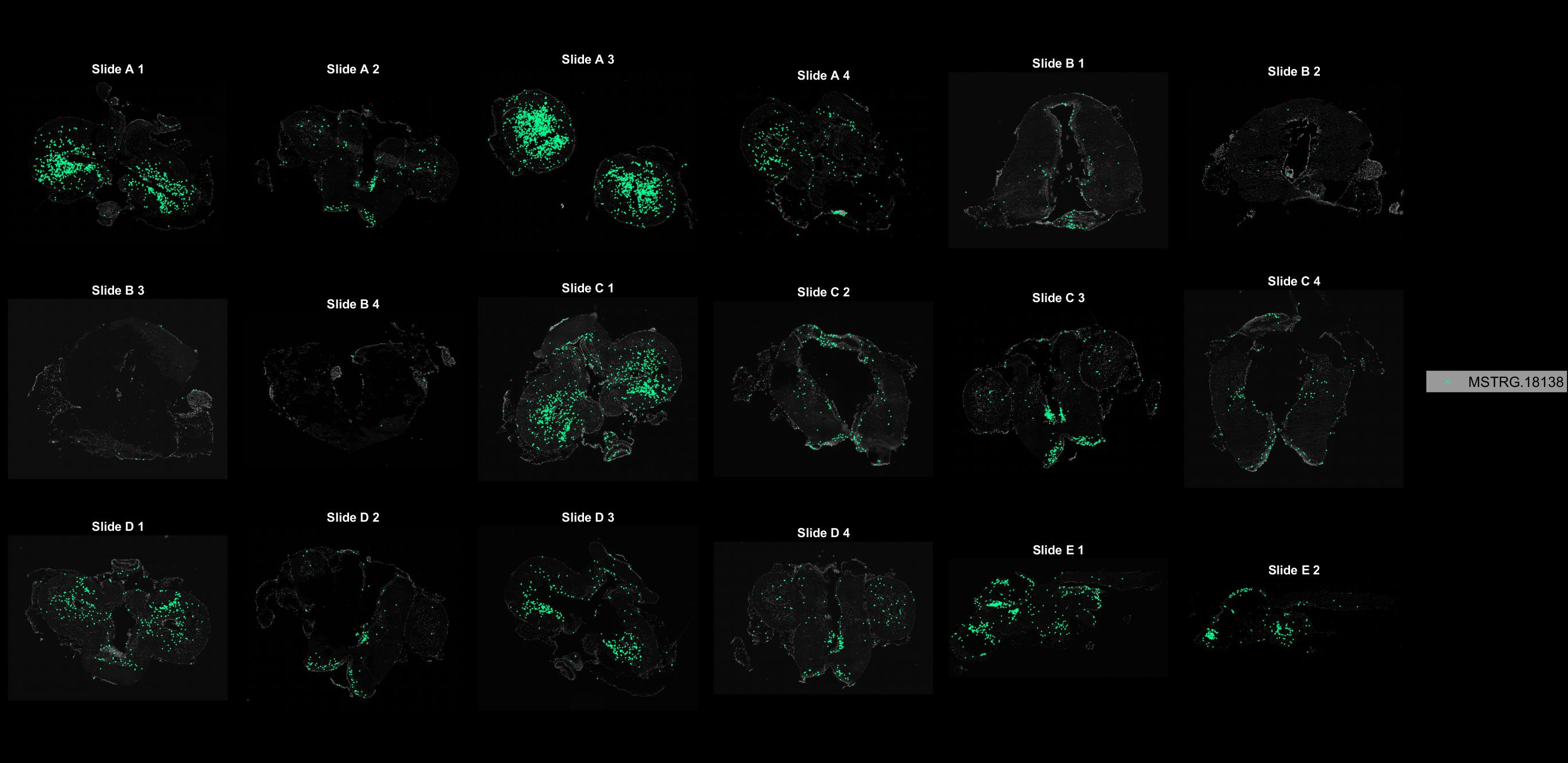

### MSTRG.19810.jpg

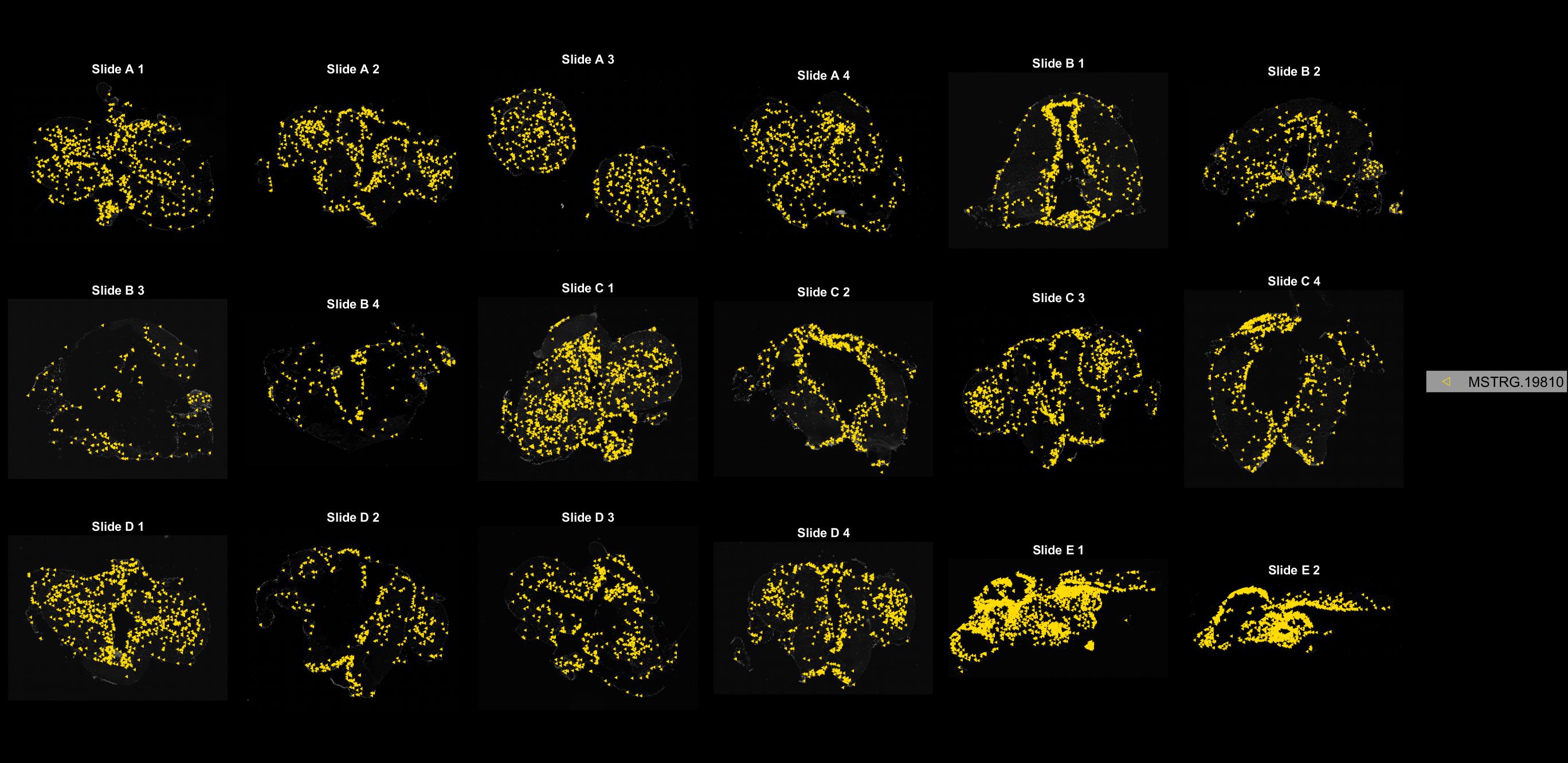

### MSTRG.20425.jpg

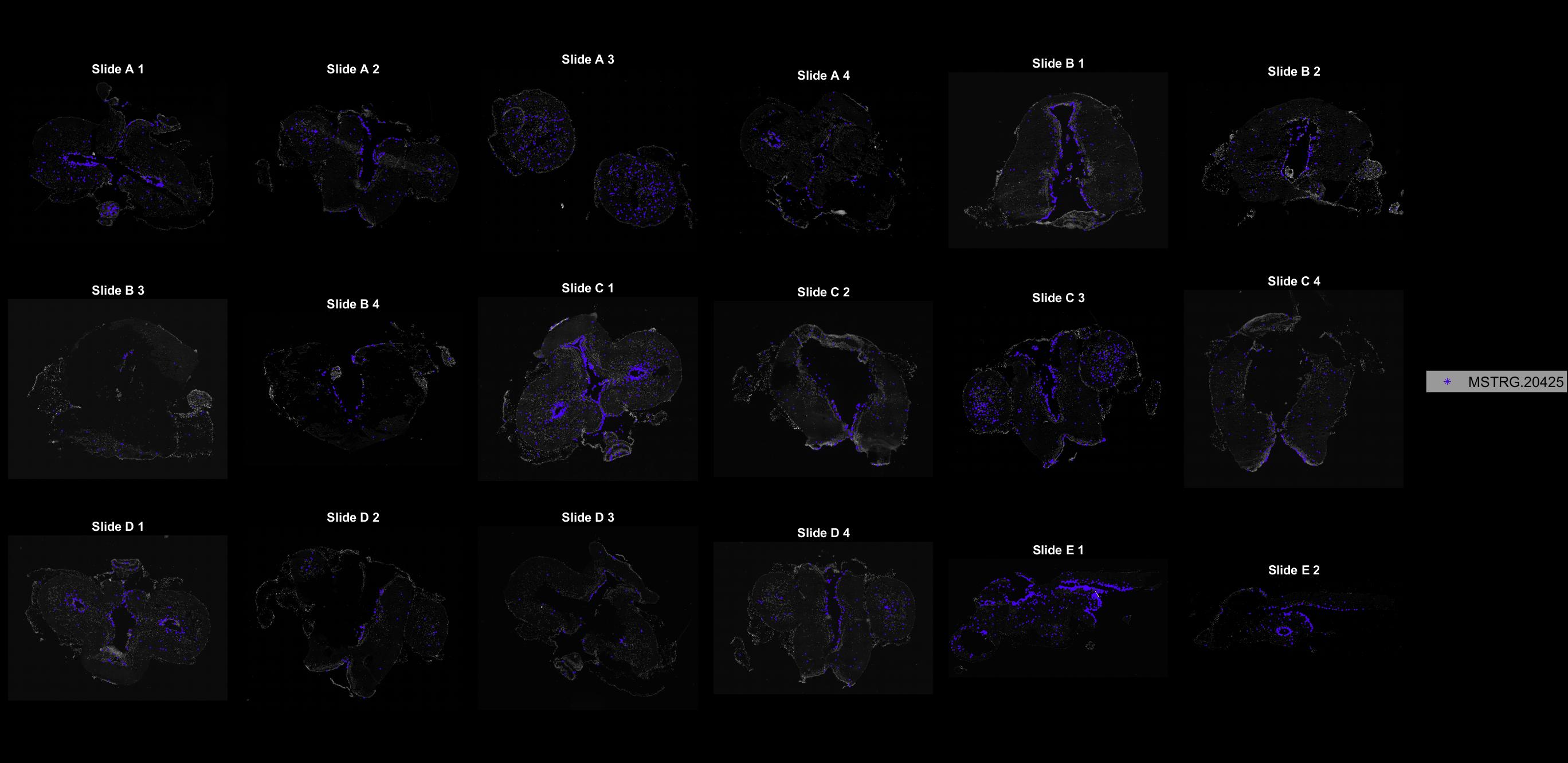

### MSTRG.23085.jpg

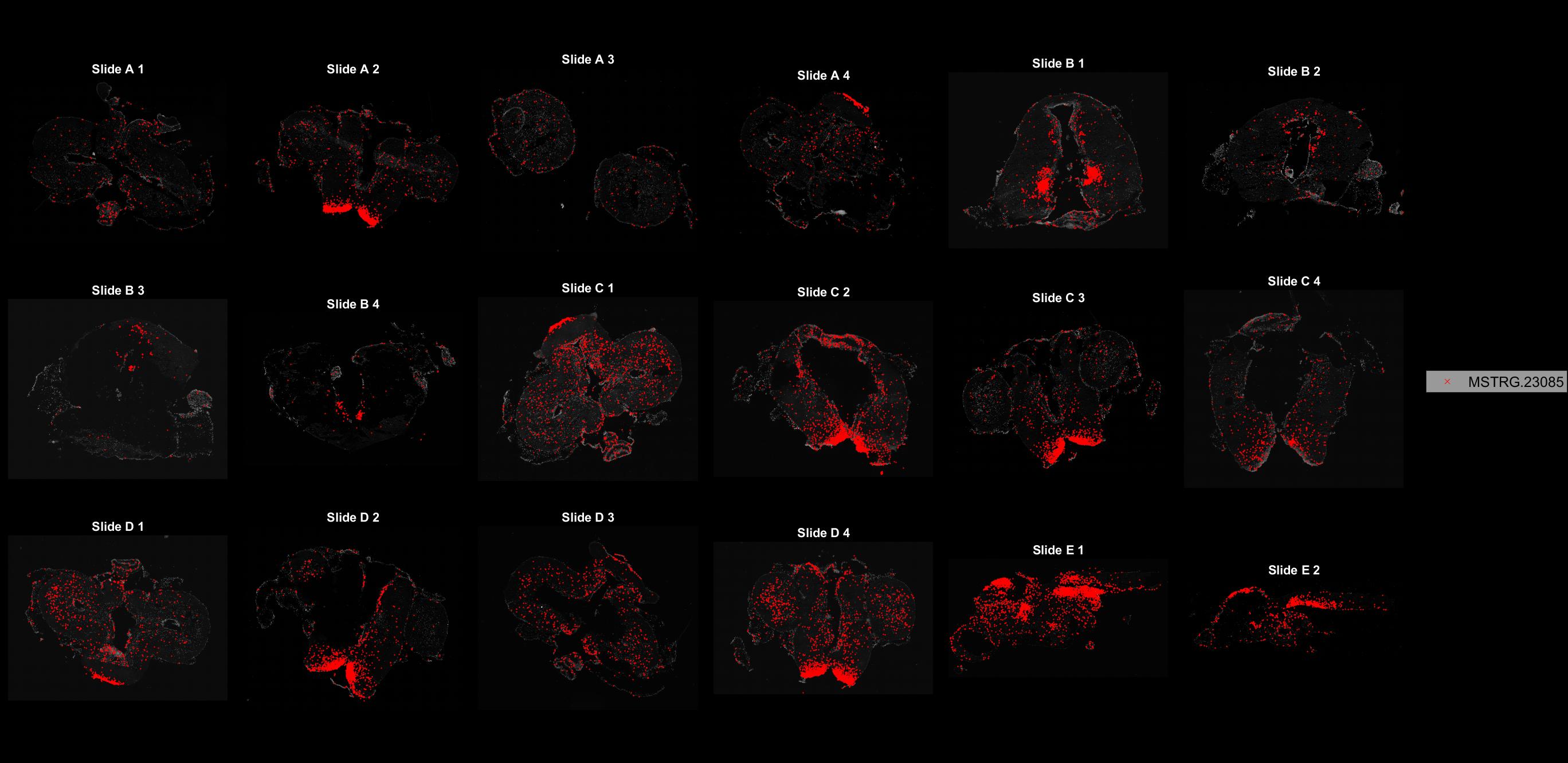
